# Exploring metal resistance genes and mechanisms in copper enriched metal ore metagenome

**DOI:** 10.1101/2020.07.02.184564

**Authors:** Esmaeil Forouzan, Ali Asghar Karkhane, Bagher Yakhchali

## Abstract

Heavy metal pollution is a major global health challenge. In order to develop bioremediation solution for decontamination of environment from heavy metals one appropriate step is to investigate heavy metal resistance strategies used by microbial communities in the metal contaminated environments. The aim of the present study was to understand detailed mechanisms by which long time heavy metal (HM) exposed microbial community use to cope with excess of HMs. We exploited the Illumina high throughput metagenomic approach to examine taxonomical and functional diversity of copper enriched soil metagenome. Three enriched metagenomes were compared against 94 metagenomes derived from non-contaminated soils. Taxonomic composition analysis showed that phylogenetic profile of metal contaminated soils were enriched with *γ-Proteobacteria*. Comparison of functional profile of the two group reveled significant difference with potential role in HM resistance (HMR). Enriched SEED categories were “Membrane Transport”, “Cell Wall and Capsule”, “Stress Response”, “Iron acquisition and metabolism” and “virulence and defense mechanisms”. Raw metagenomic reads were assembled into scaffolds and predicted Open Reading Frames (ORFs) were searched against metal resistance gene database (BacMet). Based on enriched genes and gene categories and search of known HMR genes we concluded the microbial community cope with HM using at least 10 different mechanisms. Copper resistance genes were more abundant in the metagenome relative to other metals and pumping metals out of the cell were more abundant relative to other HMR mechanism. Results of the present study could be very helpful in understanding of HMR mechanism used by microbial communities.

## Introduction

As a result of industrialization and changes in the environment, Heavy metal (HM) pollution is a major global health challenge, especially in developing countries ^1^ and situation is worsening steadily^2^. Electroplating, alloy manufacturing, smelting, fertilizers, fungicides, pigment, plastic, refining processes, battery and mining are some of anthropogenic sources for heavy metals ^3,4^.

Copper (Cu) has been known as an essential element but also a potent toxic heavy metal at high concentration. Cu could cause toxicity in the cell by replacing other metals used as a part of enzymes or other protein complexes, Reactive oxygen species (ROS) production, damage membrane integrity and denaturing of proteins^5,6^. Currently some approaches are available to decontaminate HM contaminated aqueous solution, such as ion exchange resins and chemical precipitation ^7^ however these treatments are just efficient at high concentration of metals and produce secondary contamination which require additional treatment ^8^. HM concentration in polluted water is often around 1 mg/l or even less and requires to be reduced to less than acceptable limit. According to World Health Organization (WHO) heavy metal concentration limits (in mg\L) in drinking water are 0.010 for Hg, As, and Pb, 3.0 for Zn and 0.003 for Cd^9^, This task, i.e. reaching this very low level of metal concentration is not trivial. Physicochemical procedure to remove HM from water are inefficient in this concentration and costly ^10^.

Unlike organic contaminants, which can be degraded into benign molecules, metals are not degradable in nature, hence remediation of all heavy metals are much more difficult than other organic pollutants. Therefore, detoxification of metal-polluted environments has to rely on their removal from contaminated sites. In order to develop bioremediation solution for decontamination of environment from heavy metals one appropriate step is to study HMR strategies used by microbial communities in the metal containing/contaminated environments. There are some studies on this subject^11–17^. One of main source of heavy metal contamination is metal ore mining and steel industry ^18^ so metal resistance genes have to be more prevalent in such metal contaminated sites relative to other less contaminated ones ^19^.

The recent development of high-throughput DNA sequencing technologies, combined with the development of bioinformatics and public databases, has drastically changed our perception of microbial communities to different stresses. NGS technology made functional and taxonomical profiling a routine task, which was very expensive, and time consuming before NGS ^20^. Thanks to this technology nowadays even small biological laboratories are able to launch metagenomic project and study environmental microbial community in great detail ^20^. NGS has been used as a powerful tool to explore microbial community response to different stresses including HM contamination ^12^. In a recent study on soil bacterial community structure in different iron mining area ^21^ 16s amplicon sequencing reveals that HM polluted area had significant higher bacterial alpha diversity than those from unpolluted area and contaminated area have significantly different bacterial composition relative to non-contaminated sites. Some other studies show different strategies used by different microbial communities in response to HM contamination ^12^, including HM precipitation by sulfides production, secretion of extracellular phosphatases and metallophores, change in cell wall and capsule structure, modification of lipopolysaccharides, periplasmic copper oxidation, modification of transporters, methylation/volatilization, ROS detoxification and cytoplasmic metal accumulation ^12^. As HM bioremediation have to include metal removal from environment, among these strategies, metal accumulation and HM precipitation have higher potential as a bioremediation method.

In the present study, we analyzed and compared Cu enriched soil metagenome to metagenomes derived from non-contaminated soils and reveal the variations of the functional and taxonomical between these two groups. Our findings could be useful in better understanding of how microbial community cope with HM contamination and pave the way to reach a bioremediation method to HM pollution.

## Methods

### Sample collection and DNA extraction

Three samples (S1, S2, and S3) were gathered from top surface soil (5-15cm) from Hmyard iron ore, Semnan province, Iran on 2013-07-15. S1, S2, and S3 were sampled from different part of Hmayard iron ore, an idle soil, a sample from near a tree grown in the ore and a sample from near an annual plant grown in the ore respectively. For each sample, approximately 500 g of soil gathered and kept on ice in sterile plastic bags until reaching to the NIGEB lab. After returning to the lab, 200 g of each sample soil stored at −80 C for subsequent molecular analysis. Immediately the remaining soil were sieved through a 100-mesh to remove stones and visible plant fragments and after enrichment DNA extraction was done according to Lee and Hallam^22^. The remaining was air-dried at room temperature for one week and stored at 4 °C for further chemical analysis.

### Soil physicochemical analysis

In order to measure the soil HMs content we closely followed the methods reported by ^23^. Based on Akcay and coworkers, a fraction that contains exchangeable HM species is obtained by the following procedure: each sample is extracted at room temperature for an hour with 10 ml of magnesium chloride solution (1M MgCl2 buffered at pH 7.0) with continuous shaking. HM concentration in the supernatant was measured after centrifugation at 3000g using atomic absorption spectrophotometry (Shimadzu AA670/G).

### Copper enrichment

An enrichment process was used as described by Li et al ^24^. It was assumed that copper resistant microbial fraction of the community could be enriched under high copper concentration. Briefly two grams of fresh soil sample was mixed with 5 ml sodium phosphate buffer (0.1 M) and 50 μl of the suspension spreaded over three LB agar plates containing copper (200, 400, 800, 1,000, 1500, 2000 mg L^−1^). The LB agar plates were incubated at room for 72h. The results showed that on the 1,000 mg L^−1^ copper concentration 9 colonies were found and just 2 on 1500 mg L^−1^. According to this observation, 1,000 mg L^−1^ was chosen as final concentration in enrichment process. For enrichment 5 gr of soil was added to 5 ml of enrichment solution (containing of sucrose and tryptose and starch, each 200 mg L^−1^) supplemented with 100 mg L^−1^ copper under sterile condition and incubated for 2 days room temperature and again enrichment solution was added. This cycle were repeated for 10 times in such a way that after the last cycle (20th day) the final concentration of copper in the soil was 1,000 mg L^−1^.

### Library construction and metagenomic sequencing

After DNA extraction, quantity and quality of extracted DNA were measured using spectrophotometry (NanoDrop spectrophotometer) and agarose gel electrophoresis (1%). DNA extraction was done in triplicate and pooled to minimize DNA extraction bias. Subsequently, DNA was sheared into fragments (^~^400 bp) using ultrasonication. Paired-end library were built using the TruSeq2 Kit according to the manufacturer’s instructions (http://www.illumina.com/). Resulting libraries were sequenced using Illumina HiSeq 2000 (Pair-end sequencing (2 * 100 bp) according to standard protocol (http://www.illumina.com/).

### Bioinformatics analysis

The adaptors was removed and raw reads were trimmed based on two criteria, quality score ≥20 and length ≥50 bp using trimmomatic ^25^. Clean paired reads were merged using FLASH ^26^ and submitted to mg-rast sever for further analysis ^27^. Reads data and the analysis results are available with MG-RAST project number mgp83855. Merged reads were analyzed using MG-RAST pipeline with default parameters. Clean reads were assembled using metaSPAdes software ^28^. MetaSPAdes was chosen because previous studies showed its good performance relative to other assemblers ^29,30^. Assembly was done with multiple k-mer approach (k-mers = 21,31,41,51 and 61). Genes were predicted in the resulting contigs using meta Gene Mark ^31^.

For comparative part of the study a search was done for all shotgun soil metagenomes in MG-RAST ^27^ database. The resulted metagenomes were manually checked and a collection of 94 metagenomes from non-contaminated soil was obtained. Functional and taxonomical profile for all soil metagenomes were retrieved using MG-RAST API ^32^ and used as control group in comparative analysis. Properties of metagenomes used in the study are available in Table S-DatasetStat. Comparative metagenomics statistical test was done using STAMP ^33^. For comparison, white’s non-parametric t-test was used with Benjamini-Hochberg as multiple test error correction.

For HM resistance gene analysis, metal resistance genes database were downloaded from BacMet database ^34^. All of the predicted protein were searched against the database using blastp. Blast results output were filtered based on three criteria, Evalue ≤E-5, coverage ≥ 90% and identity ≥ 70. Best hit for each query was accounted for downstream analysis. Frequency of resistant genes for each HM metal were calculated using self-written python script.

## Results

Previous studies demonstrated Hmyard iron ore contains high level of copper and could be a source of metal contamination in the region (unpublished data). Metal concentration of iron ore soil for Fe, Cu, Cr and Cd were 919, 256, 11 and 0.5 mg/kg respectively. This shows copper and iron are in high concentration relative to other studied soils ^35,36^ and could cause health problem to organisms living in there. Therefore we hypothesized microbial community of such community have to be resistant to HMs (at least to iron and copper).

### Quality control

Clean paired reads were merged using flash^26^. Flash output was 24 million merged reads with average length of 160 bp. The merged reads were submitted to MG-RAST server for further analysis. Based on the DRISEE ^37^ submitted reads error content was ^~^1% and ambiguous character content was 0.003. 28% of reads failed to pass MG-RAST QC criteria, however there were still enough reads to evaluate taxonomical and functional profile^38^. Detailed statistics are available in table S-DatasetStat. Dataset and resulted assembly properties are briefly overviewed in Table 1.

**Table 1:**
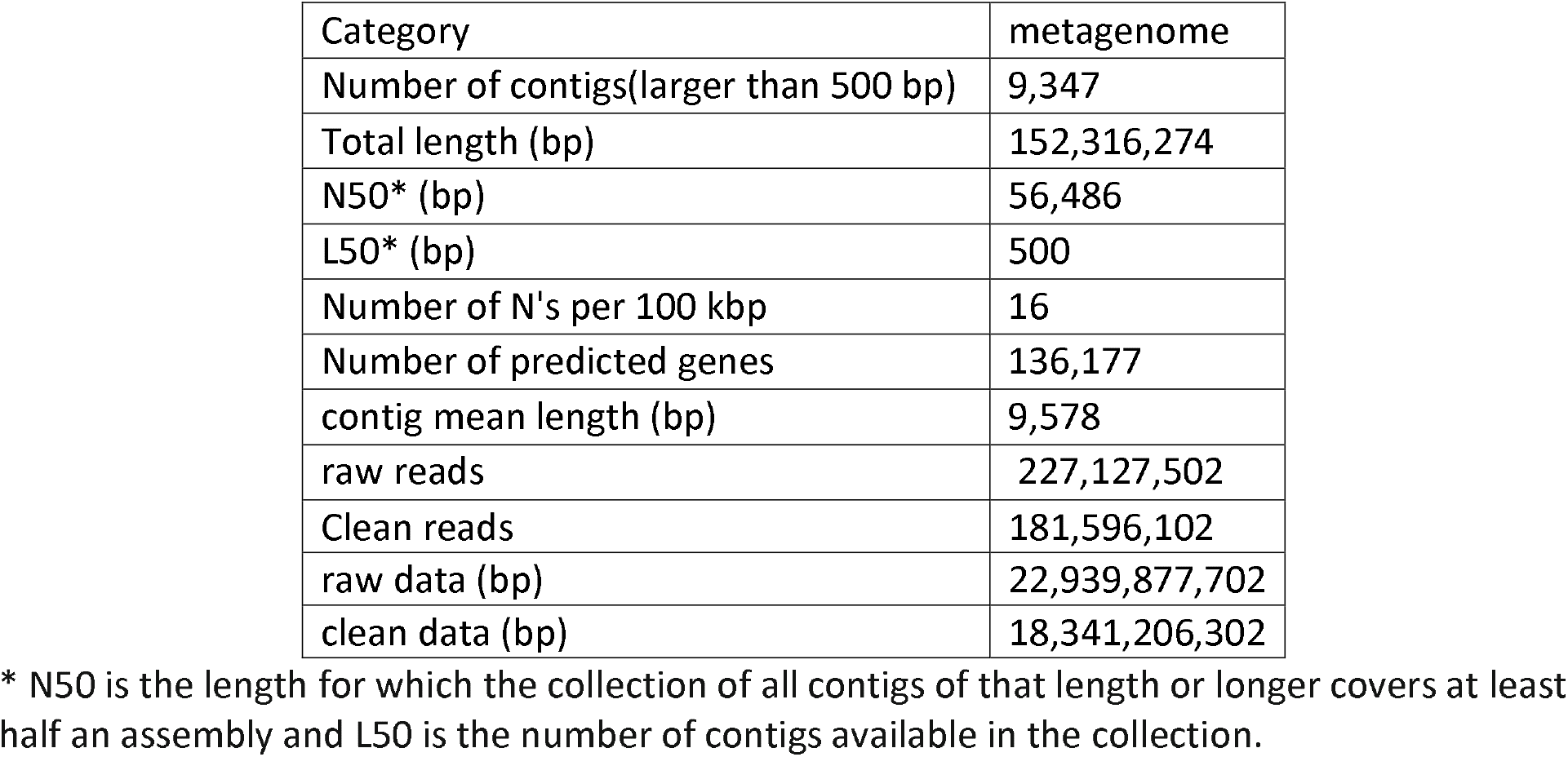
Overview of dataset and resulted assembly

### Taxonomic composition

Shanon diversity index for the enriched metagenome (EM) was 31, which is 16 fold smaller than that for average of control soils (Table S-DatasetStat), indicating that enrichment severely reduce population diversity. Rarefraction curve becomes almost flat to the right meaning that sequencing depth was enough to detect majority of the diversity available in the sample (figure S-Rarefraction). At domain level, EM contained 98% Bacteria, 1.8% Eukaryota, 0.01% Viruses and negligible Archea. This means EM was enriched in bacteria and depleted in Eukaryota and Archea (figure S-domain). At phylum level, there are 13 taxa available in the EM with 0.01% or more frequency. *Proteobacteria* and *Firmicutes* dominated in EM and comprised 90% of population (figure 1). Comparative analysis showed that EM is highly enriched in *Proteobacteria* and depleted in *actinobacteria*. The *Proteobacteria* embraced mainly *γ-Proteobacteria* and Firmicutes mainly embraced bacilli. Compared to other non-contaminated environment *γ-Proteobacteria* are noticeably more prevalent in EM, in contrast some taxa such as *α-Proteobacteria, β-Proteobacteria, Actinobacteria* and *Acidobacteria* are depleted (Table S-class). At the genus level, 15 genus obtained with relative abundance greater than 0.01% of the total community microorganisms. The 5 primary genera of the EM, namely *Serratia, Bacillus, Staphylococcus, Arthrobacter* and *Pseudomonas* consist more than 75% of total population with *Serratia* alone consisting of more than 50% of the community (figure 1).

**Figure 1:**
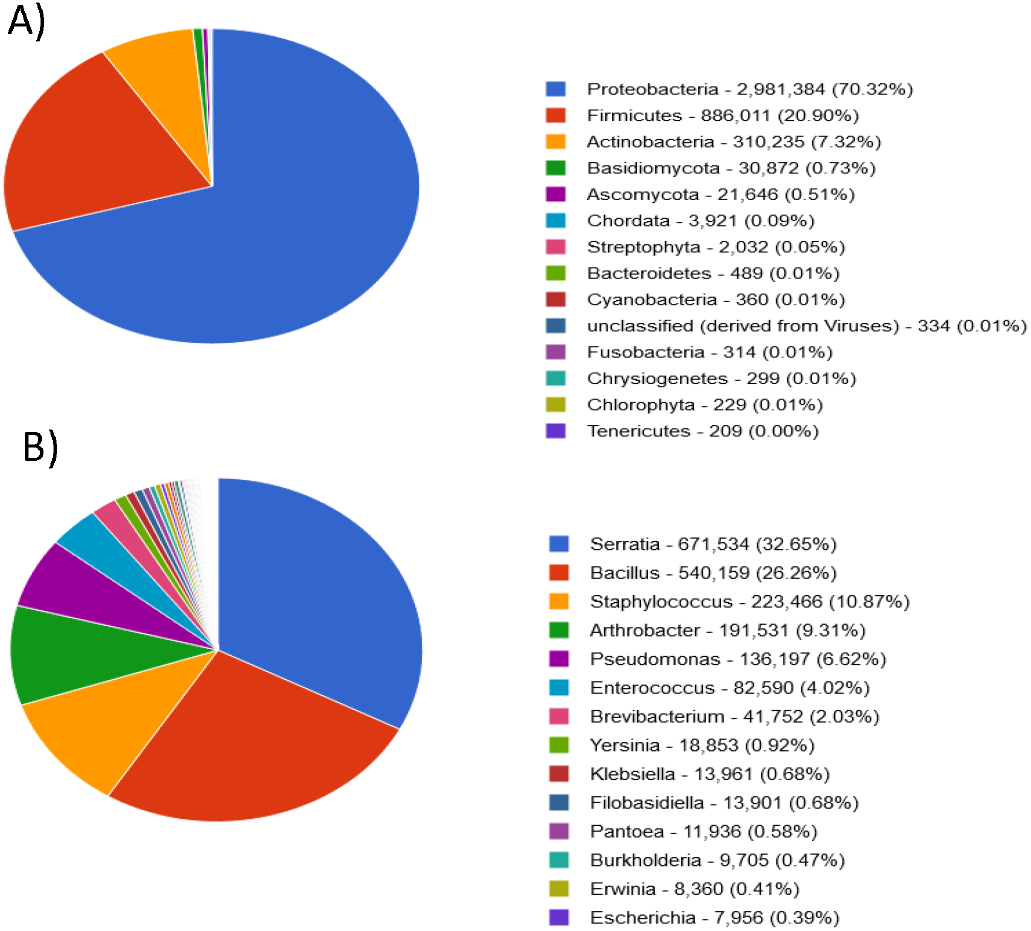
Taxonomic composition of copper enriched metagenome at phylum (A) and genus (B) level.

### Functional composition

From the QC passed reads 86% were annotated as protein features, 10% as unknown protein and 1% annotated as rRNA features. Figure 2 summarizes functional profile for level 1 of the SEED subsystems. According to it, genes related to “Clustering-based subsystems” are the most abundant gene category in EM comprising 14% of total genes. “Carbohydrates” is the second prevailing gene category following by “Amino Acids and Derivatives”. Photosynthesis were the least frequent category in the EM followed by “Secondary Metabolism”. However, mentioning mere frequency of gene categories does not make biological sense of the community. In order to reveal enriched subsystems relevant to a particular environment comparative study is required. Comparing to metagenomic profile derived from non-contaminated soil sample reveals interesting difference that points to mechanism underneath of HMR. We use available metagenomes derived from many (94 metagenomes, MG-RAST metagenome IDs are available in Table S-mgmID) different uncontaminated soils as control. Comparative study revealed that genes related to “iron acquisition and metabolism” were most enriched category in the EM relative to control metagenomes (Figure 3). “Membrane Transport”, “Cell Wall and Capsule”, “phosphorus metabolism”, “potassium metabolism” and “Virulence Disease and Defense” are among enriched subsystems. “Protein Metabolism”, “Cofactors, Vitamins, Prosthetic Groups, Pigments”, and “Respiration” are among depleted subsystems in EM. Majority of the enriched categories are associated to known HMR processes. Comparative study for level 2 and 3 of SEED subsystem revealed difference in more details. Membrane transport encompass 9 subcategories at level 2 of the subsystems hierarchy including “Sym/Antiporters” and “ABC transporters”, both of which are enriched in the EM. From “Iron acquisition and metabolism” group mainly genes related to siderophore production and secretion were enriched at level 2 (Table S-level2 and figure S-level2). “Quorum sensing and biofilm formation” is another enriched level 2 category with demonstrated role in HMR. Osmotic stress, periplasmic Stress, Detoxification, Desiccation stress were enriched level 2 sub-categories of level 1 “Stress Response” main category. “Resistance to antibiotics and toxic compounds” which is also enriched in EM have many subgroups related to HMR, such as Copper homeostasis, Mercury resistance operon and multidrug resistance efflux pumps. As expected, many of subcategories bellow “Resistance to antibiotics and toxic compounds” are enriched in the EM but ironically two group which we expected to be enriched were among the depleted categories, namely “Cobalt-zinc-cadmium resistance” and “Zinc resistance”. “Sugar alcohols” and “Proline and 4-hydroxyproline” are two enriched group with known function in abiotic stress. Isoprenoid/cell wall biosynthesis is another enriched category with known function in heat and oxidative stress ^39^(Table-S-level2).

**Figure 2:**
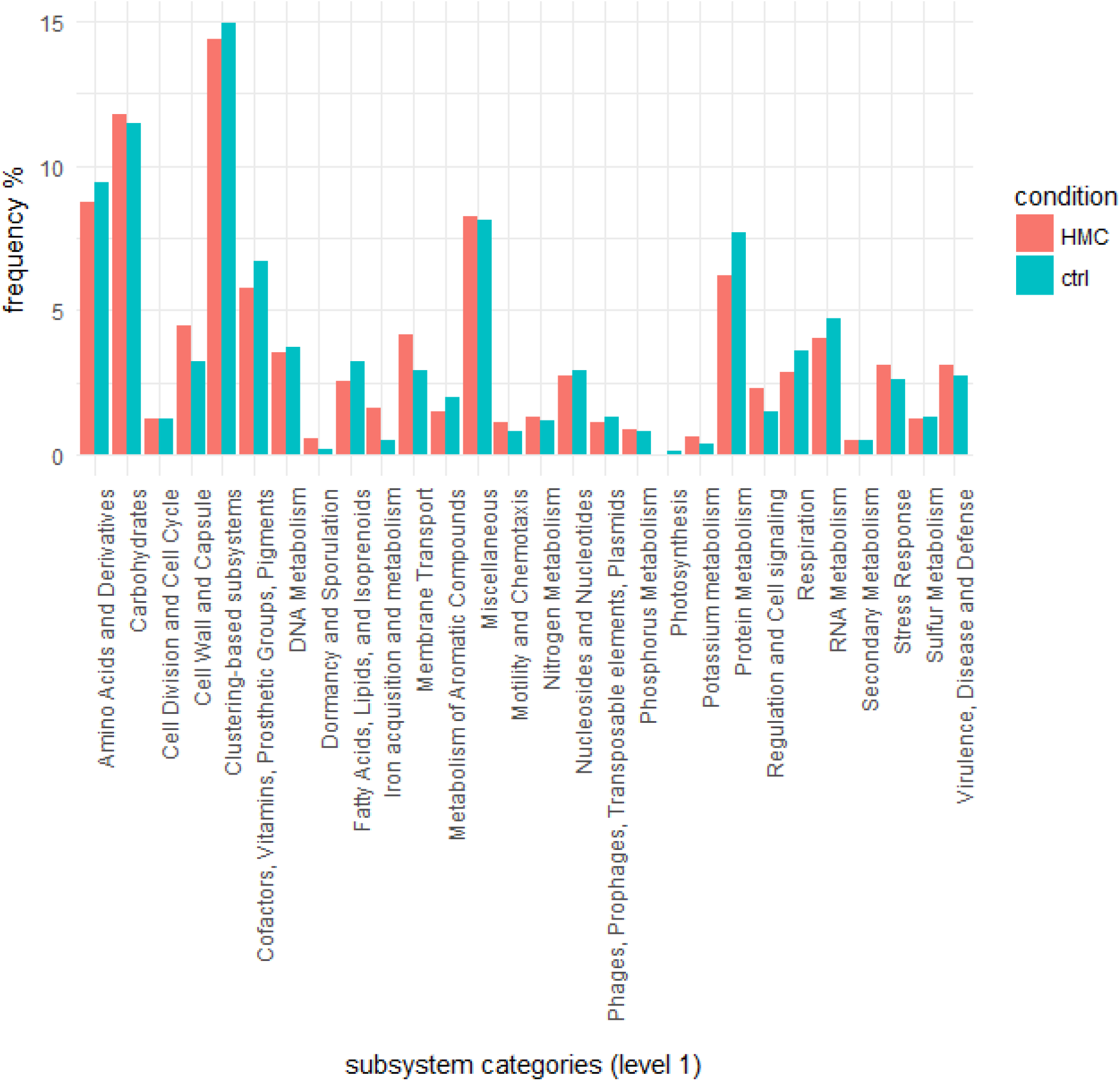
SEED subsystem (level1) distribution differences between the HMC (heavy metal contaminated) and control (HM uncontaminated) soil metagenomes based on MG-RAST annotation. The cutoff parameters for read annotation were default of MG-RAST (identity ≥ 60% and e-value ≤ 1×10-5). Comparison for level2, 3 and function level are available as supporting information (Figures and Tables names with prefix S-level2, S-level3 and S-Function).

**Figure 3:**
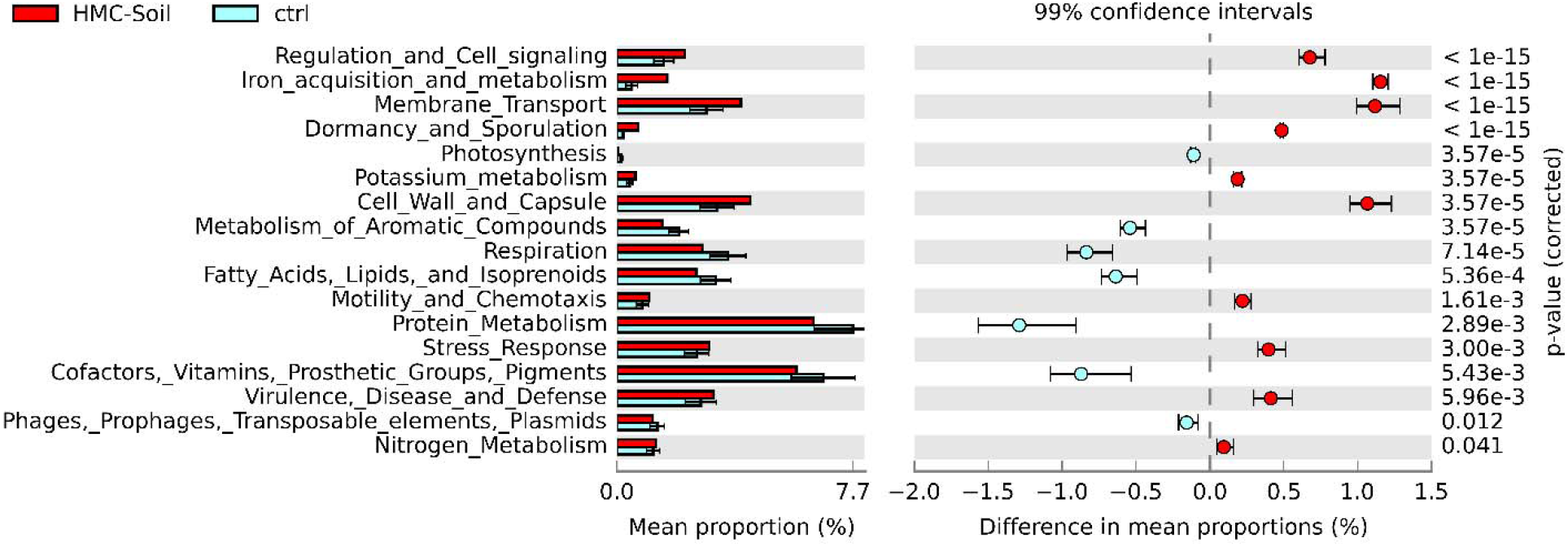
Functional comparative study of enriched heavy metal contaminated (HMC) soils with metagenomes derived from non-contaminated soils. Enriched gene categories in the EM metagenome has a positive difference between proportions (red circles), while depleted gene categories in the EM metagenome has a negative difference between proportions (blue circles). Bars on the left represent the proportion of each gene categories in the data. P value of ≥ 0.05 were considered significant.

### HM resistant genes analysis

Raw reads were assembled and genes in resulted metagenome were searched against HM resistance database^34^. This revealed available metal resistance genes in high resolution and their frequencies (Table 2, for more detailed Table refer to S-BacMet).

**Table 2:**
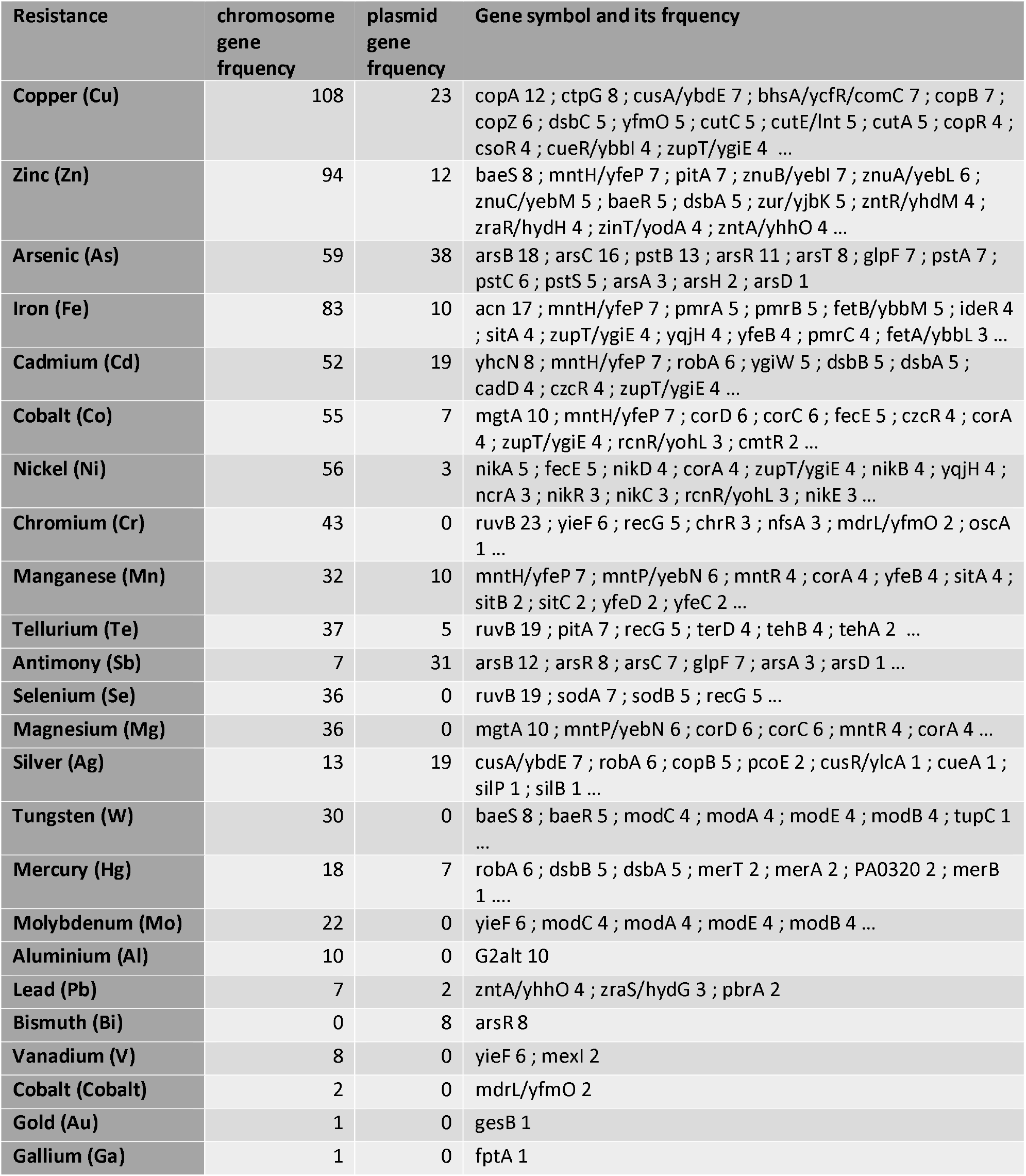
Available metal resistance genes in HMC soil metagenome, frequencies for each resistance categories and location of the genes. More information is available in the table S-BacMet.

Majority of HM resistant genes are located on the chromosome (Table 2). According to the table, copper resistance genes were most abundant in the metagenome, consisting 131 genes from 47 ortholog groups. From theses Cu resistance gene, CopA was more presented in the metagenome followed by ctpG, which both encode heavy metal efflux P-type ATPases. For zinc resistant genes top three most aboundant gene were baeS, yfeP and pitA, which encode two-component regulatory system, Divalent metal cation transporter and Low-affinity inorganic phosphate transporter respectively. An arsenic pump (arsB) is the most prevalent As resistant protein and arsenate reductase (arsC) being the second most prevalent resistance gene. Regarding Fe resistance genes, 17 aconitase (acn) were detected in the metagenome and 7 divalent metal cation transporter. In addition, 5 member of the PmrA-PmrB two-component system were present in the metagenome which are involved in controlling of bacterial response to external pH and iron. PmrA-PmrB regulates genes that modify lipopolysaccharide. yhcN, yfeP 7 and robA were main cadmium resistant genes available in the metagenome that encode proteins with unknown function, divalent metal cation transporter and transcriptional activator. RobA can confer resistance to antibiotics, silver, mercury, and cadmium. Among detected HM resistance genes, most frequently detected gene was ruvB. It encode for ATP-dependent DNA helicase, which is involved in repairing DNA damage. ArsB, which encode cytoplasmic arsenic pump protein, was the second most abundant HM resistant gene. RuvB and arsB are followed by acn, copA, mgtA and G2alt. MgtA encode for magnesium-transporting ATPase and G2alt has ATPase activity^40^.

Available HMR genes in EM revealed that different HMR gens are available in iron ore microbial community, namely different family of metal extrusion pumps(such as P-type ATPases, RNDs, MFS, Cation diffusion facilitators), metal chaperones, HM reductase, HM sensors, phosphatase, siderophore reductase, transcription activator/repressor proteins and kinase proteins.

## Discussion

### Taxonomic composition

Based in our results, taxonomic composition of the EM was drastically different from that of a typical soil, with a great enrichment of *Proteobacteria* mainly *γ-Proteobacteria*. There are many studies in which HM contaminated environments are taxonomically investigated. A previous study concluded arsenic contamination changed taxonomic composition of mediterranean marine sediments in favor of *Proteobacteria^38^*. Another study revealed that cadmium contamination of soil resulted in increase of *Proteobacteria* (from 39 to 59%) and decrease of *Acidobacteria* (18 to 6%)^41^. Christopher and coworkers showed that metal-contaminated (mainly uranium contaminated) groundwater microbial community was composed primarily of *γ-Proteobacteria*^42^. These three studies are in line with the present study. In addition, Li et al studied copper stress effect on microbial community in activated sludge. They showed after exposure to high copper stress, eukaryote portion of the community decreased significantly, which is also in line with the present study. Li and coworkers shoed that copper stress resulted in enrichment of *Firmicutes, Chloroflexi and Spirochaetes; however, Proteobacteria* were depleted after copper enrichment. Both community, i.e. before and after copper enrichment *Proteobacteria* were most abundant phyllum. ^24^. Depletion of eukaryote portion of the community after copper enrichment reported by Li is in accordance with the present study; therefore, we could conclude that in a community metal resistance genes are mainly carried out by prokaryotic part of the community. Gillan and coworkers evaluated long-term effect of metal contamination on sediment and reported that taxonomical profile did not changed significantly upon metal contamination at phylum level, but slight change was seen at genus level, seven bacterial genera were significantly more abundant in metal contaminated sites^13^. Very similar taxonomic composition between contaminated and uncontaminated sediment in Gillan study could be due to low HM concentration and longtime of contamination, enabling microbial community to adapt to new condition over long time. Pei et al compared soil sample from chromium contaminated and uncontaminated region and revealed that Cr contaminated sampling sites were taxonomically different from the uncontaminated ones. Particularly, the relative abundances of *Firmicutes* and *Bacteroidetes* were higher while *Actinobacteria* was lower in the contaminated group than uncontaminated group. In the Pei study most abundant phyla present with the contaminated group were *Proteobacteria, Bacteroidetes*, and *Firmicutes* respectively^43^. Similar to Pei study, taxonomic profile of the EM were dominated by *Proteobacteria* and *Firmicutes* but not *Bacteroidetes*.

There are different (sometimes even contrasting) results on the HM effect on taxonomical composition of the soil. Some studies claim that HM contamination has insignificant effect on taxonomical composition^13,21^, some reporting HM contamination results in enrichment of *Proteobacteria^38,41,44^* and in contrast some reports it results in depletion of *Proteobacteria*, others reports it results in enrichment of *Firmicutes^24,43^, Bacteroidetes^43^* or other phyllums. However even those studies reporting HM contamination leads to depletion of *Proteobacteria*, simultaneously exclaimed that after HM contamination *Proteobacteria* still comprise main portion of the community^13,21^. Therefore, we may conclude that HM contamination results in different taxonomic change based on the evolutionary history of the community, type/concentration of contaminated metal, physicochemical properties of environment and …; however, it seems that *Proteobacteria* are better adopted to HM contamination than other bacterial phylum. This could be explained by *Proteobacteria* highly plastic genome, displaying 10-fold genome size variation and genome plasticity^45^. At lower taxonomic levels, the situation is more complicated and hence generalization is more difficult.

### Functional composition

Enrichment of “Cell Wall and Capsule” subsystem in EM revealed that microbial community possibly cope with excess metal concentration by changing in their cell wall structure and secretion of capsules. In line with our study, it was shown that the most prominent difference between a HM contaminated and a less contaminated sediment is related to ‘cell wall and capsule’ subsystem which is more represented in contaminated sediment ^13^. Therefore non-specific binding of metals by extracytoplasmic part of the cell (outer membrane, cell wall, S-layer, lipopolysaccharide, extracellular polymeric substances) is present in the iron ore microbial community as a passive mechanism for metal exclusion. Deeper investigation showed that many genes related to “Cell Wall and Capsule” subsystem are enriched in the iom, such as peptidoglycan biosynthesis, lipopolysaccharide assembly, lipid A modifications and capsular polysaccharides Biosynthesis (Figure S-level3). This showed that several components of extracytoplasmic part of the cell are modified in response to HM. In addition, “Quorum sensing and biofilm formation” gene category is enriched in the EM. Generally biofilms are composed of extracellular polymeric substances and are capable of increasing HMR of attached communities^46^. There is evidence that planktonic *Pseudomonas aeruginosa* is more susceptible to HM relative to it’s biofilm form^47^. Therefore, we could conclude biofilm formation is also HMR strategy used by the microbial community. It has been demonstrate that quorum sensing increases resistance to osmotic, thermal and heavy metal stress^48^. Considering the iron ore metagenome was located in a hot semiarid region with high salinity, quorum sensing could be very useful to the bacterial community.

Enrichment of “Membrane Transport” category in the EM discloses another mechanism of HMR i.e. extrusion of extra HM ion out of cell. Many HM transporters from different classes are among the enriched genes including divalent metal transporter 1 (DMT1), cadmium-transporting ATPase, RND efflux pumps, CzcD (a cation diffusion facilitator) and ECF transporters(Table S-enrichedGenes), all with known function in cell metal homeostasis^49–51^. Therefore, efflux mechanism is obviously present in the EM as a HM resistance method. Metagenomic studies have found high levels of czcD in an acid mine drainage metagenome and in a zinc contaminated sediment community^13,52^. In activated sludge of a Cu-contaminated tannery wastewater copA was the most abundant of the Cu resistance systems ^53^. It also has been shown that in a cd contaminated soil most abundant pathway observed is ko02010, which is assigned to ABC transporters^41^.

Many of gene categories related to stress response, specifically oxidative stress response are enriched in EM and this shows that “ROS scavenging and DNA reparation” being present in EM as another resistance mechanism. Some of the enriched genes are “Glutathione Redox cycle”, “Universal stress response” and superoxide dismutase (Table S-enrichedGenes). Several reports indicate that microbes exposed to high concentration of metals upregulate genes that are involved in ROS scavenging, DNA reparation and hydrolysis of abnormally folded proteins^54–56^. Such systems are indirect HMR mechanism that repair damages induced by metals. At the community level genes involved in response to ROS were highly expressed in acid mine drainage communities ^52,57^.

Redox enzymes such as multicopper oxidase (CueO) and ferroxidase are also enriched in the EM revealing another mechanism of HM tolerance, i.e. transforming metal ion to less harmful ones. CueO expression confers copper tolerance in *Escherichia coli*^58^. Transformation of metal into less harmful ones is an important resistance strategy and periplasmic copper oxidation were shown to be more important mechanism than efflux in an extremely Cu-contaminated activated sludge^24^.

Iron- bounded siderophores moves into the cell efficiently but siderophores bound to other HM do not enter the cell readily and siderophore synthesis and secretion help the cell to tolerate HM in this way^12,59^. According to the present state of knowledge siderophore mediated resistance were not seen in microbial community and metallophore genes were not enriched in the studied metal contaminated communities^12^. In contrast, the EM contain plenty of genes related to siderophore synthesis and secretion and is significantly enriched relative to uncontaminated soils(Table enrichedGenes), indicating that one of the main mechanism of resistance by the community is siderophore synthesis and secretion.

Enrichment of “Potassium metabolism” category genes could be explained in two ways. First, as the iron ore was located in a semiarid region with high salinity, enrichment of potassium metabolism genes coud be a response to cope with high salinity of the environment. As shown by many studies potassium metabolism has a close relationship with salinity stress tolerance both in bacteria and plants^60,61^. Another explanation for the enrichment of potassium metabolism in the EM could be positive tolerance against elevated dose of HMs, as in the case for bacterial acidophiles. Acidophiles generating a chemiosmotic gradient by keeping up an inside positive cytoplasmic transmembrane potential and hindering metal cation entering the cytoplasm in this way ^62,63^. We could hypothesize that potassium ion could be used for similar strategy to cope with high level of salt and HMs simultaneously. Moreover some of metal efflux pumps use potassium gradient of plasma membrane as driving force for metal extrusion and potassium gradient may help the cell to cope with excess metal in this way too ^64^. Enrichment of glutathione-gated potassium efflux systems (Kef) that play role in protection against toxic electrophilic compounds^65^ supports our hypothesis and makes it more interesting. Cytoplasmic glutathione acts as a scavenger of electrophiles and its role in HM stress response is known ^66^. Glutathione mediated activation of Kef systems integrates K^+^ efflux and H+ influx, and the consequent acidification of cytoplasm protects against electrophile-mediated stress ^65^.

Amino acid metabolism related genes did not differ significantly between EM and control, but genes related to metabolism of “Lysine, threonine, methionine, and cysteine” were enriched in the EM (Tables S-level2 and S-level3). It is well known that sulfur containing molecule play key role in response to HMs, such as glutathione, H2S, cysteine and phytochelatine ^67,68^. Therefore, enrichment of cysteine metabolism could be justified as a response to need for sulfur containing molecules. Moreover, it has been shown that lysine catabolism in bacteria is related to osmotic stress resistance^69^.

Analysis of available metal resistance genes in EM metagenome showed that resistant genes for 24 metal were present in the metagenome even for those metals that were exist in the mine soil in very low content, like silver, zinc and arsenic (Table S-BacMet). Enrichment allowed us to detect many different resistance strategies that were used by the microbial community. Among HMR mechanism, metal ion export seems to be more prevailed (Figure 2 and Figure S-level3); meaning generally metal extrusion is the primary strategy for metal resistance in EM. Regarding to copper, chromium and arsenic it seems that transformation to less harmful ion is an important strategy beside extrusion. It has been reported that in a activated sludge genes related to copper oxidase after copper-enrichment were more prevalent relative to other copper resistant genes indicating that periplasmic copper oxidation is more important mechanism than efflux ^24^. This is in line with our study. Presence of extracellular phosphatase in the enriched genes(Table S-BacMet) suggest phosphate mediated precipitation of HMs is used by the microbial iron ore community against HMs. Presence of PmrG (encodes for lipopolysaccharide phosphatase) in EM metagenome support this claim. It has been shown that dephosphorylation of the Hep(II) phosphate in the core region of lipopolysaccharide by PmrG results in resistance against Fe^3+^ and Al^3+ 70^.

Based on available known HMR genes and comparative metagenomics we could concluded that 10 mechanism of HMR were available in the EM microbial community. Mechanism of HMR which were present in EM were: 1) metal extrusion, 2) cell wall modification, 3) secretion of exopolysaccharides, 4) modification of lipopolysaccharides, 5) detoxification of metals by change in oxidation state using redox enzymes, 6) precipitation of metals by extracellular phosphatases, 7) decreasing HM import into the cytoplasm by using metallophores, 8) scavenging ROS produced by HM, 9) DNA reparation, and 10) detoxification of metals by complexing with metal-binding proteins.

### Conclusion

Here we showed that high-throughput metagenomic sequencing of enriched metagenome could reveal resistance mechanism in high resolution. Bacteria cope with excess of HMs in their environments in different ways. Bacterial mechanism of adaptation to high concentrations of heavy metals have been shown in many cultivated species^71^. There are 22 known metal resistance mechanism in isolated bacteria and concern different cell partitions, i.e. the extracellular environment, outer membrane, periplasm, cytoplasmic membrane and the cytoplasm^12^. Of the 22 known metal-resistance mechanism, 14 were found in complex communities^12^. Based on the enriched genes and subsystems in the present study, we demonstrate that microbial community of iron ore use at least 10 mechanisms of HMR. The results of our study expand the current knowledge of the mechanism of HM resistance in microbial community of metal contaminated soil.

## Supporting information

https://ggenomics.ir/2020/07/xploring-metal-resistance-genes-and-mechanisms-in-copper-enriched-metal-ore-metagenome/

## Data Accessibility

- Raw DNA reads are available on the MG-RAST database under the project number mgp83855.
- Final DNA sequence assembly: uploaded to GenBank (accession number wil be added soon)
- Sampling locations and related information: available online (on the MG-RAST database)

## Author Contributions Statement

Esmaeil Forouzan: “wrote the main manuscript and prepared figures. All authors reviewed the manuscript.”

Ali Asghar Karkhane^1^, Bagher Yakhchali^1^*

## Supplementary information

<u>Table S-PCA:</u> principle component analysis for soil metagenomes used in this study at the level 1 of SEED subsystems.

<u>Table S-BacMet:</u> Available metal resistance genes in HMC soil metagenome, frequencies for each resistance categories and location of the genes.

<u>Table S-DatasetStat:</u> properties for metagenomes used in the study.

<u>Table S-enrichedGenes:</u> Enriched and depleted genes in the copper enriched metagenome

<u>Table S-level2:</u> Enriched and depleted level2 SEED subsystems in the copper enriched metagenome

<u>Table S-level3:</u> Enriched and depleted level3 SEED subsystems in the copper enriched metagenome

<u>Table S-mgmID:</u> 94 IDs for shotgun metagenomes downloaded from MG-RAST used as control for comparative study.

Figure S-class: taxonomic comparison of the copper enriched metagenome and the control group at class level.

Figure S-domain: taxonomic comparison of the copper enriched metagenome and the control group at domain level.

Figure S-level2: functional comparison of the copper enriched metagenome and the control group at level 2 of the SEED subsystem.

Figure S-level3: functional comparison of the copper enriched metagenome and the control group at level 3 of the SEED subsystem.

Figure S-Rarefraction: Rarefraction for the copper enriched metagenome

## References

1. Chowdhury, S., Mazumder, M. A. J., Al-Attas, O. & Husain, T. Heavy metals in drinking water: Occurrences, implications, and future needs in developing countries. Sci. Total Environ. 569–570, 476–488 (2016).

2. Su, C., Jiang, L. & Zhang, W. A review on heavy metal contamination in the soil worldwide: Situation, impact and remediation techniques. Environ. Skept. Crit. (2014).

3. Abdel-Aty, A. M., Ammar, N. S., Abdel Ghafar, H. H. & Ali, R. K. Biosorption of cadmium and lead from aqueous solution by fresh water alga Anabaena sphaerica biomass. J. Adv. Res. 4, 367–374 (2013).

4. Roberts, T. L. Cadmium and Phosphorous Fertilizers: The Issues and the Science. Procedia Eng. 83, 52–59 (2014).

5. Liu, J., Qu, W. & Kadiiska, M. B. Role of oxidative stress in cadmium toxicity and carcinogenesis. Toxicol. Appl. Pharmacol. 238, 209–214 (2009).

6. Nies, D. H. Microbial heavy-metal resistance. Appl. Microbiol. Biotechnol. 51, 730–750 (1999).

7. Aziz, H. A., Yusoff, M. S., Adlan, M. N., Adnan, N. H. & Alias, S. Physico-chemical removal of iron from semi-aerobic landfill leachate by limestone filter. Waste Manag. 24, 353–358 (2004).

8. Feng, D., Aldrich, C. & Tan, H. Treatment of acid mine water by use of heavy metal precipitation and ion exchange. Miner. Eng. 13, 623–642 (2000).

9. Cobbina, S. J., Duwiejuah, A. B., Quansah, R., Obiri, S. & Bakobie, N. Comparative Assessment of Heavy Metals in Drinking Water Sources in Two Small-Scale Mining Communities in Northern Ghana. Int. J. Environ. Res. Public. Health 12, 10620–10634 (2015).

10. Barakat, M. A. New trends in removing heavy metals from industrial wastewater. Arab. J. Chem. 4, 361–377 (2011).

11. Chen, L.-X. et al. Shifts in microbial community composition and function in the acidification of a lead/zinc mine tailings. Environ. Microbiol. 15, 2431–2444 (2013).

12. Gillan, D. C. Metal resistance systems in cultivated bacteria: are they found in complex communities? Curr. Opin. Biotechnol. 38, 123–130 (2016).

13. Gillan, D. C., Roosa, S., Kunath, B., Billon, G. & Wattiez, R. The long-term adaptation of bacterial communities in metal-contaminated sediments: a metaproteogenomic study. Environ. Microbiol. 17, 1991–2005 (2015).

14. Hemme, C. L. et al. Lateral Gene Transfer in a Heavy Metal-Contaminated-Groundwater Microbial Community. mBio 7, e02234–02215 (2016).

15. Li, A.-D., Li, L.-G. & Zhang, T. Exploring antibiotic resistance genes and metal resistance genes in plasmid metagenomes from wastewater treatment plants. Front. Microbiol. 6, 1025 (2015).

16. Luo, J. et al. Structural and functional variability in root-associated bacterial microbiomes of Cd/Zn hyperaccumulator Sedum alfredii. Appl. Microbiol. Biotechnol. 101, 7961–7976 (2017).

17. Zhang, X., Niu, J., Liang, Y., Liu, X. & Yin, H. Metagenome-scale analysis yields insights into the structure and function of microbial communities in a copper bioleaching heap. BMC Genet. 17, 21 (2016).

18. Li, Z., Ma, Z., van der Kuijp, T. J., Yuan, Z. & Huang, L. A review of soil heavy metal pollution from mines in China: Pollution and health risk assessment. Sci. Total Environ. 468–469, 843–853 (2014).

19. Zhuang, P., McBride, M. B., Xia, H., Li, N. & Li, Z. Health risk from heavy metals via consumption of food crops in the vicinity of Dabaoshan mine, South China. Sci. Total Environ. 407, 1551–1561 (2009).

20. Boughner, L. A. & Singh, P. Microbial Ecology: Where are we now? Postdoc J. J. Postdr. Res. Postdr. Aff. 4, 3–17 (2016).

21. Hong, C., Si, Y., Xing, Y. & Li, Y. Illumina MiSeq sequencing investigation on the contrasting soil bacterial community structures in different iron mining areas. Environ. Sci. Pollut. Res. Int. 22, 10788–10799 (2015).

22. Lee, S. & Hallam, S. J. Extraction of high molecular weight genomic DNA from soils and sediments. J. Vis. Exp. JoVE (2009). doi:10.3791/1569

23. Akcay, H., Oguz, A. & Karapire, C. Study of heavy metal pollution and speciation in Buyak Menderes and Gediz river sediments. Water Res. 37, 813–822 (2003).

24. Li, L.-G., Cai, L., Zhang, X.-X. & Zhang, T. Potentially novel copper resistance genes in copper-enriched activated sludge revealed by metagenomic analysis. Appl. Microbiol. Biotechnol. 98, 10255–10266 (2014).

25. Bolger, A. M., Lohse, M. & Usadel, B. Trimmomatic: a flexible trimmer for Illumina sequence data. Bioinforma. Oxf. Engl. 30, 2114–2120 (2014).

26. Magoč, T. & Salzberg, S. L. FLASH: fast length adjustment of short reads to improve genome assemblies. Bioinforma. Oxf. Engl. 27, 2957–2963 (2011).

27. Keegan, K. P., Glass, E. M. & Meyer, F. MG-RAST, a Metagenomics Service for Analysis of Microbial Community Structure and Function. Methods Mol. Biol. Clifton NJ 1399, 207–233 (2016).

28. Nurk, S., Meleshko, D., Korobeynikov, A. & Pevzner, P. A. metaSPAdes: a new versatile metagenomic assembler. Genome Res. 27, 824–834 (2017).

29. Forouzan, E., Maleki, M. S. M., Karkhane, A. A. & Yakhchali, B. Evaluation of nine popular de novo assemblers in microbial genome assembly. J. Microbiol. Methods (2017). doi:10.1016/j.mimet.2017.09.008

30. Forouzan, E., Shariati, P., Maleki, M. S. M., Karkhane, A. A. & Yakhchali, B. Practical evaluation of 11 de novo assemblers in metagenome assembly. J. Microbiol. Methods (2018). doi:10.1016/j.mimet.2018.06.007

31. Zhu, W., Lomsadze, A. & Borodovsky, M. Ab initio gene identification in metagenomic sequences. Nucleic Acids Res. 38, e132–e132 (2010).

32. Wilke, A. et al. A RESTful API for accessing microbial community data for MG-RAST. PLoS Comput. Biol. 11, e1004008 (2015).

33. Parks, D. H., Tyson, G. W., Hugenholtz, P. & Beiko, R. G. STAMP: statistical analysis of taxonomic and functional profiles. Bioinformatics 30, 3123–3124 (2014).

34. Pal, C., Bengtsson-Palme, J., Rensing, C., Kristiansson, E. & Larsson, D. G. J. BacMet: antibacterial biocide and metal resistance genes database. Nucleic Acids Res. 42, D737–D743 (2014).

35. Qing, X., Yutong, Z. & Shenggao, L. Assessment of heavy metal pollution and human health risk in urban soils of steel industrial city (Anshan), Liaoning, Northeast China. Ecotoxicol. Environ. Saf. 120, 377–385 (2015).

36. Jaffar, S. T. A. et al. The Extent of Heavy Metal Pollution and Their Potential Health Risk in Topsoils of the Massively Urbanized District of Shanghai. Arch. Environ. Contam. Toxicol. 73, 362–376 (2017).

37. Keegan, K. P. et al. A Platform-Independent Method for Detecting Errors in Metagenomic Sequencing Data: DRISEE. PLOS Comput. Biol. 8, e1002541 (2012).

38. Plewniak, F. et al. Metagenomic insights into microbial metabolism affecting arsenic dispersion in Mediterranean marine sediments. Mol. Ecol. 22, 4870–4883 (2013).

39. Vickers, C. E. et al. Isoprene synthesis protects transgenic tobacco plants from oxidative stress. Plant Cell Environ. 32, 520–531 (2009).

40. Nakajima, H., Kobayashi, K., Kobayashi, M., Asako, H. & Aono, R. Overexpression of the robA gene increases organic solvent tolerance and multiple antibiotic and heavy metal ion resistance in Escherichia coli. Appl. Environ. Microbiol. 61, 2302–2307 (1995).

41. Feng, G. et al. Metagenomic analysis of microbial community and function involved in cd-contaminated soil., Metagenomic analysis of microbial community and function involved in cd-contaminated soil. BMC Microbiol. BMC Microbiol. 18, 18, 11–11 (2018).

42. Hemme, C. L. et al. Metagenomic insights into evolution of a heavy metal-contaminated groundwater microbial community. ISME J. 4, 660–672 (2010).

43. Pei, Y., Yu, Z., Ji, J., Khan, A. & Li, X. Microbial Community Structure and Function Indicate the Severity of Chromium Contamination of the Yellow River. Front. Microbiol. 9, (2018).

44. Li, X. et al. Response of soil microbial communities and microbial interactions to long-term heavy metal contamination. Environ. Pollut. Barking Essex 1987 231, 908–917 (2017).

45. Boussau, B., Karlberg, E. O., Frank, A. C., Legault, B.-A. & Andersson, S. G. E. Computational inference of scenarios for α-proteobacterial genome evolution. Proc. Natl. Acad. Sci. 101, 9722–9727 (2004).

46. Harrison, J. J., Ceri, H. & Turner, R. J. Multimetal resistance and tolerance in microbial biofilms. Nat. Rev. Microbiol. 5, 928–938 (2007).

47. Teitzel, G. M. & Parsek, M. R. Heavy Metal Resistance of Biofilm and Planktonic Pseudomonas aeruginosa. Appl. Environ. Microbiol. 69, 2313–2320 (2003).

48. García-Contreras, R. et al. Quorum sensing enhancement of the stress response promotes resistance to quorum quenching and prevents social cheating. ISME J. 9, 115–125 (2015).

49. Garrick, M. D. et al. DMT1: Which metals does it transport? Biol. Res. 39, (2006).

50. Herrou, J. et al. Conserved ABC Transport System Regulated by the General Stress Response Pathways of Alpha-and Gammaproteobacteria. J. Bacteriol. 199, e00746–16 (2017).

51. Rodionov, D. A., Hebbeln, P., Gelfand, M. S. & Eitinger, T. Comparative and Functional Genomic Analysis of Prokaryotic Nickel and Cobalt Uptake Transporters: Evidence for a Novel Group of ATP-Binding Cassette Transporters. J. Bacteriol. 188, 317–327 (2006).

52. Chen, L. et al. Comparative metagenomic and metatranscriptomic analyses of microbial communities in acid mine drainage. ISME J. 9, 1579–1592 (2015).

53. Jia, S. et al. Metagenomic analysis of cadmium and copper resistance genes in activated sludge of a tannery wastewater treatment plant. J. Environ. Biol. 34, 375–380 (2013).

54. Miller, C. D. et al. Copper and cadmium: responses in Pseudomonas putida KT2440. Lett. Appl. Microbiol. 49, 775–783 (2009).

55. Poirier, I., Hammann, P., Kuhn, L. & Bertrand, M. Strategies developed by the marine bacterium Pseudomonas fluorescens BA3SM1 to resist metals: A proteome analysis. Aquat. Toxicol. Amst. Neth. 128–129, 215–232 (2013).

56. Zhang, X. et al. Global transcriptome analysis of hexavalent chromium stress responses in Staphylococcus aureus LZ-01. Ecotoxicol. Lond. Engl. 23, 1534–1545 (2014).

57. Bertin, P. N. et al. Metabolic diversity among main microorganisms inside an arsenic-rich ecosystem revealed by meta-and proteo-genomics. ISME J. 5, 1735–1747 (2011).

58. Grass, G. & Rensing, C. CueO Is a Multi-copper Oxidase That Confers Copper Tolerance in Escherichia coli. Biochem. Biophys. Res. Commun. 286, 902–908 (2001).

59. Neilands, J. B. Siderophores: structure and function of microbial iron transport compounds. J. Biol. Chem. 270, 26723–26726 (1995).

60. Wang, M., Zheng, Q., Shen, Q. & Guo, S. The Critical Role of Potassium in Plant Stress Response. Int. J. Mol. Sci. 14, 7370–7390 (2013).

61. Yaakop, A. S. et al. Characterization of the mechanism of prolonged adaptation to osmotic stress of *Jeotgalibacillus malaysiensis* via genome and transcriptome sequencing analyses. Sci. Rep. 6, 33660 (2016).

62. Dopson, M., Ossandon, F. J., Lövgren, L. & Holmes, D. S. Metal resistance or tolerance? Acidophiles confront high metal loads via both abiotic and biotic mechanisms. Front. Microbiol. 5, 157 (2014).

63. Wheaton, G., Counts, J., Mukherjee, A., Kruh, J. & Kelly, R. The Confluence of Heavy Metal Biooxidation and Heavy Metal Resistance: Implications for Bioleaching by Extreme Thermoacidophiles. Minerals 5, 397–451 (2015).

64. Nies, D. H. Efflux-mediated heavy metal resistance in prokaryotes. FEMS Microbiol. Rev. 27, 313–339

65. Roosild, T. P. et al. Mechanism of ligand-gated potassium efflux in bacterial pathogens. Proc. Natl. Acad. Sci. U. S. A. 107, 19784–19789 (2010).

66. Jozefczak, M., Remans, T., Vangronsveld, J. & Cuypers, A. Glutathione Is a Key Player in Metal-Induced Oxidative Stress Defenses. Int. J. Mol. Sci. 13, 3145–3175 (2012).

67. Giles, N. M. et al. Metal and Redox Modulation of Cysteine Protein Function. Chem. Biol. 10, 677–693 (2003).

68. Jia, H. et al. Hydrogen sulfide - cysteine cycle system enhances cadmium tolerance through alleviating cadmium-induced oxidative stress and ion toxicity in Arabidopsis roots. Sci. Rep. 6, 39702 (2016).

69. Neshich, I. A., Kiyota, E. & Arruda, P. Genome-wide analysis of lysine catabolism in bacteria reveals new connections with osmotic stress resistance. ISME J. 7, 2400–2410 (2013).

70. Nishino, K. et al. Identification of the lipopolysaccharide modifications controlled by the Salmonella PmrA/PmrB system mediating resistance to Fe(III) and Al(III). Mol. Microbiol. 61, 645–654 (2006).

71. Lemire, J. A., Harrison, J. J. & Turner, R. J. Antimicrobial activity of metals: mechanisms, molecular targets and applications. Nat. Rev. Microbiol. 11, 371–384 (2013).

