## Supplementary material for "Exploring metal resistance genes and mechanisms in copper enriched metal ore metagenome": https://ggenomics.ir/2020/07/xploring-metal-resistance-genes-and-mechanisms-in-copper-enriched-metal-ore-metagenome/: Exploring metal resistance genes Scientific Reports.docx

### Author Contributions Statement

**Esmaeil Forouzan: " wrote the main manuscript and prepared figures. All authors reviewed the manuscript."**

**Ali Asghar Karkhane^1^,**

**Bagher Yakhchali^1^***

### Competing interests

The authors declare no competing interests.

### Figure legend:

Figure 1: Taxonomic composition of copper enriched metagenome at phylum and genus level.

Figure 2: SEED subsystem (level1) distribution differences between the HMC (heavy metal contaminated) and control (HM uncontaminated) soil metagenomes.

Figure 3: Functional comparative study of enriched heavy metal contaminated (HMC) soils with metagenomes derived from non-contaminated soils.

Supplementary information.

Table S-PCA: principle component analysis for soil metagenomes used in this study at the level 1 of SEED subsystems.

Table S-BacMet: Available metal resistance genes in HMC soil metagenome, frequencies for each resistance categories and location of the genes.

Table S-DatasetStat: properties for metagenomes used in the study.

Table S-enrichedGenes: Enriched and depleted genes in the copper enriched metagenome

Table S-level2: Enriched and depleted level2 SEED subsystems in the copper enriched metagenome

Table S-level3: Enriched and depleted level3 SEED subsystems in the copper enriched metagenome

Table S-mgmID: 94 IDs for shotgun metagenomes downloaded from MG-RAST used as control for comparative study.

Figure S-class: taxonomic comparison of the copper enriched metagenome and the control group at class level.

Figure S-domain: taxonomic comparison of the copper enriched metagenome and the control group at domain level.

Figure S-level2: functional comparison of the copper enriched metagenome and the control group at level 2 of the SEED subsystem.

Figure S-level3: functional comparison of the copper enriched metagenome and the control group at level 3 of the SEED subsystem.

Figure S-Rarefraction: Rarefraction for the copper enriched metagenome

Table 1: Overview of dataset and resulted assembly

| Category | metagenome |
| --- | --- |
| Number of contigs(larger than 500 bp) | 9,347 |
| Total length (bp) | 152,316,274 |
| N50* (bp) | 56,486 |
| L50* (bp) | 500 |
| Number of N's per 100 kbp | 16 |
| Number of predicted genes | 136,177 |
| contig mean length (bp) | 9,578 |
| raw reads | 227,127,502 |
| Clean reads | 181,596,102 |
| raw data (bp) | 22,939,877,702 |
| clean data (bp) | 18,341,206,302 |

* N50 is the length for which the collection of all contigs of that length or longer covers at least half an assembly and L50 is the number of contigs available in the collection.

Figure 1: Taxonomic composition of copper enriched metagenome at phylum (A) and genus (B) level.

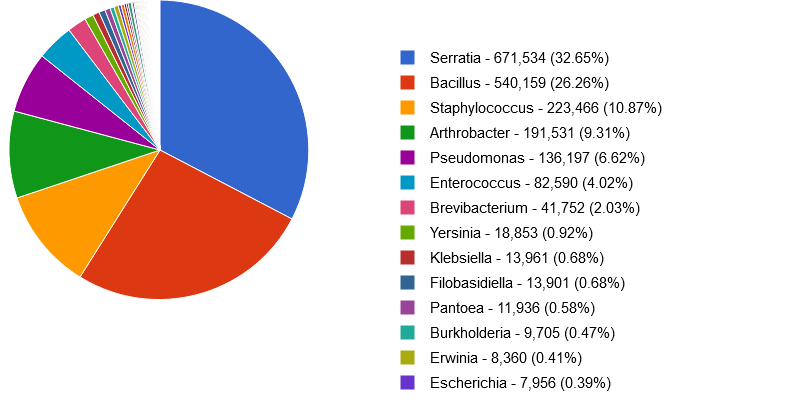

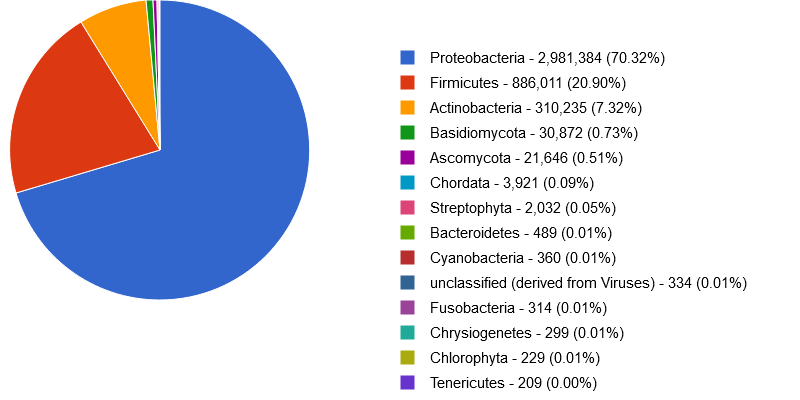

A)

B)

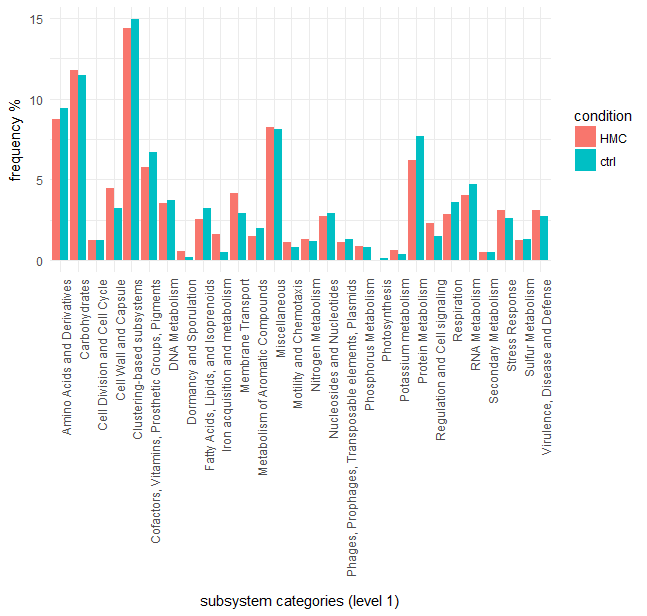

Figure 2: SEED subsystem (level1) distribution differences between the HMC (heavy metal contaminated) and control (HM uncontaminated) soil metagenomes based on MG-RAST annotation. The cutoff parameters for read annotation were default of MG-RAST (identity ≥ 60% and e-value ≤ 1x10-5). Comparison for level2, 3 and function level are available as supporting information (Figures and Tables names with prefix S-level2, S-level3 and S-Function).

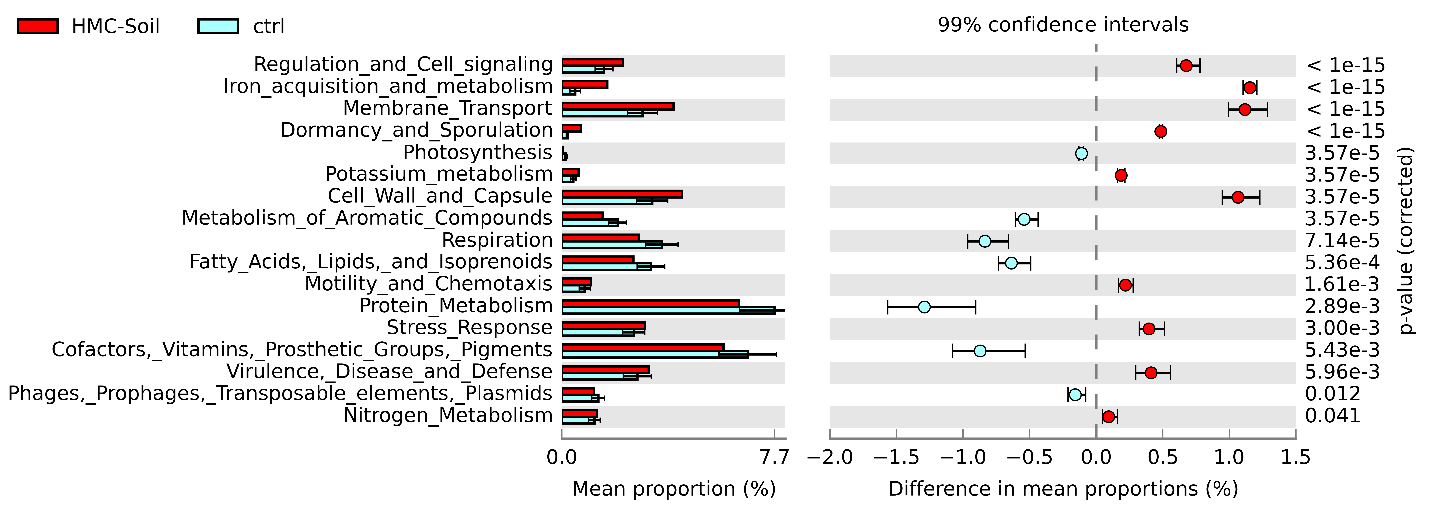

Figure 3: Functional comparative study of enriched heavy metal contaminated (HMC) soils with metagenomes derived from non-contaminated soils. Enriched gene categories in the EM metagenome has a positive difference between proportions (red circles), while depleted gene categories in the EM metagenome has a negative difference between proportions (blue circles). Bars on the left represent the proportion of each gene categories in the data. P value of ≥ 0.05 were considered significant.

Table 2: Available metal resistance genes in HMC soil metagenome, frequencies for each resistance categories and location of the genes. More information is available in the table S-BacMet.

| Resistance | chromosome gene frquency | plasmid gene frquency | Gene symbol and its frquency |
| --- | --- | --- | --- |
| Copper (Cu) | 108 | 23 | copA 12 ; ctpG 8 ; cusA/ybdE 7 ; bhsA/ycfR/comC 7 ; copB 7 ; copZ 6 ; dsbC 5 ; yfmO 5 ; cutC 5 ; cutE/lnt 5 ; cutA 5 ; copR 4 ; csoR 4 ; cueR/ybbI 4 ; zupT/ygiE 4 … |
| Zinc (Zn) | 94 | 12 | baeS 8 ; mntH/yfeP 7 ; pitA 7 ; znuB/yebI 7 ; znuA/yebL 6 ; znuC/yebM 5 ; baeR 5 ; dsbA 5 ; zur/yjbK 5 ; zntR/yhdM 4 ; zraR/hydH 4 ; zinT/yodA 4 ; zntA/yhhO 4 … |
| Arsenic (As) | 59 | 38 | arsB 18 ; arsC 16 ; pstB 13 ; arsR 11 ; arsT 8 ; glpF 7 ; pstA 7 ; pstC 6 ; pstS 5 ; arsA 3 ; arsH 2 ; arsD 1 |
| Iron (Fe) | 83 | 10 | acn 17 ; mntH/yfeP 7 ; pmrA 5 ; pmrB 5 ; fetB/ybbM 5 ; ideR 4 ; sitA 4 ; zupT/ygiE 4 ; yqjH 4 ; yfeB 4 ; pmrC 4 ; fetA/ybbL 3 … |
| Cadmium (Cd) | 52 | 19 | yhcN 8 ; mntH/yfeP 7 ; robA 6 ; ygiW 5 ; dsbB 5 ; dsbA 5 ; cadD 4 ; czcR 4 ; zupT/ygiE 4 … |
| Cobalt (Co) | 55 | 7 | mgtA 10 ; mntH/yfeP 7 ; corD 6 ; corC 6 ; fecE 5 ; czcR 4 ; corA 4 ; zupT/ygiE 4 ; rcnR/yohL 3 ; cmtR 2 … |
| Nickel (Ni) | 56 | 3 | nikA 5 ; fecE 5 ; nikD 4 ; corA 4 ; zupT/ygiE 4 ; nikB 4 ; yqjH 4 ; ncrA 3 ; nikR 3 ; nikC 3 ; rcnR/yohL 3 ; nikE 3 … |
| Chromium (Cr) | 43 | 0 | ruvB 23 ; yieF 6 ; recG 5 ; chrR 3 ; nfsA 3 ; mdrL/yfmO 2 ; oscA 1 … |
| Manganese (Mn) | 32 | 10 | mntH/yfeP 7 ; mntP/yebN 6 ; mntR 4 ; corA 4 ; yfeB 4 ; sitA 4 ; sitB 2 ; sitC 2 ; yfeD 2 ; yfeC 2 … |
| Tellurium (Te) | 37 | 5 | ruvB 19 ; pitA 7 ; recG 5 ; terD 4 ; tehB 4 ; tehA 2 … |
| Antimony (Sb) | 7 | 31 | arsB 12 ; arsR 8 ; arsC 7 ; glpF 7 ; arsA 3 ; arsD 1 … |
| Selenium (Se) | 36 | 0 | ruvB 19 ; sodA 7 ; sodB 5 ; recG 5 … |
| Magnesium (Mg) | 36 | 0 | mgtA 10 ; mntP/yebN 6 ; corD 6 ; corC 6 ; mntR 4 ; corA 4 … |
| Silver (Ag) | 13 | 19 | cusA/ybdE 7 ; robA 6 ; copB 5 ; pcoE 2 ; cusR/ylcA 1 ; cueA 1 ; silP 1 ; silB 1 … |
| Tungsten (W) | 30 | 0 | baeS 8 ; baeR 5 ; modC 4 ; modA 4 ; modE 4 ; modB 4 ; tupC 1 … |
| Mercury (Hg) | 18 | 7 | robA 6 ; dsbB 5 ; dsbA 5 ; merT 2 ; merA 2 ; PA0320 2 ; merB 1 …. |
| Molybdenum (Mo) | 22 | 0 | yieF 6 ; modC 4 ; modA 4 ; modE 4 ; modB 4 … |
| Aluminium (Al) | 10 | 0 | G2alt 10 |
| Lead (Pb) | 7 | 2 | zntA/yhhO 4 ; zraS/hydG 3 ; pbrA 2 |
| Bismuth (Bi) | 0 | 8 | arsR 8 |
| Vanadium (V) | 8 | 0 | yieF 6 ; mexI 2 |
| Cobalt (Cobalt) | 2 | 0 | mdrL/yfmO 2 |
| Gold (Au) | 1 | 0 | gesB 1 |
| Gallium (Ga) | 1 | 0 | fptA 1 |
