## Supplementary material for "Exploring metal resistance genes and mechanisms in copper enriched metal ore metagenome": https://ggenomics.ir/2020/07/xploring-metal-resistance-genes-and-mechanisms-in-copper-enriched-metal-ore-metagenome/: Figure S-domain.pdf

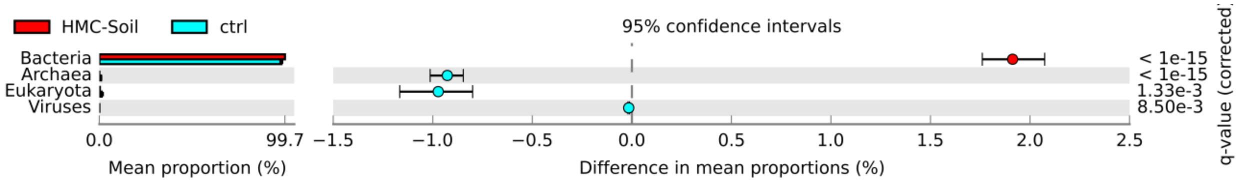


Figure S-domain: Comparative study of copper enriched metagenome and control group at domain level. As figure shows, enriched metagenome contained 98% Bacteria, 1.8% Eukaryota, 0.01% Viruses and negligible Archea. Enriched metagenome was enriched in bacteria and depleted in Eukaryota and Archea.
