## Supplementary material for "Exploring metal resistance genes and mechanisms in copper enriched metal ore metagenome": https://ggenomics.ir/2020/07/xploring-metal-resistance-genes-and-mechanisms-in-copper-enriched-metal-ore-metagenome/: Figure S-PCA.pdf

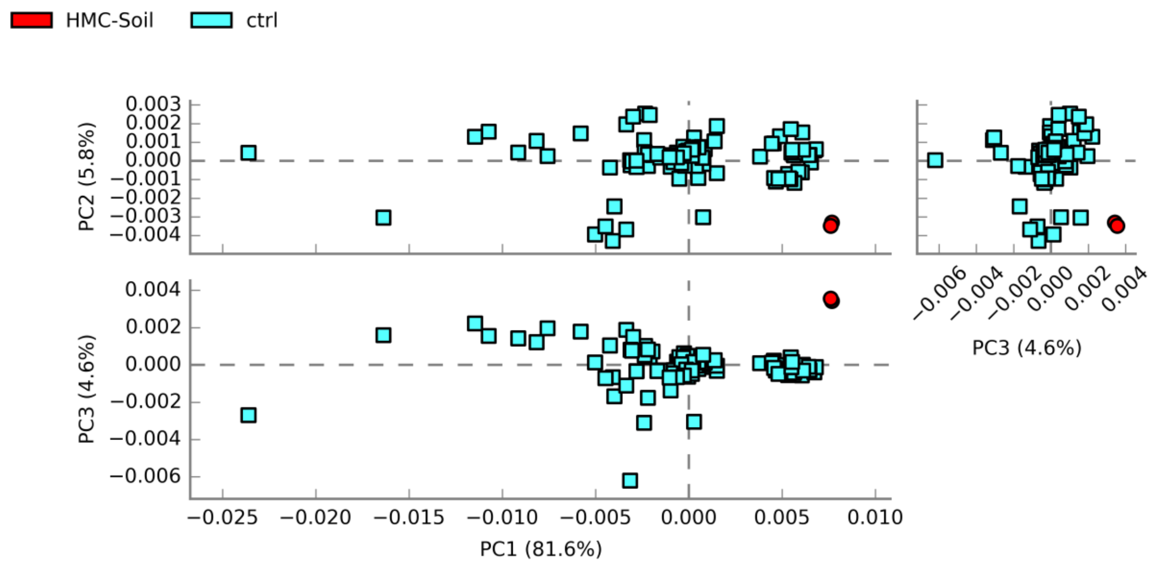


Figure S-PCA: principle component analysis for soil metagenomes used in this study at the level 1 of SEED subsystems.
