## Supplementary material for "Exploring metal resistance genes and mechanisms in copper enriched metal ore metagenome": https://ggenomics.ir/2020/07/xploring-metal-resistance-genes-and-mechanisms-in-copper-enriched-metal-ore-metagenome/: Figure S-Rarefraction.pdf

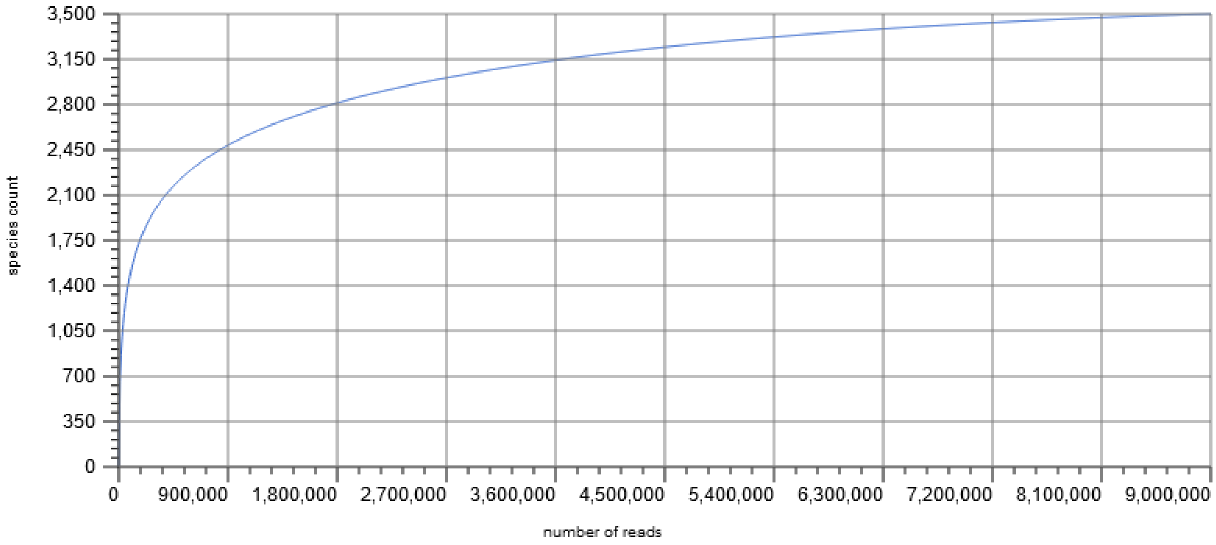


Figure S-Rarefraction: The plot shows the rarefaction curve of annotated species richness. This curve is a plot of the total number of distinct species annotations as a function of the number of sequences sampled. On the left, a steep slope indicates that a large fraction of the species diversity remains to be discovered. Rarefraction curve becomes almost flat to the right meaning that sequencing depth was enough to detect majority of the diversity available in the sample.
