## Supplementary material for "Exploring metal resistance genes and mechanisms in copper enriched metal ore metagenome": https://ggenomics.ir/2020/07/xploring-metal-resistance-genes-and-mechanisms-in-copper-enriched-metal-ore-metagenome/: Table S-enrichedGenes.pdf

| Table S-enrichedGenes: Enriched and depleted genes in the copper enriched metagenome | | | |
|---|---|---|---|
| Level | HMC-Soil freq (%) | ctrl freq (%) | p-values (corrected) |
| SS00121 | 0.134456458 | 0.016212986 | 0 |
| SS12009 | 0.134456458 | 0.016212986 | 0 |
| SS11792 | 0.143895653 | 0.050129421 | 0 |
| SS02740 | 0.113899619 | 0.029959073 | 0 |
| SS03480 | 0.113899619 | 0.029959073 | 0 |
| SS07927 | 0.113899619 | 0.029959073 | 0 |
| SS08330 | 0.113899619 | 0.029959073 | 0 |
| SS09057 | 0.113899619 | 0.029959073 | 0 |
| SS04452 | 0.100500727 | 0.031322855 | 0 |
| SS10830 | 0.100500727 | 0.031322855 | 0 |
| SS01490 | 0.072595514 | 0.008091818 | 0 |
| SS01781 | 0.072595514 | 0.008091818 | 0 |
| SS05380 | 0.072595514 | 0.008091818 | 0 |
| SS05957 | 0.076409256 | 0.012858654 | 0 |
| SS09992 | 0.076409256 | 0.012858654 | 0 |
| SS09993 | 0.076409256 | 0.012858654 | 0 |
| SS10198 | 0.085515591 | 0.024499646 | 0 |
| SS06798 | 0.071563466 | 0.011119842 | 0 |
| SS13839 | 0.071563466 | 0.011119842 | 0 |
| SS01426 | 0.062221687 | 0.006284228 | 0 |
| SS08338 | 0.06894163 | 0.016048806 | 0 |
| SS10959 | 0.06894163 | 0.016048806 | 0 |
| SS15494 | 0.06894163 | 0.016048806 | 0 |
| SS00858 | 0.14777678 | 0.096026678 | 0 |
| SS05827 | 0.14777678 | 0.096026678 | 0 |
| SS10169 | 0.14777678 | 0.096026678 | 0 |
| SS13543 | 0.14777678 | 0.096026678 | 0 |
| SS08615 | 0.068589776 | 0.01917396 | 0 |
| SS00068 | 0.114522084 | 0.066427626 | 0 |
| SS00173 | 0.114522084 | 0.066427626 | 0 |
| SS05372 | 0.114522084 | 0.066427626 | 0 |
| SS06014 | 0.114522084 | 0.066427626 | 0 |
| SS12702 | 0.114522084 | 0.066427626 | 0 |
| SS10936 | 0.059188032 | 0.013350795 | 0 |
| SS01983 | 0.060149356 | 0.015573616 | 0 |
| SS03471 | 0.060149356 | 0.015573616 | 0 |
| SS01687 | 0.132531586 | 0.08797307 | 0 |
| SS02192 | 0.132531586 | 0.08797307 | 0 |
| SS04009 | 0.132531586 | 0.08797307 | 0 |
| SS00916 | 0.057451312 | 0.014733657 | 0 |
| SS02005 | 0.057451312 | 0.014733657 | 0 |
| SS06021 | 0.057451312 | 0.014733657 | 0 |
| SS04682 | 0.041490022 | 0.000166064 | 0 |
| SS08268 | 0.041490022 | 0.000166064 | 0 |

| | | | |
|---|---|---|---|
| SS00002 | 0.078645162 | 0.039191746 | 0 |
| SS01189 | 0.078645162 | 0.039191746 | 0 |
| SS01253 | 0.078645162 | 0.039191746 | 0 |
| SS02705 | 0.078645162 | 0.039191746 | 0 |
| SS05411 | 0.078645162 | 0.039191746 | 0 |
| SS09053 | 0.078645162 | 0.039191746 | 0 |
| SS08273 | 0.078579817 | 0.039361638 | 0 |
| SS08289 | 0.078579817 | 0.039361638 | 0 |
| SS10465 | 0.049746291 | 0.011501295 | 0 |
| SS10665 | 0.049746291 | 0.011501295 | 0 |
| SS08601 | 0.059461912 | 0.021364434 | 0 |
| SS05593 | 0.061679351 | 0.023649324 | 0 |
| SS07725 | 0.037680776 | 0.000142779 | 0 |
| SS07726 | 0.037680776 | 0.000142779 | 0 |
| SS07587 | 0.08070491 | 0.045596059 | 0 |
| SS10171 | 0.08070491 | 0.045596059 | 0 |
| SS13545 | 0.08070491 | 0.045596059 | 0 |
| SS03872 | 0.046344609 | 0.011687586 | 0 |
| SS04498 | 0.046344609 | 0.011687586 | 0 |
| SS04699 | 0.046344609 | 0.011687586 | 0 |
| SS05632 | 0.036275528 | 0.002085633 | 0 |
| SS12900 | 0.036275528 | 0.002085633 | 0 |
| SS12909 | 0.036275528 | 0.002085633 | 0 |
| SS15514 | 0.10206246 | 0.068448318 | 0 |
| SS00108 | 0.0567535 | 0.023207188 | 0 |
| SS01221 | 0.0567535 | 0.023207188 | 0 |
| SS01859 | 0.045864934 | 0.013155998 | 0 |
| SS02196 | 0.045864934 | 0.013155998 | 0 |
| SS06820 | 0.045864934 | 0.013155998 | 0 |
| SS13846 | 0.045864934 | 0.013155998 | 0 |
| SS02890 | 0.034341158 | 0.001970169 | 0 |
| SS02891 | 0.034341158 | 0.001970169 | 0 |
| SS12049 | 0.034341158 | 0.001970169 | 0 |
| SS12050 | 0.034341158 | 0.001970169 | 0 |
| SS08124 | 0.065270357 | 0.033117085 | 0 |
| SS10180 | 0.065270357 | 0.033117085 | 0 |
| SS13546 | 0.065270357 | 0.033117085 | 0 |
| SS11213 | 0.032893047 | 0.000948138 | 0 |
| SS04564 | 0.038140551 | 0.006997826 | 0 |
| SS04666 | 0.038140551 | 0.006997826 | 0 |
| SS09837 | 0.035933573 | 0.004880928 | 0 |
| SS00920 | 0.03268926 | 0.002209014 | 0 |
| SS03779 | 0.03268926 | 0.002209014 | 0 |
| SS07255 | 0.03268926 | 0.002209014 | 0 |
| SS05642 | 0.04821344 | 0.01814247 | 0 |
| SS05711 | 0.04821344 | 0.01814247 | 0 |
| SS07748 | 0.04821344 | 0.01814247 | 0 |

| | | | |
|---|---|---|---|
| SS12898 | 0.04821344 | 0.01814247 | 0 |
| SS05663 | 0.032576165 | 0.002530272 | 0 |
| SS13096 | 0.032576165 | 0.002530272 | 0 |
| SS07276 | 0.036406046 | 0.006731775 | 0 |
| SS05662 | 0.032162419 | 0.002549485 | 0 |
| SS13095 | 0.032162419 | 0.002549485 | 0 |
| SS04132 | 0.03342701 | 0.003943683 | 0 |
| SS06703 | 0.03342701 | 0.003943683 | 0 |
| SS10376 | 0.040671947 | 0.011269845 | 0 |
| SS06999 | 0.043499268 | 0.014401685 | 0 |
| SS08779 | 0.045511187 | 0.016516168 | 0 |
| SS12740 | 0.045511187 | 0.016516168 | 0 |
| SS02852 | 0.051127039 | 0.02251845 | 0 |
| SS08380 | 0.051127039 | 0.02251845 | 0 |
| SS12953 | 0.051127039 | 0.02251845 | 0 |
| SS13468 | 0.03329834 | 0.004840785 | 0 |
| SS00820 | 0.029564486 | 0.001701549 | 0 |
| SS04825 | 0.036585436 | 0.00891885 | 0 |
| SS01984 | 0.028422716 | 0.000824839 | 0 |
| SS03472 | 0.028422716 | 0.000824839 | 0 |
| SS14508 | 0.027859023 | 0.000269902 | 0 |
| SS00125 | 0.041512572 | 0.014018159 | 0 |
| SS00747 | 0.032110185 | 0.004788739 | 0 |
| SS00791 | 0.032110185 | 0.004788739 | 0 |
| SS09838 | 0.032110185 | 0.004788739 | 0 |
| SS03356 | 0.030739026 | 0.004137862 | 0 |
| SS03023 | 0.035506166 | 0.008958581 | 0 |
| SS03744 | 0.035506166 | 0.008958581 | 0 |
| SS06691 | 0.030458114 | 0.003942269 | 0 |
| SS04537 | 0.033151994 | 0.006639188 | 0 |
| SS05488 | 0.048959025 | 0.022592171 | 0 |
| SS10992 | 0.026676832 | 0.000388679 | 0 |
| SS04315 | 0.043626104 | 0.017673974 | 0 |
| SS07689 | 0.043626104 | 0.017673974 | 0 |
| SS02255 | 0.087369488 | 0.061570525 | 0 |
| SS00460 | 0.037811488 | 0.01223895 | 0 |
| SS04830 | 0.037811488 | 0.01223895 | 0 |
| SS09966 | 0.037811488 | 0.01223895 | 0 |
| SS14995 | 0.037811488 | 0.01223895 | 0 |
| SS05829 | 0.033575799 | 0.008100957 | 0 |
| SS00153 | 0.025908629 | 0.000513796 | 0 |
| SS00465 | 0.025908629 | 0.000513796 | 0 |
| SS09067 | 0.109913528 | 0.084754017 | 0 |
| SS13226 | 0.030090719 | 0.005732131 | 0 |
| SS00454 | 0.030988568 | 0.006641362 | 0 |
| SS04294 | 0.030988568 | 0.006641362 | 0 |
| SS05971 | 0.030988568 | 0.006641362 | 0 |

| | | | |
|---|---|---|---|
| SS06027 | 0.030988568 | 0.006641362 | 0 |
| SS06047 | 0.030988568 | 0.006641362 | 0 |
| SS09635 | 0.030988568 | 0.006641362 | 0 |
| SS11925 | 0.030988568 | 0.006641362 | 0 |
| SS08128 | 0.04392531 | 0.019604708 | 0 |
| SS08151 | 0.04392531 | 0.019604708 | 0 |
| SS05990 | 0.026237566 | 0.002305709 | 0 |
| SS12907 | 0.023858285 | 0.000149222 | 0 |
| SS12908 | 0.023858285 | 0.000149222 | 0 |
| SS08537 | 0.029720398 | 0.006260835 | 0 |
| SS09568 | 0.029720398 | 0.006260835 | 0 |
| SS04512 | 0.023695079 | 0.000254079 | 0 |
| SS03061 | 0.030136025 | 0.007341579 | 0 |
| SS05777 | 0.030136025 | 0.007341579 | 0 |
| SS09473 | 0.030136025 | 0.007341579 | 0 |
| SS09761 | 0.030136025 | 0.007341579 | 0 |
| SS11807 | 0.030136025 | 0.007341579 | 0 |
| SS07141 | 0.031741023 | 0.009261352 | 0 |
| SS08116 | 0.031741023 | 0.009261352 | 0 |
| SS13414 | 0.025238494 | 0.002910953 | 0 |
| SS13434 | 0.025238494 | 0.002910953 | 0 |
| SS06055 | 0.033309844 | 0.011417812 | 0 |
| SS10748 | 0.033309844 | 0.011417812 | 0 |
| SS05379 | 0.023401951 | 0.001583328 | 0 |
| SS04772 | 0.044166673 | 0.022714672 | 0 |
| SS13105 | 0.044166673 | 0.022714672 | 0 |
| SS01090 | 0.038407963 | 0.017010293 | 0 |
| SS00459 | 0.031720274 | 0.010453633 | 0 |
| SS04829 | 0.031720274 | 0.010453633 | 0 |
| SS09965 | 0.031720274 | 0.010453633 | 0 |
| SS14994 | 0.031720274 | 0.010453633 | 0 |
| SS01929 | 0.024202569 | 0.003375491 | 0 |
| SS02324 | 0.024202569 | 0.003375491 | 0 |
| SS03722 | 0.024202569 | 0.003375491 | 0 |
| SS04242 | 0.054515691 | 0.0337556 | 0 |
| SS08616 | 0.030114794 | 0.009468364 | 0 |
| SS09979 | 0.024956825 | 0.004347087 | 0 |
| SS03498 | 0.032809029 | 0.012215467 | 0 |
| SS05683 | 0.032809029 | 0.012215467 | 0 |
| SS13908 | 0.032809029 | 0.012215467 | 0 |
| SS15497 | 0.032809029 | 0.012215467 | 0 |
| SS06849 | 0.020935499 | 0.000416763 | 0 |
| SS10812 | 0.028945875 | 0.008473894 | 0 |
| SS07898 | 0.027207882 | 0.007160127 | 0 |
| SS11547 | 0.022799283 | 0.002840438 | 0 |
| SS10519 | 0.030736698 | 0.011155607 | 0 |
| SS08109 | 0.031870737 | 0.012378748 | 0 |

| | | | |
|---|---|---|---|
| SS13333 | 0.031870737 | 0.012378748 | 0 |
| SS00154 | 0.070017746 | 0.050526833 | 0 |
| SS01703 | 0.070017746 | 0.050526833 | 0 |
| SS08169 | 0.02124793 | 0.001764584 | 0 |
| SS08849 | 0.02124793 | 0.001764584 | 0 |
| SS01230 | 0.024840221 | 0.00536891 | 0 |
| SS01069 | 0.024798952 | 0.005395038 | 0 |
| SS03088 | 0.031224276 | 0.011836909 | 0 |
| SS05084 | 0.031224276 | 0.011836909 | 0 |
| SS05368 | 0.031224276 | 0.011836909 | 0 |
| SS05376 | 0.031224276 | 0.011836909 | 0 |
| SS08215 | 0.031224276 | 0.011836909 | 0 |
| SS10095 | 0.031224276 | 0.011836909 | 0 |
| SS10804 | 0.031224276 | 0.011836909 | 0 |
| SS13536 | 0.031224276 | 0.011836909 | 0 |
| SS01361 | 0.026978654 | 0.00760392 | 0 |
| SS11550 | 0.026978654 | 0.00760392 | 0 |
| SS11754 | 0.026978654 | 0.00760392 | 0 |
| SS02726 | 0.019618949 | 0.000389113 | 0 |
| SS04596 | 0.019618949 | 0.000389113 | 0 |
| SS11700 | 0.048285357 | 0.029370991 | 0 |
| SS09457 | 0.028138524 | 0.009296174 | 0 |
| SS06044 | 0.019077886 | 0.000252934 | 0 |
| SS10744 | 0.019077886 | 0.000252934 | 0 |
| SS08280 | 0.021720449 | 0.003162986 | 0 |
| SS06286 | 0.02906889 | 0.010681477 | 0 |
| SS06782 | 0.021254101 | 0.002913142 | 0 |
| SS11704 | 0.020587453 | 0.002691846 | 0 |
| SS04267 | 0.020290116 | 0.002443598 | 0 |
| SS10719 | 0.020290116 | 0.002443598 | 0 |
| SS05978 | 0.029408242 | 0.011565556 | 0 |
| SS06013 | 0.029408242 | 0.011565556 | 0 |
| SS11100 | 0.029408242 | 0.011565556 | 0 |
| SS06340 | 0.018720664 | 0.000978497 | 0 |
| SS06435 | 0.018720664 | 0.000978497 | 0 |
| SS12152 | 0.018720664 | 0.000978497 | 0 |
| SS13634 | 0.018720664 | 0.000978497 | 0 |
| SS10947 | 0.021830251 | 0.004130519 | 0 |
| SS01306 | 0.026979491 | 0.009641976 | 0 |
| SS03937 | 0.026979491 | 0.009641976 | 0 |
| SS13405 | 0.026979491 | 0.009641976 | 0 |
| SS13424 | 0.026979491 | 0.009641976 | 0 |
| SS04414 | 0.018008228 | 0.000782553 | 0 |
| SS10954 | 0.018008228 | 0.000782553 | 0 |
| SS08278 | 0.035539842 | 0.018461563 | 0 |
| SS13404 | 0.035474796 | 0.018700073 | 0 |
| SS13423 | 0.035474796 | 0.018700073 | 0 |

| | | | |
|---|---|---|---|
| SS04127 | 0.034519035 | 0.017763267 | 0 |
| SS09540 | 0.021790634 | 0.005063123 | 0 |
| SS03083 | 0.022040496 | 0.005417949 | 0 |
| SS05443 | 0.022040496 | 0.005417949 | 0 |
| SS08304 | 0.023974786 | 0.007521747 | 0 |
| SS07251 | 0.017046651 | 0.000651762 | 0 |
| SS04375 | 0.024355074 | 0.008014128 | 0 |
| SS09869 | 0.023315628 | 0.007096793 | 0 |
| SS10574 | 0.033112491 | 0.016933415 | 0 |
| SS00948 | 0.022533477 | 0.006428654 | 0 |
| SS06404 | 0.017924783 | 0.001830245 | 0 |
| SS14801 | 0.025181006 | 0.009301545 | 0 |
| SS15058 | 0.025181006 | 0.009301545 | 0 |
| SS10310 | 0.020529048 | 0.004807186 | 0 |
| SS06280 | 0.026143626 | 0.010498265 | 0 |
| SS11566 | 0.016480916 | 0.000881144 | 0 |
| SS11771 | 0.016480916 | 0.000881144 | 0 |
| SS06190 | 0.018869694 | 0.003470055 | 0 |
| SS14806 | 0.018587704 | 0.003249268 | 0 |
| SS15063 | 0.018587704 | 0.003249268 | 0 |
| SS08715 | 0.022106953 | 0.006912485 | 0 |
| SS14999 | 0.022106953 | 0.006912485 | 0 |
| SS15012 | 0.022106953 | 0.006912485 | 0 |
| SS10508 | 0.015526107 | 0.000347994 | 0 |
| SS09570 | 0.017794863 | 0.002729641 | 0 |
| SS09608 | 0.017794863 | 0.002729641 | 0 |
| SS09336 | 0.023922586 | 0.009008357 | 0 |
| SS01865 | 0.03120129 | 0.016406063 | 0 |
| SS11720 | 0.016312777 | 0.001644264 | 0 |
| SS11775 | 0.016312777 | 0.001644264 | 0 |
| SS01924 | 0.01507747 | 0.000479045 | 0 |
| SS03721 | 0.01507747 | 0.000479045 | 0 |
| SS09830 | 0.026540064 | 0.011966626 | 0 |
| SS01579 | 0.021644173 | 0.007072105 | 0 |
| SS09138 | 0.021644173 | 0.007072105 | 0 |
| SS08279 | 0.01913597 | 0.004679217 | 0 |
| SS06728 | 0.019956098 | 0.005517975 | 0 |
| SS03952 | 0.023416174 | 0.009000179 | 0 |
| SS11565 | 0.023416174 | 0.009000179 | 0 |
| SS11770 | 0.023416174 | 0.009000179 | 0 |
| SS00765 | 0.029442216 | 0.01516316 | 0 |
| SS09846 | 0.029442216 | 0.01516316 | 0 |
| SS06266 | 0.020626038 | 0.006355415 | 0 |
| SS15367 | 0.020626038 | 0.006355415 | 0 |
| SS08230 | 0.043427959 | 0.02920785 | 0 |
| SS11584 | 0.043427959 | 0.02920785 | 0 |
| SS14269 | 0.043427959 | 0.02920785 | 0 |

| | | | |
|---|---|---|---|
| SS14291 | 0.043427959 | 0.02920785 | 0 |
| SS14349 | 0.043427959 | 0.02920785 | 0 |
| SS01077 | 0.015802729 | 0.00159797 | 0 |
| SS08224 | 0.028670686 | 0.014568647 | 0 |
| SS11582 | 0.028670686 | 0.014568647 | 0 |
| SS11585 | 0.028670686 | 0.014568647 | 0 |
| SS14217 | 0.028670686 | 0.014568647 | 0 |
| SS14276 | 0.028670686 | 0.014568647 | 0 |
| SS14336 | 0.028670686 | 0.014568647 | 0 |
| SS14876 | 0.028670686 | 0.014568647 | 0 |
| SS15399 | 0.028670686 | 0.014568647 | 0 |
| SS00778 | 0.032524848 | 0.01845134 | 0 |
| SS06761 | 0.014527276 | 0.000457671 | 0 |
| SS08432 | 0.014527276 | 0.000457671 | 0 |
| SS08469 | 0.014527276 | 0.000457671 | 0 |
| SS02982 | 0.017633055 | 0.003566418 | 0 |
| SS05374 | 0.016120047 | 0.002064118 | 0 |
| SS07706 | 0.02995059 | 0.015955938 | 0 |
| SS10076 | 0.014241203 | 0.000287449 | 0 |
| SS13415 | 0.026545799 | 0.012641913 | 0 |
| SS13425 | 0.026545799 | 0.012641913 | 0 |
| SS05980 | 0.015720684 | 0.001877488 | 0 |
| SS06018 | 0.015720684 | 0.001877488 | 0 |
| SS11101 | 0.015720684 | 0.001877488 | 0 |
| SS04538 | 0.017443491 | 0.003610098 | 0 |
| SS08447 | 0.014761872 | 0.000982392 | 0 |
| SS02784 | 0.024346024 | 0.010574327 | 0 |
| SS14260 | 0.024346024 | 0.010574327 | 0 |
| SS14287 | 0.024346024 | 0.010574327 | 0 |
| SS14367 | 0.024346024 | 0.010574327 | 0 |
| SS06468 | 0.037924617 | 0.024187827 | 0 |
| SS09556 | 0.037924617 | 0.024187827 | 0 |
| SS10462 | 0.037924617 | 0.024187827 | 0 |
| SS09532 | 0.02633256 | 0.012720901 | 0 |
| SS10455 | 0.02633256 | 0.012720901 | 0 |
| SS03912 | 0.018256725 | 0.004660058 | 0 |
| SS04932 | 0.018256725 | 0.004660058 | 0 |
| SS10192 | 0.016778804 | 0.003253269 | 0 |
| SS10648 | 0.016778804 | 0.003253269 | 0 |
| SS08275 | 0.019105047 | 0.005591731 | 0 |
| SS08291 | 0.019105047 | 0.005591731 | 0 |
| SS06852 | 0.014633879 | 0.001138321 | 0 |
| SS02397 | 0.030782291 | 0.017486618 | 0 |
| SS02500 | 0.030782291 | 0.017486618 | 0 |
| SS00361 | 0.028072548 | 0.014815901 | 0 |
| SS10739 | 0.028072548 | 0.014815901 | 0 |
| SS05804 | 0.015931285 | 0.002710432 | 0 |

| | | | |
|---|---|---|---|
| SS15539 | 0.015931285 | 0.002710432 | 0 |
| SS07661 | 0.014329178 | 0.001133173 | 0 |
| SS06895 | 0.022885732 | 0.009695948 | 0 |
| SS11868 | 0.022885732 | 0.009695948 | 0 |
| SS07628 | 0.013190012 | 1.57E-05 | 0 |
| SS09114 | 0.018796905 | 0.005642552 | 0 |
| SS05445 | 0.01731045 | 0.004184647 | 0 |
| SS04654 | 0.021646375 | 0.008553667 | 0 |
| SS03529 | 0.02408234 | 0.01101315 | 0 |
| SS11236 | 0.02408234 | 0.01101315 | 0 |
| SS11706 | 0.02408234 | 0.01101315 | 0 |
| SS04367 | 0.013734756 | 0.000716105 | 0 |
| SS01200 | 0.029646003 | 0.016642985 | 0 |
| SS02707 | 0.030904802 | 0.018041328 | 0 |
| SS09145 | 0.030904802 | 0.018041328 | 0 |
| SS02173 | 0.036207385 | 0.023371906 | 0 |
| SS06803 | 0.013220419 | 0.000416231 | 0 |
| SS12093 | 0.013220419 | 0.000416231 | 0 |
| SS12130 | 0.013220419 | 0.000416231 | 0 |
| SS02115 | 0.040573385 | 0.0277926 | 0 |
| SS12870 | 0.040573385 | 0.0277926 | 0 |
| SS09591 | 0.014957092 | 0.00224016 | 0 |
| SS00124 | 0.041096348 | 0.028497283 | 0 |
| SS08857 | 0.014018467 | 0.001442746 | 0 |
| SS15536 | 0.014018467 | 0.001442746 | 0 |
| SS04212 | 0.013686766 | 0.001125587 | 0 |
| SS08312 | 0.013337195 | 0.00079197 | 0 |
| SS03012 | 0.014030683 | 0.001610424 | 0 |
| SS06448 | 0.013788516 | 0.001426434 | 0 |
| SS00806 | 0.014160523 | 0.001800414 | 0 |
| SS09436 | 0.014160523 | 0.001800414 | 0 |
| SS11557 | 0.013333788 | 0.000984913 | 0 |
| SS02376 | 0.035319629 | 0.023009576 | 0 |
| SS04514 | 0.035319629 | 0.023009576 | 0 |
| SS14804 | 0.013695816 | 0.001419159 | 0 |
| SS15061 | 0.013695816 | 0.001419159 | 0 |
| SS13411 | 0.023110235 | 0.010863373 | 0 |
| SS13431 | 0.023110235 | 0.010863373 | 0 |
| SS08719 | 0.015461049 | 0.003296084 | 0 |
| SS10698 | 0.015461049 | 0.003296084 | 0 |
| SS11551 | 0.012346175 | 0.000229599 | 0 |
| SS11755 | 0.012346175 | 0.000229599 | 0 |
| SS05418 | 0.012471324 | 0.000466395 | 0 |
| SS14786 | 0.01276709 | 0.000835934 | 0 |
| SS15051 | 0.01276709 | 0.000835934 | 0 |
| SS08154 | 0.014818718 | 0.002973892 | 0 |
| SS09601 | 0.01324017 | 0.001445075 | 0 |

| | | | |
|---|---|---|---|
| SS10782 | 0.032699032 | 0.02091672 | 0 |
| SS01317 | 0.024796268 | 0.013030261 | 0 |
| SS03958 | 0.024796268 | 0.013030261 | 0 |
| SS10008 | 0.024796268 | 0.013030261 | 0 |
| SS11546 | 0.012239894 | 0.000485713 | 0 |
| SS11752 | 0.012239894 | 0.000485713 | 0 |
| SS08322 | 0.01285214 | 0.001118005 | 0 |
| SS13852 | 0.014069141 | 0.002352314 | 0 |
| SS01194 | 0.044176606 | 0.032479793 | 0 |
| SS01259 | 0.044176606 | 0.032479793 | 0 |
| SS00427 | 0.023985751 | 0.01233614 | 0 |
| SS02400 | 0.013602679 | 0.001964481 | 0 |
| SS07278 | 0.013602679 | 0.001964481 | 0 |
| SS09544 | 0.017071128 | 0.005490056 | 0 |
| SS01240 | 0.012157206 | 0.000605805 | 0 |
| SS05543 | 0.012157206 | 0.000605805 | 0 |
| SS01639 | 0.016032038 | 0.004490886 | 0 |
| SS01656 | 0.016032038 | 0.004490886 | 0 |
| SS07716 | 0.011899338 | 0.00038244 | 0 |
| SS05455 | 0.018624522 | 0.007128428 | 0 |
| SS04616 | 0.016854999 | 0.005378265 | 0 |
| SS14987 | 0.016854999 | 0.005378265 | 0 |
| SS13859 | 0.012583214 | 0.001172462 | 0 |
| SS07896 | 0.017879477 | 0.006527015 | 0 |
| SS00326 | 0.018249189 | 0.006898658 | 0 |
| SS14887 | 0.018249189 | 0.006898658 | 0 |
| SS08437 | 0.03311982 | 0.02177934 | 0 |
| SS08472 | 0.03311982 | 0.02177934 | 0 |
| SS10983 | 0.03311982 | 0.02177934 | 0 |
| SS02866 | 0.022516811 | 0.011197028 | 0 |
| SS12033 | 0.022516811 | 0.011197028 | 0 |
| SS13026 | 0.022516811 | 0.011197028 | 0 |
| SS10408 | 0.012037987 | 0.000804953 | 0 |
| SS01632 | 0.016963207 | 0.005800416 | 0 |
| SS02374 | 0.016963207 | 0.005800416 | 0 |
| SS06743 | 0.016963207 | 0.005800416 | 0 |
| SS00769 | 0.013186685 | 0.002079733 | 0 |
| SS04578 | 0.030874911 | 0.019768676 | 0 |
| SS04615 | 0.030874911 | 0.019768676 | 0 |
| SS04680 | 0.030874911 | 0.019768676 | 0 |
| SS04354 | 0.019967557 | 0.008889871 | 0 |
| SS13128 | 0.019967557 | 0.008889871 | 0 |
| SS06309 | 0.011697993 | 0.000661911 | 0 |
| SS06799 | 0.011697993 | 0.000661911 | 0 |
| SS01881 | 0.014933854 | 0.003933456 | 0 |
| SS13966 | 0.014933854 | 0.003933456 | 0 |
| SS06326 | 0.011327673 | 0.000359587 | 0 |

| | | | |
|---|---|---|---|
| SS06369 | 0.011327673 | 0.000359587 | 0 |
| SS06426 | 0.011327673 | 0.000359587 | 0 |
| SS12134 | 0.011327673 | 0.000359587 | 0 |
| SS00160 | 0.011628887 | 0.000676816 | 0 |
| SS00466 | 0.011628887 | 0.000676816 | 0 |
| SS05653 | 0.015671971 | 0.004734777 | 0 |
| SS12922 | 0.015671971 | 0.004734777 | 0 |
| SS06325 | 0.011327673 | 0.000468607 | 0 |
| SS06425 | 0.011327673 | 0.000468607 | 0 |
| SS08307 | 0.016389936 | 0.005530935 | 0 |
| SS00461 | 0.028547785 | 0.017712118 | 0 |
| SS03066 | 0.01191324 | 0.001108766 | 0 |
| SS13229 | 0.014425962 | 0.003633915 | 0 |
| SS06480 | 0.017221397 | 0.006454049 | 0 |
| SS08599 | 0.032496403 | 0.021732086 | 0 |
| SS02431 | 0.041703868 | 0.030954542 | 0 |
| SS13829 | 0.041703868 | 0.030954542 | 0 |
| SS04292 | 0.01112252 | 0.000409929 | 0 |
| SS05970 | 0.01112252 | 0.000409929 | 0 |
| SS04774 | 0.015084317 | 0.004379243 | 0 |
| SS13107 | 0.015084317 | 0.004379243 | 0 |
| SS12298 | 0.011402583 | 0.000722334 | 0 |
| SS01205 | 0.013550801 | 0.002986908 | 0 |
| SS01270 | 0.013550801 | 0.002986908 | 0 |
| SS02613 | 0.013550801 | 0.002986908 | 0 |
| SS15319 | 0.012276667 | 0.001718864 | 0 |
| SS04133 | 0.025085943 | 0.014624577 | 0 |
| SS06704 | 0.025085943 | 0.014624577 | 0 |
| SS15439 | 0.017325992 | 0.006905751 | 0 |
| SS02180 | 0.025498886 | 0.015106728 | 0 |
| SS06884 | 0.025498886 | 0.015106728 | 0 |
| SS14223 | 0.025498886 | 0.015106728 | 0 |
| SS02561 | 0.020211971 | 0.009973517 | 0 |
| SS02731 | 0.020211971 | 0.009973517 | 0 |
| SS11235 | 0.020211971 | 0.009973517 | 0 |
| SS11556 | 0.020211971 | 0.009973517 | 0 |
| SS03507 | 0.021363387 | 0.011158098 | 0 |
| SS13911 | 0.021363387 | 0.011158098 | 0 |
| SS00061 | 0.019210525 | 0.009008956 | 0 |
| SS00814 | 0.019210525 | 0.009008956 | 0 |
| SS02837 | 0.019210525 | 0.009008956 | 0 |
| SS03673 | 0.014046029 | 0.003846584 | 0 |
| SS07081 | 0.014046029 | 0.003846584 | 0 |
| SS09107 | 0.011580334 | 0.001410804 | 0 |
| SS09973 | 0.011580334 | 0.001410804 | 0 |
| SS00250 | 0.01049401 | 0.000435273 | 0 |
| SS07145 | 0.01049401 | 0.000435273 | 0 |

| | | | |
|---|---|---|---|
| SS08248 | 0.015986043 | 0.005947309 | 0 |
| SS04245 | 0.013419882 | 0.003390273 | 0 |
| SS03941 | 0.037611222 | 0.027584032 | 0 |
| SS04668 | 0.037611222 | 0.027584032 | 0 |
| SS05846 | 0.021490062 | 0.011481578 | 0 |
| SS07142 | 0.021222088 | 0.011218131 | 0 |
| SS08120 | 0.021222088 | 0.011218131 | 0 |
| SS15368 | 0.021222088 | 0.011218131 | 0 |
| SS00377 | 0.010374309 | 0.000396508 | 0 |
| SS01718 | 0.010374309 | 0.000396508 | 0 |
| SS10200 | 0.010374309 | 0.000396508 | 0 |
| SS10753 | 0.010374309 | 0.000396508 | 0 |
| SS00101 | 0.01457623 | 0.004616693 | 0 |
| SS08915 | 0.011275393 | 0.001383978 | 0 |
| SS03506 | 0.019417283 | 0.009533243 | 0 |
| SS13910 | 0.019417283 | 0.009533243 | 0 |
| SS02369 | 0.014805573 | 0.005049327 | 0 |
| SS04562 | 0.014805573 | 0.005049327 | 0 |
| SS04664 | 0.014805573 | 0.005049327 | 0 |
| SS12468 | 0.009862093 | 0.000115433 | 0 |
| SS12487 | 0.009862093 | 0.000115433 | 0 |
| SS08618 | 0.012947765 | 0.003275625 | 0 |
| SS08133 | 0.009867461 | 0.000201754 | 0 |
| SS08155 | 0.009867461 | 0.000201754 | 0 |
| SS05364 | 0.01313 | 0.003465612 | 0 |
| SS05373 | 0.01313 | 0.003465612 | 0 |
| SS06467 | 0.03221642 | 0.022560207 | 0 |
| SS09555 | 0.03221642 | 0.022560207 | 0 |
| SS10461 | 0.03221642 | 0.022560207 | 0 |
| SS12471 | 0.01067316 | 0.001038435 | 0 |
| SS05444 | 0.011713856 | 0.00208782 | 0 |
| SS13416 | 0.012133452 | 0.002508608 | 0 |
| SS13426 | 0.012133452 | 0.002508608 | 0 |
| SS10477 | 0.012262008 | 0.002666038 | 0 |
| SS10478 | 0.012262008 | 0.002666038 | 0 |
| SS00513 | 0.01023566 | 0.000756907 | 0 |
| SS00256 | 0.011447374 | 0.002004896 | 0 |
| SS00264 | 0.011447374 | 0.002004896 | 0 |
| SS02532 | 0.011447374 | 0.002004896 | 0 |
| SS02535 | 0.011447374 | 0.002004896 | 0 |
| SS08137 | 0.011447374 | 0.002004896 | 0 |
| SS08158 | 0.011447374 | 0.002004896 | 0 |
| SS08167 | 0.011447374 | 0.002004896 | 0 |
| SS08847 | 0.011447374 | 0.002004896 | 0 |
| SS09829 | 0.011447374 | 0.002004896 | 0 |
| SS00295 | 0.010167436 | 0.000742884 | 0 |
| SS02205 | 0.018123926 | 0.008767732 | 0 |

| | | | |
|---|---|---|---|
| SS04555 | 0.018123926 | 0.008767732 | 0 |
| SS09171 | 0.00940452 | 6.50E-05 | 0 |
| SS08617 | 0.010823669 | 0.001505474 | 0 |
| SS05981 | 0.009415336 | 0.000136613 | 0 |
| SS06019 | 0.009415336 | 0.000136613 | 0 |
| SS11103 | 0.009415336 | 0.000136613 | 0 |
| SS05448 | 0.016529067 | 0.00726588 | 0 |
| SS00056 | 0.02541176 | 0.016154827 | 0 |
| SS00798 | 0.02541176 | 0.016154827 | 0 |
| SS04641 | 0.031053901 | 0.021884422 | 0 |
| SS04652 | 0.031053901 | 0.021884422 | 0 |
| SS04482 | 0.011083499 | 0.001916265 | 0 |
| SS04999 | 0.022308332 | 0.013253929 | 0 |
| SS06151 | 0.022308332 | 0.013253929 | 0 |
| SS06181 | 0.022308332 | 0.013253929 | 0 |
| SS09148 | 0.011734204 | 0.002688153 | 0 |
| SS10338 | 0.011734204 | 0.002688153 | 0 |
| SS00263 | 0.021829012 | 0.012789017 | 0 |
| SS04278 | 0.021829012 | 0.012789017 | 0 |
| SS08774 | 0.021829012 | 0.012789017 | 0 |
| SS10498 | 0.021829012 | 0.012789017 | 0 |
| SS10684 | 0.021829012 | 0.012789017 | 0 |
| SS09867 | 0.020262198 | 0.01125453 | 0 |
| SS02674 | 0.01360752 | 0.004641401 | 0 |
| SS00076 | 0.02020135 | 0.011268613 | 0 |
| SS03704 | 0.02020135 | 0.011268613 | 0 |
| SS05810 | 0.02020135 | 0.011268613 | 0 |
| SS08869 | 0.02020135 | 0.011268613 | 0 |
| SS02743 | 0.014208628 | 0.005276205 | 0 |
| SS05223 | 0.014208628 | 0.005276205 | 0 |
| SS14810 | 0.014208628 | 0.005276205 | 0 |
| SS10965 | 0.017348829 | 0.008459349 | 0 |
| SS14858 | 0.017348829 | 0.008459349 | 0 |
| SS00079 | 0.010963041 | 0.002106455 | 0 |
| SS03583 | 0.010963041 | 0.002106455 | 0 |
| SS13489 | 0.010963041 | 0.002106455 | 0 |
| SS06961 | 0.027463181 | 0.018677573 | 0 |
| SS07301 | 0.027463181 | 0.018677573 | 0 |
| SS04271 | 0.024326388 | 0.01558171 | 0 |
| SS08769 | 0.024326388 | 0.01558171 | 0 |
| SS00130 | 0.009469096 | 0.000755063 | 0 |
| SS09150 | 0.009142844 | 0.000450319 | 0 |
| SS13750 | 0.009142844 | 0.000450319 | 0 |
| SS12299 | 0.011039638 | 0.002382667 | 0 |
| SS03269 | 0.009524221 | 0.000870382 | 0 |
| SS07675 | 0.010407159 | 0.00179667 | 0 |
| SS01025 | 0.008921507 | 0.000322053 | 0 |

| | | | |
|---|---|---|---|
| SS00358 | 0.02121945 | 0.012669924 | 0 |
| SS02140 | 0.02121945 | 0.012669924 | 0 |
| SS02699 | 0.02121945 | 0.012669924 | 0 |
| SS10089 | 0.02121945 | 0.012669924 | 0 |
| SS10193 | 0.02121945 | 0.012669924 | 0 |
| SS15450 | 0.02121945 | 0.012669924 | 0 |
| SS05836 | 0.017390327 | 0.008854832 | 0 |
| SS09812 | 0.012730753 | 0.004204169 | 0 |
| SS09269 | 0.010173527 | 0.001654715 | 0 |
| SS10143 | 0.010173527 | 0.001654715 | 0 |
| SS06725 | 0.010073451 | 0.001570628 | 0 |
| SS08108 | 0.029424713 | 0.021046057 | 0 |
| SS09086 | 0.029424713 | 0.021046057 | 0 |
| SS09098 | 0.029424713 | 0.021046057 | 0 |
| SS09080 | 0.0265998 | 0.018259109 | 0 |
| SS14861 | 0.0265998 | 0.018259109 | 0 |
| SS00097 | 0.012560859 | 0.0042687 | 0 |
| SS00859 | 0.01077163 | 0.002496088 | 0 |
| SS03550 | 0.017359886 | 0.009122544 | 0 |
| SS06024 | 0.017359886 | 0.009122544 | 0 |
| SS07900 | 0.011960955 | 0.003723863 | 0 |
| SS00463 | 0.016100422 | 0.007907772 | 0 |
| SS04837 | 0.016100422 | 0.007907772 | 0 |
| SS00487 | 0.012418367 | 0.004255099 | 0 |
| SS02093 | 0.012418367 | 0.004255099 | 0 |
| SS02981 | 0.021964702 | 0.01380811 | 0 |
| SS00039 | 0.008317554 | 0.00016348 | 0 |
| SS01568 | 0.010801796 | 0.00267997 | 0 |
| SS04012 | 0.010801796 | 0.00267997 | 0 |
| SS04333 | 0.015815473 | 0.007722803 | 0 |
| SS13017 | 0.015815473 | 0.007722803 | 0 |
| SS01633 | 0.023355681 | 0.015266877 | 0 |
| SS01649 | 0.023355681 | 0.015266877 | 0 |
| SS15364 | 0.023355681 | 0.015266877 | 0 |
| SS01713 | 0.010890127 | 0.002852454 | 0 |
| SS05848 | 0.010890127 | 0.002852454 | 0 |
| SS13967 | 0.00857166 | 0.000603197 | 0 |
| SS08918 | 0.00795821 | 1.91E-05 | 0 |
| SS07277 | 0.008666001 | 0.00078323 | 0 |
| SS14928 | 0.007928124 | 6.33E-05 | 0 |
| SS01065 | 0.009309055 | 0.001460376 | 0 |
| SS12058 | 0.00819549 | 0.000363039 | 0 |
| SS02139 | 0.008001716 | 0.000177878 | 0 |
| SS04366 | 0.008255019 | 0.000441543 | 0 |
| SS03084 | 0.015535765 | 0.007731772 | 0 |
| SS05394 | 0.015535765 | 0.007731772 | 0 |
| SS05450 | 0.015535765 | 0.007731772 | 0 |

| | | | |
|---|---|---|---|
| SS04982 | 0.016844343 | 0.009051775 | 0 |
| SS06142 | 0.016844343 | 0.009051775 | 0 |
| SS09118 | 0.016844343 | 0.009051775 | 0 |
| SS12981 | 0.016844343 | 0.009051775 | 0 |
| SS03692 | 0.007988376 | 0.000233975 | 0 |
| SS15317 | 0.007988376 | 0.000233975 | 0 |
| SS06716 | 0.015440059 | 0.007732513 | 0 |
| SS01386 | 0.009636031 | 0.001955158 | 0 |
| SS00054 | 0.010734731 | 0.003088786 | 0 |
| SS00810 | 0.010734731 | 0.003088786 | 0 |
| SS01392 | 0.010734731 | 0.003088786 | 0 |
| SS02831 | 0.010734731 | 0.003088786 | 0 |
| SS08809 | 0.008951593 | 0.001315207 | 0 |
| SS03082 | 0.010398385 | 0.002795899 | 0 |
| SS05392 | 0.010398385 | 0.002795899 | 0 |
| SS05438 | 0.010398385 | 0.002795899 | 0 |
| SS14902 | 0.014437822 | 0.006837869 | 0 |
| SS03676 | 0.011947569 | 0.004362374 | 0 |
| SS07084 | 0.011947569 | 0.004362374 | 0 |
| SS07110 | 0.011947569 | 0.004362374 | 0 |
| SS13008 | 0.011947569 | 0.004362374 | 0 |
| SS01988 | 0.015273366 | 0.007740608 | 0 |
| SS00048 | 0.008263473 | 0.000761556 | 0 |
| SS02236 | 0.009275001 | 0.0018274 | 0 |
| SS07279 | 0.008604832 | 0.001214038 | 0 |
| SS05796 | 0.016988166 | 0.009656676 | 0 |
| SS10761 | 0.016988166 | 0.009656676 | 0 |
| SS01224 | 0.010709497 | 0.003389387 | 0 |
| SS05397 | 0.010709497 | 0.003389387 | 0 |
| SS05454 | 0.010709497 | 0.003389387 | 0 |
| SS00757 | 0.007699619 | 0.000409916 | 0 |
| SS09448 | 0.009703853 | 0.002461645 | 0 |
| SS08173 | 0.008732504 | 0.00153741 | 0 |
| SS08852 | 0.008732504 | 0.00153741 | 0 |
| SS11576 | 0.014891873 | 0.007696821 | 0 |
| SS14256 | 0.014891873 | 0.007696821 | 0 |
| SS15452 | 0.014891873 | 0.007696821 | 0 |
| SS02558 | 0.016541845 | 0.009391303 | 0 |
| SS04030 | 0.016541845 | 0.009391303 | 0 |
| SS10672 | 0.016541845 | 0.009391303 | 0 |
| SS11052 | 0.009457363 | 0.002311638 | 0 |
| SS01749 | 0.015845432 | 0.00874671 | 0 |
| SS10862 | 0.015845432 | 0.00874671 | 0 |
| SS06281 | 0.011964717 | 0.004874738 | 0 |
| SS13306 | 0.00779256 | 0.000723846 | 0 |
| SS02327 | 0.018073446 | 0.011009128 | 0 |
| SS02725 | 0.018073446 | 0.011009128 | 0 |

| | | | |
|---|---|---|---|
| SS09159 | 0.018073446 | 0.011009128 | 0 |
| SS08770 | 0.007331581 | 0.000285115 | 0 |
| SS04115 | 0.009069653 | 0.002076397 | 0 |
| SS04457 | 0.009069653 | 0.002076397 | 0 |
| SS06967 | 0.008299489 | 0.001315736 | 0 |
| SS06306 | 0.00813588 | 0.001232554 | 0 |
| SS06410 | 0.00813588 | 0.001232554 | 0 |
| SS12127 | 0.00813588 | 0.001232554 | 0 |
| SS04683 | 0.006906938 | 2.62E-05 | 0 |
| SS08269 | 0.006906938 | 2.62E-05 | 0 |
| SS01793 | 0.007517303 | 0.000646021 | 0 |
| SS06136 | 0.007517303 | 0.000646021 | 0 |
| SS05388 | 0.009197888 | 0.002334815 | 0 |
| SS05420 | 0.009197888 | 0.002334815 | 0 |
| SS07494 | 0.008033924 | 0.00117995 | 0 |
| SS15105 | 0.008033924 | 0.00117995 | 0 |
| SS06322 | 0.008322601 | 0.001469303 | 0 |
| SS06422 | 0.008322601 | 0.001469303 | 0 |
| SS06822 | 0.008322601 | 0.001469303 | 0 |
| SS10548 | 0.008322601 | 0.001469303 | 0 |
| SS12133 | 0.008322601 | 0.001469303 | 0 |
| SS02763 | 0.022841986 | 0.015991352 | 0 |
| SS03573 | 0.022841986 | 0.015991352 | 0 |
| SS09082 | 0.022841986 | 0.015991352 | 0 |
| SS06801 | 0.007431771 | 0.000598041 | 0 |
| SS12091 | 0.007431771 | 0.000598041 | 0 |
| SS12128 | 0.007431771 | 0.000598041 | 0 |
| SS03172 | 0.009459244 | 0.002672755 | 0 |
| SS11060 | 0.009459244 | 0.002672755 | 0 |
| SS06454 | 0.01396855 | 0.007219155 | 0 |
| SS11863 | 0.01396855 | 0.007219155 | 0 |
| SS07281 | 0.008052952 | 0.001336712 | 0 |
| SS04282 | 0.013756951 | 0.007044859 | 0 |
| SS08775 | 0.013756951 | 0.007044859 | 0 |
| SS10687 | 0.013756951 | 0.007044859 | 0 |
| SS04528 | 0.007911699 | 0.001224095 | 0 |
| SS03617 | 0.007307907 | 0.000621708 | 0 |
| SS08625 | 0.007307907 | 0.000621708 | 0 |
| SS10441 | 0.007307907 | 0.000621708 | 0 |
| SS01223 | 0.012786118 | 0.006138003 | 0 |
| SS05396 | 0.012786118 | 0.006138003 | 0 |
| SS05453 | 0.012786118 | 0.006138003 | 0 |
| SS03860 | 0.015845432 | 0.009206003 | 0 |
| SS15485 | 0.008556474 | 0.001941342 | 0 |
| SS11212 | 0.006811233 | 0.000379302 | 0 |
| SS10474 | 0.013311674 | 0.00690078 | 0 |
| SS13472 | 0.007298731 | 0.000910143 | 0 |

| | | | |
|---|---|---|---|
| SS05937 | 0.011273351 | 0.004912336 | 0 |
| SS01243 | 0.007216525 | 0.00087767 | 0 |
| SS05538 | 0.007216525 | 0.00087767 | 0 |
| SS05552 | 0.007216525 | 0.00087767 | 0 |
| SS14881 | 0.012728436 | 0.006393612 | 0 |
| SS00735 | 0.006427171 | 9.91E-05 | 0 |
| SS05654 | 0.015347519 | 0.009033272 | 0 |
| SS07739 | 0.015347519 | 0.009033272 | 0 |
| SS12903 | 0.015347519 | 0.009033272 | 0 |
| SS12916 | 0.015347519 | 0.009033272 | 0 |
| SS12978 | 0.015347519 | 0.009033272 | 0 |
| SS11574 | 0.01137743 | 0.005078505 | 0 |
| SS14250 | 0.01137743 | 0.005078505 | 0 |
| SS14285 | 0.01137743 | 0.005078505 | 0 |
| SS02425 | 0.009161953 | 0.002865485 | 0 |
| SS02238 | 0.01008515 | 0.003796566 | 0 |
| SS13738 | 0.01008515 | 0.003796566 | 0 |
| SS08231 | 0.00777321 | 0.001584375 | 0 |
| SS13864 | 0.00777321 | 0.001584375 | 0 |
| SS01628 | 0.025071364 | 0.018889987 | 0 |
| SS01646 | 0.025071364 | 0.018889987 | 0 |
| SS12365 | 0.025071364 | 0.018889987 | 0 |
| SS04454 | 0.007018106 | 0.000843981 | 0 |
| SS07271 | 0.006367561 | 0.000210288 | 0 |
| SS08938 | 0.015124221 | 0.008980743 | 0 |
| SS15434 | 0.015124221 | 0.008980743 | 0 |
| SS00006 | 0.01703275 | 0.010892506 | 0 |
| SS02709 | 0.01703275 | 0.010892506 | 0 |
| SS09072 | 0.01703275 | 0.010892506 | 0 |
| SS04459 | 0.014757663 | 0.008646733 | 0 |
| SS08937 | 0.014757663 | 0.008646733 | 0 |
| SS13527 | 0.014757663 | 0.008646733 | 0 |
| SS07662 | 0.011604008 | 0.005538113 | 0 |
| SS08913 | 0.011604008 | 0.005538113 | 0 |
| SS08919 | 0.011604008 | 0.005538113 | 0 |
| SS06850 | 0.006323654 | 0.000267278 | 0 |
| SS01437 | 0.007339231 | 0.001324091 | 0 |
| SS11548 | 0.017960593 | 0.011961557 | 0 |
| SS11753 | 0.017960593 | 0.011961557 | 0 |
| SS01803 | 0.007442507 | 0.001472193 | 0 |
| SS10335 | 0.01007906 | 0.004109014 | 0 |
| SS07284 | 0.007540897 | 0.001571918 | 0 |
| SS01022 | 0.01099705 | 0.005139767 | 0 |
| SS08331 | 0.01099705 | 0.005139767 | 0 |
| SS12338 | 0.016185185 | 0.010345342 | 0 |
| SS15296 | 0.007070466 | 0.001237558 | 0 |
| SS08934 | 0.014805539 | 0.008980743 | 0 |

| | | | |
|---|---|---|---|
| SS08944 | 0.014805539 | 0.008980743 | 0 |
| SS06449 | 0.007976918 | 0.002156447 | 0 |
| SS02691 | 0.009625696 | 0.003805619 | 0 |
| SS12123 | 0.006862308 | 0.001051803 | 0 |
| SS06656 | 0.006775698 | 0.001010603 | 0 |
| SS09999 | 0.006775698 | 0.001010603 | 0 |
| SS12830 | 0.007245613 | 0.001502688 | 0 |
| SS02744 | 0.012460106 | 0.006731009 | 0 |
| SS05224 | 0.012460106 | 0.006731009 | 0 |
| SS14811 | 0.012460106 | 0.006731009 | 0 |
| SS05800 | 0.008105955 | 0.002385814 | 0 |
| SS05496 | 0.006501244 | 0.00078345 | 0 |
| SS12125 | 0.006501244 | 0.00078345 | 0 |
| SS03646 | 0.012173069 | 0.006455368 | 0 |
| SS09433 | 0.007609763 | 0.001910903 | 0 |
| SS04502 | 0.009115843 | 0.00344707 | 0 |
| SS14896 | 0.005651317 | 1.39E-05 | 0 |
| SS09600 | 0.007433813 | 0.001886331 | 0 |
| SS07664 | 0.006770812 | 0.001242978 | 0 |
| SS08924 | 0.006770812 | 0.001242978 | 0 |
| SS09146 | 0.006770812 | 0.001242978 | 0 |
| SS09160 | 0.007380374 | 0.001876913 | 0 |
| SS12150 | 0.005815166 | 0.000316529 | 0 |
| SS08329 | 0.01056119 | 0.005066696 | 0 |
| SS10964 | 0.01056119 | 0.005066696 | 0 |
| SS08299 | 0.005893323 | 0.000469299 | 0 |
| SS01219 | 0.00583127 | 0.000411245 | 0 |
| SS03470 | 0.005433467 | 2.67E-05 | 0 |
| SS15489 | 0.005433467 | 2.67E-05 | 0 |
| SS01614 | 0.009132945 | 0.003736627 | 0 |
| SS06685 | 0.009132945 | 0.003736627 | 0 |
| SS06705 | 0.01020445 | 0.004816832 | 0 |
| SS00083 | 0.018772623 | 0.013404417 | 0 |
| SS03585 | 0.018772623 | 0.013404417 | 0 |
| SS13494 | 0.018772623 | 0.013404417 | 0 |
| SS02973 | 0.005684247 | 0.000329428 | 0 |
| SS12428 | 0.005534461 | 0.000185716 | 0 |
| SS05909 | 0.008540772 | 0.003204557 | 0 |
| SS15416 | 0.008540772 | 0.003204557 | 0 |
| SS05381 | 0.006628274 | 0.001305095 | 0 |
| SS06495 | 0.006433743 | 0.001111584 | 0 |
| SS07200 | 0.007861701 | 0.002552247 | 0 |
| SS07224 | 0.007861701 | 0.002552247 | 0 |
| SS02356 | 0.008065374 | 0.002758598 | 0 |
| SS08946 | 0.008065374 | 0.002758598 | 0 |
| SS15480 | 0.008459208 | 0.003162225 | 0 |
| SS03674 | 0.007066819 | 0.001780625 | 0 |

| | | | |
|---|---|---|---|
| SS07082 | 0.007066819 | 0.001780625 | 0 |
| SS14800 | 0.007839312 | 0.002570582 | 0 |
| SS15057 | 0.007839312 | 0.002570582 | 0 |
| SS02381 | 0.006650951 | 0.001389508 | 0 |
| SS02732 | 0.006650951 | 0.001389508 | 0 |
| SS08192 | 0.006650951 | 0.001389508 | 0 |
| SS04911 | 0.005343772 | 0.000107718 | 0 |
| SS08888 | 0.010647238 | 0.005453126 | 0 |
| SS03651 | 0.007095505 | 0.001915389 | 0 |
| SS07063 | 0.007095505 | 0.001915389 | 0 |
| SS11707 | 0.006484257 | 0.00130872 | 0 |
| SS03523 | 0.005430944 | 0.000262485 | 0 |
| SS06071 | 0.00853444 | 0.003371989 | 0 |
| SS08883 | 0.00853444 | 0.003371989 | 0 |
| SS00046 | 0.009959118 | 0.004827434 | 0 |
| SS06127 | 0.009959118 | 0.004827434 | 0 |
| SS09660 | 0.009959118 | 0.004827434 | 0 |
| SS04288 | 0.007353775 | 0.002240439 | 0 |
| SS04321 | 0.007353775 | 0.002240439 | 0 |
| SS00662 | 0.00728499 | 0.002189626 | 0 |
| SS06940 | 0.00728499 | 0.002189626 | 0 |
| SS09952 | 0.00728499 | 0.002189626 | 0 |
| SS00433 | 0.005692299 | 0.000627327 | 0 |
| SS02568 | 0.009525506 | 0.004501393 | 0 |
| SS14187 | 0.009525506 | 0.004501393 | 0 |
| SS14249 | 0.009525506 | 0.004501393 | 0 |
| SS06550 | 0.005845252 | 0.000837015 | 0 |
| SS03531 | 0.011921051 | 0.00691848 | 0 |
| SS11239 | 0.011921051 | 0.00691848 | 0 |
| SS11716 | 0.011921051 | 0.00691848 | 0 |
| SS11762 | 0.011921051 | 0.00691848 | 0 |
| SS00044 | 0.006312677 | 0.001328842 | 0 |
| SS01049 | 0.010644634 | 0.005680282 | 0 |
| SS01310 | 0.010644634 | 0.005680282 | 0 |
| SS03943 | 0.010644634 | 0.005680282 | 0 |
| SS11233 | 0.010644634 | 0.005680282 | 0 |
| SS15453 | 0.010644634 | 0.005680282 | 0 |
| SS15468 | 0.010644634 | 0.005680282 | 0 |
| SS15477 | 0.010644634 | 0.005680282 | 0 |
| SS04515 | 0.005049646 | 9.12E-05 | 0 |
| SS03013 | 0.005177399 | 0.000239276 | 0 |
| SS06828 | 0.005177399 | 0.000239276 | 0 |
| SS12140 | 0.005177399 | 0.000239276 | 0 |
| SS04214 | 0.00636568 | 0.001428895 | 0 |
| SS00963 | 0.005024929 | 0.00014442 | 0 |
| SS06751 | 0.005024929 | 0.00014442 | 0 |
| SS09590 | 0.005376381 | 0.000538925 | 0 |

| | | | |
|---|---|---|---|
| SS00751 | 0.005828666 | 0.001025798 | 0 |
| SS05313 | 0.01097442 | 0.006179762 | 0 |
| SS12348 | 0.004875302 | 8.84E-05 | 0 |
| SS02282 | 0.00517756 | 0.000391234 | 0 |
| SS08332 | 0.00517756 | 0.000391234 | 0 |
| SS09064 | 0.00517756 | 0.000391234 | 0 |
| SS06551 | 0.006758276 | 0.002030223 | 0 |
| SS00316 | 0.004904987 | 0.000219712 | 0 |
| SS07273 | 0.004904987 | 0.000219712 | 0 |
| SS07314 | 0.004904987 | 0.000219712 | 0 |
| SS07236 | 0.010457077 | 0.005771918 | 0 |
| SS07267 | 0.011477299 | 0.00681557 | 0 |
| SS07311 | 0.011477299 | 0.00681557 | 0 |
| SS06754 | 0.008605829 | 0.003954385 | 0 |
| SS11234 | 0.008605829 | 0.003954385 | 0 |
| SS11555 | 0.008605829 | 0.003954385 | 0 |
| SS11757 | 0.008605829 | 0.003954385 | 0 |
| SS14226 | 0.008605829 | 0.003954385 | 0 |
| SS11221 | 0.010378392 | 0.005730976 | 0 |
| SS11244 | 0.010378392 | 0.005730976 | 0 |
| SS08234 | 0.008422034 | 0.003781662 | 0 |
| SS13865 | 0.008422034 | 0.003781662 | 0 |
| SS02260 | 0.004665264 | 5.18E-05 | 0 |
| SS02522 | 0.005289851 | 0.000686139 | 0 |
| SS15427 | 0.010346586 | 0.005747147 | 0 |
| SS00930 | 0.006684604 | 0.002096889 | 0 |
| SS01587 | 0.006684604 | 0.002096889 | 0 |
| SS09141 | 0.006684604 | 0.002096889 | 0 |
| SS14507 | 0.004935153 | 0.000357163 | 0 |
| SS09879 | 0.004670632 | 0.000102506 | 0 |
| SS10955 | 0.0121925 | 0.007669878 | 0 |
| SS00775 | 0.004651523 | 0.000138247 | 0 |
| SS11581 | 0.005256118 | 0.000744558 | 0 |
| SS14484 | 0.005256118 | 0.000744558 | 0 |
| SS00062 | 0.012401655 | 0.007906021 | 0 |
| SS00815 | 0.012401655 | 0.007906021 | 0 |
| SS05673 | 0.006201671 | 0.001758437 | 0 |
| SS09109 | 0.006201671 | 0.001758437 | 0 |
| SS09974 | 0.006201671 | 0.001758437 | 0 |
| SS04865 | 0.005374259 | 0.000931874 | 0 |
| SS07283 | 0.005155847 | 0.000741882 | 0 |
| SS13870 | 0.004553936 | 0.000151464 | 0 |
| SS11573 | 0.009346069 | 0.004956021 | 0 |
| SS11750 | 0.009346069 | 0.004956021 | 0 |
| SS14186 | 0.009346069 | 0.004956021 | 0 |
| SS14248 | 0.009346069 | 0.004956021 | 0 |
| SS00053 | 0.012382626 | 0.007995506 | 0 |

| | | | |
|---|---|---|---|
| SS00808 | 0.012382626 | 0.007995506 | 0 |
| SS01578 | 0.008374239 | 0.004001292 | 0 |
| SS02185 | 0.008374239 | 0.004001292 | 0 |
| SS00804 | 0.00703882 | 0.002734977 | 0 |
| SS12894 | 0.006557447 | 0.002258759 | 0 |
| SS09582 | 0.0060897 | 0.001808427 | 0 |
| SS00876 | 0.006744615 | 0.002501627 | 0 |
| SS11590 | 0.006744615 | 0.002501627 | 0 |
| SS14361 | 0.006744615 | 0.002501627 | 0 |
| SS03597 | 0.007135283 | 0.002945995 | 0 |
| SS00663 | 0.00655669 | 0.002371439 | 0 |
| SS06941 | 0.00655669 | 0.002371439 | 0 |
| SS08246 | 0.006121105 | 0.001946121 | 0 |
| SS00010 | 0.005016555 | 0.000861463 | 0 |
| SS02712 | 0.005016555 | 0.000861463 | 0 |
| SS12543 | 0.005016555 | 0.000861463 | 0 |
| SS09444 | 0.005844208 | 0.001711035 | 0 |
| SS05423 | 0.007949917 | 0.003817406 | 0 |
| SS06819 | 0.00466282 | 0.000546421 | 0 |
| SS13845 | 0.00466282 | 0.000546421 | 0 |
| SS08810 | 0.004335845 | 0.000240548 | 0 |
| SS02976 | 0.00588347 | 0.001790874 | 0 |
| SS04475 | 0.004776672 | 0.000688152 | 0 |
| SS14208 | 0.007722731 | 0.003641565 | 0 |
| SS15476 | 0.007722731 | 0.003641565 | 0 |
| SS02100 | 0.006065946 | 0.002002481 | 0 |
| SS07464 | 0.006065946 | 0.002002481 | 0 |
| SS07470 | 0.006065946 | 0.002002481 | 0 |
| SS15152 | 0.006065946 | 0.002002481 | 0 |
| SS04381 | 0.006579722 | 0.002526135 | 0 |
| SS07196 | 0.006170862 | 0.002122286 | 0 |
| SS07220 | 0.006170862 | 0.002122286 | 0 |
| SS12716 | 0.004709653 | 0.000662985 | 0 |
| SS13286 | 0.004709653 | 0.000662985 | 0 |
| SS08235 | 0.013597071 | 0.009573231 | 0 |
| SS13866 | 0.013597071 | 0.009573231 | 0 |
| SS05949 | 0.009287182 | 0.005283625 | 0 |
| SS08400 | 0.00396268 | 1.49E-05 | 0 |
| SS05390 | 0.010672598 | 0.00673775 | 0 |
| SS05428 | 0.010672598 | 0.00673775 | 0 |
| SS07674 | 0.006415873 | 0.002512208 | 0 |
| SS05434 | 0.005033543 | 0.001155789 | 0 |
| SS01888 | 0.008771158 | 0.004914208 | 0 |
| SS07483 | 0.008771158 | 0.004914208 | 0 |
| SS00047 | 0.005457382 | 0.001626059 | 0 |
| SS09013 | 0.005125842 | 0.001298876 | 0 |
| SS02413 | 0.00396276 | 0.000150664 | 0 |

| | | | |
|---|---|---|---|
| SS06045 | 0.00396276 | 0.000150664 | 0 |
| SS02103 | 0.003859404 | 5.51E-05 | 0 |
| SS12578 | 0.003859404 | 5.51E-05 | 0 |
| SS05391 | 0.005584057 | 0.001789893 | 0 |
| SS05431 | 0.005584057 | 0.001789893 | 0 |
| SS13632 | 0.005127642 | 0.001337754 | 0 |
| SS05393 | 0.007529358 | 0.003751082 | 0 |
| SS05449 | 0.007529358 | 0.003751082 | 0 |
| SS02985 | 0.004126449 | 0.000372091 | 0 |
| SS12881 | 0.003971535 | 0.000239962 | 0 |
| SS03042 | 0.004957267 | 0.001227977 | 0 |
| SS09469 | 0.004957267 | 0.001227977 | 0 |
| SS08959 | 0.006342362 | 0.002613552 | 0 |
| SS10283 | 0.006342362 | 0.002613552 | 0 |
| SS03003 | 0.0104209 | 0.006740611 | 0 |
| SS03514 | 0.0104209 | 0.006740611 | 0 |
| SS05503 | 0.0104209 | 0.006740611 | 0 |
| SS09967 | 0.0104209 | 0.006740611 | 0 |
| SS15360 | 0.0104209 | 0.006740611 | 0 |
| SS15426 | 0.0104209 | 0.006740611 | 0 |
| SS06722 | 0.007858582 | 0.00419011 | 0 |
| SS13021 | 0.007858582 | 0.00419011 | 0 |
| SS06962 | 0.006150595 | 0.002510107 | 0 |
| SS07265 | 0.006150595 | 0.002510107 | 0 |
| SS07289 | 0.006150595 | 0.002510107 | 0 |
| SS01252 | 0.003867697 | 0.000254269 | 0 |
| SS02234 | 0.003867697 | 0.000254269 | 0 |
| SS13418 | 0.005016636 | 0.001406976 | 0 |
| SS13428 | 0.005016636 | 0.001406976 | 0 |
| SS12828 | 0.0072388 | 0.003647591 | 0 |
| SS10511 | 0.009208223 | 0.005664189 | 0 |
| SS09858 | 0.004723474 | 0.001179967 | 0 |
| SS12209 | 0.003805002 | 0.000293617 | 0 |
| SS05628 | 0.005436472 | 0.001962333 | 0 |
| SS05589 | 0.004251839 | 0.000780779 | 0 |
| SS02259 | 0.003567882 | 0.000170029 | 0 |
| SS11250 | 0.003567882 | 0.000170029 | 0 |
| SS11913 | 0.004586063 | 0.001191418 | 0 |
| SS15527 | 0.00501183 | 0.001636683 | 0 |
| SS12582 | 0.003489084 | 0.000136455 | 0 |
| SS00828 | 0.003722877 | 0.000381196 | 0 |
| SS07272 | 0.006210606 | 0.002884789 | 0 |
| SS07313 | 0.006210606 | 0.002884789 | 0 |
| SS00947 | 0.008031802 | 0.00472441 | 0 |
| SS08794 | 0.005591948 | 0.002306874 | 0 |
| SS09605 | 0.003663508 | 0.000409832 | 0 |
| SS00177 | 0.003399227 | 0.000154519 | 0 |

| | | | |
|---|---|---|---|
| SS01778 | 0.003399227 | 0.000154519 | 0 |
| SS05275 | 0.004131656 | 0.000887331 | 0 |
| SS13581 | 0.004109542 | 0.000938545 | 0 |
| SS06025 | 0.005547318 | 0.002415611 | 0 |
| SS05870 | 0.005906662 | 0.002780277 | 0 |
| SS06540 | 0.005906662 | 0.002780277 | 0 |
| SS01309 | 0.003145683 | 6.93E-05 | 0 |
| SS03942 | 0.003145683 | 6.93E-05 | 0 |
| SS10824 | 0.003145683 | 6.93E-05 | 0 |
| SS08308 | 0.003856078 | 0.000800487 | 0 |
| SS03762 | 0.003055827 | 1.08E-05 | 0 |
| SS13318 | 0.005488753 | 0.002460917 | 0 |
| SS04168 | 0.003210821 | 0.000249333 | 0 |
| SS03982 | 0.002976868 | 1.87E-05 | 0 |
| SS05385 | 0.006661205 | 0.003707279 | 0 |
| SS06737 | 0.003418337 | 0.000466049 | 0 |
| SS08306 | 0.004019927 | 0.001070069 | 0 |
| SS01068 | 0.003116079 | 0.000205646 | 0 |
| SS04643 | 0.003388331 | 0.000478089 | 0 |
| SS06660 | 0.003388331 | 0.000478089 | 0 |
| SS12161 | 0.002968735 | 7.71E-05 | 0 |
| SS05963 | 0.002881563 | 2.40E-05 | 0 |
| SS13395 | 0.002865138 | 1.09E-05 | 0 |
| SS06687 | 0.003480389 | 0.000630678 | 0 |
| SS04269 | 0.002990368 | 0.000150545 | 0 |
| SS06867 | 0.003718231 | 0.000894373 | 0 |
| SS10294 | 0.003718231 | 0.000894373 | 0 |
| SS14207 | 0.003718231 | 0.000894373 | 0 |
| SS06511 | 0.005751829 | 0.002930752 | 0 |
| SS10415 | 0.005751829 | 0.002930752 | 0 |
| SS10428 | 0.005751829 | 0.002930752 | 0 |
| SS15425 | 0.003774595 | 0.001009492 | 0 |
| SS00163 | 0.003748317 | 0.00098371 | 0 |
| SS07684 | 0.003205453 | 0.000442409 | 0 |
| SS12417 | 0.002851477 | 9.42E-05 | 0 |
| SS01785 | 0.003063879 | 0.000320641 | 0 |
| SS08310 | 0.0035386 | 0.000832072 | 0 |
| SS08428 | 0.005003537 | 0.002340978 | 0 |
| SS03298 | 0.006522716 | 0.003872453 | 0 |
| SS04413 | 0.007026238 | 0.004383681 | 0 |
| SS00320 | 0.002963528 | 0.000343397 | 0 |
| SS09916 | 0.006373652 | 0.0037658 | 0 |
| SS09589 | 0.002827322 | 0.000251129 | 0 |
| SS02643 | 0.005215698 | 0.002640069 | 0 |
| SS01615 | 0.003026303 | 0.000468189 | 0 |
| SS06879 | 0.003026303 | 0.000468189 | 0 |
| SS00439 | 0.00308355 | 0.000539394 | 0 |

| | | | |
|---|---|---|---|
| SS01201 | 0.004240701 | 0.001702615 | 0 |
| SS07029 | 0.002570933 | 4.95E-05 | 0 |
| SS09926 | 0.00494629 | 0.002444759 | 0 |
| SS12185 | 0.002707139 | 0.00020808 | 0 |
| SS09207 | 0.00301777 | 0.000518876 | 0 |
| SS07205 | 0.003398264 | 0.000914113 | 0 |
| SS15262 | 0.002955396 | 0.000511382 | 0 |
| SS00011 | 0.00247018 | 2.68E-05 | 0 |
| SS02329 | 0.004093358 | 0.001657915 | 0 |
| SS06891 | 0.003374831 | 0.000962379 | 0 |
| SS08680 | 0.003374831 | 0.000962379 | 0 |
| SS00246 | 0.002500186 | 0.000102801 | 0 |
| SS01138 | 0.002426835 | 5.31E-05 | 0 |
| SS12488 | 0.002394065 | 2.81E-05 | 0 |
| SS12486 | 0.002385853 | 2.35E-05 | 0 |
| SS12522 | 0.002385853 | 2.35E-05 | 0 |
| SS14586 | 0.003753284 | 0.001391244 | 0 |
| SS14677 | 0.003753284 | 0.001391244 | 0 |
| SS02278 | 0.004254523 | 0.001903984 | 0 |
| SS02295 | 0.002598174 | 0.000248471 | 0 |
| SS14959 | 0.002598174 | 0.000248471 | 0 |
| SS00457 | 0.003616836 | 0.001311994 | 0 |
| SS12888 | 0.005294175 | 0.003018694 | 0 |
| SS02730 | 0.00336858 | 0.001131236 | 0 |
| SS00464 | 0.002963769 | 0.000739566 | 0 |
| SS10280 | 0.005232524 | 0.003020098 | 0 |
| SS06489 | 0.002685828 | 0.000478089 | 0 |
| SS07886 | 0.005389319 | 0.003214947 | 0 |
| SS14974 | 0.003736377 | 0.001564689 | 0 |
| SS12320 | 0.005294175 | 0.003126741 | 0 |
| SS02288 | 0.003756048 | 0.001591448 | 0 |
| SS02480 | 0.003756048 | 0.001591448 | 0 |
| SS12506 | 0.002235986 | 9.60E-05 | 0 |
| SS13745 | 0.002235986 | 9.60E-05 | 0 |
| SS05150 | 0.003121688 | 0.000987904 | 0 |
| SS06655 | 0.002606708 | 0.000474265 | 0 |
| SS09998 | 0.002606708 | 0.000474265 | 0 |
| SS00965 | 0.003660824 | 0.001533212 | 0 |
| SS06208 | 0.003660824 | 0.001533212 | 0 |
| SS07122 | 0.003660824 | 0.001533212 | 0 |
| SS09014 | 0.002135313 | 1.86E-05 | 0 |
| SS00113 | 0.003139592 | 0.001023811 | 0 |
| SS08069 | 0.003210901 | 0.001131435 | 0 |
| SS07826 | 0.003211464 | 0.001138466 | 0 |
| SS09414 | 0.003211464 | 0.001138466 | 0 |
| SS14002 | 0.003085431 | 0.001060297 | 0 |
| SS00835 | 0.003739141 | 0.001714999 | 0 |

| | | | |
|---|---|---|---|
| SS03500 | 0.003739141 | 0.001714999 | 0 |
| SS05461 | 0.003418417 | 0.001401856 | 0 |
| SS07332 | 0.002187112 | 0.000180449 | 0 |
| SS10365 | 0.002047981 | 4.79E-05 | 0 |
| SS02991 | 0.002669162 | 0.00068191 | 0 |
| SS05818 | 0.002669162 | 0.00068191 | 0 |
| SS08568 | 0.002669162 | 0.00068191 | 0 |
| SS11860 | 0.002669162 | 0.00068191 | 0 |
| SS09293 | 0.005079331 | 0.003095229 | 0 |
| SS10227 | 0.005079331 | 0.003095229 | 0 |
| SS08589 | 0.002974184 | 0.001029797 | 0 |
| SS13983 | 0.002974184 | 0.001029797 | 0 |
| SS04473 | 0.003600251 | 0.001659719 | 0 |
| SS01595 | 0.004499935 | 0.002588911 | 0 |
| SS02190 | 0.004499935 | 0.002588911 | 0 |
| SS05435 | 0.002127101 | 0.000236921 | 0 |
| SS04318 | 0.002807651 | 0.000940353 | 0 |
| SS13948 | 0.001868349 | 2.39E-06 | 0 |
| SS02080 | 0.00235064 | 0.00051053 | 0 |
| SS05152 | 0.003787854 | 0.001964127 | 0 |
| SS05262 | 0.003787854 | 0.001964127 | 0 |
| SS12779 | 0.003787854 | 0.001964127 | 0 |
| SS10766 | 0.002518973 | 0.000722827 | 0 |
| SS07054 | 0.00180297 | 1.71E-05 | 0 |
| SS14592 | 0.002072779 | 0.000298134 | 0 |
| SS14686 | 0.002072779 | 0.000298134 | 0 |
| SS14046 | 0.001805734 | 5.24E-05 | 0 |
| SS11713 | 0.00299101 | 0.00124952 | 0 |
| SS07208 | 0.001748487 | 2.60E-05 | 0 |
| SS03995 | 0.003660342 | 0.001939112 | 0 |
| SS12375 | 0.001773045 | 7.59E-05 | 0 |
| SS14497 | 0.001985527 | 0.000296328 | 0 |
| SS02695 | 0.001925515 | 0.000242729 | 0 |
| SS02333 | 0.002352922 | 0.000698489 | 0 |
| SS12447 | 0.001694005 | 4.83E-05 | 0 |
| SS07498 | 0.002393744 | 0.000758771 | 0 |
| SS09009 | 0.001852085 | 0.000235459 | 0 |
| SS09865 | 0.003109908 | 0.001503988 | 0 |
| SS09331 | 0.001985125 | 0.00040011 | 0 |
| SS06786 | 0.001718321 | 0.000147383 | 0 |
| SS02773 | 0.004155732 | 0.002596495 | 0 |
| SS06256 | 0.004155732 | 0.002596495 | 0 |
| SS12420 | 0.001522425 | 8.61E-06 | 0 |
| SS04925 | 0.002938971 | 0.00142549 | 0 |
| SS10083 | 0.002222084 | 0.000720996 | 0 |
| SS10070 | 0.002222084 | 0.000723875 | 0 |
| SS03009 | 0.001604149 | 0.000111306 | 0 |

| | | | |
|---|---|---|---|
| SS06826 | 0.001604149 | 0.000111306 | 0 |
| SS08411 | 0.001604149 | 0.000111306 | 0 |
| SS12139 | 0.001604149 | 0.000111306 | 0 |
| SS02216 | 0.001713355 | 0.000245526 | 0 |
| SS04022 | 0.001544298 | 0.000112088 | 0 |
| SS02666 | 0.001772884 | 0.000355756 | 0 |
| SS13354 | 0.001884533 | 0.000502954 | 0 |
| SS13421 | 0.001884533 | 0.000502954 | 0 |
| SS04471 | 0.001448914 | 9.47E-05 | 0 |
| SS10863 | 0.001405328 | 5.35E-05 | 0 |
| SS02584 | 0.001378247 | 3.87E-05 | 0 |
| SS05417 | 0.0024836 | 0.00115112 | 0 |
| SS05469 | 0.0024836 | 0.00115112 | 0 |
| SS06864 | 0.001571299 | 0.000243352 | 0 |
| SS01072 | 0.001514132 | 0.000188276 | 0 |
| SS01472 | 0.001337265 | 3.54E-05 | 0 |
| SS06359 | 0.002377319 | 0.001117338 | 0 |
| SS09216 | 0.002097176 | 0.000852481 | 0 |
| SS10928 | 0.002097176 | 0.000852481 | 0 |
| SS15203 | 0.00166408 | 0.000452188 | 0 |
| SS07936 | 0.00195512 | 0.000777288 | 0 |
| SS07937 | 0.00195512 | 0.000777288 | 0 |
| SS13605 | 0.001942181 | 0.000794855 | 0 |
| SS09528 | 0.001351086 | 0.000260146 | 0 |
| SS13756 | 0.001351086 | 0.000260146 | 0 |
| SS01840 | 0.001904044 | 0.000830523 | 0 |
| SS14591 | 0.001255621 | 0.000239121 | 0 |
| SS14685 | 0.001255621 | 0.000239121 | 0 |
| SS04428 | 0.001165846 | 0.000185656 | 0 |
| SS08415 | 0.001260748 | 0.0002887 | 0 |
| SS14465 | 0.001260748 | 0.0002887 | 0 |
| SS01378 | 0.001171133 | 0.000217125 | 0 |
| SS01092 | 0.001062249 | 0.000113132 | 0 |
| SS12233 | 0.001258145 | 0.000320323 | 0 |
| SS09122 | 0.001059404 | 0.000128168 | 0 |
| SS02903 | 0.001007605 | 0.000116942 | 0 |
| SS02758 | 0.001124703 | 0.000249072 | 0 |
| SS03569 | 0.001124703 | 0.000249072 | 0 |
| SS10173 | 0.001124703 | 0.000249072 | 0 |
| SS12372 | 0.00112234 | 0.0002485 | 0 |
| SS00146 | 0.001239437 | 0.000387119 | 0 |
| SS00151 | 0.001239437 | 0.000387119 | 0 |
| SS04426 | 0.000966703 | 0.000152106 | 0 |
| SS06302 | 0.000966703 | 0.000152106 | 0 |
| SS04023 | 0.000841715 | 3.23E-05 | 0 |
| SS12401 | 0.001457207 | 0.0006693 | 0 |
| SS12228 | 0.000882456 | 0.000147977 | 0 |

| | | | |
|---|---|---|---|
| SS00306 | 0.001053956 | 0.000329223 | 0 |
| SS09856 | 0.000762514 | 5.35E-05 | 0 |
| SS12416 | 0.000912462 | 0.000228921 | 0 |
| SS04470 | 0.000710796 | 3.75E-05 | 0 |
| SS03626 | 0.000819922 | 0.000199015 | 0 |
| SS01076 | 0.000855215 | 0.000237886 | 0 |
| SS02977 | 0.00066729 | 6.29E-05 | 0 |
| SS05278 | 0.000661762 | 7.11E-05 | 0 |
| SS13888 | 0.000623704 | 5.16E-05 | 0 |
| SS11543 | 0.001013134 | 0.000458495 | 0 |
| SS13836 | 0.001013134 | 0.000458495 | 0 |
| SS08209 | 0.000789916 | 0.000240441 | 0 |
| SS05328 | 0.000561009 | 1.90E-05 | 0 |
| SS10571 | 0.000798048 | 0.00026945 | 0 |
| SS09120 | 0.000621101 | 9.99E-05 | 0 |
| SS12202 | 0.000552957 | 3.23E-05 | 0 |
| SS02857 | 0.000762675 | 0.000288224 | 0 |
| SS10901 | 0.000522872 | 5.28E-05 | 0 |
| SS13886 | 0.000492946 | 2.84E-05 | 0 |
| SS06046 | 0.000468469 | 8.85E-06 | 0 |
| SS07201 | 0.000531084 | 8.11E-05 | 0 |
| SS07509 | 0.000531084 | 8.11E-05 | 0 |
| SS12210 | 0.000490343 | 4.35E-05 | 0 |
| SS02246 | 0.000536452 | 0.000103661 | 0 |
| SS04054 | 0.000536452 | 0.000103661 | 0 |
| SS12208 | 0.000490343 | 6.13E-05 | 0 |
| SS10522 | 0.000512055 | 8.31E-05 | 0 |
| SS11106 | 0.000443912 | 4.31E-05 | 0 |
| SS07209 | 0.000599147 | 0.000200688 | 0 |
| SS12406 | 0.000408539 | 1.26E-05 | 0 |
| SS12497 | 0.000386665 | 2.22E-05 | 0 |
| SS04218 | 0.000498314 | 0.000135388 | 0 |
| SS01557 | 0.000405855 | 5.27E-05 | 0 |
| SS09220 | 0.000381297 | 2.95E-05 | 0 |
| SS03813 | 0.000468389 | 0.000118458 | 0 |
| SS07115 | 0.00034316 | 1.06E-06 | 0 |
| SS03129 | 0.000351372 | 1.34E-05 | 0 |
| SS11666 | 0.000419435 | 8.74E-05 | 0 |
| SS09921 | 0.00035682 | 2.74E-05 | 0 |
| SS05535 | 0.000340476 | 2.85E-05 | 0 |
| SS05548 | 0.000340476 | 2.85E-05 | 0 |
| SS05279 | 0.000315918 | 2.07E-05 | 0 |
| SS04881 | 0.000302338 | 1.32E-05 | 0 |
| SS10523 | 0.000321367 | 3.77E-05 | 0 |
| SS11701 | 0.00029689 | 2.67E-05 | 0 |
| SS03273 | 0.000332343 | 7.10E-05 | 0 |
| SS04671 | 0.000343079 | 8.19E-05 | 0 |

| | | | |
|---|---|---|---|
| SS04425 | 0.000332183 | 7.95E-05 | 0 |
| SS06301 | 0.000332183 | 7.95E-05 | 0 |
| SS03649 | 0.000288677 | 5.43E-05 | 0 |
| SS07062 | 0.000288677 | 5.43E-05 | 0 |
| SS00292 | 0.000231511 | 2.85E-06 | 0 |
| SS12512 | 0.000253304 | 2.88E-05 | 0 |
| SS14190 | 0.000253223 | 4.09E-05 | 0 |
| SS02498 | 0.000215166 | 3.08E-06 | 0 |
| SS04295 | 0.000209718 | 5.01E-06 | 0 |
| SS01996 | 0.000204269 | 4.50E-06 | 0 |
| SS03162 | 0.000204269 | 4.50E-06 | 0 |
| SS10537 | 0.000193373 | 4.37E-07 | 0 |
| SS11657 | 0.000247855 | 5.67E-05 | 0 |
| SS05085 | 0.000256068 | 6.71E-05 | 0 |
| SS12245 | 0.000220614 | 4.08E-05 | 0 |
| SS05078 | 0.00017158 | 9.31E-06 | 0 |
| SS13783 | 0.00017158 | 3.13E-05 | 0 |
| SS07320 | 0.000179712 | 4.28E-05 | 0 |
| SS09317 | 0.00013889 | 1.32E-05 | 0 |
| SS14540 | 9.53E-05 | 6.55E-06 | 0 |
| SS09226 | 9.26E-05 | 4.41E-06 | 0 |
| SS08608 | 9.26E-05 | 7.60E-06 | 0 |
| SS10243 | 9.53E-05 | 1.41E-05 | 0 |
| SS03442 | 8.72E-05 | 6.12E-06 | 0 |
| SS04182 | 8.72E-05 | 6.12E-06 | 0 |
| SS14859 | 6.54E-05 | 1.43E-05 | 0 |
| SS11654 | 4.36E-05 | 5.77E-06 | 0 |
| SS04479 | 3.81E-05 | 2.40E-06 | 0 |
| SS05135 | 3.81E-05 | 4.03E-06 | 0 |
| SS08767 | 3.81E-05 | 4.03E-06 | 0 |
| SS14844 | 3.81E-05 | 4.03E-06 | 0 |
| SS02010 | 3.81E-05 | 4.32E-06 | 0 |
| SS04735 | 2.72E-05 | 2.73E-06 | 0 |
| SS10263 | 2.72E-05 | 4.16E-06 | 0 |
| SS11513 | 1.63E-05 | 1.63E-06 | 0 |
| SS11674 | 1.63E-05 | 1.76E-06 | 0 |
| SS11686 | 1.63E-05 | 1.76E-06 | 0 |
| SS11699 | 1.63E-05 | 1.76E-06 | 0 |
| SS10315 | 1.09E-05 | 7.43E-07 | 0 |
| SS11631 | 1.09E-05 | 8.24E-07 | 0 |
| SS10230 | 1.09E-05 | 1.05E-06 | 0 |
| SS14839 | 1.09E-05 | 1.24E-06 | 0 |
| SS11489 | 1.09E-05 | 1.24E-06 | 0 |
| SS10258 | 5.45E-06 | 1.87E-07 | 0 |
| SS13540 | 5.45E-06 | 2.61E-07 | 0 |
| SS13456 | 5.45E-06 | 3.04E-07 | 0 |
| SS14379 | 5.45E-06 | 3.37E-07 | 0 |

| | | | |
|---|---|---|---|
| SS05104 | 5.45E-06 | 3.51E-07 | 0 |
| SS08737 | 5.45E-06 | 3.51E-07 | 0 |
| SS11880 | 5.45E-06 | 4.19E-07 | 0 |
| SS11383 | 5.45E-06 | 4.83E-07 | 0 |
| SS11663 | 5.45E-06 | 5.61E-07 | 0 |
| SS08988 | 5.45E-06 | 5.77E-07 | 0 |
| SS11211 | 0.002069177 | 0.093844573 | 1.56E-58 |
| SS11916 | 0.000866433 | 0.111700178 | 5.87E-55 |
| SS05674 | 0.000855536 | 0.10448766 | 1.34E-54 |
| SS10163 | 0.000855536 | 0.10448766 | 1.34E-54 |
| SS11944 | 0.000855536 | 0.10448766 | 1.34E-54 |
| SS01714 | 2.68E-06 | 0.027467228 | 7.66E-54 |
| SS04599 | 0.007437908 | 0.127571444 | 1.07E-53 |
| SS04627 | 0.007437908 | 0.127571444 | 1.07E-53 |
| SS04601 | 0.001958606 | 0.033607471 | 1.20E-53 |
| SS08698 | 0.001958606 | 0.033607471 | 1.20E-53 |
| SS06117 | 0.000727703 | 0.016325221 | 1.49E-53 |
| SS09648 | 0.000727703 | 0.016325221 | 1.49E-53 |
| SS06159 | 0.008114087 | 0.071986934 | 2.09E-53 |
| SS06184 | 0.008114087 | 0.071986934 | 2.09E-53 |
| SS10786 | 0.008114087 | 0.071986934 | 2.09E-53 |
| SS11110 | 0.000141895 | 0.048705619 | 2.17E-52 |
| SS09564 | 0 | 0.062800705 | 2.61E-52 |
| SS14989 | 5.45E-06 | 0.011450974 | 6.09E-52 |
| SS00928 | 0.000933612 | 0.013481865 | 1.08E-51 |
| SS04692 | 0.000915868 | 0.016971706 | 2.23E-51 |
| SS13773 | 0.001705624 | 0.037634552 | 2.80E-51 |
| SS13777 | 0.001705624 | 0.037634552 | 2.80E-51 |
| SS02625 | 0.001326884 | 0.03508151 | 4.58E-51 |
| SS00941 | 0.018326348 | 0.055288788 | 5.80E-51 |
| SS14011 | 0.018326348 | 0.055288788 | 5.80E-51 |
| SS00944 | 0.018370015 | 0.055359574 | 6.97E-51 |
| SS14031 | 0.018370015 | 0.055359574 | 6.97E-51 |
| SS01726 | 0.012603929 | 0.042596508 | 1.88E-50 |
| SS04194 | 0.012603929 | 0.042596508 | 1.88E-50 |
| SS06913 | 0.012603929 | 0.042596508 | 1.88E-50 |
| SS07180 | 0.012603929 | 0.042596508 | 1.88E-50 |
| SS15076 | 0.012603929 | 0.042596508 | 1.88E-50 |
| SS08626 | 0.007089575 | 0.057685106 | 8.07E-50 |
| SS14990 | 0 | 0.00985422 | 9.35E-50 |
| SS00204 | 0.001574625 | 0.021415089 | 9.85E-50 |
| SS00570 | 0.001574625 | 0.021415089 | 9.85E-50 |
| SS01731 | 0.001574625 | 0.021415089 | 9.85E-50 |
| SS02667 | 0.017246631 | 0.053846638 | 3.29E-49 |
| SS03534 | 0.017246631 | 0.053846638 | 3.29E-49 |
| SS10693 | 0.017246631 | 0.053846638 | 3.29E-49 |
| SS02420 | 0.008184192 | 0.05180438 | 5.36E-49 |

| | | | |
|---|---|---|---|
| SS05597 | 0 | 0.008682516 | 5.59E-49 |
| SS15228 | 0.008233949 | 0.032577543 | 6.63E-49 |
| SS04985 | 0.002700567 | 0.024755468 | 9.24E-49 |
| SS14014 | 0.009531872 | 0.04227121 | 1.69E-48 |
| SS05906 | 2.99E-05 | 0.020119231 | 4.78E-48 |
| SS05976 | 0 | 0.017623586 | 1.12E-47 |
| SS09666 | 0 | 0.017623586 | 1.12E-47 |
| SS11958 | 0 | 0.017623586 | 1.12E-47 |
| SS02020 | 0.001662359 | 0.031652295 | 3.52E-47 |
| SS10341 | 0.001662359 | 0.031652295 | 3.52E-47 |
| SS05895 | 0.000500918 | 0.021048372 | 3.99E-47 |
| SS01640 | 0.094999851 | 0.23297907 | 5.01E-47 |
| SS05338 | 0.094999851 | 0.23297907 | 5.01E-47 |
| SS05349 | 0.094999851 | 0.23297907 | 5.01E-47 |
| SS08726 | 0.094999851 | 0.23297907 | 5.01E-47 |
| SS01737 | 0.007351412 | 0.032939298 | 8.87E-47 |
| SS06506 | 0.007351412 | 0.032939298 | 8.87E-47 |
| SS07190 | 0.007351412 | 0.032939298 | 8.87E-47 |
| SS11939 | 0.007351412 | 0.032939298 | 8.87E-47 |
| SS05596 | 0 | 0.015353234 | 1.02E-46 |
| SS03343 | 0.002228656 | 0.013766115 | 1.33E-46 |
| SS05670 | 0.002228656 | 0.013766115 | 1.33E-46 |
| SS06001 | 0.002228656 | 0.013766115 | 1.33E-46 |
| SS10092 | 0.002228656 | 0.013766115 | 1.33E-46 |
| SS04291 | 0.000882777 | 0.009796222 | 1.60E-46 |
| SS04979 | 0.000882777 | 0.009796222 | 1.60E-46 |
| SS05968 | 0.000882777 | 0.009796222 | 1.60E-46 |
| SS12966 | 0.000882777 | 0.009796222 | 1.60E-46 |
| SS08258 | 0.001495746 | 0.038113664 | 1.61E-46 |
| SS02465 | 0.016090489 | 0.067974567 | 2.43E-46 |
| SS04114 | 0.01166254 | 0.033305077 | 2.78E-46 |
| SS06676 | 0.01166254 | 0.033305077 | 2.78E-46 |
| SS08858 | 0.01166254 | 0.033305077 | 2.78E-46 |
| SS02045 | 0.000935941 | 0.02194034 | 3.14E-46 |
| SS02534 | 0.000935941 | 0.02194034 | 3.14E-46 |
| SS03641 | 0.000935941 | 0.02194034 | 3.14E-46 |
| SS10347 | 0.000935941 | 0.02194034 | 3.14E-46 |
| SS01734 | 0.006516866 | 0.022024917 | 1.01E-45 |
| SS06504 | 0.006516866 | 0.022024917 | 1.01E-45 |
| SS07185 | 0.006516866 | 0.022024917 | 1.01E-45 |
| SS13929 | 0.003827517 | 0.027614824 | 1.19E-45 |
| SS02463 | 0.017692436 | 0.08706687 | 1.34E-45 |
| SS13730 | 0.017692436 | 0.08706687 | 1.34E-45 |
| SS13748 | 2.76E-06 | 0.011409365 | 1.79E-45 |
| SS08113 | 0.009794156 | 0.067620191 | 3.47E-45 |
| SS08148 | 0.009794156 | 0.067620191 | 3.47E-45 |
| SS04304 | 0.000470993 | 0.007972454 | 3.57E-45 |

| | | | |
|---|---|---|---|
| SS01281 | 0.002090603 | 0.032757562 | 4.10E-45 |
| SS04600 | 0.004037877 | 0.041470039 | 7.62E-45 |
| SS08697 | 0.004037877 | 0.041470039 | 7.62E-45 |
| SS02213 | 0.011823464 | 0.032129032 | 1.24E-44 |
| SS03610 | 0.007995946 | 0.019067812 | 1.38E-44 |
| SS06512 | 0.007995946 | 0.019067812 | 1.38E-44 |
| SS10416 | 0.007995946 | 0.019067812 | 1.38E-44 |
| SS10435 | 0.007995946 | 0.019067812 | 1.38E-44 |
| SS11811 | 0.007995946 | 0.019067812 | 1.38E-44 |
| SS07688 | 0.003603256 | 0.020817086 | 1.47E-44 |
| SS13923 | 0.003603256 | 0.020817086 | 1.47E-44 |
| SS02207 | 0.011809723 | 0.031842069 | 1.53E-44 |
| SS00537 | 0.005713656 | 0.031266296 | 1.68E-44 |
| SS04815 | 0.000166132 | 0.011428451 | 2.30E-44 |
| SS01956 | 2.76E-06 | 0.0063795 | 2.83E-44 |
| SS03285 | 2.76E-06 | 0.0063795 | 2.83E-44 |
| SS11869 | 8.13E-06 | 0.005669358 | 2.90E-44 |
| SS06115 | 0.000182396 | 0.008293225 | 3.63E-44 |
| SS09631 | 0.000182396 | 0.008293225 | 3.63E-44 |
| SS08257 | 0.000353815 | 0.01139624 | 5.52E-44 |
| SS00103 | 0.01316425 | 0.068645547 | 6.17E-44 |
| SS07909 | 0 | 0.01100097 | 8.92E-44 |
| SS00194 | 0.002407485 | 0.013562899 | 9.32E-44 |
| SS09909 | 0.002407485 | 0.013562899 | 9.32E-44 |
| SS11905 | 0.002407485 | 0.013562899 | 9.32E-44 |
| SS01407 | 2.68E-06 | 0.013033508 | 1.63E-43 |
| SS11542 | 0.001708629 | 0.022213773 | 2.06E-43 |
| SS14173 | 0.001708629 | 0.022213773 | 2.06E-43 |
| SS01889 | 0 | 0.002962563 | 2.35E-43 |
| SS01917 | 0 | 0.002962563 | 2.35E-43 |
| SS06688 | 2.76E-06 | 0.00875134 | 3.50E-43 |
| SS03479 | 1.63E-05 | 0.007758994 | 4.83E-43 |
| SS04028 | 1.63E-05 | 0.007758994 | 4.83E-43 |
| SS05326 | 0.007595666 | 0.029703861 | 6.37E-43 |
| SS08349 | 0.007595666 | 0.029703861 | 6.37E-43 |
| SS07912 | 5.45E-06 | 0.012156571 | 8.78E-43 |
| SS08256 | 0.000544504 | 0.015442682 | 1.45E-42 |
| SS10509 | 0.000544504 | 0.015442682 | 1.45E-42 |
| SS00106 | 0.017482752 | 0.06330441 | 1.59E-42 |
| SS07693 | 0.002360699 | 0.051735816 | 5.04E-42 |
| SS13926 | 0.002360699 | 0.051735816 | 5.04E-42 |
| SS07004 | 0.001996744 | 0.009372895 | 6.78E-42 |
| SS10675 | 0.007582682 | 0.018471931 | 9.33E-42 |
| SS00942 | 0.017660905 | 0.046514638 | 1.11E-41 |
| SS14012 | 0.017660905 | 0.046514638 | 1.11E-41 |
| SS00945 | 0.017726284 | 0.046525569 | 1.28E-41 |
| SS14032 | 0.017726284 | 0.046525569 | 1.28E-41 |

| | | | |
|---|---|---|---|
| SS06015 | 0.008896709 | 0.024173131 | 1.67E-41 |
| SS08186 | 0.008896709 | 0.024173131 | 1.67E-41 |
| SS00070 | 0.038752498 | 0.109948776 | 1.94E-41 |
| SS00174 | 0.038752498 | 0.109948776 | 1.94E-41 |
| SS00201 | 0.038752498 | 0.109948776 | 1.94E-41 |
| SS00558 | 0.038752498 | 0.109948776 | 1.94E-41 |
| SS01776 | 0.038752498 | 0.109948776 | 1.94E-41 |
| SS05335 | 0.038752498 | 0.109948776 | 1.94E-41 |
| SS06903 | 0.038752498 | 0.109948776 | 1.94E-41 |
| SS06916 | 0.038752498 | 0.109948776 | 1.94E-41 |
| SS07636 | 0.038752498 | 0.109948776 | 1.94E-41 |
| SS15079 | 0.038752498 | 0.109948776 | 1.94E-41 |
| SS03474 | 0.000141494 | 0.006490013 | 2.20E-41 |
| SS09923 | 0.005000211 | 0.031414211 | 3.88E-41 |
| SS02810 | 0.000911097 | 0.014774206 | 4.41E-41 |
| SS05825 | 0.000911097 | 0.014774206 | 4.41E-41 |
| SS08111 | 0.048512451 | 0.106554038 | 4.68E-41 |
| SS04377 | 0.000563212 | 0.012757537 | 4.72E-41 |
| SS04983 | 0.000563212 | 0.012757537 | 4.72E-41 |
| SS00411 | 0.001520062 | 0.01643579 | 5.38E-41 |
| SS08168 | 0.000602393 | 0.014126164 | 6.50E-41 |
| SS08848 | 0.000602393 | 0.014126164 | 6.50E-41 |
| SS00528 | 9.56E-05 | 0.015480358 | 6.73E-41 |
| SS01398 | 9.56E-05 | 0.015480358 | 6.73E-41 |
| SS00511 | 0 | 0.016400311 | 7.69E-41 |
| SS05845 | 0 | 0.016400311 | 7.69E-41 |
| SS05857 | 0 | 0.016400311 | 7.69E-41 |
| SS09063 | 0 | 0.016400311 | 7.69E-41 |
| SS00104 | 0.015949798 | 0.069133752 | 8.00E-41 |
| SS01954 | 0 | 0.014920179 | 8.66E-41 |
| SS02441 | 0 | 0.014920179 | 8.66E-41 |
| SS03281 | 0 | 0.014920179 | 8.66E-41 |
| SS10162 | 0 | 0.006416361 | 9.34E-41 |
| SS15302 | 6.80E-05 | 0.006605996 | 1.04E-40 |
| SS02060 | 0.000672899 | 0.01166486 | 1.06E-40 |
| SS06548 | 0.001214719 | 0.014504993 | 1.50E-40 |
| SS08707 | 0.001214719 | 0.014504993 | 1.50E-40 |
| SS10704 | 0.001214719 | 0.014504993 | 1.50E-40 |
| SS08409 | 0.016907073 | 0.049718349 | 2.17E-40 |
| SS10216 | 0.040807602 | 0.116435667 | 2.43E-40 |
| SS10330 | 0.040807602 | 0.116435667 | 2.43E-40 |
| SS01829 | 0.003298475 | 0.022810599 | 2.54E-40 |
| SS02313 | 0.003298475 | 0.022810599 | 2.54E-40 |
| SS01396 | 3.27E-05 | 0.005581868 | 2.79E-40 |
| SS13933 | 0.00490603 | 0.052708839 | 3.11E-40 |
| SS04036 | 0.000861902 | 0.043694641 | 3.24E-40 |
| SS06557 | 0.000861902 | 0.043694641 | 3.24E-40 |

| | | | |
|---|---|---|---|
| SS15227 | 0.000564657 | 0.007436756 | 3.28E-40 |
| SS09477 | 0 | 0.007799429 | 3.73E-40 |
| SS00309 | 0.000888707 | 0.012067824 | 3.88E-40 |
| SS02360 | 0 | 0.014559724 | 4.59E-40 |
| SS03436 | 0.042649926 | 0.089256431 | 4.73E-40 |
| SS06042 | 0.042649926 | 0.089256431 | 4.73E-40 |
| SS10642 | 0.042649926 | 0.089256431 | 4.73E-40 |
| SS11890 | 0.042649926 | 0.089256431 | 4.73E-40 |
| SS15522 | 0.000233873 | 0.009685776 | 5.67E-40 |
| SS02678 | 2.68E-06 | 0.006172545 | 6.29E-40 |
| SS13551 | 2.68E-06 | 0.006172545 | 6.29E-40 |
| SS06116 | 0.000675262 | 0.008900435 | 7.74E-40 |
| SS09647 | 0.000675262 | 0.008900435 | 7.74E-40 |
| SS01735 | 0.009735866 | 0.0331563 | 8.44E-40 |
| SS06505 | 0.009735866 | 0.0331563 | 8.44E-40 |
| SS07188 | 0.009735866 | 0.0331563 | 8.44E-40 |
| SS08958 | 5.45E-06 | 0.011352541 | 8.81E-40 |
| SS11801 | 5.45E-06 | 0.011352541 | 8.81E-40 |
| SS02418 | 0.028781934 | 0.072313726 | 1.35E-39 |
| SS04812 | 0.028781934 | 0.072313726 | 1.35E-39 |
| SS11149 | 0.028781934 | 0.072313726 | 1.35E-39 |
| SS10329 | 0.047287616 | 0.125659409 | 1.56E-39 |
| SS14849 | 0.003679371 | 0.009922159 | 1.86E-39 |
| SS08591 | 0 | 0.00561268 | 2.19E-39 |
| SS09659 | 4.62E-05 | 0.009188622 | 2.26E-39 |
| SS11936 | 4.62E-05 | 0.009188622 | 2.26E-39 |
| SS04037 | 0 | 0.017668636 | 3.70E-39 |
| SS08564 | 0 | 0.017668636 | 3.70E-39 |
| SS02679 | 1.09E-05 | 0.00726174 | 4.19E-39 |
| SS11081 | 0.013366672 | 0.041067437 | 4.65E-39 |
| SS15406 | 0.013366672 | 0.041067437 | 4.65E-39 |
| SS12886 | 0 | 0.00567583 | 6.11E-39 |
| SS05684 | 0.000305664 | 0.019999346 | 6.26E-39 |
| SS02475 | 0.00870282 | 0.029075078 | 6.45E-39 |
| SS13370 | 0.00870282 | 0.029075078 | 6.45E-39 |
| SS05668 | 0 | 0.006339078 | 7.63E-39 |
| SS10157 | 0 | 0.006339078 | 7.63E-39 |
| SS15513 | 3.01E-05 | 0.005896859 | 7.64E-39 |
| SS03143 | 0.016208504 | 0.037542124 | 7.94E-39 |
| SS09048 | 0.016208504 | 0.037542124 | 7.94E-39 |
| SS10555 | 0.016208504 | 0.037542124 | 7.94E-39 |
| SS02507 | 1.09E-05 | 0.011848882 | 8.86E-39 |
| SS10567 | 1.09E-05 | 0.011848882 | 8.86E-39 |
| SS06465 | 0.009475267 | 0.027476543 | 9.22E-39 |
| SS09522 | 0.009475267 | 0.027476543 | 9.22E-39 |
| SS09549 | 0.009475267 | 0.027476543 | 9.22E-39 |
| SS10458 | 0.009475267 | 0.027476543 | 9.22E-39 |

| | | | |
|---|---|---|---|
| SS09566 | 2.45E-05 | 0.004838294 | 1.54E-38 |
| SS15328 | 0.002018618 | 0.017560452 | 1.63E-38 |
| SS06189 | 0.025574324 | 0.063686478 | 1.88E-38 |
| SS11911 | 0.025574324 | 0.063686478 | 1.88E-38 |
| SS13264 | 0.025574324 | 0.063686478 | 1.88E-38 |
| SS06050 | 0.008925716 | 0.029619497 | 1.91E-38 |
| SS06074 | 0.008925716 | 0.029619497 | 1.91E-38 |
| SS09645 | 0.008925716 | 0.029619497 | 1.91E-38 |
| SS09347 | 0.086484347 | 0.190289121 | 2.08E-38 |
| SS11007 | 0.086484347 | 0.190289121 | 2.08E-38 |
| SS11117 | 0 | 0.018540744 | 2.28E-38 |
| SS07700 | 0.003953114 | 0.03366295 | 2.39E-38 |
| SS13928 | 0.003953114 | 0.03366295 | 2.39E-38 |
| SS06975 | 0.051731738 | 0.119308763 | 2.84E-38 |
| SS10286 | 0.051731738 | 0.119308763 | 2.84E-38 |
| SS10795 | 0.051731738 | 0.119308763 | 2.84E-38 |
| SS04199 | 0.001467621 | 0.017068353 | 2.99E-38 |
| SS13266 | 0.001467621 | 0.017068353 | 2.99E-38 |
| SS05480 | 0.035891822 | 0.087055215 | 3.02E-38 |
| SS07649 | 0.035891822 | 0.087055215 | 3.02E-38 |
| SS13724 | 0.035891822 | 0.087055215 | 3.02E-38 |
| SS13732 | 0.035891822 | 0.087055215 | 3.02E-38 |
| SS03511 | 0.032368008 | 0.063302493 | 3.18E-38 |
| SS05314 | 0.032368008 | 0.063302493 | 3.18E-38 |
| SS08342 | 0.032368008 | 0.063302493 | 3.18E-38 |
| SS02125 | 0 | 0.005957691 | 4.14E-38 |
| SS03615 | 0.007993182 | 0.024212438 | 5.32E-38 |
| SS06513 | 0.007993182 | 0.024212438 | 5.32E-38 |
| SS10418 | 0.007993182 | 0.024212438 | 5.32E-38 |
| SS10439 | 0.007993182 | 0.024212438 | 5.32E-38 |
| SS03546 | 0.001272287 | 0.008411695 | 5.36E-38 |
| SS03620 | 0.001272287 | 0.008411695 | 5.36E-38 |
| SS10411 | 0.001272287 | 0.008411695 | 5.36E-38 |
| SS10422 | 0.001272287 | 0.008411695 | 5.36E-38 |
| SS10444 | 0.001272287 | 0.008411695 | 5.36E-38 |
| SS10453 | 0.001272287 | 0.008411695 | 5.36E-38 |
| SS00494 | 0.023621453 | 0.051545311 | 5.46E-38 |
| SS04034 | 0.0008345 | 0.059940639 | 5.87E-38 |
| SS06555 | 0.0008345 | 0.059940639 | 5.87E-38 |
| SS06093 | 0.022773487 | 0.058958166 | 5.96E-38 |
| SS06109 | 0.022773487 | 0.058958166 | 5.96E-38 |
| SS13921 | 0.022773487 | 0.058958166 | 5.96E-38 |
| SS00550 | 0.003685978 | 0.022056864 | 6.14E-38 |
| SS02042 | 0 | 0.006440901 | 6.50E-38 |
| SS02533 | 0 | 0.006440901 | 6.50E-38 |
| SS03675 | 5.70E-05 | 0.008476838 | 6.51E-38 |
| SS07083 | 5.70E-05 | 0.008476838 | 6.51E-38 |

| | | | |
|---|---|---|---|
| SS01395 | 0.000255747 | 0.007159489 | 8.47E-38 |
| SS00860 | 0 | 0.004561056 | 1.10E-37 |
| SS14848 | 0.010430351 | 0.035549153 | 1.16E-37 |
| SS04119 | 0.003706291 | 0.012344542 | 1.19E-37 |
| SS01409 | 2.68E-06 | 0.011005143 | 1.47E-37 |
| SS03786 | 0 | 0.003848918 | 1.58E-37 |
| SS02445 | 4.38E-05 | 0.106513782 | 2.07E-37 |
| SS03286 | 4.38E-05 | 0.106513782 | 2.07E-37 |
| SS10864 | 0.000735514 | 0.006107969 | 2.58E-37 |
| SS10496 | 0.000125069 | 0.00831849 | 2.60E-37 |
| SS10527 | 0.000125069 | 0.00831849 | 2.60E-37 |
| SS05365 | 0.004546687 | 0.078183086 | 2.80E-37 |
| SS05369 | 0.004546687 | 0.078183086 | 2.80E-37 |
| SS09036 | 0.004546687 | 0.078183086 | 2.80E-37 |
| SS09046 | 0.004546687 | 0.078183086 | 2.80E-37 |
| SS02426 | 0.048476493 | 0.104238489 | 3.02E-37 |
| SS07013 | 0.048476493 | 0.104238489 | 3.02E-37 |
| SS07050 | 0.048476493 | 0.104238489 | 3.02E-37 |
| SS08695 | 0.048476493 | 0.104238489 | 3.02E-37 |
| SS11157 | 0.048476493 | 0.104238489 | 3.02E-37 |
| SS13125 | 0.048476493 | 0.104238489 | 3.02E-37 |
| SS06156 | 0 | 0.005450831 | 3.24E-37 |
| SS06182 | 0 | 0.005450831 | 3.24E-37 |
| SS06194 | 0.01784751 | 0.039650861 | 3.80E-37 |
| SS11934 | 0.01784751 | 0.039650861 | 3.80E-37 |
| SS13281 | 0.01784751 | 0.039650861 | 3.80E-37 |
| SS13922 | 0.010664672 | 0.029248702 | 3.89E-37 |
| SS01738 | 0.010755492 | 0.049429226 | 3.90E-37 |
| SS06507 | 0.010755492 | 0.049429226 | 3.90E-37 |
| SS07191 | 0.010755492 | 0.049429226 | 3.90E-37 |
| SS11940 | 0.010755492 | 0.049429226 | 3.90E-37 |
| SS02289 | 0.016267712 | 0.032566681 | 4.00E-37 |
| SS04240 | 0.016267712 | 0.032566681 | 4.00E-37 |
| SS02817 | 0 | 0.003010342 | 4.25E-37 |
| SS15068 | 0.000695369 | 0.013736444 | 6.15E-37 |
| SS02448 | 2.18E-05 | 0.021707531 | 6.19E-37 |
| SS03289 | 2.18E-05 | 0.021707531 | 6.19E-37 |
| SS01657 | 0 | 0.003571356 | 6.33E-37 |
| SS05863 | 0 | 0.003571356 | 6.33E-37 |
| SS01276 | 0.011567958 | 0.035299161 | 6.59E-37 |
| SS12273 | 2.68E-06 | 0.007754357 | 8.47E-37 |
| SS02359 | 0 | 0.020128876 | 8.52E-37 |
| SS05318 | 0.023175373 | 0.054562938 | 8.88E-37 |
| SS01700 | 0.025027584 | 0.05295439 | 9.47E-37 |
| SS07167 | 0.025027584 | 0.05295439 | 9.47E-37 |
| SS02450 | 0.000147825 | 0.02814796 | 9.58E-37 |
| SS03291 | 0.000147825 | 0.02814796 | 9.58E-37 |

| | | | |
|---|---|---|---|
| SS13451 | 0 | 0.001956937 | 1.41E-36 |
| SS01403 | 2.68E-06 | 0.007030577 | 1.42E-36 |
| SS01953 | 0 | 0.017994566 | 1.57E-36 |
| SS02440 | 0 | 0.017994566 | 1.57E-36 |
| SS03280 | 0 | 0.017994566 | 1.57E-36 |
| SS00829 | 0.000520509 | 0.003799392 | 1.80E-36 |
| SS01416 | 0.000520509 | 0.003799392 | 1.80E-36 |
| SS01217 | 0.012031186 | 0.075962069 | 1.84E-36 |
| SS05522 | 0.001916064 | 0.015715641 | 1.85E-36 |
| SS06118 | 0.003988006 | 0.026311802 | 2.05E-36 |
| SS09649 | 0.003988006 | 0.026311802 | 2.05E-36 |
| SS02361 | 0 | 0.005022762 | 2.71E-36 |
| SS01733 | 0.002391221 | 0.009086792 | 2.77E-36 |
| SS06905 | 0.002391221 | 0.009086792 | 2.77E-36 |
| SS15083 | 0.002391221 | 0.009086792 | 2.77E-36 |
| SS01204 | 0.011599684 | 0.038654843 | 2.80E-36 |
| SS01269 | 0.011599684 | 0.038654843 | 2.80E-36 |
| SS04589 | 8.13E-06 | 0.013875951 | 3.23E-36 |
| SS08196 | 0.008399954 | 0.023501763 | 3.54E-36 |
| SS11218 | 0.008399954 | 0.023501763 | 3.54E-36 |
| SS06736 | 0.001128511 | 0.0238248 | 4.00E-36 |
| SS04737 | 0.000356981 | 0.004264219 | 4.20E-36 |
| SS07529 | 0.000579476 | 0.009915941 | 4.49E-36 |
| SS00933 | 2.19E-05 | 0.005242788 | 5.09E-36 |
| SS01265 | 0.015064566 | 0.039327312 | 5.64E-36 |
| SS09802 | 0.015064566 | 0.039327312 | 5.64E-36 |
| SS11579 | 0.015064566 | 0.039327312 | 5.64E-36 |
| SS14482 | 0.015064566 | 0.039327312 | 5.64E-36 |
| SS00935 | 2.17E-05 | 0.007263279 | 5.95E-36 |
| SS02501 | 2.17E-05 | 0.007263279 | 5.95E-36 |
| SS02681 | 2.17E-05 | 0.007263279 | 5.95E-36 |
| SS15500 | 0.00048205 | 0.033118342 | 7.46E-36 |
| SS13152 | 0.001088928 | 0.013604811 | 7.83E-36 |
| SS04977 | 0.001346039 | 0.008109584 | 7.85E-36 |
| SS09035 | 0.001346039 | 0.008109584 | 7.85E-36 |
| SS12926 | 0.001346039 | 0.008109584 | 7.85E-36 |
| SS08583 | 0.011671475 | 0.034587493 | 8.43E-36 |
| SS00408 | 0.000150188 | 0.010836625 | 9.44E-36 |
| SS00762 | 0.005817288 | 0.015509932 | 1.13E-35 |
| SS04409 | 0.005817288 | 0.015509932 | 1.13E-35 |
| SS10187 | 0.005817288 | 0.015509932 | 1.13E-35 |
| SS06724 | 0.010339452 | 0.035311863 | 1.22E-35 |
| SS08126 | 0.011183495 | 0.029709024 | 1.50E-35 |
| SS10181 | 0.011183495 | 0.029709024 | 1.50E-35 |
| SS13277 | 8.73E-05 | 0.019024098 | 2.12E-35 |
| SS10166 | 1.09E-05 | 0.011015613 | 3.04E-35 |
| SS11951 | 1.09E-05 | 0.011015613 | 3.04E-35 |

| | | | |
|---|---|---|---|
| SS02795 | 8.21E-06 | 0.00482511 | 3.08E-35 |
| SS15467 | 0.000779421 | 0.007115051 | 3.24E-35 |
| SS00838 | 0.002844745 | 0.02684401 | 3.27E-35 |
| SS02357 | 0 | 0.005948094 | 3.60E-35 |
| SS02358 | 0 | 0.005948094 | 3.60E-35 |
| SS01399 | 2.76E-06 | 0.002337114 | 4.21E-35 |
| SS00574 | 0 | 0.009588416 | 4.38E-35 |
| SS06584 | 0 | 0.009588416 | 4.38E-35 |
| SS08245 | 0 | 0.002468042 | 4.57E-35 |
| SS08255 | 0 | 0.002468042 | 4.57E-35 |
| SS03663 | 0.000204189 | 0.015659926 | 4.98E-35 |
| SS04323 | 0.000204189 | 0.015659926 | 4.98E-35 |
| SS12970 | 0.000204189 | 0.015659926 | 4.98E-35 |
| SS15286 | 0.000204189 | 0.015659926 | 4.98E-35 |
| SS03672 | 5.37E-06 | 0.004321665 | 6.26E-35 |
| SS07080 | 5.37E-06 | 0.004321665 | 6.26E-35 |
| SS13006 | 5.37E-06 | 0.004321665 | 6.26E-35 |
| SS08251 | 0 | 0.007164845 | 6.81E-35 |
| SS00166 | 0.015649261 | 0.03371294 | 7.38E-35 |
| SS00191 | 0.015649261 | 0.03371294 | 7.38E-35 |
| SS00524 | 0.015649261 | 0.03371294 | 7.38E-35 |
| SS01772 | 0.015649261 | 0.03371294 | 7.38E-35 |
| SS09906 | 0.015649261 | 0.03371294 | 7.38E-35 |
| SS15069 | 8.05E-06 | 0.006766309 | 7.81E-35 |
| SS15229 | 0.000949154 | 0.009237452 | 8.44E-35 |
| SS03697 | 0.000485537 | 0.006185392 | 8.52E-35 |
| SS10228 | 0.000485537 | 0.006185392 | 8.52E-35 |
| SS01955 | 8.48E-05 | 0.006218682 | 9.33E-35 |
| SS02444 | 8.48E-05 | 0.006218682 | 9.33E-35 |
| SS03284 | 8.48E-05 | 0.006218682 | 9.33E-35 |
| SS07913 | 0 | 0.008258519 | 9.60E-35 |
| SS02969 | 0.003552581 | 0.01426647 | 9.89E-35 |
| SS02995 | 0.003552581 | 0.01426647 | 9.89E-35 |
| SS13147 | 0.003552581 | 0.01426647 | 9.89E-35 |
| SS04617 | 0.090454208 | 0.168137756 | 1.20E-34 |
| SS05704 | 0.004865209 | 0.019204895 | 1.42E-34 |
| SS09860 | 0.004865209 | 0.019204895 | 1.42E-34 |
| SS09480 | 2.68E-06 | 0.004229815 | 1.45E-34 |
| SS10776 | 0.007220459 | 0.03593009 | 1.47E-34 |
| SS03252 | 0.00119405 | 0.010501316 | 1.60E-34 |
| SS03353 | 0.00119405 | 0.010501316 | 1.60E-34 |
| SS01414 | 0.002761059 | 0.009422572 | 1.75E-34 |
| SS01739 | 0.002761059 | 0.009422572 | 1.75E-34 |
| SS06508 | 0.002761059 | 0.009422572 | 1.75E-34 |
| SS07192 | 0.002761059 | 0.009422572 | 1.75E-34 |
| SS04097 | 0 | 0.013882432 | 1.84E-34 |
| SS02052 | 0 | 0.019860424 | 1.85E-34 |

| | | | |
|---|---|---|---|
| SS02536 | 0 | 0.019860424 | 1.85E-34 |
| SS05319 | 0.022282376 | 0.053228661 | 2.04E-34 |
| SS06224 | 0.045136829 | 0.11176975 | 2.17E-34 |
| SS12597 | 0.045136829 | 0.11176975 | 2.17E-34 |
| SS01717 | 0.003962473 | 0.021632231 | 2.24E-34 |
| SS13562 | 0.003962473 | 0.021632231 | 2.24E-34 |
| SS00761 | 0.001132835 | 0.010965187 | 2.33E-34 |
| SS09844 | 0.001132835 | 0.010965187 | 2.33E-34 |
| SS10714 | 7.35E-05 | 0.0044084 | 2.38E-34 |
| SS00175 | 0.010568726 | 0.026566406 | 2.76E-34 |
| SS00202 | 0.010568726 | 0.026566406 | 2.76E-34 |
| SS01729 | 0.010568726 | 0.026566406 | 2.76E-34 |
| SS07637 | 0.010568726 | 0.026566406 | 2.76E-34 |
| SS13174 | 0 | 0.004228269 | 2.83E-34 |
| SS06100 | 0.014772562 | 0.074337118 | 3.27E-34 |
| SS06112 | 0.014772562 | 0.074337118 | 3.27E-34 |
| SS13927 | 0.014772562 | 0.074337118 | 3.27E-34 |
| SS01388 | 8.21E-06 | 0.003616013 | 3.62E-34 |
| SS10792 | 0.00103136 | 0.0174506 | 4.29E-34 |
| SS05882 | 8.13E-06 | 0.002289162 | 5.04E-34 |
| SS05951 | 8.13E-06 | 0.002289162 | 5.04E-34 |
| SS02442 | 0 | 0.008536481 | 5.66E-34 |
| SS03282 | 0 | 0.008536481 | 5.66E-34 |
| SS02595 | 0.109999267 | 0.171291973 | 6.00E-34 |
| SS04651 | 0.109999267 | 0.171291973 | 6.00E-34 |
| SS00176 | 0.00683275 | 0.020649162 | 7.54E-34 |
| SS00203 | 0.00683275 | 0.020649162 | 7.54E-34 |
| SS01730 | 0.00683275 | 0.020649162 | 7.54E-34 |
| SS06308 | 0.00683275 | 0.020649162 | 7.54E-34 |
| SS07638 | 0.00683275 | 0.020649162 | 7.54E-34 |
| SS10367 | 0.002672729 | 0.010686255 | 1.06E-33 |
| SS01071 | 5.37E-06 | 0.005450252 | 1.11E-33 |
| SS02632 | 0.05235262 | 0.090546176 | 1.14E-33 |
| SS04585 | 0.05235262 | 0.090546176 | 1.14E-33 |
| SS02429 | 0.021463578 | 0.037442395 | 1.24E-33 |
| SS04257 | 0.021463578 | 0.037442395 | 1.24E-33 |
| SS07647 | 0.021463578 | 0.037442395 | 1.24E-33 |
| SS00824 | 1.64E-05 | 0.003259212 | 1.37E-33 |
| SS02237 | 0.005987308 | 0.021195099 | 1.41E-33 |
| SS13622 | 0.005987308 | 0.021195099 | 1.41E-33 |
| SS14171 | 0.005987308 | 0.021195099 | 1.41E-33 |
| SS05096 | 0 | 0.007019796 | 1.62E-33 |
| SS01494 | 0.03255537 | 0.089950831 | 1.67E-33 |
| SS13271 | 0.028379646 | 0.050554223 | 1.79E-33 |
| SS06651 | 4.08E-05 | 0.005081432 | 1.81E-33 |
| SS02389 | 0 | 0.004232078 | 1.82E-33 |
| SS03024 | 0.008267395 | 0.02235671 | 2.14E-33 |

| | | | |
|---|---|---|---|
| SS03745 | 0.008267395 | 0.02235671 | 2.14E-33 |
| SS13971 | 0.000389831 | 0.00414014 | 2.26E-33 |
| SS06547 | 1.37E-05 | 0.00480654 | 2.41E-33 |
| SS02723 | 0.005933341 | 0.0443906 | 2.62E-33 |
| SS11950 | 0.005933341 | 0.0443906 | 2.62E-33 |
| SS02122 | 0.000824969 | 0.005921034 | 2.68E-33 |
| SS07007 | 0.004056344 | 0.018099473 | 2.79E-33 |
| SS07528 | 0.004056344 | 0.018099473 | 2.79E-33 |
| SS05744 | 0.004500497 | 0.030378949 | 3.34E-33 |
| SS01498 | 0.013299458 | 0.047582302 | 3.44E-33 |
| SS01005 | 0.010188151 | 0.024877413 | 3.84E-33 |
| SS09049 | 0.010188151 | 0.024877413 | 3.84E-33 |
| SS14918 | 0.010188151 | 0.024877413 | 3.84E-33 |
| SS00205 | 0.071707323 | 0.116041197 | 4.16E-33 |
| SS01777 | 0.071707323 | 0.116041197 | 4.16E-33 |
| SS01784 | 0.071707323 | 0.116041197 | 4.16E-33 |
| SS05336 | 0.071707323 | 0.116041197 | 4.16E-33 |
| SS05347 | 0.071707323 | 0.116041197 | 4.16E-33 |
| SS06919 | 0.071707323 | 0.116041197 | 4.16E-33 |
| SS08724 | 0.071707323 | 0.116041197 | 4.16E-33 |
| SS09917 | 0.071707323 | 0.116041197 | 4.16E-33 |
| SS15082 | 0.071707323 | 0.116041197 | 4.16E-33 |
| SS02287 | 0.002135474 | 0.010276594 | 4.69E-33 |
| SS08903 | 0 | 0.009186534 | 4.97E-33 |
| SS04987 | 0.03013965 | 0.069102503 | 5.16E-33 |
| SS09037 | 0.03013965 | 0.069102503 | 5.16E-33 |
| SS12984 | 0.03013965 | 0.069102503 | 5.16E-33 |
| SS01402 | 2.68E-06 | 0.00308578 | 5.37E-33 |
| SS07244 | 0.028851488 | 0.055035923 | 5.58E-33 |
| SS08778 | 0.028851488 | 0.055035923 | 5.58E-33 |
| SS12739 | 0.028851488 | 0.055035923 | 5.58E-33 |
| SS13341 | 0.028851488 | 0.055035923 | 5.58E-33 |
| SS02417 | 0 | 0.003278495 | 5.95E-33 |
| SS08687 | 0 | 0.003278495 | 5.95E-33 |
| SS03465 | 0.000555722 | 0.004513947 | 7.87E-33 |
| SS08902 | 2.16E-05 | 0.009373657 | 8.35E-33 |
| SS01050 | 0.001270165 | 0.017473725 | 9.55E-33 |
| SS01311 | 0.001270165 | 0.017473725 | 9.55E-33 |
| SS02040 | 0.002564441 | 0.015447603 | 9.57E-33 |
| SS02286 | 0.062121313 | 0.127278685 | 1.01E-32 |
| SS03510 | 0.062121313 | 0.127278685 | 1.01E-32 |
| SS05311 | 0.062121313 | 0.127278685 | 1.01E-32 |
| SS08340 | 0.062121313 | 0.127278685 | 1.01E-32 |
| SS03307 | 0.004291181 | 0.013757699 | 1.13E-32 |
| SS04316 | 5.45E-06 | 0.0024297 | 1.43E-32 |
| SS12937 | 5.45E-06 | 0.0024297 | 1.43E-32 |
| SS09560 | 0.00942387 | 0.028434238 | 1.46E-32 |

| | | | |
|---|---|---|---|
| SS05554 | 0.003320475 | 0.022231084 | 1.54E-32 |
| SS15389 | 0.003320475 | 0.022231084 | 1.54E-32 |
| SS08926 | 0.00921663 | 0.024503225 | 1.57E-32 |
| SS11794 | 0.00921663 | 0.024503225 | 1.57E-32 |
| SS13526 | 0.00921663 | 0.024503225 | 1.57E-32 |
| SS08709 | 0.000128154 | 0.003093351 | 1.62E-32 |
| SS00150 | 0.003540721 | 0.018213248 | 1.81E-32 |
| SS04192 | 0.003540721 | 0.018213248 | 1.81E-32 |
| SS07942 | 0 | 0.007939708 | 1.82E-32 |
| SS10515 | 0.021878654 | 0.057580997 | 1.93E-32 |
| SS10545 | 0.021878654 | 0.057580997 | 1.93E-32 |
| SS02461 | 0.001310987 | 0.021029974 | 2.14E-32 |
| SS11822 | 9.55E-05 | 0.002974364 | 2.46E-32 |
| SS11830 | 9.55E-05 | 0.002974364 | 2.46E-32 |
| SS03417 | 8.21E-06 | 0.010244515 | 2.62E-32 |
| SS03475 | 1.63E-05 | 0.002463534 | 2.73E-32 |
| SS15534 | 0 | 0.003685622 | 3.00E-32 |
| SS04612 | 0.005087543 | 0.023494708 | 3.10E-32 |
| SS08699 | 0.005087543 | 0.023494708 | 3.10E-32 |
| SS03448 | 0.005298546 | 0.029946978 | 3.36E-32 |
| SS07917 | 0.005298546 | 0.029946978 | 3.36E-32 |
| SS02813 | 0 | 0.003425732 | 3.70E-32 |
| SS15070 | 1.10E-05 | 0.002759281 | 4.09E-32 |
| SS01710 | 0.04322377 | 0.07468163 | 4.30E-32 |
| SS03322 | 2.18E-05 | 0.008637896 | 4.62E-32 |
| SS00840 | 0 | 0.007069968 | 5.39E-32 |
| SS06663 | 0 | 0.007069968 | 5.39E-32 |
| SS03321 | 0.000783585 | 0.006430383 | 5.48E-32 |
| SS02318 | 0.014643089 | 0.037607247 | 6.23E-32 |
| SS02626 | 0.014643089 | 0.037607247 | 6.23E-32 |
| SS05656 | 0.000346084 | 0.00455789 | 6.97E-32 |
| SS07225 | 0.000814714 | 0.005040611 | 7.06E-32 |
| SS03262 | 0.028906819 | 0.193846706 | 7.98E-32 |
| SS02044 | 0 | 0.004123709 | 8.60E-32 |
| SS01693 | 0.000934014 | 0.007004607 | 8.83E-32 |
| SS03256 | 0.028906819 | 0.192895933 | 8.89E-32 |
| SS05809 | 0.015470191 | 0.042025695 | 9.56E-32 |
| SS05852 | 0.015470191 | 0.042025695 | 9.56E-32 |
| SS10068 | 0.015470191 | 0.042025695 | 9.56E-32 |
| SS12276 | 0.000155155 | 0.0026318 | 1.22E-31 |
| SS00133 | 0 | 0.003088766 | 1.46E-31 |
| SS03249 | 0.002179221 | 0.012580363 | 1.54E-31 |
| SS03347 | 0.002179221 | 0.012580363 | 1.54E-31 |
| SS09762 | 0.004089469 | 0.013382473 | 1.54E-31 |
| SS02455 | 7.91E-05 | 0.004122158 | 1.64E-31 |
| SS05590 | 0 | 0.005019876 | 1.67E-31 |
| SS01887 | 0 | 0.008936605 | 1.69E-31 |

| | | | |
|---|---|---|---|
| SS00522 | 0.000163769 | 0.004423249 | 1.76E-31 |
| SS08660 | 0.000315196 | 0.007118576 | 2.05E-31 |
| SS04408 | 0.000953284 | 0.006254143 | 2.05E-31 |
| SS08694 | 1.89E-05 | 0.002156536 | 2.05E-31 |
| SS14488 | 0.006366678 | 0.013628179 | 2.12E-31 |
| SS02090 | 0.003974219 | 0.010250416 | 2.21E-31 |
| SS03486 | 0.003974219 | 0.010250416 | 2.21E-31 |
| SS06756 | 0.003974219 | 0.010250416 | 2.21E-31 |
| SS11271 | 0.003974219 | 0.010250416 | 2.21E-31 |
| SS01031 | 2.18E-05 | 0.004081714 | 3.24E-31 |
| SS07910 | 0 | 0.002509001 | 3.32E-31 |
| SS05091 | 0 | 0.017497358 | 3.39E-31 |
| SS06119 | 2.46E-05 | 0.005553721 | 3.46E-31 |
| SS09650 | 2.46E-05 | 0.005553721 | 3.46E-31 |
| SS13223 | 5.47E-05 | 0.006385058 | 3.49E-31 |
| SS06129 | 1.91E-05 | 0.002759245 | 3.70E-31 |
| SS13230 | 0.000245091 | 0.003084673 | 3.70E-31 |
| SS06097 | 0.010616762 | 0.029376069 | 3.87E-31 |
| SS06111 | 0.010616762 | 0.029376069 | 3.87E-31 |
| SS11776 | 0.004231204 | 0.01237896 | 3.91E-31 |
| SS14237 | 0.004231204 | 0.01237896 | 3.91E-31 |
| SS02172 | 0.014550996 | 0.038153429 | 4.05E-31 |
| SS10311 | 6.82E-05 | 0.002789791 | 4.64E-31 |
| SS01723 | 0.010724202 | 0.043208987 | 5.00E-31 |
| SS06500 | 0.010724202 | 0.043208987 | 5.00E-31 |
| SS07174 | 0.010724202 | 0.043208987 | 5.00E-31 |
| SS09910 | 0.010724202 | 0.043208987 | 5.00E-31 |
| SS10427 | 0.010724202 | 0.043208987 | 5.00E-31 |
| SS05090 | 0 | 0.012002726 | 5.30E-31 |
| SS01891 | 0.000215407 | 0.006067497 | 5.54E-31 |
| SS03716 | 0.000215407 | 0.006067497 | 5.54E-31 |
| SS04816 | 0.016783094 | 0.032642917 | 5.69E-31 |
| SS11150 | 0.016783094 | 0.032642917 | 5.69E-31 |
| SS13366 | 0.03701141 | 0.108600783 | 6.28E-31 |
| SS06932 | 0.018871139 | 0.038501172 | 6.93E-31 |
| SS09950 | 0.018871139 | 0.038501172 | 6.93E-31 |
| SS00812 | 0.011771952 | 0.030612308 | 6.94E-31 |
| SS03602 | 0.011771952 | 0.030612308 | 6.94E-31 |
| SS06510 | 0.011771952 | 0.030612308 | 6.94E-31 |
| SS09808 | 0.011771952 | 0.030612308 | 6.94E-31 |
| SS10414 | 0.011771952 | 0.030612308 | 6.94E-31 |
| SS10425 | 0.011771952 | 0.030612308 | 6.94E-31 |
| SS13575 | 0.011771952 | 0.030612308 | 6.94E-31 |
| SS09981 | 0.008538971 | 0.023033189 | 7.71E-31 |
| SS02446 | 5.45E-06 | 0.023149454 | 8.04E-31 |
| SS03287 | 5.45E-06 | 0.023149454 | 8.04E-31 |
| SS10514 | 0.01182523 | 0.026128625 | 8.81E-31 |

| | | | |
|---|---|---|---|
| SS10544 | 0.01182523 | 0.026128625 | 8.81E-31 |
| SS05586 | 1.63E-05 | 0.013791651 | 8.97E-31 |
| SS04618 | 4.90E-05 | 0.01536438 | 9.36E-31 |
| SS08406 | 2.19E-05 | 0.002157986 | 1.01E-30 |
| SS10856 | 2.19E-05 | 0.002157986 | 1.01E-30 |
| SS14458 | 2.19E-05 | 0.002157986 | 1.01E-30 |
| SS13357 | 0 | 0.005473159 | 1.01E-30 |
| SS13169 | 0 | 0.00231023 | 1.03E-30 |
| SS06679 | 0.00601635 | 0.013159951 | 1.12E-30 |
| SS01599 | 0.000503763 | 0.003934274 | 1.12E-30 |
| SS02476 | 0.000503763 | 0.003934274 | 1.12E-30 |
| SS02508 | 1.63E-05 | 0.007021079 | 1.16E-30 |
| SS10570 | 1.63E-05 | 0.007021079 | 1.16E-30 |
| SS13173 | 0 | 0.002562131 | 1.28E-30 |
| SS05935 | 0 | 0.00253669 | 1.50E-30 |
| SS00105 | 0.023470703 | 0.048466798 | 1.74E-30 |
| SS05807 | 0.040593836 | 0.0688093 | 1.74E-30 |
| SS05850 | 0.040593836 | 0.0688093 | 1.74E-30 |
| SS03153 | 0.002361777 | 0.007422619 | 1.90E-30 |
| SS12039 | 0.002361777 | 0.007422619 | 1.90E-30 |
| SS12775 | 0.002361777 | 0.007422619 | 1.90E-30 |
| SS00495 | 2.68E-06 | 0.00125224 | 2.54E-30 |
| SS02474 | 0.015495345 | 0.026695951 | 2.56E-30 |
| SS13369 | 0.015495345 | 0.026695951 | 2.56E-30 |
| SS00572 | 0.010865282 | 0.035512124 | 2.57E-30 |
| SS00982 | 0 | 0.003021212 | 2.60E-30 |
| SS03234 | 0.000394958 | 0.003984468 | 2.90E-30 |
| SS00510 | 1.37E-05 | 0.004149779 | 3.38E-30 |
| SS05844 | 1.37E-05 | 0.004149779 | 3.38E-30 |
| SS05856 | 1.37E-05 | 0.004149779 | 3.38E-30 |
| SS09062 | 1.37E-05 | 0.004149779 | 3.38E-30 |
| SS08854 | 0.002424231 | 0.015147976 | 3.47E-30 |
| SS01188 | 0.050274932 | 0.10757712 | 3.89E-30 |
| SS02767 | 0.050274932 | 0.10757712 | 3.89E-30 |
| SS02542 | 0 | 0.00250178 | 4.21E-30 |
| SS01922 | 0.00095669 | 0.005004099 | 5.01E-30 |
| SS02410 | 0.00095669 | 0.005004099 | 5.01E-30 |
| SS03639 | 0.00095669 | 0.005004099 | 5.01E-30 |
| SS01892 | 0.000354458 | 0.005067236 | 5.03E-30 |
| SS03717 | 0.000354458 | 0.005067236 | 5.03E-30 |
| SS09152 | 0.000354458 | 0.005067236 | 5.03E-30 |
| SS15219 | 0.001920549 | 0.007215928 | 5.85E-30 |
| SS04105 | 0.038236016 | 0.076649596 | 6.34E-30 |
| SS05797 | 0.038236016 | 0.076649596 | 6.34E-30 |
| SS05823 | 0.038236016 | 0.076649596 | 6.34E-30 |
| SS02054 | 0.005066312 | 0.012674491 | 6.39E-30 |
| SS02894 | 0.005066312 | 0.012674491 | 6.39E-30 |

| | | | |
|---|---|---|---|
| SS10355 | 0.005066312 | 0.012674491 | 6.39E-30 |
| SS00843 | 0 | 0.002305586 | 6.40E-30 |
| SS02527 | 0.019486424 | 0.031775093 | 6.48E-30 |
| SS04252 | 0.019486424 | 0.031775093 | 6.48E-30 |
| SS15363 | 0.019486424 | 0.031775093 | 6.48E-30 |
| SS06752 | 0.010102699 | 0.023050483 | 6.56E-30 |
| SS09539 | 0.010102699 | 0.023050483 | 6.56E-30 |
| SS03315 | 0.000394958 | 0.00384545 | 8.47E-30 |
| SS00483 | 0.000356258 | 0.004748865 | 8.87E-30 |
| SS01725 | 0.012078098 | 0.03664709 | 9.47E-30 |
| SS04193 | 0.012078098 | 0.03664709 | 9.47E-30 |
| SS06912 | 0.012078098 | 0.03664709 | 9.47E-30 |
| SS07179 | 0.012078098 | 0.03664709 | 9.47E-30 |
| SS15075 | 0.012078098 | 0.03664709 | 9.47E-30 |
| SS06672 | 0.003435438 | 0.011712062 | 9.99E-30 |
| SS14060 | 0.0027748 | 0.01075899 | 1.01E-29 |
| SS14148 | 3.54E-05 | 0.002839519 | 1.09E-29 |
| SS03653 | 5.53E-06 | 0.005285369 | 1.21E-29 |
| SS07064 | 5.53E-06 | 0.005285369 | 1.21E-29 |
| SS00509 | 0.033756165 | 0.079348982 | 1.25E-29 |
| SS05814 | 0.033756165 | 0.079348982 | 1.25E-29 |
| SS05843 | 0.033756165 | 0.079348982 | 1.25E-29 |
| SS05855 | 0.033756165 | 0.079348982 | 1.25E-29 |
| SS09061 | 0.033756165 | 0.079348982 | 1.25E-29 |
| SS02682 | 0 | 0.002495686 | 1.27E-29 |
| SS13151 | 0.002198089 | 0.02978291 | 1.30E-29 |
| SS01661 | 4.36E-05 | 0.003104649 | 1.41E-29 |
| SS05866 | 4.36E-05 | 0.003104649 | 1.41E-29 |
| SS06099 | 0 | 0.007076725 | 1.52E-29 |
| SS13172 | 0.000147023 | 0.005150586 | 1.57E-29 |
| SS06225 | 0.007966457 | 0.02271928 | 1.63E-29 |
| SS02484 | 0.001896634 | 0.010677019 | 1.72E-29 |
| SS08505 | 2.18E-05 | 0.004006714 | 1.74E-29 |
| SS10160 | 2.18E-05 | 0.004006714 | 1.74E-29 |
| SS08901 | 2.76E-06 | 0.003428752 | 1.79E-29 |
| SS10196 | 2.68E-06 | 0.003636858 | 1.90E-29 |
| SS11090 | 0.024946777 | 0.041721992 | 2.00E-29 |
| SS14992 | 0 | 0.009746596 | 2.16E-29 |
| SS03414 | 0.000754864 | 0.009839861 | 2.45E-29 |
| SS03415 | 0.000754864 | 0.009839861 | 2.45E-29 |
| SS00169 | 0.000136287 | 0.004646727 | 2.62E-29 |
| SS00193 | 0.000136287 | 0.004646727 | 2.62E-29 |
| SS08874 | 0.000136287 | 0.004646727 | 2.62E-29 |
| SS11427 | 0.007038614 | 0.01535359 | 2.69E-29 |
| SS05740 | 4.08E-05 | 0.005726308 | 3.14E-29 |
| SS01400 | 0 | 0.001716267 | 3.25E-29 |
| SS04098 | 3.55E-05 | 0.007770754 | 3.40E-29 |

| | | | |
|---|---|---|---|
| SS08474 | 0 | 0.002403938 | 3.74E-29 |
| SS11076 | 0.008139597 | 0.027207522 | 4.16E-29 |
| SS05093 | 2.76E-06 | 0.005466379 | 4.20E-29 |
| SS08953 | 0.003549014 | 0.018867703 | 4.32E-29 |
| SS08974 | 0.003549014 | 0.018867703 | 4.32E-29 |
| SS09543 | 0.015844744 | 0.032461917 | 5.36E-29 |
| SS09155 | 0.001777048 | 0.009669877 | 5.42E-29 |
| SS04523 | 0.020775699 | 0.038476616 | 5.71E-29 |
| SS00132 | 2.68E-06 | 0.004862928 | 5.77E-29 |
| SS13412 | 0.001862098 | 0.007112942 | 7.39E-29 |
| SS13432 | 0.001862098 | 0.007112942 | 7.39E-29 |
| SS11075 | 0.006013666 | 0.0135463 | 7.40E-29 |
| SS04930 | 0 | 0.003088412 | 7.49E-29 |
| SS09095 | 8.69E-05 | 0.001933987 | 8.13E-29 |
| SS03412 | 2.76E-06 | 0.030762005 | 8.60E-29 |
| SS06144 | 0.033440912 | 0.056657865 | 9.55E-29 |
| SS06174 | 0.033440912 | 0.056657865 | 9.55E-29 |
| SS01404 | 0 | 0.001894142 | 1.05E-28 |
| SS03413 | 2.76E-06 | 0.031057468 | 1.06E-28 |
| SS06240 | 0.045078516 | 0.098191933 | 1.16E-28 |
| SS10292 | 0.045078516 | 0.098191933 | 1.16E-28 |
| SS06108 | 0 | 0.008348272 | 1.18E-28 |
| SS11085 | 0.007015227 | 0.020396317 | 1.25E-28 |
| SS03482 | 0.009793984 | 0.02188818 | 1.32E-28 |
| SS03543 | 0.009793984 | 0.02188818 | 1.32E-28 |
| SS03612 | 0.009793984 | 0.02188818 | 1.32E-28 |
| SS03705 | 0.009793984 | 0.02188818 | 1.32E-28 |
| SS05211 | 0.009793984 | 0.02188818 | 1.32E-28 |
| SS05514 | 0.009793984 | 0.02188818 | 1.32E-28 |
| SS08062 | 0.009793984 | 0.02188818 | 1.32E-28 |
| SS08232 | 0.009793984 | 0.02188818 | 1.32E-28 |
| SS10409 | 0.009793984 | 0.02188818 | 1.32E-28 |
| SS10437 | 0.009793984 | 0.02188818 | 1.32E-28 |
| SS10450 | 0.009793984 | 0.02188818 | 1.32E-28 |
| SS10822 | 0.009793984 | 0.02188818 | 1.32E-28 |
| SS14218 | 0.009793984 | 0.02188818 | 1.32E-28 |
| SS15492 | 0.009793984 | 0.02188818 | 1.32E-28 |
| SS06618 | 0 | 0.004107095 | 1.33E-28 |
| SS00308 | 0.000656233 | 0.006198454 | 1.34E-28 |
| SS00149 | 0.00357582 | 0.016107527 | 1.46E-28 |
| SS04191 | 0.00357582 | 0.016107527 | 1.46E-28 |
| SS06673 | 0.000767481 | 0.004456756 | 1.52E-28 |
| SS05181 | 0.000634762 | 0.005632809 | 1.68E-28 |
| SS07297 | 0 | 0.001727622 | 1.95E-28 |
| SS15324 | 0 | 0.002432591 | 1.95E-28 |
| SS04205 | 0.012868083 | 0.021320153 | 1.95E-28 |
| SS10808 | 0.012868083 | 0.021320153 | 1.95E-28 |

| | | | |
|---|---|---|---|
| SS06245 | 0.008730222 | 0.018836667 | 1.97E-28 |
| SS02411 | 0.000348528 | 0.00614219 | 2.00E-28 |
| SS03642 | 0.000348528 | 0.00614219 | 2.00E-28 |
| SS00131 | 5.45E-06 | 0.001436628 | 2.12E-28 |
| SS15298 | 0.000109286 | 0.001932676 | 2.30E-28 |
| SS07593 | 0 | 0.006490209 | 2.50E-28 |
| SS02824 | 0 | 0.0019322 | 2.86E-28 |
| SS08345 | 0.006454412 | 0.016929713 | 2.88E-28 |
| SS04281 | 0.000196217 | 0.00313119 | 3.12E-28 |
| SS01123 | 0.023480762 | 0.044560616 | 3.31E-28 |
| SS03020 | 0.023480762 | 0.044560616 | 3.31E-28 |
| SS14475 | 0.023480762 | 0.044560616 | 3.31E-28 |
| SS03261 | 0.071192205 | 0.224125855 | 3.33E-28 |
| SS07594 | 0 | 0.003454451 | 3.52E-28 |
| SS10184 | 0 | 0.003454451 | 3.52E-28 |
| SS02452 | 0 | 0.002807185 | 3.65E-28 |
| SS03295 | 0 | 0.002807185 | 3.65E-28 |
| SS11263 | 0.007517303 | 0.014374395 | 3.80E-28 |
| SS05675 | 0 | 0.005309041 | 3.83E-28 |
| SS11945 | 0 | 0.005309041 | 3.83E-28 |
| SS06101 | 0.02416044 | 0.064745296 | 4.13E-28 |
| SS06113 | 0.02416044 | 0.064745296 | 4.13E-28 |
| SS07694 | 0.02416044 | 0.064745296 | 4.13E-28 |
| SS07505 | 0.002070657 | 0.012656118 | 4.15E-28 |
| SS02622 | 0.000102794 | 0.003354303 | 4.59E-28 |
| SS09545 | 1.10E-05 | 0.001594351 | 4.95E-28 |
| SS13448 | 5.45E-06 | 0.005278447 | 5.12E-28 |
| SS03025 | 0.006815282 | 0.018542098 | 5.34E-28 |
| SS03746 | 0.006815282 | 0.018542098 | 5.34E-28 |
| SS02262 | 0.003819064 | 0.012617843 | 5.77E-28 |
| SS10022 | 0.003819064 | 0.012617843 | 5.77E-28 |
| SS02063 | 0 | 0.001399343 | 6.02E-28 |
| SS11931 | 0.000247293 | 0.004416287 | 7.56E-28 |
| SS13554 | 0.000247293 | 0.004416287 | 7.56E-28 |
| SS08547 | 0.00034933 | 0.008039757 | 7.99E-28 |
| SS05680 | 0.003115918 | 0.013419943 | 8.30E-28 |
| SS08578 | 0.003115918 | 0.013419943 | 8.30E-28 |
| SS05216 | 1.63E-05 | 0.002355142 | 8.93E-28 |
| SS10165 | 8.17E-05 | 0.004347959 | 9.49E-28 |
| SS04697 | 5.53E-06 | 0.005866706 | 9.65E-28 |
| SS03250 | 0.001316275 | 0.008477911 | 1.04E-27 |
| SS03348 | 0.001316275 | 0.008477911 | 1.04E-27 |
| SS04215 | 0.000370481 | 0.010507616 | 1.04E-27 |
| SS05498 | 0.000370481 | 0.010507616 | 1.04E-27 |
| SS08681 | 0.000370481 | 0.010507616 | 1.04E-27 |
| SS00862 | 2.76E-06 | 0.001473551 | 1.20E-27 |
| SS13222 | 0.000379256 | 0.004140795 | 1.23E-27 |

| | | | |
|---|---|---|---|
| SS11968 | 0.023133071 | 0.047453966 | 1.26E-27 |
| SS13292 | 0.023133071 | 0.047453966 | 1.26E-27 |
| SS03536 | 5.45E-06 | 0.002686921 | 1.34E-27 |
| SS03603 | 5.45E-06 | 0.002686921 | 1.34E-27 |
| SS05199 | 5.45E-06 | 0.002686921 | 1.34E-27 |
| SS08046 | 5.45E-06 | 0.002686921 | 1.34E-27 |
| SS10405 | 5.45E-06 | 0.002686921 | 1.34E-27 |
| SS10426 | 5.45E-06 | 0.002686921 | 1.34E-27 |
| SS10448 | 5.45E-06 | 0.002686921 | 1.34E-27 |
| SS12857 | 0.000275739 | 0.003460391 | 1.36E-27 |
| SS13362 | 0 | 0.004446157 | 1.44E-27 |
| SS07533 | 0.002625495 | 0.011522984 | 1.49E-27 |
| SS01418 | 0 | 0.002375982 | 1.61E-27 |
| SS02703 | 0.019588622 | 0.030377313 | 1.66E-27 |
| SS06926 | 0.019588622 | 0.030377313 | 1.66E-27 |
| SS09945 | 0.019588622 | 0.030377313 | 1.66E-27 |
| SS00917 | 0.002142241 | 0.007842187 | 1.72E-27 |
| SS08236 | 0.025518878 | 0.043248672 | 2.01E-27 |
| SS15405 | 0.025518878 | 0.043248672 | 2.01E-27 |
| SS03435 | 0.002242076 | 0.010787935 | 2.05E-27 |
| SS06968 | 1.09E-05 | 0.001881402 | 2.14E-27 |
| SS06250 | 0.004981423 | 0.011129514 | 2.21E-27 |
| SS08195 | 0.004981423 | 0.011129514 | 2.21E-27 |
| SS11217 | 0.004981423 | 0.011129514 | 2.21E-27 |
| SS09493 | 0 | 0.00219558 | 2.28E-27 |
| SS02872 | 0 | 0.005967747 | 2.35E-27 |
| SS07261 | 0.007793719 | 0.014687914 | 2.92E-27 |
| SS03481 | 0.027710085 | 0.044723926 | 2.93E-27 |
| SS04258 | 0.027710085 | 0.044723926 | 2.93E-27 |
| SS15376 | 0.027710085 | 0.044723926 | 2.93E-27 |
| SS15429 | 0.027710085 | 0.044723926 | 2.93E-27 |
| SS15491 | 0.027710085 | 0.044723926 | 2.93E-27 |
| SS05626 | 0.004797065 | 0.01405788 | 3.03E-27 |
| SS07717 | 0.004797065 | 0.01405788 | 3.03E-27 |
| SS00077 | 0 | 0.004144904 | 3.22E-27 |
| SS05189 | 0 | 0.004144904 | 3.22E-27 |
| SS06492 | 0 | 0.004144904 | 3.22E-27 |
| SS10494 | 0.000617534 | 0.003809549 | 3.38E-27 |
| SS01666 | 0.000879451 | 0.003811392 | 3.62E-27 |
| SS01945 | 0.000879451 | 0.003811392 | 3.62E-27 |
| SS02827 | 0.000879451 | 0.003811392 | 3.62E-27 |
| SS06261 | 0.038253932 | 0.069705523 | 3.71E-27 |
| SS06766 | 0.038253932 | 0.069705523 | 3.71E-27 |
| SS13778 | 0.038253932 | 0.069705523 | 3.71E-27 |
| SS14893 | 0.038253932 | 0.069705523 | 3.71E-27 |
| SS09485 | 5.45E-06 | 0.003620864 | 3.97E-27 |
| SS01670 | 0.000321848 | 0.002767753 | 4.23E-27 |

| | | | |
|---|---|---|---|
| SS02838 | 0.000321848 | 0.002767753 | 4.23E-27 |
| SS04796 | 0.020954447 | 0.036207016 | 4.64E-27 |
| SS06133 | 0.020954447 | 0.036207016 | 4.64E-27 |
| SS06164 | 0.020954447 | 0.036207016 | 4.64E-27 |
| SS08378 | 0.020954447 | 0.036207016 | 4.64E-27 |
| SS01401 | 8.21E-06 | 0.001526904 | 4.73E-27 |
| SS00851 | 0.00242984 | 0.017875037 | 5.44E-27 |
| SS10828 | 0.011093559 | 0.018966538 | 5.71E-27 |
| SS00370 | 0.003915527 | 0.009461356 | 6.68E-27 |
| SS06875 | 0.003915527 | 0.009461356 | 6.68E-27 |
| SS12332 | 0.003915527 | 0.009461356 | 6.68E-27 |
| SS14213 | 0.003915527 | 0.009461356 | 6.68E-27 |
| SS04347 | 0 | 0.003109195 | 6.88E-27 |
| SS13068 | 0 | 0.003109195 | 6.88E-27 |
| SS02819 | 0 | 0.001856698 | 6.95E-27 |
| SS08422 | 0.026458409 | 0.053186676 | 7.00E-27 |
| SS08464 | 0.026458409 | 0.053186676 | 7.00E-27 |
| SS02443 | 0 | 0.006416428 | 7.65E-27 |
| SS03283 | 0 | 0.006416428 | 7.65E-27 |
| SS14056 | 0.059546091 | 0.088009397 | 9.30E-27 |
| SS09567 | 0.000299654 | 0.005149386 | 1.14E-26 |
| SS02704 | 0.023080585 | 0.038513431 | 1.14E-26 |
| SS03548 | 0.023080585 | 0.038513431 | 1.14E-26 |
| SS06928 | 0.023080585 | 0.038513431 | 1.14E-26 |
| SS13171 | 0.010439768 | 0.020845645 | 1.16E-26 |
| SS14982 | 0.000506768 | 0.004439441 | 1.23E-26 |
| SS00376 | 2.68E-06 | 0.003435132 | 1.51E-26 |
| SS13176 | 0 | 0.001443428 | 1.55E-26 |
| SS00782 | 0.011302794 | 0.026395185 | 1.58E-26 |
| SS03109 | 0 | 0.003108345 | 1.60E-26 |
| SS13861 | 0 | 0.001087776 | 1.76E-26 |
| SS02422 | 0.000130838 | 0.002750985 | 1.81E-26 |
| SS02481 | 0 | 0.002003599 | 2.02E-26 |
| SS06282 | 0.000362269 | 0.003301282 | 2.49E-26 |
| SS01511 | 0.001432248 | 0.004526586 | 2.49E-26 |
| SS13247 | 0.001432248 | 0.004526586 | 2.49E-26 |
| SS00402 | 0.024359938 | 0.05072302 | 2.57E-26 |
| SS07784 | 0.024359938 | 0.05072302 | 2.57E-26 |
| SS09158 | 0.024359938 | 0.05072302 | 2.57E-26 |
| SS09612 | 0 | 0.002816591 | 2.58E-26 |
| SS04566 | 0 | 0.003102809 | 2.70E-26 |
| SS14042 | 0.015679702 | 0.032563094 | 2.74E-26 |
| SS02134 | 0 | 0.001288602 | 2.85E-26 |
| SS00953 | 0.002094973 | 0.006469688 | 3.35E-26 |
| SS05918 | 0.002094973 | 0.006469688 | 3.35E-26 |
| SS06910 | 0.043998534 | 0.07053741 | 3.41E-26 |
| SS02473 | 0 | 0.001359762 | 3.65E-26 |

| | | | |
|---|---|---|---|
| SS06662 | 0 | 0.00115211 | 4.02E-26 |
| SS09449 | 0 | 0.001483589 | 4.13E-26 |
| SS00502 | 0 | 0.003589467 | 4.33E-26 |
| SS05812 | 0 | 0.003589467 | 4.33E-26 |
| SS05834 | 0 | 0.003589467 | 4.33E-26 |
| SS01372 | 1.62E-05 | 0.001484845 | 4.61E-26 |
| SS08265 | 0.000103918 | 0.001923792 | 4.62E-26 |
| SS15301 | 4.64E-05 | 0.005259742 | 4.62E-26 |
| SS03561 | 0 | 0.005067922 | 4.76E-26 |
| SS14139 | 0 | 0.005067922 | 4.76E-26 |
| SS00084 | 0.034764642 | 0.052366259 | 5.84E-26 |
| SS04237 | 0.034764642 | 0.052366259 | 5.84E-26 |
| SS03497 | 0 | 0.001527474 | 5.88E-26 |
| SS03370 | 0 | 0.001790691 | 5.96E-26 |
| SS13553 | 2.46E-05 | 0.001298446 | 6.48E-26 |
| SS01596 | 0 | 0.002904029 | 6.78E-26 |
| SS00833 | 9.82E-05 | 0.001452181 | 7.08E-26 |
| SS05190 | 0 | 0.003676566 | 7.17E-26 |
| SS03247 | 0.000525716 | 0.00373876 | 7.52E-26 |
| SS03341 | 0.000525716 | 0.00373876 | 7.52E-26 |
| SS03512 | 0.020254387 | 0.034110485 | 7.71E-26 |
| SS05315 | 0.020254387 | 0.034110485 | 7.71E-26 |
| SS06474 | 0.003193077 | 0.007993474 | 8.21E-26 |
| SS15216 | 7.86E-05 | 0.003324658 | 8.30E-26 |
| SS01127 | 0.022064973 | 0.047853722 | 8.34E-26 |
| SS08950 | 0.022064973 | 0.047853722 | 8.34E-26 |
| SS10282 | 0.022064973 | 0.047853722 | 8.34E-26 |
| SS11003 | 0.022064973 | 0.047853722 | 8.34E-26 |
| SS02815 | 0 | 0.001732803 | 8.42E-26 |
| SS04233 | 0.034786435 | 0.052301724 | 8.94E-26 |
| SS03425 | 5.45E-06 | 0.001181444 | 9.70E-26 |
| SS15218 | 0.000470832 | 0.003237684 | 9.97E-26 |
| SS01387 | 0.003509076 | 0.009918933 | 1.08E-25 |
| SS10010 | 5.46E-05 | 0.002846009 | 1.10E-25 |
| SS12321 | 0 | 0.00276825 | 1.11E-25 |
| SS02559 | 0.009538123 | 0.016646297 | 1.18E-25 |
| SS04602 | 0.009538123 | 0.016646297 | 1.18E-25 |
| SS08551 | 0 | 0.001912356 | 1.20E-25 |
| SS09619 | 0 | 0.001912356 | 1.20E-25 |
| SS02859 | 0.002056916 | 0.008180379 | 1.29E-25 |
| SS12983 | 0.002056916 | 0.008180379 | 1.29E-25 |
| SS08277 | 0.047004903 | 0.113713618 | 1.34E-25 |
| SS06479 | 0.002320313 | 0.007801297 | 1.38E-25 |
| SS01702 | 0.024093524 | 0.038069315 | 1.48E-25 |
| SS02583 | 0.024093524 | 0.038069315 | 1.48E-25 |
| SS07169 | 0.024093524 | 0.038069315 | 1.48E-25 |
| SS05677 | 0.000120022 | 0.002059727 | 1.64E-25 |

| | | | |
|---|---|---|---|
| SS08575 | 0.000120022 | 0.002059727 | 1.64E-25 |
| SS05794 | 0.000446837 | 0.003302929 | 1.64E-25 |
| SS11732 | 0.000348367 | 0.004435332 | 1.77E-25 |
| SS03101 | 0.00386429 | 0.008131031 | 1.89E-25 |
| SS03103 | 0.00386429 | 0.008131031 | 1.89E-25 |
| SS10321 | 0.00386429 | 0.008131031 | 1.89E-25 |
| SS02468 | 0 | 0.001001314 | 1.94E-25 |
| SS05776 | 0 | 0.002668103 | 1.95E-25 |
| SS05681 | 0.000311193 | 0.004301584 | 2.13E-25 |
| SS08579 | 0.000311193 | 0.004301584 | 2.13E-25 |
| SS12272 | 0.000195976 | 0.00261917 | 2.17E-25 |
| SS04813 | 6.24E-05 | 0.002294807 | 2.19E-25 |
| SS00955 | 0.000703065 | 0.006102125 | 2.20E-25 |
| SS02651 | 0.00141346 | 0.004330857 | 2.35E-25 |
| SS03441 | 8.05E-06 | 0.002282217 | 2.36E-25 |
| SS13600 | 5.37E-06 | 0.001808551 | 2.59E-25 |
| SS01012 | 0.041608358 | 0.058995638 | 2.68E-25 |
| SS14015 | 0.041608358 | 0.058995638 | 2.68E-25 |
| SS03276 | 0 | 0.00113198 | 2.76E-25 |
| SS14162 | 0.001765394 | 0.006790675 | 2.78E-25 |
| SS10715 | 0 | 0.002815569 | 2.84E-25 |
| SS04109 | 0.001720925 | 0.015021075 | 3.05E-25 |
| SS07959 | 0 | 0.002108638 | 3.75E-25 |
| SS01812 | 0 | 0.001106682 | 4.13E-25 |
| SS02004 | 0 | 0.001106682 | 4.13E-25 |
| SS03189 | 0 | 0.001106682 | 4.13E-25 |
| SS09665 | 0 | 0.001106682 | 4.13E-25 |
| SS04259 | 0.003634787 | 0.01087561 | 4.27E-25 |
| SS15377 | 0.003634787 | 0.01087561 | 4.27E-25 |
| SS05927 | 0.002658506 | 0.006529249 | 4.65E-25 |
| SS04644 | 0.000179632 | 0.002876053 | 4.79E-25 |
| SS03654 | 0.005668545 | 0.013295868 | 4.80E-25 |
| SS07065 | 0.005668545 | 0.013295868 | 4.80E-25 |
| SS12940 | 0.005668545 | 0.013295868 | 4.80E-25 |
| SS11172 | 0 | 0.002006164 | 5.11E-25 |
| SS08939 | 0 | 0.001981378 | 5.13E-25 |
| SS13849 | 0 | 0.001981378 | 5.13E-25 |
| SS11260 | 0.008893669 | 0.015302231 | 5.43E-25 |
| SS09054 | 1.09E-05 | 0.001714312 | 5.82E-25 |
| SS09087 | 1.09E-05 | 0.001714312 | 5.82E-25 |
| SS10359 | 0.005018276 | 0.011286787 | 6.44E-25 |
| SS06712 | 9.00E-05 | 0.002045102 | 6.47E-25 |
| SS13221 | 0.000491386 | 0.0051566 | 6.69E-25 |
| SS00836 | 0.000503281 | 0.004398074 | 6.82E-25 |
| SS13175 | 0 | 0.001347433 | 7.03E-25 |
| SS09937 | 0.000762274 | 0.00347484 | 7.75E-25 |
| SS02816 | 0 | 0.002095656 | 7.76E-25 |

| | | | |
|---|---|---|---|
| SS05553 | 4.35E-05 | 0.002201393 | 7.85E-25 |
| SS01823 | 0 | 0.003100751 | 8.42E-25 |
| SS05592 | 0 | 0.002152166 | 8.64E-25 |
| SS01788 | 0.000101315 | 0.003098036 | 8.66E-25 |
| SS05481 | 0.000101315 | 0.003098036 | 8.66E-25 |
| SS07229 | 0.000101315 | 0.003098036 | 8.66E-25 |
| SS09506 | 0.000101315 | 0.003098036 | 8.66E-25 |
| SS09763 | 0.000101315 | 0.003098036 | 8.66E-25 |
| SS05322 | 0.032638184 | 0.055680125 | 9.96E-25 |
| SS12275 | 0.000379095 | 0.006899067 | 9.98E-25 |
| SS05746 | 0.001217805 | 0.003971811 | 9.99E-25 |
| SS12329 | 1.89E-05 | 0.00180251 | 1.04E-24 |
| SS04525 | 0.001680906 | 0.008273485 | 1.08E-24 |
| SS11089 | 0.027415914 | 0.043997959 | 1.11E-24 |
| SS05000 | 0.001524467 | 0.008754962 | 1.16E-24 |
| SS06152 | 0.001524467 | 0.008754962 | 1.16E-24 |
| SS06107 | 0.001541454 | 0.007879118 | 1.16E-24 |
| SS13170 | 2.68E-06 | 0.002417245 | 1.39E-24 |
| SS04842 | 5.40E-05 | 0.002640418 | 1.57E-24 |
| SS13876 | 5.40E-05 | 0.002640418 | 1.57E-24 |
| SS01890 | 0.007628952 | 0.014332171 | 1.60E-24 |
| SS04810 | 0.007628952 | 0.014332171 | 1.60E-24 |
| SS11147 | 0.007628952 | 0.014332171 | 1.60E-24 |
| SS02362 | 0 | 0.001815752 | 1.62E-24 |
| SS07631 | 2.76E-06 | 0.002542117 | 1.68E-24 |
| SS04099 | 0 | 0.001488741 | 1.94E-24 |
| SS05679 | 0.00168351 | 0.00851996 | 1.98E-24 |
| SS08577 | 0.00168351 | 0.00851996 | 1.98E-24 |
| SS04348 | 0 | 0.001819725 | 2.04E-24 |
| SS13069 | 0 | 0.001819725 | 2.04E-24 |
| SS04239 | 0.009626098 | 0.015508491 | 2.07E-24 |
| SS07723 | 1.09E-05 | 0.002852814 | 2.16E-24 |
| SS09458 | 0.011067361 | 0.019019438 | 2.19E-24 |
| SS02427 | 1.11E-05 | 0.00151863 | 2.22E-24 |
| SS11967 | 0.017849839 | 0.038049187 | 2.39E-24 |
| SS13291 | 0.017849839 | 0.038049187 | 2.39E-24 |
| SS00102 | 0.008112642 | 0.018111783 | 2.56E-24 |
| SS03076 | 0.008112642 | 0.018111783 | 2.56E-24 |
| SS06154 | 0 | 0.003733592 | 2.78E-24 |
| SS11804 | 0.013530808 | 0.025927987 | 2.82E-24 |
| SS07949 | 0 | 0.004144481 | 2.89E-24 |
| SS09928 | 8.13E-06 | 0.001028491 | 3.04E-24 |
| SS13177 | 0 | 0.001684162 | 3.18E-24 |
| SS04204 | 0.003039081 | 0.010812273 | 3.19E-24 |
| SS08138 | 0.027555527 | 0.048881184 | 3.30E-24 |
| SS08159 | 0.027555527 | 0.048881184 | 3.30E-24 |
| SS10870 | 0.027555527 | 0.048881184 | 3.30E-24 |

| | | | |
|---|---|---|---|
| SS01720 | 0.005748307 | 0.012241947 | 3.31E-24 |
| SS06899 | 0.005748307 | 0.012241947 | 3.31E-24 |
| SS15072 | 0.005748307 | 0.012241947 | 3.31E-24 |
| SS03274 | 0.005853063 | 0.031073541 | 3.32E-24 |
| SS15017 | 0.000139131 | 0.002931738 | 3.41E-24 |
| SS07303 | 0 | 0.001516862 | 3.70E-24 |
| SS00217 | 0.000667772 | 0.010110888 | 3.97E-24 |
| SS02392 | 0 | 0.001368338 | 4.18E-24 |
| SS01658 | 0 | 0.001041029 | 4.69E-24 |
| SS05865 | 0 | 0.001041029 | 4.69E-24 |
| SS05877 | 0 | 0.001041029 | 4.69E-24 |
| SS05946 | 0 | 0.001041029 | 4.69E-24 |
| SS04224 | 0.000591497 | 0.002536652 | 5.08E-24 |
| SS05561 | 0.000591256 | 0.014792347 | 5.61E-24 |
| SS03096 | 0 | 0.001516408 | 7.16E-24 |
| SS05148 | 0.010148602 | 0.020954673 | 7.21E-24 |
| SS03092 | 0.006485301 | 0.014019423 | 7.60E-24 |
| SS05406 | 0.006485301 | 0.014019423 | 7.60E-24 |
| SS05479 | 0.006485301 | 0.014019423 | 7.60E-24 |
| SS13731 | 0.006485301 | 0.014019423 | 7.60E-24 |
| SS05562 | 0 | 0.00438626 | 7.61E-24 |
| SS00412 | 0.001597221 | 0.0062316 | 8.08E-24 |
| SS01036 | 0 | 0.000794552 | 8.70E-24 |
| SS03449 | 0 | 0.003090824 | 8.77E-24 |
| SS04275 | 0 | 0.001488206 | 8.85E-24 |
| SS04765 | 0 | 0.001488206 | 8.85E-24 |
| SS06187 | 0.057261587 | 0.096914499 | 8.86E-24 |
| SS10152 | 0.057261587 | 0.096914499 | 8.86E-24 |
| SS11908 | 0.057261587 | 0.096914499 | 8.86E-24 |
| SS13257 | 0.057261587 | 0.096914499 | 8.86E-24 |
| SS15357 | 0.057261587 | 0.096914499 | 8.86E-24 |
| SS00576 | 2.46E-05 | 0.002190788 | 9.19E-24 |
| SS05551 | 2.46E-05 | 0.002190788 | 9.19E-24 |
| SS04255 | 0 | 0.000531705 | 1.02E-23 |
| SS06645 | 2.75E-05 | 0.002163255 | 1.11E-23 |
| SS13735 | 0 | 0.001434734 | 1.15E-23 |
| SS11253 | 0.013718893 | 0.024036989 | 1.16E-23 |
| SS11575 | 0.013718893 | 0.024036989 | 1.16E-23 |
| SS14191 | 0.013718893 | 0.024036989 | 1.16E-23 |
| SS14255 | 0.013718893 | 0.024036989 | 1.16E-23 |
| SS14318 | 0.013718893 | 0.024036989 | 1.16E-23 |
| SS14343 | 0.013718893 | 0.024036989 | 1.16E-23 |
| SS15238 | 0.013718893 | 0.024036989 | 1.16E-23 |
| SS01370 | 0.000863668 | 0.005203342 | 1.17E-23 |
| SS02811 | 0 | 0.001905809 | 1.19E-23 |
| SS00907 | 0.000593297 | 0.003209078 | 1.25E-23 |
| SS03229 | 0.000593297 | 0.003209078 | 1.25E-23 |

| | | | |
|---|---|---|---|
| SS03319 | 0.000593297 | 0.003209078 | 1.25E-23 |
| SS00503 | 0.024351404 | 0.045727125 | 1.27E-23 |
| SS05835 | 0.024351404 | 0.045727125 | 1.27E-23 |
| SS15396 | 0.024351404 | 0.045727125 | 1.27E-23 |
| SS05266 | 0.036107481 | 0.058132942 | 1.30E-23 |
| SS13633 | 0.014209833 | 0.032342167 | 1.34E-23 |
| SS10007 | 0.000143937 | 0.003193288 | 1.41E-23 |
| SS02923 | 0 | 0.001575535 | 1.47E-23 |
| SS13273 | 0 | 0.00264166 | 1.47E-23 |
| SS01664 | 0.000947675 | 0.003729373 | 1.51E-23 |
| SS10783 | 0.001384257 | 0.006245634 | 1.58E-23 |
| SS15033 | 0 | 0.001264927 | 1.63E-23 |
| SS01719 | 0.000198741 | 0.002462637 | 1.73E-23 |
| SS06898 | 0.000198741 | 0.002462637 | 1.73E-23 |
| SS09476 | 2.76E-06 | 0.003410449 | 1.83E-23 |
| SS02624 | 0.010008026 | 0.024181703 | 1.92E-23 |
| SS06199 | 0.003499819 | 0.0100347 | 2.01E-23 |
| SS09658 | 0.003499819 | 0.0100347 | 2.01E-23 |
| SS01397 | 0.000378774 | 0.002246892 | 2.04E-23 |
| SS14460 | 0.000378774 | 0.002246892 | 2.04E-23 |
| SS07306 | 0 | 0.000936493 | 2.05E-23 |
| SS06264 | 0.003315496 | 0.011514887 | 2.07E-23 |
| SS03422 | 0.000149867 | 0.003478202 | 2.15E-23 |
| SS02454 | 0 | 0.001003713 | 2.50E-23 |
| SS11080 | 0.005186851 | 0.01215432 | 2.84E-23 |
| SS13565 | 0.021647305 | 0.052715685 | 2.89E-23 |
| SS05569 | 0.000288677 | 0.01038251 | 3.16E-23 |
| SS03357 | 0 | 0.001446956 | 3.74E-23 |
| SS11258 | 0.008234064 | 0.016728834 | 3.88E-23 |
| SS03040 | 6.82E-05 | 0.001291537 | 3.90E-23 |
| SS09467 | 6.82E-05 | 0.001291537 | 3.90E-23 |
| SS05587 | 0 | 0.00609602 | 3.92E-23 |
| SS03292 | 0.00046863 | 0.004088522 | 4.33E-23 |
| SS05983 | 0.011005871 | 0.025079125 | 4.59E-23 |
| SS03345 | 0.00243569 | 0.018879224 | 4.60E-23 |
| SS03476 | 0.00243569 | 0.018879224 | 4.60E-23 |
| SS04691 | 0.002337541 | 0.006178687 | 4.88E-23 |
| SS07579 | 0 | 0.001051262 | 5.13E-23 |
| SS10860 | 0.000953846 | 0.003463318 | 5.22E-23 |
| SS15034 | 1.37E-05 | 0.001505553 | 6.03E-23 |
| SS13915 | 0 | 0.000578779 | 6.14E-23 |
| SS00165 | 0.005106652 | 0.011179627 | 6.16E-23 |
| SS00190 | 0.005106652 | 0.011179627 | 6.16E-23 |
| SS09905 | 0.005106652 | 0.011179627 | 6.16E-23 |
| SS14184 | 2.76E-06 | 0.00153898 | 6.29E-23 |
| SS07599 | 0.004092199 | 0.009488449 | 6.34E-23 |
| SS00653 | 0.0132674 | 0.03087201 | 6.34E-23 |

| | | | |
|---|---|---|---|
| SS06935 | 0.0132674 | 0.03087201 | 6.34E-23 |
| SS08605 | 0 | 0.001865871 | 6.72E-23 |
| SS15035 | 2.68E-06 | 0.001077477 | 6.86E-23 |
| SS15327 | 0.001282301 | 0.006830353 | 7.14E-23 |
| SS02957 | 0 | 0.001764604 | 7.19E-23 |
| SS05719 | 1.09E-05 | 0.004407666 | 7.22E-23 |
| SS03625 | 0.000236879 | 0.005385701 | 7.55E-23 |
| SS01287 | 0 | 0.001341766 | 7.62E-23 |
| SS00821 | 0.00699575 | 0.015912518 | 7.90E-23 |
| SS01108 | 0.00699575 | 0.015912518 | 7.90E-23 |
| SS03230 | 0.001551949 | 0.00459627 | 8.13E-23 |
| SS03320 | 0.001551949 | 0.00459627 | 8.13E-23 |
| SS15358 | 0.001551949 | 0.00459627 | 8.13E-23 |
| SS05079 | 0 | 0.001433986 | 8.17E-23 |
| SS06633 | 3.51E-05 | 0.001488789 | 8.20E-23 |
| SS01826 | 0.022564412 | 0.04402127 | 9.00E-23 |
| SS10212 | 0.022564412 | 0.04402127 | 9.00E-23 |
| SS10325 | 0.022564412 | 0.04402127 | 9.00E-23 |
| SS06470 | 0.041695082 | 0.0604423 | 9.46E-23 |
| SS09558 | 0.041695082 | 0.0604423 | 9.46E-23 |
| SS10495 | 0.041695082 | 0.0604423 | 9.46E-23 |
| SS02332 | 0 | 0.002645471 | 1.00E-22 |
| SS08048 | 2.76E-06 | 0.001973975 | 1.02E-22 |
| SS08073 | 2.76E-06 | 0.001973975 | 1.02E-22 |
| SS02453 | 0.002893504 | 0.008154169 | 1.09E-22 |
| SS03236 | 0.00068648 | 0.002558244 | 1.09E-22 |
| SS03333 | 0.00068648 | 0.002558244 | 1.09E-22 |
| SS01501 | 0.01276881 | 0.024771464 | 1.15E-22 |
| SS01684 | 0.01276881 | 0.024771464 | 1.15E-22 |
| SS13568 | 0.01276881 | 0.024771464 | 1.15E-22 |
| SS01905 | 0 | 0.00072813 | 1.17E-22 |
| SS02127 | 0.028445587 | 0.045183853 | 1.23E-22 |
| SS03102 | 0.028445587 | 0.045183853 | 1.23E-22 |
| SS10322 | 0.028445587 | 0.045183853 | 1.23E-22 |
| SS03527 | 0.018213495 | 0.028791776 | 1.27E-22 |
| SS05330 | 0.018213495 | 0.028791776 | 1.27E-22 |
| SS08351 | 0.018213495 | 0.028791776 | 1.27E-22 |
| SS13881 | 2.76E-06 | 0.002971196 | 1.33E-22 |
| SS15036 | 9.52E-05 | 0.001091374 | 1.34E-22 |
| SS08711 | 5.20E-05 | 0.002213572 | 1.36E-22 |
| SS14836 | 0.003030995 | 0.012951652 | 1.44E-22 |
| SS13201 | 0 | 0.000926842 | 1.45E-22 |
| SS07531 | 0.000277941 | 0.002899721 | 1.48E-22 |
| SS04280 | 0.00021793 | 0.004308891 | 1.52E-22 |
| SS05678 | 0.000367556 | 0.013301469 | 1.53E-22 |
| SS08576 | 0.000367556 | 0.013301469 | 1.53E-22 |
| SS02778 | 0.032934431 | 0.050025444 | 1.59E-22 |

| | | | |
|---|---|---|---|
| SS11436 | 0.032934431 | 0.050025444 | 1.59E-22 |
| SS13931 | 0.004726318 | 0.009719717 | 1.63E-22 |
| SS15231 | 0 | 0.001892489 | 1.72E-22 |
| SS03935 | 0 | 0.003775658 | 1.74E-22 |
| SS02403 | 0.000114092 | 0.002627848 | 1.80E-22 |
| SS03632 | 0.000114092 | 0.002627848 | 1.80E-22 |
| SS11257 | 0.010442292 | 0.016547175 | 1.81E-22 |
| SS04210 | 0 | 0.004903082 | 1.85E-22 |
| SS14088 | 0 | 0.00200663 | 1.99E-22 |
| SS05556 | 0 | 0.00373789 | 2.02E-22 |
| SS11082 | 0 | 0.00373789 | 2.02E-22 |
| SS06304 | 0.000327056 | 0.002151594 | 2.06E-22 |
| SS06484 | 0.014335349 | 0.024705306 | 2.27E-22 |
| SS05144 | 0 | 0.000778924 | 2.29E-22 |
| SS10489 | 0.042893388 | 0.064575177 | 2.42E-22 |
| SS05908 | 3.53E-05 | 0.002036745 | 2.68E-22 |
| SS15397 | 3.53E-05 | 0.002036745 | 2.68E-22 |
| SS13219 | 0.010026378 | 0.016969862 | 2.79E-22 |
| SS00172 | 0.084412199 | 0.121844121 | 2.96E-22 |
| SS00197 | 0.084412199 | 0.121844121 | 2.96E-22 |
| SS00527 | 0.084412199 | 0.121844121 | 2.96E-22 |
| SS00651 | 0.084412199 | 0.121844121 | 2.96E-22 |
| SS01774 | 0.084412199 | 0.121844121 | 2.96E-22 |
| SS01783 | 0.084412199 | 0.121844121 | 2.96E-22 |
| SS06933 | 0.084412199 | 0.121844121 | 2.96E-22 |
| SS07633 | 0.084412199 | 0.121844121 | 2.96E-22 |
| SS08211 | 0.084412199 | 0.121844121 | 2.96E-22 |
| SS09911 | 0.084412199 | 0.121844121 | 2.96E-22 |
| SS11906 | 0.084412199 | 0.121844121 | 2.96E-22 |
| SS07769 | 5.37E-06 | 0.00248773 | 3.59E-22 |
| SS12973 | 5.37E-06 | 0.00248773 | 3.59E-22 |
| SS05594 | 0 | 0.001308493 | 3.80E-22 |
| SS09474 | 0 | 0.001778336 | 3.92E-22 |
| SS02992 | 2.72E-05 | 0.000766823 | 3.96E-22 |
| SS02323 | 0.000117338 | 0.003576175 | 4.10E-22 |
| SS00405 | 0.004145397 | 0.017712575 | 4.12E-22 |
| SS11274 | 0.017670241 | 0.031062349 | 4.30E-22 |
| SS04276 | 0 | 0.001497232 | 4.35E-22 |
| SS04766 | 0 | 0.001497232 | 4.35E-22 |
| SS02762 | 0.069445668 | 0.092587288 | 4.46E-22 |
| SS14052 | 0.069445668 | 0.092587288 | 4.46E-22 |
| SS11079 | 0.024149428 | 0.056515466 | 4.73E-22 |
| SS11073 | 0 | 0.001952021 | 4.82E-22 |
| SS08417 | 0 | 0.001015021 | 5.33E-22 |
| SS12061 | 0.000155074 | 0.002081786 | 5.36E-22 |
| SS02390 | 0 | 0.000916781 | 5.37E-22 |
| SS00505 | 0 | 0.004936278 | 5.49E-22 |

| | | | |
|---|---|---|---|
| SS05813 | 0 | 0.004936278 | 5.49E-22 |
| SS05838 | 0 | 0.004936278 | 5.49E-22 |
| SS10492 | 0.009399795 | 0.015718648 | 5.50E-22 |
| SS10681 | 0.009399795 | 0.015718648 | 5.50E-22 |
| SS08413 | 4.10E-05 | 0.00302034 | 5.63E-22 |
| SS14463 | 4.10E-05 | 0.00302034 | 5.63E-22 |
| SS11130 | 0.016454478 | 0.034952611 | 5.79E-22 |
| SS14742 | 0.016454478 | 0.034952611 | 5.79E-22 |
| SS13359 | 0 | 0.00202421 | 5.87E-22 |
| SS01612 | 0.00152956 | 0.007475718 | 5.91E-22 |
| SS02814 | 0 | 0.001588156 | 6.05E-22 |
| SS13168 | 0 | 0.000842058 | 6.22E-22 |
| SS04014 | 0.008643577 | 0.024165845 | 6.31E-22 |
| SS12063 | 0.0007521 | 0.002501546 | 6.52E-22 |
| SS08412 | 0.006380224 | 0.012578331 | 8.11E-22 |
| SS14462 | 0.006380224 | 0.012578331 | 8.11E-22 |
| SS04254 | 2.76E-06 | 0.001300775 | 8.95E-22 |
| SS12927 | 0.000315677 | 0.001657273 | 9.35E-22 |
| SS15278 | 0.000315677 | 0.001657273 | 9.35E-22 |
| SS03191 | 0 | 0.000876755 | 9.70E-22 |
| SS01249 | 0.008183722 | 0.025150963 | 9.79E-22 |
| SS02487 | 0.008183722 | 0.025150963 | 9.79E-22 |
| SS03631 | 0 | 0.001987815 | 9.95E-22 |
| SS04559 | 0 | 0.001987815 | 9.95E-22 |
| SS04832 | 0 | 0.001987815 | 9.95E-22 |
| SS08967 | 0.001680505 | 0.008520259 | 1.03E-21 |
| SS09481 | 0.011722792 | 0.019582897 | 1.04E-21 |
| SS00839 | 0 | 0.002004471 | 1.06E-21 |
| SS02818 | 0 | 0.001349614 | 1.13E-21 |
| SS14747 | 1.37E-05 | 0.003416906 | 1.20E-21 |
| SS15230 | 0.002551743 | 0.007893691 | 1.22E-21 |
| SS01668 | 0.000261436 | 0.002439735 | 1.27E-21 |
| SS03054 | 0.000261436 | 0.002439735 | 1.27E-21 |
| SS14457 | 0.000261436 | 0.002439735 | 1.27E-21 |
| SS01627 | 0.074936668 | 0.108196025 | 1.28E-21 |
| SS02285 | 0.074936668 | 0.108196025 | 1.28E-21 |
| SS05334 | 0.074936668 | 0.108196025 | 1.28E-21 |
| SS05345 | 0.074936668 | 0.108196025 | 1.28E-21 |
| SS06908 | 0.074936668 | 0.108196025 | 1.28E-21 |
| SS08722 | 0.074936668 | 0.108196025 | 1.28E-21 |
| SS09908 | 0.074936668 | 0.108196025 | 1.28E-21 |
| SS11901 | 0.074936668 | 0.108196025 | 1.28E-21 |
| SS09592 | 0 | 0.003247944 | 1.29E-21 |
| SS10014 | 0 | 0.009493828 | 1.30E-21 |
| SS15501 | 0.000310631 | 0.002814716 | 1.35E-21 |
| SS02847 | 0 | 0.001142307 | 1.37E-21 |
| SS03669 | 0 | 0.002037355 | 1.37E-21 |

| | | | |
|---|---|---|---|
| SS07077 | 0 | 0.002037355 | 1.37E-21 |
| SS13356 | 0 | 0.002041814 | 1.38E-21 |
| SS05231 | 0.000296568 | 0.002157292 | 1.40E-21 |
| SS03243 | 0.000920915 | 0.004336325 | 1.43E-21 |
| SS03336 | 0.000920915 | 0.004336325 | 1.43E-21 |
| SS02009 | 7.35E-05 | 0.000994493 | 1.47E-21 |
| SS10512 | 0.006948437 | 0.013999764 | 1.50E-21 |
| SS10543 | 0.006948437 | 0.013999764 | 1.50E-21 |
| SS02840 | 0.001548508 | 0.006760093 | 1.54E-21 |
| SS10794 | 5.53E-06 | 0.000867492 | 1.55E-21 |
| SS00934 | 0.000417153 | 0.002055756 | 1.55E-21 |
| SS02680 | 0.000417153 | 0.002055756 | 1.55E-21 |
| SS15090 | 0 | 0.002190292 | 1.64E-21 |
| SS02591 | 0.00356248 | 0.013374505 | 1.68E-21 |
| SS02592 | 0.00356248 | 0.013374505 | 1.68E-21 |
| SS05977 | 0.00356248 | 0.013374505 | 1.68E-21 |
| SS02587 | 0.00356248 | 0.013374221 | 1.68E-21 |
| SS10763 | 0.00356248 | 0.013374221 | 1.68E-21 |
| SS04797 | 5.45E-06 | 0.001566124 | 1.71E-21 |
| SS01698 | 0.00102387 | 0.003789671 | 1.73E-21 |
| SS05572 | 2.73E-05 | 0.003309641 | 1.74E-21 |
| SS07592 | 0.01668054 | 0.026543891 | 1.78E-21 |
| SS03293 | 0 | 0.000968545 | 1.82E-21 |
| SS02550 | 3.55E-05 | 0.001164766 | 2.20E-21 |
| SS00868 | 2.68E-06 | 0.001190166 | 2.23E-21 |
| SS03192 | 2.68E-06 | 0.001190166 | 2.23E-21 |
| SS09918 | 0 | 0.001870369 | 2.65E-21 |
| SS03426 | 2.68E-06 | 0.00136032 | 2.92E-21 |
| SS12892 | 0 | 0.002647516 | 2.93E-21 |
| SS01797 | 0.013691526 | 0.02493473 | 3.14E-21 |
| SS06140 | 0.013691526 | 0.02493473 | 3.14E-21 |
| SS14931 | 0.013691526 | 0.02493473 | 3.14E-21 |
| SS08703 | 1.88E-05 | 0.001259895 | 3.43E-21 |
| SS08107 | 0.022281378 | 0.032138646 | 3.44E-21 |
| SS09085 | 0.022281378 | 0.032138646 | 3.44E-21 |
| SS09093 | 0.022281378 | 0.032138646 | 3.44E-21 |
| SS05462 | 0.00186422 | 0.00721255 | 3.64E-21 |
| SS05031 | 0 | 0.000734796 | 3.69E-21 |
| SS03021 | 0.002482843 | 0.014554779 | 3.84E-21 |
| SS04360 | 0 | 0.000592038 | 4.12E-21 |
| SS13133 | 0 | 0.000592038 | 4.12E-21 |
| SS10301 | 0.007575674 | 0.01794201 | 4.18E-21 |
| SS05010 | 0 | 0.000556981 | 4.21E-21 |
| SS10211 | 0.027782185 | 0.054373086 | 4.61E-21 |
| SS10324 | 0.027782185 | 0.054373086 | 4.61E-21 |
| SS08067 | 0 | 0.003463515 | 4.65E-21 |
| SS10446 | 0 | 0.003463515 | 4.65E-21 |

| | | | |
|---|---|---|---|
| SS05097 | 0 | 0.013229618 | 4.67E-21 |
| SS04274 | 0.000748773 | 0.002803684 | 4.86E-21 |
| SS04764 | 0.000748773 | 0.002803684 | 4.86E-21 |
| SS07575 | 0.000103436 | 0.000903328 | 5.06E-21 |
| SS14983 | 0.001031199 | 0.004593772 | 5.16E-21 |
| SS05795 | 0.001059886 | 0.005156411 | 5.30E-21 |
| SS02642 | 0.004330959 | 0.00892826 | 5.35E-21 |
| SS12291 | 0 | 0.00194069 | 5.35E-21 |
| SS15284 | 0 | 0.000887351 | 5.40E-21 |
| SS04110 | 0 | 0.000989457 | 5.72E-21 |
| SS01820 | 0.008270527 | 0.014489235 | 6.33E-21 |
| SS10468 | 0.008270527 | 0.014489235 | 6.33E-21 |
| SS03193 | 0 | 0.001032287 | 6.58E-21 |
| SS03242 | 0.001198776 | 0.006407878 | 6.96E-21 |
| SS03335 | 0.001198776 | 0.006407878 | 6.96E-21 |
| SS05268 | 0 | 0.001440631 | 7.49E-21 |
| SS09133 | 0.000267125 | 0.005552932 | 8.16E-21 |
| SS01816 | 0.054280097 | 0.076915375 | 8.57E-21 |
| SS09044 | 0.054280097 | 0.076915375 | 8.57E-21 |
| SS13121 | 0.054280097 | 0.076915375 | 8.57E-21 |
| SS14075 | 0.020726022 | 0.030603285 | 9.59E-21 |
| SS01773 | 0 | 0.000564063 | 1.00E-20 |
| SS05059 | 0 | 0.000564063 | 1.00E-20 |
| SS04702 | 0 | 0.004034521 | 1.03E-20 |
| SS00137 | 0 | 0.000665441 | 1.09E-20 |
| SS02371 | 0.01785219 | 0.025308418 | 1.18E-20 |
| SS03297 | 0.01785219 | 0.025308418 | 1.18E-20 |
| SS04032 | 0 | 0.000836027 | 1.19E-20 |
| SS02118 | 0 | 0.001231432 | 1.20E-20 |
| SS05573 | 2.46E-05 | 0.002971944 | 1.22E-20 |
| SS00656 | 0.013405567 | 0.032872727 | 1.36E-20 |
| SS02028 | 0.013405567 | 0.032872727 | 1.36E-20 |
| SS09936 | 0.013405567 | 0.032872727 | 1.36E-20 |
| SS07981 | 0 | 0.002224045 | 1.38E-20 |
| SS08873 | 0 | 0.002224045 | 1.38E-20 |
| SS11946 | 0 | 0.002224045 | 1.38E-20 |
| SS01672 | 0.000796007 | 0.004025546 | 1.43E-20 |
| SS00168 | 8.21E-06 | 0.003647393 | 1.44E-20 |
| SS01496 | 0 | 0.000744743 | 1.51E-20 |
| SS07532 | 0.000264441 | 0.0025744 | 1.52E-20 |
| SS05227 | 0.021566097 | 0.040561017 | 1.67E-20 |
| SS07905 | 0.021566097 | 0.040561017 | 1.67E-20 |
| SS14814 | 0.021566097 | 0.040561017 | 1.67E-20 |
| SS12738 | 0 | 0.000494746 | 1.77E-20 |
| SS12821 | 0 | 0.000494746 | 1.77E-20 |
| SS09288 | 0.004774917 | 0.019798372 | 1.82E-20 |
| SS13655 | 0.000147103 | 0.002293398 | 1.92E-20 |

| | | | |
|---|---|---|---|
| SS02000 | 2.76E-06 | 0.006224312 | 1.95E-20 |
| SS03171 | 2.76E-06 | 0.006224312 | 1.95E-20 |
| SS03484 | 2.76E-06 | 0.006224312 | 1.95E-20 |
| SS00303 | 0.000583204 | 0.003726369 | 1.96E-20 |
| SS01073 | 3.25E-05 | 0.001660104 | 1.97E-20 |
| SS05003 | 0 | 0.000821902 | 1.99E-20 |
| SS07547 | 0.002904354 | 0.008154823 | 2.00E-20 |
| SS13101 | 0.002904354 | 0.008154823 | 2.00E-20 |
| SS05483 | 0.001037852 | 0.00338728 | 2.06E-20 |
| SS02248 | 0.000356579 | 0.003019376 | 2.23E-20 |
| SS04058 | 0.000356579 | 0.003019376 | 2.23E-20 |
| SS00471 | 0.005844885 | 0.014097648 | 2.30E-20 |
| SS02393 | 0.005844885 | 0.014097648 | 2.30E-20 |
| SS10754 | 0 | 0.003062415 | 2.34E-20 |
| SS02211 | 0 | 0.001120879 | 2.34E-20 |
| SS08656 | 0 | 0.001122695 | 2.36E-20 |
| SS06247 | 0.000855938 | 0.003689788 | 2.44E-20 |
| SS10299 | 0.000855938 | 0.003689788 | 2.44E-20 |
| SS02331 | 0.002375197 | 0.008850119 | 2.46E-20 |
| SS06239 | 0.022900609 | 0.044520538 | 2.58E-20 |
| SS10291 | 0.022900609 | 0.044520538 | 2.58E-20 |
| SS15213 | 0 | 0.002394222 | 2.59E-20 |
| SS04277 | 0 | 0.00129623 | 2.70E-20 |
| SS04767 | 0 | 0.00129623 | 2.70E-20 |
| SS03108 | 0 | 0.001673895 | 2.73E-20 |
| SS06249 | 0 | 0.002313789 | 2.89E-20 |
| SS06277 | 0 | 0.002313789 | 2.89E-20 |
| SS11956 | 0 | 0.000529098 | 2.93E-20 |
| SS13288 | 0 | 0.000529098 | 2.93E-20 |
| SS05578 | 0 | 0.001534623 | 2.96E-20 |
| SS13552 | 6.04E-05 | 0.001679152 | 2.97E-20 |
| SS10302 | 1.09E-05 | 0.001810522 | 3.03E-20 |
| SS05116 | 2.45E-05 | 0.001852723 | 3.04E-20 |
| SS08388 | 2.45E-05 | 0.001852723 | 3.04E-20 |
| SS08749 | 2.45E-05 | 0.001852723 | 3.04E-20 |
| SS13150 | 2.45E-05 | 0.001852723 | 3.04E-20 |
| SS15042 | 0 | 0.0011376 | 3.28E-20 |
| SS07765 | 0.000149947 | 0.006422393 | 3.32E-20 |
| SS05692 | 0.00214028 | 0.004876327 | 3.32E-20 |
| SS08718 | 0.00214028 | 0.004876327 | 3.32E-20 |
| SS15025 | 0.00214028 | 0.004876327 | 3.32E-20 |
| SS15032 | 0.00214028 | 0.004876327 | 3.32E-20 |
| SS00861 | 0 | 0.00119677 | 3.55E-20 |
| SS00493 | 8.29E-06 | 0.000714935 | 3.58E-20 |
| SS04670 | 0.020109854 | 0.035543062 | 3.74E-20 |
| SS11806 | 0.020109854 | 0.035543062 | 3.74E-20 |
| SS04619 | 6.27E-05 | 0.004132108 | 3.80E-20 |

| | | | |
|---|---|---|---|
| SS02395 | 0 | 0.001013486 | 3.84E-20 |
| SS05841 | 0.008013415 | 0.014188387 | 4.03E-20 |
| SS03661 | 0 | 0.002471908 | 4.17E-20 |
| SS10650 | 0.00840304 | 0.014021794 | 4.47E-20 |
| SS06595 | 0 | 0.001371002 | 5.08E-20 |
| SS01907 | 2.72E-05 | 0.001067102 | 5.08E-20 |
| SS02648 | 0.070655134 | 0.097696627 | 5.08E-20 |
| SS06541 | 0.070655134 | 0.097696627 | 5.08E-20 |
| SS15542 | 0 | 0.000819542 | 5.61E-20 |
| SS05566 | 0.000223378 | 0.002170219 | 5.67E-20 |
| SS10042 | 0 | 0.001104226 | 5.78E-20 |
| SS02812 | 0 | 0.001237801 | 5.84E-20 |
| SS08233 | 0.018597797 | 0.05103519 | 6.14E-20 |
| SS11169 | 0.018597797 | 0.05103519 | 6.14E-20 |
| SS15400 | 0.018597797 | 0.05103519 | 6.14E-20 |
| SS00891 | 0 | 0.003000862 | 6.24E-20 |
| SS05564 | 0.000656876 | 0.003880095 | 6.60E-20 |
| SS02319 | 0.011353997 | 0.023579418 | 7.45E-20 |
| SS05700 | 0.011353997 | 0.023579418 | 7.45E-20 |
| SS10749 | 0.011353997 | 0.023579418 | 7.45E-20 |
| SS04496 | 0 | 0.000437635 | 7.65E-20 |
| SS13163 | 0 | 0.001037913 | 7.75E-20 |
| SS06297 | 0 | 0.003946045 | 7.76E-20 |
| SS06298 | 0 | 0.003946045 | 7.76E-20 |
| SS14886 | 0.002369749 | 0.006848512 | 7.82E-20 |
| SS02713 | 0.020256073 | 0.030491904 | 8.04E-20 |
| SS03560 | 0.020256073 | 0.030491904 | 8.04E-20 |
| SS06945 | 0.020256073 | 0.030491904 | 8.04E-20 |
| SS09944 | 0.020256073 | 0.030491904 | 8.04E-20 |
| SS13654 | 0.000734952 | 0.003680051 | 8.87E-20 |
| SS01872 | 0 | 0.001223391 | 9.53E-20 |
| SS13855 | 0 | 0.001223391 | 9.53E-20 |
| SS02485 | 0.000665329 | 0.003205866 | 1.11E-19 |
| SS09075 | 0.000665329 | 0.003205866 | 1.11E-19 |
| SS00069 | 4.37E-05 | 0.000728905 | 1.20E-19 |
| SS05501 | 4.37E-05 | 0.000728905 | 1.20E-19 |
| SS13726 | 0 | 0.000857807 | 1.22E-19 |
| SS04603 | 0.007553078 | 0.012071378 | 1.22E-19 |
| SS08872 | 0 | 0.002115871 | 1.26E-19 |
| SS11943 | 0 | 0.002115871 | 1.26E-19 |
| SS07611 | 0 | 0.001746889 | 1.26E-19 |
| SS07640 | 0 | 0.001746889 | 1.26E-19 |
| SS06293 | 0 | 0.00387823 | 1.29E-19 |
| SS06294 | 0 | 0.00387823 | 1.29E-19 |
| SS07527 | 0.008458291 | 0.020564712 | 1.29E-19 |
| SS07543 | 0.008458291 | 0.020564712 | 1.29E-19 |
| SS05658 | 0.000621663 | 0.002776582 | 1.32E-19 |

| | | | |
|---|---|---|---|
| SS03339 | 0.00097011 | 0.005377765 | 1.37E-19 |
| SS02437 | 0.01184185 | 0.024135051 | 1.38E-19 |
| SS08542 | 0.01184185 | 0.024135051 | 1.38E-19 |
| SS06516 | 0 | 0.000787483 | 1.38E-19 |
| SS03598 | 0 | 0.003584519 | 1.51E-19 |
| SS07610 | 0 | 0.004150775 | 1.56E-19 |
| SS07639 | 0 | 0.004150775 | 1.56E-19 |
| SS09487 | 0.000214764 | 0.003130222 | 1.57E-19 |
| SS00776 | 0.02283921 | 0.043634416 | 1.65E-19 |
| SS10167 | 0.02283921 | 0.043634416 | 1.65E-19 |
| SS05563 | 0.000427407 | 0.005045305 | 1.69E-19 |
| SS00450 | 0.001029077 | 0.004662494 | 1.69E-19 |
| SS01436 | 0 | 0.00149712 | 1.73E-19 |
| SS12962 | 0 | 0.00149712 | 1.73E-19 |
| SS13523 | 5.45E-06 | 0.001606243 | 1.76E-19 |
| SS11127 | 0.00114005 | 0.005998269 | 1.83E-19 |
| SS11071 | 1.63E-05 | 0.001001251 | 1.84E-19 |
| SS02447 | 0 | 0.001083098 | 1.89E-19 |
| SS03288 | 0 | 0.001083098 | 1.89E-19 |
| SS13727 | 0 | 0.000891115 | 1.92E-19 |
| SS03515 | 0.030253903 | 0.049317383 | 1.92E-19 |
| SS14481 | 0.030253903 | 0.049317383 | 1.92E-19 |
| SS09134 | 0.001674976 | 0.005767262 | 2.06E-19 |
| SS02860 | 0 | 0.001223467 | 2.07E-19 |
| SS04107 | 0 | 0.002565512 | 2.13E-19 |
| SS11008 | 2.72E-05 | 0.000662899 | 2.17E-19 |
| SS02868 | 0.0002777 | 0.001584704 | 2.21E-19 |
| SS13728 | 1.10E-05 | 0.001283913 | 2.22E-19 |
| SS04077 | 0 | 0.001550545 | 2.24E-19 |
| SS04490 | 3.85E-05 | 0.002525836 | 2.44E-19 |
| SS09157 | 0 | 0.000550598 | 2.77E-19 |
| SS04123 | 0.020828977 | 0.034743068 | 2.90E-19 |
| SS06693 | 0.020828977 | 0.034743068 | 2.90E-19 |
| SS03700 | 0 | 0.000628243 | 2.96E-19 |
| SS10500 | 0 | 0.003584246 | 3.00E-19 |
| SS05447 | 0.009219681 | 0.021829738 | 3.07E-19 |
| SS10031 | 0 | 0.002028412 | 3.14E-19 |
| SS00598 | 0.001757262 | 0.004980595 | 3.15E-19 |
| SS09499 | 0.001757262 | 0.004980595 | 3.15E-19 |
| SS13225 | 7.07E-05 | 0.048490059 | 3.28E-19 |
| SS11144 | 0.001889178 | 0.004717083 | 3.29E-19 |
| SS15096 | 0.001889178 | 0.004717083 | 3.29E-19 |
| SS02124 | 0.009152375 | 0.01562137 | 3.33E-19 |
| SS06745 | 0.009152375 | 0.01562137 | 3.33E-19 |
| SS01389 | 0 | 0.000577235 | 3.40E-19 |
| SS04492 | 0 | 0.001053287 | 3.43E-19 |
| SS04190 | 0.054245607 | 0.080125698 | 3.44E-19 |

| | | | |
|---|---|---|---|
| SS13254 | 0.054245607 | 0.080125698 | 3.44E-19 |
| SS11733 | 0 | 0.000478933 | 3.44E-19 |
| SS01740 | 0.017373787 | 0.026052705 | 3.46E-19 |
| SS06906 | 0.017373787 | 0.026052705 | 3.46E-19 |
| SS15085 | 0.017373787 | 0.026052705 | 3.46E-19 |
| SS06674 | 0.000604676 | 0.00240825 | 3.46E-19 |
| SS11044 | 0 | 0.002587342 | 3.85E-19 |
| SS13382 | 0.000811709 | 0.006415003 | 4.09E-19 |
| SS07629 | 0 | 0.002067044 | 4.20E-19 |
| SS14097 | 0 | 0.000601113 | 4.27E-19 |
| SS10470 | 5.16E-05 | 0.004389496 | 4.29E-19 |
| SS07818 | 0 | 0.00106446 | 4.49E-19 |
| SS02544 | 0.001073627 | 0.003204398 | 4.63E-19 |
| SS02129 | 0.017149927 | 0.02504105 | 4.90E-19 |
| SS02434 | 0.017149927 | 0.02504105 | 4.90E-19 |
| SS11439 | 0.017149927 | 0.02504105 | 4.90E-19 |
| SS10025 | 0 | 0.000743102 | 4.91E-19 |
| SS11962 | 0.060603086 | 0.082599148 | 5.04E-19 |
| SS12893 | 0.060603086 | 0.082599148 | 5.04E-19 |
| SS13289 | 0.060603086 | 0.082599148 | 5.04E-19 |
| SS04934 | 0 | 0.000913487 | 5.46E-19 |
| SS01669 | 0.000569061 | 0.004434623 | 5.48E-19 |
| SS01947 | 0.000569061 | 0.004434623 | 5.48E-19 |
| SS03060 | 0.000569061 | 0.004434623 | 5.48E-19 |
| SS14459 | 0.000569061 | 0.004434623 | 5.48E-19 |
| SS00548 | 0 | 0.001062765 | 5.50E-19 |
| SS06677 | 0 | 0.001062765 | 5.50E-19 |
| SS15212 | 0 | 0.004100547 | 5.78E-19 |
| SS15222 | 0 | 0.004100547 | 5.78E-19 |
| SS11095 | 0 | 0.001386847 | 5.79E-19 |
| SS04040 | 5.37E-06 | 0.000747222 | 5.89E-19 |
| SS06733 | 0 | 0.002847463 | 5.98E-19 |
| SS13209 | 0 | 0.000358042 | 6.43E-19 |
| SS12325 | 2.68E-06 | 0.004090773 | 6.64E-19 |
| SS12885 | 0 | 0.003208384 | 7.55E-19 |
| SS04971 | 0 | 0.00334809 | 7.59E-19 |
| SS06161 | 0 | 0.00334809 | 7.59E-19 |
| SS04710 | 0 | 0.002536477 | 7.68E-19 |
| SS11084 | 0.027308233 | 0.039679014 | 7.86E-19 |
| SS10040 | 2.76E-06 | 0.002011651 | 8.02E-19 |
| SS00075 | 0.000149706 | 0.001875143 | 8.14E-19 |
| SS05188 | 0.000149706 | 0.001875143 | 8.14E-19 |
| SS08868 | 0.000149706 | 0.001875143 | 8.14E-19 |
| SS11918 | 0.000149706 | 0.001875143 | 8.14E-19 |
| SS03667 | 0 | 0.001442689 | 8.64E-19 |
| SS07075 | 0 | 0.001442689 | 8.64E-19 |
| SS10483 | 0.044120713 | 0.064075443 | 9.02E-19 |

| | | | |
|---|---|---|---|
| SS12596 | 0.044120713 | 0.064075443 | 9.02E-19 |
| SS14375 | 1.92E-05 | 0.000787552 | 9.03E-19 |
| SS06989 | 0.005542031 | 0.008739763 | 9.41E-19 |
| SS11432 | 0.005542031 | 0.008739763 | 9.41E-19 |
| SS01325 | 0.03010743 | 0.042227382 | 9.52E-19 |
| SS14492 | 0.03010743 | 0.042227382 | 9.52E-19 |
| SS14889 | 0.03010743 | 0.042227382 | 9.52E-19 |
| SS05525 | 0 | 0.002722415 | 9.83E-19 |
| SS15387 | 0 | 0.002722415 | 9.83E-19 |
| SS03199 | 0.003672925 | 0.008411436 | 1.06E-18 |
| SS06892 | 0.003672925 | 0.008411436 | 1.06E-18 |
| SS11867 | 0.003672925 | 0.008411436 | 1.06E-18 |
| SS13148 | 0 | 0.003532387 | 1.12E-18 |
| SS09914 | 0 | 0.001255545 | 1.13E-18 |
| SS11158 | 0.017491642 | 0.048105579 | 1.15E-18 |
| SS11179 | 0.017491642 | 0.048105579 | 1.15E-18 |
| SS02186 | 0.033743571 | 0.048467204 | 1.16E-18 |
| SS06271 | 0.033743571 | 0.048467204 | 1.16E-18 |
| SS10178 | 0.033743571 | 0.048467204 | 1.16E-18 |
| SS07156 | 0.000520348 | 0.001724277 | 1.16E-18 |
| SS04878 | 0 | 0.001715135 | 1.19E-18 |
| SS13901 | 0 | 0.001715135 | 1.19E-18 |
| SS14112 | 1.92E-05 | 0.001189511 | 1.20E-18 |
| SS14981 | 0.00104166 | 0.0056737 | 1.28E-18 |
| SS02829 | 3.04E-05 | 0.001749158 | 1.31E-18 |
| SS09919 | 0.000985812 | 0.003540106 | 1.32E-18 |
| SS01450 | 0 | 0.000631536 | 1.34E-18 |
| SS13051 | 0 | 0.000631536 | 1.34E-18 |
| SS04611 | 0.016632986 | 0.028848813 | 1.39E-18 |
| SS01802 | 0.030453595 | 0.040142892 | 1.44E-18 |
| SS04998 | 0.030453595 | 0.040142892 | 1.44E-18 |
| SS06150 | 0.030453595 | 0.040142892 | 1.44E-18 |
| SS06180 | 0.030453595 | 0.040142892 | 1.44E-18 |
| SS15408 | 0.030453595 | 0.040142892 | 1.44E-18 |
| SS04209 | 5.37E-06 | 0.000736506 | 1.46E-18 |
| SS15215 | 0 | 0.001195421 | 1.49E-18 |
| SS01428 | 2.17E-05 | 0.001153916 | 1.51E-18 |
| SS00901 | 0.002857121 | 0.011867295 | 1.51E-18 |
| SS02464 | 0 | 0.00072862 | 1.52E-18 |
| SS01028 | 1.09E-05 | 0.005521338 | 1.53E-18 |
| SS07516 | 0.000947675 | 0.002544601 | 1.59E-18 |
| SS01724 | 0.002754292 | 0.006141237 | 1.66E-18 |
| SS06902 | 0.002754292 | 0.006141237 | 1.66E-18 |
| SS06911 | 0.002754292 | 0.006141237 | 1.66E-18 |
| SS07178 | 0.002754292 | 0.006141237 | 1.66E-18 |
| SS15074 | 0.002754292 | 0.006141237 | 1.66E-18 |
| SS04590 | 2.68E-06 | 0.00089571 | 1.78E-18 |

| | | | |
|---|---|---|---|
| SS03073 | 0.002758215 | 0.005661848 | 1.85E-18 |
| SS08686 | 0.002758215 | 0.005661848 | 1.85E-18 |
| SS13144 | 0.000373085 | 0.00291501 | 1.86E-18 |
| SS15316 | 0.000373085 | 0.00291501 | 1.86E-18 |
| SS04256 | 0.089937266 | 0.12217307 | 1.88E-18 |
| SS07646 | 0.089937266 | 0.12217307 | 1.88E-18 |
| SS00365 | 0.080895439 | 0.108537892 | 1.89E-18 |
| SS01015 | 0.080895439 | 0.108537892 | 1.89E-18 |
| SS02659 | 0.080895439 | 0.108537892 | 1.89E-18 |
| SS02739 | 0.080895439 | 0.108537892 | 1.89E-18 |
| SS03579 | 0.080895439 | 0.108537892 | 1.89E-18 |
| SS06870 | 0.080895439 | 0.108537892 | 1.89E-18 |
| SS07827 | 0.080895439 | 0.108537892 | 1.89E-18 |
| SS08220 | 0.080895439 | 0.108537892 | 1.89E-18 |
| SS14151 | 0.080895439 | 0.108537892 | 1.89E-18 |
| SS14209 | 0.080895439 | 0.108537892 | 1.89E-18 |
| SS14296 | 0.080895439 | 0.108537892 | 1.89E-18 |
| SS15371 | 0.080895439 | 0.108537892 | 1.89E-18 |
| SS03340 | 0.000940907 | 0.005371621 | 1.90E-18 |
| SS06729 | 0 | 0.003581393 | 1.99E-18 |
| SS02320 | 1.10E-05 | 0.00089659 | 2.00E-18 |
| SS11139 | 0.000266884 | 0.002410183 | 2.02E-18 |
| SS11444 | 0.011808484 | 0.01789361 | 2.04E-18 |
| SS04124 | 0.011911278 | 0.028173358 | 2.09E-18 |
| SS06694 | 0.011911278 | 0.028173358 | 2.09E-18 |
| SS11730 | 0 | 0.000859126 | 2.17E-18 |
| SS08604 | 0 | 0.00061978 | 2.79E-18 |
| SS09097 | 0 | 0.000644668 | 2.88E-18 |
| SS00225 | 0.001441424 | 0.003333752 | 2.94E-18 |
| SS08485 | 0.006906984 | 0.01605156 | 3.08E-18 |
| SS09406 | 2.72E-05 | 0.001042201 | 3.14E-18 |
| SS11912 | 2.68E-06 | 0.000641309 | 3.16E-18 |
| SS03665 | 0 | 0.000729989 | 3.23E-18 |
| SS01365 | 0.039335955 | 0.05123715 | 3.25E-18 |
| SS04622 | 0.039335955 | 0.05123715 | 3.25E-18 |
| SS02212 | 5.45E-06 | 0.001335315 | 3.27E-18 |
| SS05269 | 0.03736012 | 0.055171056 | 3.28E-18 |
| SS13956 | 0 | 0.000898542 | 3.52E-18 |
| SS07630 | 0 | 0.001005068 | 3.54E-18 |
| SS13363 | 0 | 0.001070867 | 3.61E-18 |
| SS03596 | 0.05201265 | 0.071511057 | 3.65E-18 |
| SS10177 | 0.05201265 | 0.071511057 | 3.65E-18 |
| SS12031 | 0.05201265 | 0.071511057 | 3.65E-18 |
| SS14855 | 0.05201265 | 0.071511057 | 3.65E-18 |
| SS00514 | 0.007919464 | 0.017125448 | 3.79E-18 |
| SS04090 | 0 | 0.001635964 | 3.96E-18 |
| SS01196 | 0.023196237 | 0.042451549 | 4.00E-18 |

| | | | |
|---|---|---|---|
| SS01260 | 0.023196237 | 0.042451549 | 4.00E-18 |
| SS02757 | 0.023196237 | 0.042451549 | 4.00E-18 |
| SS03568 | 0.023196237 | 0.042451549 | 4.00E-18 |
| SS02469 | 0 | 0.001410719 | 4.05E-18 |
| SS00419 | 0.028443465 | 0.085377387 | 4.19E-18 |
| SS05720 | 0.028443465 | 0.085377387 | 4.19E-18 |
| SS12729 | 0.028443465 | 0.085377387 | 4.19E-18 |
| SS13146 | 0.028443465 | 0.085377387 | 4.19E-18 |
| SS01117 | 1.64E-05 | 0.000630911 | 4.21E-18 |
| SS01897 | 0 | 0.000600279 | 4.42E-18 |
| SS01902 | 0 | 0.000600279 | 4.42E-18 |
| SS07066 | 0 | 0.001395537 | 4.53E-18 |
| SS02055 | 0.004714619 | 0.008879525 | 4.63E-18 |
| SS02895 | 0.004714619 | 0.008879525 | 4.63E-18 |
| SS10356 | 0.004714619 | 0.008879525 | 4.63E-18 |
| SS05910 | 0.011472768 | 0.016705398 | 4.64E-18 |
| SS15417 | 0.011472768 | 0.016705398 | 4.64E-18 |
| SS01821 | 0.044837588 | 0.247847001 | 4.82E-18 |
| SS06741 | 0.044837588 | 0.247847001 | 4.82E-18 |
| SS02022 | 0 | 0.000791602 | 5.15E-18 |
| SS02041 | 0.003602098 | 0.011436503 | 5.22E-18 |
| SS10346 | 0.003602098 | 0.011436503 | 5.22E-18 |
| SS02421 | 0.002086245 | 0.012283469 | 5.76E-18 |
| SS08208 | 0.002446105 | 0.027860131 | 5.79E-18 |
| SS05788 | 0.000288436 | 0.002099523 | 5.82E-18 |
| SS00080 | 0.030723358 | 0.047109462 | 5.98E-18 |
| SS13278 | 0.030723358 | 0.047109462 | 5.98E-18 |
| SS06226 | 0.003175929 | 0.043569334 | 6.00E-18 |
| SS04202 | 0.027578042 | 0.048715906 | 6.21E-18 |
| SS13268 | 0.027578042 | 0.048715906 | 6.21E-18 |
| SS12326 | 0.000278102 | 0.002885307 | 6.31E-18 |
| SS06157 | 0 | 0.000902973 | 6.46E-18 |
| SS04546 | 0.000283711 | 0.001696179 | 6.57E-18 |
| SS11882 | 0.037281689 | 0.071645175 | 6.65E-18 |
| SS10056 | 0 | 0.000735561 | 6.75E-18 |
| SS11105 | 0 | 0.000735561 | 6.75E-18 |
| SS13452 | 0 | 0.007232448 | 6.79E-18 |
| SS11125 | 0.0005284 | 0.001346558 | 6.90E-18 |
| SS00551 | 0.001806698 | 0.005471881 | 7.23E-18 |
| SS15300 | 5.53E-06 | 0.001473031 | 7.26E-18 |
| SS02823 | 0 | 0.000793651 | 7.44E-18 |
| SS00726 | 0.000103115 | 0.001138209 | 8.10E-18 |
| SS14199 | 0.000103115 | 0.001138209 | 8.10E-18 |
| SS14200 | 0 | 0.000594694 | 8.19E-18 |
| SS00569 | 0 | 0.000794675 | 8.20E-18 |
| SS02599 | 0.006110014 | 0.013658543 | 8.57E-18 |
| SS03710 | 1.36E-05 | 0.000516049 | 8.59E-18 |

| | | | |
|---|---|---|---|
| SS00438 | 0.010088603 | 0.01935969 | 8.63E-18 |
| SS03563 | 0 | 0.001230899 | 8.64E-18 |
| SS14145 | 0 | 0.001230899 | 8.64E-18 |
| SS06574 | 0 | 0.000967283 | 1.01E-17 |
| SS01020 | 0.026695139 | 0.034413083 | 1.05E-17 |
| SS14047 | 0.026695139 | 0.034413083 | 1.05E-17 |
| SS10488 | 0 | 0.000381959 | 1.05E-17 |
| SS07897 | 0 | 0.000533244 | 1.06E-17 |
| SS10005 | 0.001574625 | 0.005034748 | 1.10E-17 |
| SS13166 | 0 | 0.000714279 | 1.19E-17 |
| SS01213 | 0.000144339 | 0.001933571 | 1.19E-17 |
| SS02543 | 0.000144339 | 0.001933571 | 1.19E-17 |
| SS10566 | 0 | 0.001733619 | 1.19E-17 |
| SS11067 | 0 | 0.000539449 | 1.22E-17 |
| SS06650 | 9.25E-05 | 0.017781686 | 1.24E-17 |
| SS13123 | 0 | 0.002558725 | 1.25E-17 |
| SS13960 | 0 | 0.002558725 | 1.25E-17 |
| SS15276 | 0 | 0.002558725 | 1.25E-17 |
| SS04621 | 0.053564197 | 0.080105621 | 1.26E-17 |
| SS04709 | 0 | 0.001496668 | 1.28E-17 |
| SS01277 | 0.008889586 | 0.016520304 | 1.31E-17 |
| SS08193 | 0 | 0.000745429 | 1.31E-17 |
| SS11219 | 0.008346034 | 0.014936612 | 1.35E-17 |
| SS11428 | 0.008346034 | 0.014936612 | 1.35E-17 |
| SS02254 | 0.049148715 | 0.248484861 | 1.36E-17 |
| SS00089 | 0.022719888 | 0.031579988 | 1.47E-17 |
| SS11963 | 0.022719888 | 0.031579988 | 1.47E-17 |
| SS12896 | 0.022719888 | 0.031579988 | 1.47E-17 |
| SS13290 | 0.022719888 | 0.031579988 | 1.47E-17 |
| SS15419 | 0.022719888 | 0.031579988 | 1.47E-17 |
| SS02007 | 2.76E-06 | 0.000851737 | 1.48E-17 |
| SS02008 | 2.76E-06 | 0.000851737 | 1.48E-17 |
| SS00319 | 0.02641964 | 0.035916414 | 1.52E-17 |
| SS04250 | 0.02641964 | 0.035916414 | 1.52E-17 |
| SS09040 | 0.02641964 | 0.035916414 | 1.52E-17 |
| SS15415 | 0.02641964 | 0.035916414 | 1.52E-17 |
| SS03419 | 0 | 0.000624535 | 1.56E-17 |
| SS11043 | 0 | 0.002784195 | 1.58E-17 |
| SS11795 | 0 | 0.00306578 | 1.61E-17 |
| SS01618 | 0.015932845 | 0.021406584 | 1.63E-17 |
| SS02176 | 0.015932845 | 0.021406584 | 1.63E-17 |
| SS02478 | 0.015932845 | 0.021406584 | 1.63E-17 |
| SS15433 | 0.015932845 | 0.021406584 | 1.63E-17 |
| SS05457 | 8.69E-05 | 0.000813931 | 1.66E-17 |
| SS08910 | 0.009609868 | 0.017484573 | 1.67E-17 |
| SS02334 | 0.000257788 | 0.002937533 | 1.92E-17 |
| SS13387 | 0 | 0.000456906 | 1.95E-17 |

| | | | |
|---|---|---|---|
| SS04091 | 0 | 0.002401963 | 2.09E-17 |
| SS06713 | 0.003285697 | 0.006314202 | 2.12E-17 |
| SS12342 | 0 | 0.000865926 | 2.17E-17 |
| SS10712 | 0.000187844 | 0.001370673 | 2.18E-17 |
| SS05885 | 0.001186515 | 0.005561154 | 2.29E-17 |
| SS13878 | 0.001719205 | 0.007605238 | 2.33E-17 |
| SS02439 | 0 | 0.001139774 | 2.66E-17 |
| SS05002 | 0 | 0.000505194 | 2.68E-17 |
| SS02672 | 0.011189826 | 0.022924167 | 2.71E-17 |
| SS04117 | 0.023357494 | 0.03240873 | 2.72E-17 |
| SS06041 | 0.023357494 | 0.03240873 | 2.72E-17 |
| SS08119 | 0.023357494 | 0.03240873 | 2.72E-17 |
| SS08150 | 0.023357494 | 0.03240873 | 2.72E-17 |
| SS03562 | 2.76E-06 | 0.001981303 | 2.74E-17 |
| SS14141 | 2.76E-06 | 0.001981303 | 2.74E-17 |
| SS06985 | 0 | 0.001088708 | 2.83E-17 |
| SS10757 | 0 | 0.001088708 | 2.83E-17 |
| SS13447 | 0 | 0.000542964 | 2.84E-17 |
| SS02458 | 0.000185883 | 0.009353651 | 2.89E-17 |
| SS02673 | 0.008818346 | 0.017919276 | 3.02E-17 |
| SS10660 | 0 | 0.001001345 | 3.04E-17 |
| SS06924 | 0.002417143 | 0.007086678 | 3.09E-17 |
| SS15088 | 0.002417143 | 0.007086678 | 3.09E-17 |
| SS14105 | 0.068042783 | 0.089419432 | 3.09E-17 |
| SS01508 | 5.75E-05 | 0.000904964 | 3.25E-17 |
| SS03975 | 0.012457537 | 0.019385415 | 3.38E-17 |
| SS04607 | 0.012457537 | 0.019385415 | 3.38E-17 |
| SS13502 | 0.015466498 | 0.022925896 | 3.50E-17 |
| SS12384 | 0.000464982 | 0.002607325 | 3.52E-17 |
| SS02123 | 0.000927682 | 0.003686923 | 3.69E-17 |
| SS10668 | 0.000927682 | 0.003686923 | 3.69E-17 |
| SS07198 | 0.002898195 | 0.014266129 | 3.76E-17 |
| SS07222 | 0.002898195 | 0.014266129 | 3.76E-17 |
| SS12938 | 0.002898195 | 0.014266129 | 3.76E-17 |
| SS15266 | 0.002898195 | 0.014266129 | 3.76E-17 |
| SS02335 | 0.002271198 | 0.009406806 | 3.81E-17 |
| SS05842 | 0.002271198 | 0.009406806 | 3.81E-17 |
| SS07578 | 0 | 0.001086097 | 3.83E-17 |
| SS03541 | 0 | 0.001033404 | 4.16E-17 |
| SS03608 | 0 | 0.001033404 | 4.16E-17 |
| SS05209 | 0 | 0.001033404 | 4.16E-17 |
| SS05511 | 0 | 0.001033404 | 4.16E-17 |
| SS08060 | 0 | 0.001033404 | 4.16E-17 |
| SS00995 | 0.032259215 | 0.04406302 | 4.29E-17 |
| SS04232 | 0.032259215 | 0.04406302 | 4.29E-17 |
| SS15359 | 0.032259215 | 0.04406302 | 4.29E-17 |
| SS02056 | 0.000160523 | 0.00117907 | 4.62E-17 |

| | | | |
|---|---|---|---|
| SS10357 | 0.000160523 | 0.00117907 | 4.62E-17 |
| SS07758 | 0.002040089 | 0.013530156 | 4.70E-17 |
| SS12879 | 0.001722083 | 0.006756465 | 4.88E-17 |
| SS08383 | 1.09E-05 | 0.00047772 | 4.97E-17 |
| SS00123 | 0.029180057 | 0.044959332 | 5.14E-17 |
| SS11747 | 8.21E-06 | 0.000379403 | 5.60E-17 |
| SS07935 | 0 | 0.000430315 | 5.71E-17 |
| SS10067 | 0 | 0.000430315 | 5.71E-17 |
| SS02302 | 0.024417241 | 0.04116087 | 5.87E-17 |
| SS03784 | 0.024417241 | 0.04116087 | 5.87E-17 |
| SS04542 | 0.024417241 | 0.04116087 | 5.87E-17 |
| SS04572 | 0.024417241 | 0.04116087 | 5.87E-17 |
| SS04581 | 0.024417241 | 0.04116087 | 5.87E-17 |
| SS04675 | 0.024417241 | 0.04116087 | 5.87E-17 |
| SS15224 | 0 | 0.000547281 | 6.27E-17 |
| SS08670 | 0 | 0.000337722 | 6.60E-17 |
| SS12344 | 0 | 0.000663517 | 6.91E-17 |
| SS00185 | 0 | 0.001234438 | 7.00E-17 |
| SS00603 | 0 | 0.001234438 | 7.00E-17 |
| SS07266 | 0.009209748 | 0.014608702 | 7.46E-17 |
| SS07310 | 0.009209748 | 0.014608702 | 7.46E-17 |
| SS00818 | 5.45E-05 | 0.001163889 | 7.51E-17 |
| SS05193 | 0.01943583 | 0.026224694 | 7.79E-17 |
| SS08870 | 0.01943583 | 0.026224694 | 7.79E-17 |
| SS11938 | 0.01943583 | 0.026224694 | 7.79E-17 |
| SS00183 | 0 | 0.000332459 | 8.10E-17 |
| SS00601 | 0 | 0.000332459 | 8.10E-17 |
| SS01747 | 0 | 0.000951681 | 8.12E-17 |
| SS10713 | 0.001157473 | 0.003155934 | 8.30E-17 |
| SS10755 | 0 | 0.00343957 | 8.50E-17 |
| SS05772 | 0 | 0.001586314 | 8.64E-17 |
| SS00652 | 0 | 0.001732852 | 8.93E-17 |
| SS03553 | 0 | 0.001732852 | 8.93E-17 |
| SS01814 | 0 | 0.00067942 | 9.25E-17 |
| SS10793 | 0 | 0.001021767 | 9.37E-17 |
| SS11796 | 0 | 0.001026997 | 9.44E-17 |
| SS08689 | 0 | 0.000241735 | 9.54E-17 |
| SS01183 | 0 | 0.000576453 | 9.85E-17 |
| SS06894 | 0.004980218 | 0.008159911 | 1.01E-16 |
| SS03668 | 0 | 0.000473746 | 1.02E-16 |
| SS07076 | 0 | 0.000473746 | 1.02E-16 |
| SS08508 | 0 | 0.00098742 | 1.03E-16 |
| SS00304 | 0 | 0.000866155 | 1.05E-16 |
| SS03799 | 0 | 0.002252833 | 1.06E-16 |
| SS01202 | 0.024062279 | 0.032244491 | 1.07E-16 |
| SS02610 | 0.024062279 | 0.032244491 | 1.07E-16 |
| SS11703 | 0.024062279 | 0.032244491 | 1.07E-16 |

| | | | |
|---|---|---|---|
| SS04678 | 0 | 0.000535168 | 1.08E-16 |
| SS05889 | 0.000512617 | 0.003959931 | 1.11E-16 |
| SS02513 | 0.000471314 | 0.001765315 | 1.11E-16 |
| SS13998 | 0.001720202 | 0.004446542 | 1.13E-16 |
| SS05900 | 0 | 0.001518944 | 1.20E-16 |
| SS15394 | 0 | 0.001518944 | 1.20E-16 |
| SS03549 | 0.008696729 | 0.013410627 | 1.22E-16 |
| SS03601 | 0.008696729 | 0.013410627 | 1.22E-16 |
| SS05229 | 0.008696729 | 0.013410627 | 1.22E-16 |
| SS07908 | 0.008696729 | 0.013410627 | 1.22E-16 |
| SS14816 | 0.008696729 | 0.013410627 | 1.22E-16 |
| SS06092 | 0.068288253 | 0.091540992 | 1.25E-16 |
| SS10689 | 0.068288253 | 0.091540992 | 1.25E-16 |
| SS13530 | 0.068288253 | 0.091540992 | 1.25E-16 |
| SS15237 | 0.068288253 | 0.091540992 | 1.25E-16 |
| SS02419 | 0.001392825 | 0.006080184 | 1.26E-16 |
| SS08691 | 0.001392825 | 0.006080184 | 1.26E-16 |
| SS03037 | 0.022364834 | 0.03412869 | 1.37E-16 |
| SS03753 | 0.022364834 | 0.03412869 | 1.37E-16 |
| SS15365 | 0.022364834 | 0.03412869 | 1.37E-16 |
| SS02605 | 0.004904631 | 0.011007415 | 1.39E-16 |
| SS11431 | 0.004904631 | 0.011007415 | 1.39E-16 |
| SS02131 | 0.015506322 | 0.0250224 | 1.40E-16 |
| SS02436 | 0.015506322 | 0.0250224 | 1.40E-16 |
| SS13736 | 0 | 0.000658393 | 1.50E-16 |
| SS09816 | 0 | 0.001213276 | 1.53E-16 |
| SS14353 | 0 | 0.001213276 | 1.53E-16 |
| SS08584 | 0.000278102 | 0.002614142 | 1.67E-16 |
| SS12328 | 0.000278102 | 0.002614142 | 1.67E-16 |
| SS00262 | 0 | 0.000392005 | 1.73E-16 |
| SS13782 | 0.039456975 | 0.064928424 | 1.74E-16 |
| SS14894 | 0.039456975 | 0.064928424 | 1.74E-16 |
| SS07162 | 0 | 0.00087389 | 1.74E-16 |
| SS04063 | 0 | 0.000368254 | 1.96E-16 |
| SS06336 | 0.000469031 | 0.001938515 | 2.01E-16 |
| SS06833 | 0.000469031 | 0.001938515 | 2.01E-16 |
| SS03241 | 0.000774294 | 0.004807799 | 2.04E-16 |
| SS03334 | 0.000774294 | 0.004807799 | 2.04E-16 |
| SS06682 | 0.080263303 | 0.112559773 | 2.12E-16 |
| SS12331 | 0.080263303 | 0.112559773 | 2.12E-16 |
| SS15382 | 0.080263303 | 0.112559773 | 2.12E-16 |
| SS05905 | 2.76E-06 | 0.002291453 | 2.12E-16 |
| SS02889 | 0.024826273 | 0.001176037 | 2.22E-16 |
| SS12048 | 0.024826273 | 0.001176037 | 2.22E-16 |
| SS02256 | 0.078945894 | 0.061449017 | 2.22E-16 |
| SS01815 | 0.01856302 | 0.010204601 | 2.22E-16 |
| SS09043 | 0.01856302 | 0.010204601 | 2.22E-16 |

| | | | |
|---|---|---|---|
| SS13120 | 0.01856302 | 0.010204601 | 2.22E-16 |
| SS04820 | 0.009163558 | 0.002016846 | 2.22E-16 |
| SS00417 | 0.012261285 | 0.00688791 | 2.22E-16 |
| SS00431 | 0.012261285 | 0.00688791 | 2.22E-16 |
| SS01795 | 0.011160383 | 0.005982237 | 2.22E-16 |
| SS03142 | 0.011160383 | 0.005982237 | 2.22E-16 |
| SS06138 | 0.011160383 | 0.005982237 | 2.22E-16 |
| SS06169 | 0.011160383 | 0.005982237 | 2.22E-16 |
| SS10554 | 0.011160383 | 0.005982237 | 2.22E-16 |
| SS07620 | 0.01070156 | 0.005740485 | 2.22E-16 |
| SS01195 | 0.016661592 | 0.011812684 | 2.22E-16 |
| SS02638 | 0.016661592 | 0.011812684 | 2.22E-16 |
| SS10023 | 0.016661592 | 0.011812684 | 2.22E-16 |
| SS03041 | 0.019411548 | 0.014574082 | 2.22E-16 |
| SS09468 | 0.019411548 | 0.014574082 | 2.22E-16 |
| SS01586 | 0.008139241 | 0.003391287 | 2.22E-16 |
| SS03888 | 0.008139241 | 0.003391287 | 2.22E-16 |
| SS09139 | 0.008139241 | 0.003391287 | 2.22E-16 |
| SS06671 | 0.007733547 | 0.00336803 | 2.22E-16 |
| SS13639 | 0.006308755 | 0.002385308 | 2.22E-16 |
| SS08683 | 0.004780801 | 0.001201123 | 2.22E-16 |
| SS08942 | 0.004780801 | 0.001201123 | 2.22E-16 |
| SS11728 | 0.004780801 | 0.001201123 | 2.22E-16 |
| SS03461 | 0.006638689 | 0.003157217 | 2.22E-16 |
| SS07302 | 0.006638689 | 0.003157217 | 2.22E-16 |
| SS14939 | 0.000879853 | 9.67E-05 | 2.22E-16 |
| SS04928 | 0.000969387 | 0.000347284 | 2.22E-16 |
| SS06991 | 0.000716405 | 0.000102131 | 2.22E-16 |
| SS12070 | 7.90E-05 | 1.71E-05 | 2.22E-16 |
| SS00378 | 5.45E-06 | 1.01E-06 | 2.22E-16 |
| SS06597 | 0 | 0.000950332 | 2.30E-16 |
| SS14988 | 0 | 0.000476643 | 2.35E-16 |
| SS01701 | 0.013731316 | 0.019737391 | 2.38E-16 |
| SS07168 | 0.013731316 | 0.019737391 | 2.38E-16 |
| SS09279 | 0.031178132 | 0.07254968 | 2.40E-16 |
| SS06998 | 0.018276075 | 0.024630212 | 2.41E-16 |
| SS07511 | 0.018276075 | 0.024630212 | 2.41E-16 |
| SS11114 | 0.018276075 | 0.024630212 | 2.41E-16 |
| SS02462 | 0.02310786 | 0.036302266 | 2.41E-16 |
| SS03091 | 0.02310786 | 0.036302266 | 2.41E-16 |
| SS05405 | 0.02310786 | 0.036302266 | 2.41E-16 |
| SS05478 | 0.02310786 | 0.036302266 | 2.41E-16 |
| SS07294 | 0.02310786 | 0.036302266 | 2.41E-16 |
| SS13729 | 0.02310786 | 0.036302266 | 2.41E-16 |
| SS03032 | 0.022282697 | 0.02984724 | 2.42E-16 |
| SS03751 | 0.022282697 | 0.02984724 | 2.42E-16 |
| SS03329 | 0.000533367 | 0.005028794 | 2.44E-16 |

| | | | |
|---|---|---|---|
| SS03802 | 5.45E-06 | 0.005433673 | 2.45E-16 |
| SS07163 | 0 | 0.000510775 | 2.46E-16 |
| SS12748 | 0.014422395 | 0.02598813 | 2.47E-16 |
| SS14503 | 0.014422395 | 0.02598813 | 2.47E-16 |
| SS10968 | 0 | 0.000383716 | 2.48E-16 |
| SS14093 | 0.044800288 | 0.062685926 | 2.49E-16 |
| SS01075 | 2.68E-06 | 0.001378866 | 2.49E-16 |
| SS15015 | 9.83E-05 | 0.001382552 | 2.52E-16 |
| SS08871 | 0.01943583 | 0.025967289 | 2.55E-16 |
| SS11942 | 0.01943583 | 0.025967289 | 2.55E-16 |
| SS11813 | 0 | 0.00060913 | 2.62E-16 |
| SS06629 | 0 | 0.000853279 | 2.66E-16 |
| SS00816 | 0.000394235 | 0.002125094 | 2.71E-16 |
| SS06350 | 2.45E-05 | 0.000476717 | 2.73E-16 |
| SS06856 | 2.45E-05 | 0.000476717 | 2.73E-16 |
| SS02102 | 0.017628662 | 0.029325368 | 2.75E-16 |
| SS02582 | 0.017628662 | 0.029325368 | 2.75E-16 |
| SS07465 | 0.017628662 | 0.029325368 | 2.75E-16 |
| SS14358 | 0.017628662 | 0.029325368 | 2.75E-16 |
| SS11097 | 0 | 0.000437674 | 2.80E-16 |
| SS13759 | 0 | 0.002087844 | 2.87E-16 |
| SS03443 | 0 | 0.000733674 | 2.88E-16 |
| SS08708 | 0.001399478 | 0.00512621 | 3.10E-16 |
| SS10705 | 0.001399478 | 0.00512621 | 3.10E-16 |
| SS13003 | 0.000100672 | 0.000720578 | 3.14E-16 |
| SS07774 | 0.006354382 | 0.011861062 | 3.15E-16 |
| SS00785 | 0 | 0.00081101 | 3.31E-16 |
| SS14454 | 0.004499293 | 0.009725761 | 3.39E-16 |
| SS13361 | 0 | 0.000994792 | 3.44E-16 |
| SS11131 | 0.002734059 | 0.006618579 | 3.52E-16 |
| SS14743 | 0.002734059 | 0.006618579 | 3.52E-16 |
| SS00367 | 0.000329419 | 0.002408621 | 3.60E-16 |
| SS01017 | 0.000329419 | 0.002408621 | 3.60E-16 |
| SS03581 | 0.000329419 | 0.002408621 | 3.60E-16 |
| SS06872 | 0.000329419 | 0.002408621 | 3.60E-16 |
| SS08222 | 0.000329419 | 0.002408621 | 3.60E-16 |
| SS08632 | 0.000329419 | 0.002408621 | 3.60E-16 |
| SS14153 | 0.000329419 | 0.002408621 | 3.60E-16 |
| SS15373 | 0.000329419 | 0.002408621 | 3.60E-16 |
| SS10816 | 0 | 0.000716059 | 3.72E-16 |
| SS02220 | 0.005104645 | 0.011128017 | 3.79E-16 |
| SS14489 | 0.005104645 | 0.011128017 | 3.79E-16 |
| SS04243 | 0.003556665 | 0.0085905 | 3.87E-16 |
| SS03723 | 0.006760673 | 0.011795293 | 4.04E-16 |
| SS00081 | 0.019425014 | 0.025867903 | 4.06E-16 |
| SS02336 | 0.001603231 | 0.006021206 | 4.11E-16 |
| SS04089 | 3.50E-05 | 0.001874942 | 4.30E-16 |

| | | | |
|---|---|---|---|
| SS07212 | 0.001099583 | 0.003140269 | 4.32E-16 |
| SS01564 | 0.008924397 | 0.003609255 | 4.44E-16 |
| SS02183 | 0.008924397 | 0.003609255 | 4.44E-16 |
| SS07207 | 0.00320894 | 0.000753089 | 4.44E-16 |
| SS14745 | 8.99E-05 | 1.18E-05 | 4.44E-16 |
| SS10354 | 0.002837622 | 0.009402555 | 4.53E-16 |
| SS08497 | 0 | 0.00043059 | 4.74E-16 |
| SS01895 | 0 | 0.000336394 | 4.92E-16 |
| SS07496 | 0 | 0.000336394 | 4.92E-16 |
| SS02401 | 0 | 0.001772679 | 4.93E-16 |
| SS03628 | 0 | 0.001772679 | 4.93E-16 |
| SS00180 | 0 | 0.000562675 | 5.22E-16 |
| SS05682 | 9.81E-05 | 0.0007747 | 5.36E-16 |
| SS08582 | 9.81E-05 | 0.0007747 | 5.36E-16 |
| SS09602 | 0 | 0.000757541 | 5.63E-16 |
| SS13772 | 0 | 0.000848025 | 5.67E-16 |
| SS13776 | 0 | 0.000848025 | 5.67E-16 |
| SS13549 | 0.022398933 | 0.058877294 | 5.71E-16 |
| SS06628 | 0 | 0.002112826 | 5.80E-16 |
| SS00523 | 0 | 0.000471045 | 5.83E-16 |
| SS04371 | 0.032351743 | 0.042302892 | 5.85E-16 |
| SS09033 | 0.032351743 | 0.042302892 | 5.85E-16 |
| SS12925 | 0.032351743 | 0.042302892 | 5.85E-16 |
| SS12882 | 0 | 0.001520595 | 6.01E-16 |
| SS05787 | 0.000245493 | 0.001924387 | 6.59E-16 |
| SS00416 | 0.013961817 | 0.003748648 | 6.66E-16 |
| SS00430 | 0.013961817 | 0.003748648 | 6.66E-16 |
| SS10563 | 0.013961817 | 0.003748648 | 6.66E-16 |
| SS02292 | 0.00241041 | 2.36E-05 | 6.66E-16 |
| SS04211 | 0.001587563 | 0.000147064 | 6.66E-16 |
| SS04421 | 0.001587563 | 0.000147064 | 6.66E-16 |
| SS06292 | 0.001587563 | 0.000147064 | 6.66E-16 |
| SS08963 | 0.002469056 | 0.007115506 | 6.83E-16 |
| SS11010 | 0.002469056 | 0.007115506 | 6.83E-16 |
| SS02053 | 3.28E-05 | 0.001410071 | 7.00E-16 |
| SS07260 | 0 | 0.000541696 | 7.64E-16 |
| SS08051 | 0 | 0.000550094 | 8.02E-16 |
| SS05760 | 0 | 0.001383095 | 8.07E-16 |
| SS09571 | 0 | 0.00155521 | 8.22E-16 |
| SS04729 | 0 | 0.000612506 | 8.33E-16 |
| SS06143 | 0 | 0.001536484 | 8.45E-16 |
| SS06173 | 0 | 0.001536484 | 8.45E-16 |
| SS06287 | 0.019605403 | 0.025554046 | 8.53E-16 |
| SS06620 | 0 | 0.000974313 | 8.58E-16 |
| SS00232 | 0 | 0.000501468 | 8.64E-16 |
| SS09076 | 0.021491863 | 0.032265516 | 8.86E-16 |
| SS03015 | 0.021025389 | 0.012771719 | 8.88E-16 |

| | | | |
|---|---|---|---|
| SS05233 | 0.021025389 | 0.012771719 | 8.88E-16 |
| SS05517 | 0.021025389 | 0.012771719 | 8.88E-16 |
| SS09970 | 0.021025389 | 0.012771719 | 8.88E-16 |
| SS04403 | 0.014949751 | 0.008378418 | 8.88E-16 |
| SS00323 | 0.012467883 | 0.00664134 | 8.88E-16 |
| SS06026 | 0.012467883 | 0.00664134 | 8.88E-16 |
| SS02539 | 0.019005453 | 0.013949915 | 8.88E-16 |
| SS08214 | 0.019005453 | 0.013949915 | 8.88E-16 |
| SS10485 | 0.019005453 | 0.013949915 | 8.88E-16 |
| SS09584 | 0.006208404 | 0.002038643 | 8.88E-16 |
| SS13601 | 0.009539247 | 0.006090596 | 8.88E-16 |
| SS05414 | 0.003106983 | 0.001736652 | 8.88E-16 |
| SS05446 | 0.003106983 | 0.001736652 | 8.88E-16 |
| SS01610 | 5.45E-06 | 8.01E-07 | 8.88E-16 |
| SS00997 | 0.00043594 | 0.001421016 | 8.93E-16 |
| SS03806 | 0 | 0.001785234 | 8.98E-16 |
| SS09484 | 0 | 0.000263846 | 9.28E-16 |
| SS05340 | 5.45E-06 | 0.000785276 | 9.29E-16 |
| SS02488 | 0 | 0.001365883 | 9.47E-16 |
| SS11440 | 0.016967818 | 0.024592954 | 9.65E-16 |
| SS05139 | 3.23E-05 | 0.000983787 | 9.70E-16 |
| SS15141 | 3.23E-05 | 0.000983787 | 9.70E-16 |
| SS00781 | 0.028311182 | 0.039758382 | 9.71E-16 |
| SS13930 | 0.000287954 | 0.00791601 | 1.03E-15 |
| SS00887 | 0 | 0.001043409 | 1.05E-15 |
| SS13216 | 0 | 0.001043409 | 1.05E-15 |
| SS02388 | 0 | 0.000512458 | 1.06E-15 |
| SS03267 | 0 | 0.000299078 | 1.09E-15 |
| SS11087 | 0.007364156 | 0.013370787 | 1.11E-15 |
| SS13963 | 0.014292485 | 0.005991906 | 1.11E-15 |
| SS05412 | 0.005508183 | 0.002009839 | 1.11E-15 |
| SS05421 | 0.005508183 | 0.002009839 | 1.11E-15 |
| SS09922 | 0.001206346 | 5.02E-05 | 1.11E-15 |
| SS11949 | 0.001206346 | 5.02E-05 | 1.11E-15 |
| SS08685 | 1.36E-05 | 0.000255965 | 1.13E-15 |
| SS11222 | 0.010799399 | 0.018989707 | 1.15E-15 |
| SS11442 | 0.010799399 | 0.018989707 | 1.15E-15 |
| SS01008 | 0.001737912 | 0.003916139 | 1.21E-15 |
| SS02516 | 0.001737912 | 0.003916139 | 1.21E-15 |
| SS13889 | 0 | 0.000958398 | 1.29E-15 |
| SS15004 | 0 | 0.000958398 | 1.29E-15 |
| SS03005 | 0.012538297 | 0.021821393 | 1.29E-15 |
| SS14469 | 0.012538297 | 0.021821393 | 1.29E-15 |
| SS05987 | 0.007330503 | 0.01375317 | 1.29E-15 |
| SS06030 | 0.007330503 | 0.01375317 | 1.29E-15 |
| SS11112 | 0.007330503 | 0.01375317 | 1.29E-15 |
| SS12994 | 0.007330503 | 0.01375317 | 1.29E-15 |

| | | | |
|---|---|---|---|
| SS13469 | 0.029472898 | 0.007705512 | 1.33E-15 |
| SS02579 | 0.04025976 | 0.029805349 | 1.33E-15 |
| SS04713 | 0.04025976 | 0.029805349 | 1.33E-15 |
| SS11049 | 0.04025976 | 0.029805349 | 1.33E-15 |
| SS11885 | 0.022106839 | 0.011836338 | 1.33E-15 |
| SS08862 | 0 | 0.000771888 | 1.34E-15 |
| SS08135 | 0.020198437 | 0.03487012 | 1.35E-15 |
| SS13769 | 0.097486616 | 0.13139439 | 1.38E-15 |
| SS13775 | 0.097486616 | 0.13139439 | 1.38E-15 |
| SS14892 | 0.097486616 | 0.13139439 | 1.38E-15 |
| SS14128 | 8.29E-06 | 0.000558672 | 1.39E-15 |
| SS14357 | 8.29E-06 | 0.000558672 | 1.39E-15 |
| SS02338 | 0.032047972 | 0.042645998 | 1.40E-15 |
| SS10473 | 0.032047972 | 0.042645998 | 1.40E-15 |
| SS05009 | 0 | 0.000296907 | 1.42E-15 |
| SS10044 | 0 | 0.000390621 | 1.45E-15 |
| SS11435 | 0.005754237 | 0.010830345 | 1.49E-15 |
| SS15281 | 0 | 0.000745957 | 1.55E-15 |
| SS14788 | 0.03183916 | 0.005683837 | 1.55E-15 |
| SS15052 | 0.03183916 | 0.005683837 | 1.55E-15 |
| SS10565 | 0.006561657 | 0.000460946 | 1.55E-15 |
| SS09986 | 0.033448918 | 0.04840577 | 1.64E-15 |
| SS09019 | 0 | 0.000292973 | 1.68E-15 |
| SS07479 | 0.012299779 | 0.006346621 | 1.78E-15 |
| SS15095 | 0.012299779 | 0.006346621 | 1.78E-15 |
| SS06886 | 0 | 0.000331008 | 1.79E-15 |
| SS14225 | 0 | 0.000331008 | 1.79E-15 |
| SS01834 | 0 | 0.000732078 | 1.83E-15 |
| SS02263 | 0.000808544 | 0.002917615 | 1.89E-15 |
| SS10026 | 0.000808544 | 0.002917615 | 1.89E-15 |
| SS15232 | 0 | 0.00030834 | 1.97E-15 |
| SS09815 | 0 | 0.001754166 | 1.99E-15 |
| SS14351 | 0 | 0.001754166 | 1.99E-15 |
| SS10080 | 0.002372593 | 0.001334253 | 2.00E-15 |
| SS09479 | 2.68E-06 | 0.000404108 | 2.00E-15 |
| SS15207 | 0 | 0.001439051 | 2.01E-15 |
| SS06155 | 4.35E-05 | 0.000711604 | 2.01E-15 |
| SS02337 | 8.75E-05 | 0.001191782 | 2.11E-15 |
| SS05530 | 2.76E-06 | 0.000403998 | 2.17E-15 |
| SS07790 | 0.017149606 | 0.000298097 | 2.22E-15 |
| SS12044 | 0.017149606 | 0.000298097 | 2.22E-15 |
| SS09599 | 0.0133253 | 0.001633372 | 2.22E-15 |
| SS15532 | 0.010911277 | 0.003070699 | 2.22E-15 |
| SS03128 | 0.010547013 | 0.00655431 | 2.22E-15 |
| SS12369 | 0.00013889 | 4.19E-05 | 2.22E-15 |
| SS03118 | 0.002457838 | 0.007776267 | 2.23E-15 |
| SS10209 | 0.002457838 | 0.007776267 | 2.23E-15 |

| | | | |
|---|---|---|---|
| SS10226 | 0.002457838 | 0.007776267 | 2.23E-15 |
| SS07789 | 0 | 0.00069256 | 2.26E-15 |
| SS03453 | 0 | 0.000831788 | 2.26E-15 |
| SS05786 | 0.000770807 | 0.003142759 | 2.36E-15 |
| SS01408 | 0 | 0.001006897 | 2.48E-15 |
| SS01003 | 0.014017951 | 0.006745922 | 2.66E-15 |
| SS02471 | 0.014017951 | 0.006745922 | 2.66E-15 |
| SS03624 | 0.005442402 | 0.000937065 | 2.66E-15 |
| SS10946 | 0.006172468 | 0.003634955 | 2.66E-15 |
| SS07557 | 0.017353279 | 0.024411515 | 2.69E-15 |
| SS11264 | 0.017353279 | 0.024411515 | 2.69E-15 |
| SS07696 | 4.37E-05 | 0.000737305 | 2.81E-15 |
| SS01811 | 0 | 0.003410695 | 2.89E-15 |
| SS02003 | 0 | 0.003410695 | 2.89E-15 |
| SS03188 | 0 | 0.003410695 | 2.89E-15 |
| SS09664 | 0 | 0.003410695 | 2.89E-15 |
| SS09292 | 0.018387403 | 0.030029864 | 2.93E-15 |
| SS03503 | 0 | 0.000252863 | 3.04E-15 |
| SS10717 | 0 | 0.000892001 | 3.07E-15 |
| SS00071 | 0.013217574 | 0.020406807 | 3.13E-15 |
| SS03701 | 0.013217574 | 0.020406807 | 3.13E-15 |
| SS04821 | 0.013217574 | 0.020406807 | 3.13E-15 |
| SS05504 | 0.013217574 | 0.020406807 | 3.13E-15 |
| SS06267 | 0 | 0.000384315 | 3.33E-15 |
| SS04073 | 0 | 0.00081549 | 3.49E-15 |
| SS01524 | 9.81E-05 | 0.000601758 | 3.49E-15 |
| SS07317 | 9.81E-05 | 0.000601758 | 3.49E-15 |
| SS08134 | 0.023575951 | 0.038553164 | 3.54E-15 |
| SS09407 | 8.13E-06 | 0.000529518 | 3.56E-15 |
| SS14215 | 0.001366869 | 0.003457539 | 3.65E-15 |
| SS11122 | 0 | 0.000985297 | 3.70E-15 |
| SS13137 | 6.26E-05 | 0.000578931 | 3.91E-15 |
| SS15308 | 6.26E-05 | 0.000578931 | 3.91E-15 |
| SS00904 | 5.45E-06 | 0.003573425 | 3.99E-15 |
| SS09961 | 5.45E-06 | 0.003573425 | 3.99E-15 |
| SS10525 | 0.000405935 | 0.000109418 | 4.00E-15 |
| SS03328 | 0.000441469 | 0.004560022 | 4.17E-15 |
| SS04062 | 0 | 0.00045453 | 4.23E-15 |
| SS07651 | 0.002470341 | 0.004688789 | 4.38E-15 |
| SS12219 | 0.000689244 | 0.000252463 | 4.44E-15 |
| SS02606 | 0.028362384 | 0.036707922 | 4.64E-15 |
| SS08819 | 0.028362384 | 0.036707922 | 4.64E-15 |
| SS13706 | 0.028362384 | 0.036707922 | 4.64E-15 |
| SS04584 | 0.024644669 | 0.00505959 | 4.66E-15 |
| SS06528 | 0.008607309 | 0.004511498 | 4.66E-15 |
| SS06531 | 0.008607309 | 0.004511498 | 4.66E-15 |
| SS07741 | 0.008607309 | 0.004511498 | 4.66E-15 |

| | | | |
|---|---|---|---|
| SS13449 | 0.008607309 | 0.004511498 | 4.66E-15 |
| SS00032 | 0 | 0.000545888 | 4.70E-15 |
| SS04220 | 0 | 0.001204015 | 5.20E-15 |
| SS11616 | 0.106844683 | 0.141948836 | 5.48E-15 |
| SS13932 | 0.004141635 | 0.025635941 | 5.51E-15 |
| SS13958 | 0.004141635 | 0.025635941 | 5.51E-15 |
| SS08370 | 5.45E-06 | 0.000476329 | 5.54E-15 |
| SS09862 | 0.00428805 | 0.008161624 | 5.55E-15 |
| SS06996 | 0.000811468 | 0.000389637 | 5.55E-15 |
| SS12302 | 0 | 0.000853312 | 5.71E-15 |
| SS09273 | 4.38E-05 | 0.000779016 | 5.82E-15 |
| SS08058 | 0 | 0.000567009 | 5.97E-15 |
| SS07211 | 0.002850996 | 0.000540655 | 6.00E-15 |
| SS07228 | 0.002850996 | 0.000540655 | 6.00E-15 |
| SS02231 | 0.00819237 | 0.014423866 | 6.08E-15 |
| SS06929 | 0.00819237 | 0.014423866 | 6.08E-15 |
| SS09947 | 0.00819237 | 0.014423866 | 6.08E-15 |
| SS08446 | 0.014598104 | 0.019716895 | 6.31E-15 |
| SS08477 | 0.014598104 | 0.019716895 | 6.31E-15 |
| SS10858 | 0.014598104 | 0.019716895 | 6.31E-15 |
| SS15413 | 0.014598104 | 0.019716895 | 6.31E-15 |
| SS04206 | 0.017544656 | 0.030291873 | 6.39E-15 |
| SS10809 | 0.017544656 | 0.030291873 | 6.39E-15 |
| SS09542 | 0.015303463 | 0.030193309 | 6.43E-15 |
| SS00362 | 0.025720957 | 0.038394681 | 6.79E-15 |
| SS01709 | 0.025720957 | 0.038394681 | 6.79E-15 |
| SS01727 | 0.025720957 | 0.038394681 | 6.79E-15 |
| SS06914 | 0.025720957 | 0.038394681 | 6.79E-15 |
| SS07170 | 0.025720957 | 0.038394681 | 6.79E-15 |
| SS07181 | 0.025720957 | 0.038394681 | 6.79E-15 |
| SS10740 | 0.025720957 | 0.038394681 | 6.79E-15 |
| SS15077 | 0.025720957 | 0.038394681 | 6.79E-15 |
| SS04822 | 0.018574766 | 0.014102417 | 6.88E-15 |
| SS05015 | 0.018574766 | 0.014102417 | 6.88E-15 |
| SS11231 | 0.018574766 | 0.014102417 | 6.88E-15 |
| SS11552 | 0.018574766 | 0.014102417 | 6.88E-15 |
| SS11738 | 0.018574766 | 0.014102417 | 6.88E-15 |
| SS11756 | 0.018574766 | 0.014102417 | 6.88E-15 |
| SS00100 | 0.002176296 | 0.001330121 | 6.88E-15 |
| SS02405 | 0.000138248 | 0.001635665 | 6.92E-15 |
| SS03635 | 0.000138248 | 0.001635665 | 6.92E-15 |
| SS01851 | 0.015401807 | 0.006746081 | 7.11E-15 |
| SS06770 | 0.015401807 | 0.006746081 | 7.11E-15 |
| SS06806 | 0.015401807 | 0.006746081 | 7.11E-15 |
| SS13841 | 0.015401807 | 0.006746081 | 7.11E-15 |
| SS08146 | 0 | 0.000345782 | 7.19E-15 |
| SS04230 | 0 | 0.001215574 | 7.22E-15 |

| | | | |
|---|---|---|---|
| SS05094 | 2.76E-06 | 0.002951743 | 7.23E-15 |
| SS02489 | 0.000386424 | 0.004042994 | 7.42E-15 |
| SS14065 | 0.00091799 | 0.005217398 | 7.76E-15 |
| SS03671 | 0.015297889 | 0.009501122 | 7.77E-15 |
| SS07079 | 0.015297889 | 0.009501122 | 7.77E-15 |
| SS13005 | 0.015297889 | 0.009501122 | 7.77E-15 |
| SS01427 | 0.004722751 | 4.65E-05 | 8.22E-15 |
| SS00186 | 0 | 0.000989039 | 8.41E-15 |
| SS00604 | 0 | 0.000989039 | 8.41E-15 |
| SS10657 | 0 | 0.000829797 | 8.50E-15 |
| SS10297 | 0.00868675 | 0.013702604 | 8.51E-15 |
| SS02291 | 0.055262125 | 0.070508489 | 8.61E-15 |
| SS02384 | 0.055262125 | 0.070508489 | 8.61E-15 |
| SS13999 | 0.055262125 | 0.070508489 | 8.61E-15 |
| SS11091 | 0 | 0.000575062 | 8.71E-15 |
| SS03614 | 0 | 0.000329533 | 9.03E-15 |
| SS03148 | 0.002200853 | 0.000117638 | 9.10E-15 |
| SS04359 | 0 | 0.000220306 | 9.51E-15 |
| SS13132 | 0 | 0.000220306 | 9.51E-15 |
| SS00894 | 0.003323594 | 0.001166103 | 9.77E-15 |
| SS04513 | 0.003323594 | 0.001166103 | 9.77E-15 |
| SS06692 | 0.007225655 | 0.014455921 | 9.87E-15 |
| SS09541 | 0.006600792 | 0.001153804 | 9.99E-15 |
| SS11335 | 7.35E-05 | 1.47E-05 | 9.99E-15 |
| SS02086 | 0 | 0.001548038 | 1.01E-14 |
| SS08123 | 0.012074645 | 0.001714606 | 1.04E-14 |
| SS09130 | 0.001797281 | 0.004225119 | 1.05E-14 |
| SS02133 | 0.003236697 | 0.006876912 | 1.05E-14 |
| SS03538 | 0.003236697 | 0.006876912 | 1.05E-14 |
| SS05204 | 0.003236697 | 0.006876912 | 1.05E-14 |
| SS10429 | 0.003236697 | 0.006876912 | 1.05E-14 |
| SS06122 | 0.011482828 | 0.016836511 | 1.06E-14 |
| SS05774 | 0 | 0.000500586 | 1.13E-14 |
| SS05745 | 0.003076221 | 0.006642012 | 1.17E-14 |
| SS10039 | 0 | 0.001002666 | 1.17E-14 |
| SS02385 | 0.016099493 | 0.011498629 | 1.18E-14 |
| SS02538 | 0.016099493 | 0.011498629 | 1.18E-14 |
| SS03521 | 0.016099493 | 0.011498629 | 1.18E-14 |
| SS14214 | 0.016099493 | 0.011498629 | 1.18E-14 |
| SS08712 | 0.001228541 | 0.00284571 | 1.21E-14 |
| SS10707 | 0.001228541 | 0.00284571 | 1.21E-14 |
| SS07517 | 0.000550033 | 3.01E-05 | 1.33E-14 |
| SS06471 | 0 | 0.000818496 | 1.39E-14 |
| SS09536 | 0.030426079 | 0.020548377 | 1.40E-14 |
| SS02101 | 0.012255998 | 0.008193829 | 1.40E-14 |
| SS09066 | 0.012255998 | 0.008193829 | 1.40E-14 |
| SS02993 | 0 | 0.000242042 | 1.41E-14 |

| | | | |
|---|---|---|---|
| SS07161 | 0 | 0.000329248 | 1.42E-14 |
| SS00078 | 0 | 0.000759727 | 1.46E-14 |
| SS12278 | 0.002950269 | 0.001255178 | 1.49E-14 |
| SS06134 | 2.76E-06 | 0.00049113 | 1.49E-14 |
| SS00998 | 2.68E-06 | 0.001207772 | 1.50E-14 |
| SS03680 | 5.37E-06 | 0.000442082 | 1.56E-14 |
| SS07091 | 5.37E-06 | 0.000442082 | 1.56E-14 |
| SS08305 | 0.014013386 | 0.006047681 | 1.58E-14 |
| SS04219 | 0 | 0.000726199 | 1.60E-14 |
| SS04372 | 0.000114413 | 0.000694793 | 1.66E-14 |
| SS07650 | 0.002742994 | 0.005912688 | 1.75E-14 |
| SS11438 | 0.002742994 | 0.005912688 | 1.75E-14 |
| SS08339 | 0.022808849 | 0.010415348 | 1.80E-14 |
| SS10709 | 0.000274936 | 0.000847222 | 1.80E-14 |
| SS13317 | 0.00624614 | 0.002131371 | 1.82E-14 |
| SS15153 | 0.00624614 | 0.002131371 | 1.82E-14 |
| SS03616 | 0.003704731 | 0.007154251 | 1.85E-14 |
| SS05780 | 0.003704731 | 0.007154251 | 1.85E-14 |
| SS06514 | 0.003704731 | 0.007154251 | 1.85E-14 |
| SS10419 | 0.003704731 | 0.007154251 | 1.85E-14 |
| SS10440 | 0.003704731 | 0.007154251 | 1.85E-14 |
| SS11817 | 0.003704731 | 0.007154251 | 1.85E-14 |
| SS03991 | 0.007861701 | 0.001529332 | 1.91E-14 |
| SS01024 | 2.76E-06 | 0.000548047 | 1.92E-14 |
| SS06447 | 0 | 0.000625072 | 1.95E-14 |
| SS07721 | 0 | 0.000625072 | 1.95E-14 |
| SS12336 | 2.68E-06 | 0.000457723 | 1.97E-14 |
| SS06453 | 0.011801464 | 0.006167323 | 2.00E-14 |
| SS11862 | 0.011801464 | 0.006167323 | 2.00E-14 |
| SS10327 | 0 | 0.000501571 | 2.05E-14 |
| SS00425 | 9.25E-05 | 0.000520299 | 2.07E-14 |
| SS00404 | 0 | 0.000311534 | 2.15E-14 |
| SS02654 | 0.001206427 | 2.41E-05 | 2.20E-14 |
| SS01122 | 0.013680435 | 0.018567676 | 2.22E-14 |
| SS03019 | 0.013680435 | 0.018567676 | 2.22E-14 |
| SS14474 | 0.013680435 | 0.018567676 | 2.22E-14 |
| SS03231 | 0.000421637 | 0.0015636 | 2.29E-14 |
| SS03323 | 0.000421637 | 0.0015636 | 2.29E-14 |
| SS06242 | 0.003428189 | 0.001811314 | 2.40E-14 |
| SS06877 | 0.003428189 | 0.001811314 | 2.40E-14 |
| SS15384 | 0.003428189 | 0.001811314 | 2.40E-14 |
| SS15445 | 0.003428189 | 0.001811314 | 2.40E-14 |
| SS04059 | 0 | 0.000588055 | 2.44E-14 |
| SS05136 | 0.000180033 | 0.001131413 | 2.44E-14 |
| SS15138 | 0.000180033 | 0.001131413 | 2.44E-14 |
| SS15211 | 0 | 0.000781396 | 2.49E-14 |
| SS07580 | 0.00011189 | 0.00078225 | 2.50E-14 |

| | | | |
|---|---|---|---|
| SS00957 | 0.017749201 | 0.00680366 | 2.58E-14 |
| SS04655 | 0.017749201 | 0.00680366 | 2.58E-14 |
| SS02502 | 0 | 0.00066454 | 2.63E-14 |
| SS09852 | 0.0022464 | 0.000861054 | 2.71E-14 |
| SS00744 | 0.001225776 | 9.64E-05 | 2.78E-14 |
| SS01617 | 0.005401821 | 0.009158693 | 2.80E-14 |
| SS00227 | 0.000577515 | 0.000263162 | 2.84E-14 |
| SS05384 | 0 | 0.000270882 | 2.87E-14 |
| SS06927 | 0.032296779 | 0.047097486 | 2.88E-14 |
| SS10640 | 0.032296779 | 0.047097486 | 2.88E-14 |
| SS05467 | 0.003131541 | 0.000355735 | 2.93E-14 |
| SS07603 | 0.001176501 | 0.00067916 | 2.93E-14 |
| SS11928 | 0 | 0.001924714 | 2.95E-14 |
| SS05222 | 0 | 0.000540279 | 2.98E-14 |
| SS10412 | 0 | 0.000540279 | 2.98E-14 |
| SS10447 | 0 | 0.000540279 | 2.98E-14 |
| SS02467 | 0 | 0.001139893 | 3.00E-14 |
| SS06627 | 3.28E-05 | 0.000589601 | 3.01E-14 |
| SS01644 | 0.006246381 | 0.001872643 | 3.09E-14 |
| SS04866 | 0.006246381 | 0.001872643 | 3.09E-14 |
| SS00066 | 0 | 0.000989597 | 3.09E-14 |
| SS05184 | 0 | 0.000989597 | 3.09E-14 |
| SS08864 | 0 | 0.000989597 | 3.09E-14 |
| SS11903 | 0 | 0.000989597 | 3.09E-14 |
| SS13485 | 0 | 0.000989597 | 3.09E-14 |
| SS09465 | 0.004136542 | 0.000313093 | 3.22E-14 |
| SS05087 | 2.68E-06 | 0.000339374 | 3.27E-14 |
| SS11284 | 0.013398055 | 0.021507217 | 3.27E-14 |
| SS11613 | 0 | 0.001049746 | 3.29E-14 |
| SS09421 | 0.017148769 | 0.012664687 | 3.33E-14 |
| SS10041 | 0 | 0.000341408 | 3.35E-14 |
| SS09778 | 5.53E-06 | 0.003716365 | 3.38E-14 |
| SS10591 | 5.53E-06 | 0.003716365 | 3.38E-14 |
| SS15170 | 5.53E-06 | 0.003716365 | 3.38E-14 |
| SS04353 | 0 | 0.000511383 | 3.38E-14 |
| SS07160 | 0 | 0.00078549 | 3.39E-14 |
| SS08309 | 0.001546501 | 0.000512798 | 3.42E-14 |
| SS12042 | 0.034002885 | 0.04243868 | 3.46E-14 |
| SS14860 | 0.034002885 | 0.04243868 | 3.46E-14 |
| SS07237 | 0.01383271 | 0.005061076 | 3.46E-14 |
| SS00926 | 0.026546831 | 0.034903423 | 3.47E-14 |
| SS11614 | 0.026546831 | 0.034903423 | 3.47E-14 |
| SS03089 | 0.003169839 | 0.001848028 | 3.69E-14 |
| SS05403 | 0.003169839 | 0.001848028 | 3.69E-14 |
| SS05568 | 7.63E-05 | 2.47E-06 | 3.69E-14 |
| SS10358 | 0.000144258 | 0.000698141 | 3.70E-14 |
| SS01391 | 0.001886139 | 0.005637746 | 3.71E-14 |

| | | | |
|---|---|---|---|
| SS07690 | 0.001795962 | 0.004313504 | 3.93E-14 |
| SS13934 | 0.001795962 | 0.004313504 | 3.93E-14 |
| SS02701 | 0.001045904 | 0.000574018 | 3.97E-14 |
| SS06837 | 0.001045904 | 0.000574018 | 3.97E-14 |
| SS14729 | 0.007147304 | 0.000755984 | 4.02E-14 |
| SS01894 | 0.004740335 | 0.007905726 | 4.12E-14 |
| SS07495 | 0.004740335 | 0.007905726 | 4.12E-14 |
| SS03312 | 0.019675106 | 0.025898769 | 4.13E-14 |
| SS07569 | 0 | 0.000538601 | 4.18E-14 |
| SS01052 | 0.052051269 | 0.069624537 | 4.29E-14 |
| SS01313 | 0.052051269 | 0.069624537 | 4.29E-14 |
| SS02601 | 0.052051269 | 0.069624537 | 4.29E-14 |
| SS09813 | 0.052051269 | 0.069624537 | 4.29E-14 |
| SS07911 | 0 | 0.000375714 | 4.34E-14 |
| SS08894 | 8.17E-05 | 0.000544043 | 4.43E-14 |
| SS09554 | 0.004983786 | 0.007763456 | 4.47E-14 |
| SS04593 | 2.68E-06 | 0.00021285 | 4.53E-14 |
| SS06984 | 0 | 0.000360123 | 4.82E-14 |
| SS10020 | 0 | 0.000293509 | 4.97E-14 |
| SS02399 | 0.00791342 | 0.001546496 | 5.02E-14 |
| SS06038 | 0.028615206 | 0.022219477 | 5.04E-14 |
| SS06063 | 0.028615206 | 0.022219477 | 5.04E-14 |
| SS01841 | 0 | 0.00047309 | 5.09E-14 |
| SS05657 | 0.000519866 | 0.003286195 | 5.09E-14 |
| SS06149 | 0.031146406 | 0.021580922 | 5.15E-14 |
| SS06179 | 0.031146406 | 0.021580922 | 5.15E-14 |
| SS10780 | 0.031146406 | 0.021580922 | 5.15E-14 |
| SS08059 | 0 | 0.000429167 | 5.19E-14 |
| SS09489 | 0 | 0.000686235 | 5.21E-14 |
| SS00899 | 0 | 0.001297461 | 5.23E-14 |
| SS04228 | 0.000678428 | 0.00342631 | 5.34E-14 |
| SS08684 | 0.000678428 | 0.00342631 | 5.34E-14 |
| SS08943 | 0.000678428 | 0.00342631 | 5.34E-14 |
| SS04570 | 0.022231173 | 0.014177019 | 5.55E-14 |
| SS04642 | 0.022231173 | 0.014177019 | 5.55E-14 |
| SS06659 | 0.022231173 | 0.014177019 | 5.55E-14 |
| SS06537 | 0.032889124 | 0.041936031 | 5.86E-14 |
| SS07327 | 0.002175974 | 3.92E-05 | 6.06E-14 |
| SS15303 | 2.76E-06 | 0.000411666 | 6.14E-14 |
| SS01199 | 0.013237715 | 0.022057113 | 6.39E-14 |
| SS02639 | 0.013237715 | 0.022057113 | 6.39E-14 |
| SS10024 | 0.013237715 | 0.022057113 | 6.39E-14 |
| SS00373 | 0.003424267 | 0.001660763 | 6.46E-14 |
| SS06880 | 0.003424267 | 0.001660763 | 6.46E-14 |
| SS14220 | 0.003424267 | 0.001660763 | 6.46E-14 |
| SS15401 | 0.003424267 | 0.001660763 | 6.46E-14 |
| SS09096 | 0.007146983 | 0.000309014 | 6.53E-14 |

| | | | |
|---|---|---|---|
| SS10997 | 0 | 0.000536841 | 6.55E-14 |
| SS13853 | 0.001197973 | 0.004568109 | 6.66E-14 |
| SS11053 | 0 | 0.000478395 | 6.73E-14 |
| SS05253 | 5.45E-06 | 0.000300375 | 6.77E-14 |
| SS12770 | 5.45E-06 | 0.000300375 | 6.77E-14 |
| SS03989 | 0.018095446 | 0.011758391 | 6.88E-14 |
| SS14123 | 0.018095446 | 0.011758391 | 6.88E-14 |
| SS00501 | 0.086595882 | 0.11164748 | 6.92E-14 |
| SS05811 | 0.086595882 | 0.11164748 | 6.92E-14 |
| SS05833 | 0.086595882 | 0.11164748 | 6.92E-14 |
| SS15225 | 2.20E-05 | 0.000891685 | 6.95E-14 |
| SS05837 | 5.53E-06 | 0.000346436 | 7.09E-14 |
| SS04355 | 0 | 0.001806759 | 7.50E-14 |
| SS10383 | 0 | 0.000521946 | 8.17E-14 |
| SS11900 | 0.012176751 | 0.019138746 | 8.22E-14 |
| SS02693 | 0.009085918 | 0.006518059 | 8.24E-14 |
| SS09551 | 0.009085918 | 0.006518059 | 8.24E-14 |
| SS14126 | 0.009085918 | 0.006518059 | 8.24E-14 |
| SS05691 | 0.001736387 | 0.004424954 | 8.24E-14 |
| SS08717 | 0.001736387 | 0.004424954 | 8.24E-14 |
| SS15024 | 0.001736387 | 0.004424954 | 8.24E-14 |
| SS15031 | 0.001736387 | 0.004424954 | 8.24E-14 |
| SS04055 | 0.00103737 | 0.00244187 | 8.32E-14 |
| SS00863 | 2.47E-05 | 0.001078259 | 8.36E-14 |
| SS12041 | 7.92E-05 | 0.001852739 | 8.36E-14 |
| SS14027 | 0.02026861 | 0.030913133 | 8.37E-14 |
| SS03179 | 0.000433738 | 0.002621777 | 8.41E-14 |
| SS08421 | 0.014524512 | 1.48E-05 | 8.42E-14 |
| SS08463 | 0.014524512 | 1.48E-05 | 8.42E-14 |
| SS00181 | 2.76E-06 | 0.000672011 | 8.44E-14 |
| SS14897 | 0.003194717 | 7.86E-07 | 8.57E-14 |
| SS09323 | 0.000550273 | 0.000114982 | 8.86E-14 |
| SS11113 | 0.005731917 | 0.001530512 | 8.93E-14 |
| SS00213 | 0.001865424 | 0.000123673 | 8.97E-14 |
| SS00736 | 0.001865424 | 0.000123673 | 8.97E-14 |
| SS09834 | 0.001865424 | 0.000123673 | 8.97E-14 |
| SS06760 | 0 | 0.000303926 | 9.05E-14 |
| SS08431 | 0 | 0.000303926 | 9.05E-14 |
| SS08468 | 0 | 0.000303926 | 9.05E-14 |
| SS05770 | 0 | 0.002854763 | 9.16E-14 |
| SS01787 | 0.005865278 | 0.002064818 | 9.19E-14 |
| SS03255 | 0.005865278 | 0.002064818 | 9.19E-14 |
| SS02631 | 0.001671329 | 0.003717509 | 9.20E-14 |
| SS04583 | 0.001671329 | 0.003717509 | 9.20E-14 |
| SS13580 | 0.004310405 | 0.001649541 | 9.73E-14 |
| SS06734 | 0 | 0.000157326 | 9.95E-14 |
| SS06502 | 0.002137516 | 0.000921156 | 1.01E-13 |

| | | | |
|---|---|---|---|
| SS07176 | 0.002137516 | 0.000921156 | 1.01E-13 |
| SS09229 | 0.005271063 | 0.002846876 | 1.03E-13 |
| SS10932 | 0.005271063 | 0.002846876 | 1.03E-13 |
| SS08673 | 0.000566538 | 0.001475348 | 1.04E-13 |
| SS04322 | 0.019375452 | 0.005318317 | 1.05E-13 |
| SS12968 | 0.019375452 | 0.005318317 | 1.05E-13 |
| SS06708 | 0.021085573 | 0.00746404 | 1.06E-13 |
| SS13585 | 0 | 0.000420122 | 1.17E-13 |
| SS14949 | 0 | 0.000351219 | 1.17E-13 |
| SS13548 | 0 | 0.001926515 | 1.19E-13 |
| SS06576 | 0 | 0.000571124 | 1.19E-13 |
| SS10463 | 0.055144064 | 0.003947702 | 1.21E-13 |
| SS10663 | 0.055144064 | 0.003947702 | 1.21E-13 |
| SS03656 | 0.000384222 | 0.001399095 | 1.23E-13 |
| SS07068 | 0.000384222 | 0.001399095 | 1.23E-13 |
| SS09103 | 0.005147554 | 1.76E-05 | 1.24E-13 |
| SS10711 | 0.000452366 | 0.00171765 | 1.25E-13 |
| SS00396 | 0.002026348 | 0.004515894 | 1.25E-13 |
| SS02801 | 0 | 0.000198481 | 1.27E-13 |
| SS01852 | 0.022688483 | 0.00847033 | 1.30E-13 |
| SS06771 | 0.022688483 | 0.00847033 | 1.30E-13 |
| SS06807 | 0.022688483 | 0.00847033 | 1.30E-13 |
| SS13842 | 0.022688483 | 0.00847033 | 1.30E-13 |
| SS15521 | 0.013864436 | 0.009951626 | 1.33E-13 |
| SS08054 | 0 | 0.000388191 | 1.34E-13 |
| SS06675 | 0.00061337 | 0.001963325 | 1.35E-13 |
| SS03951 | 0.020810831 | 0.026875173 | 1.37E-13 |
| SS11564 | 0.020810831 | 0.026875173 | 1.37E-13 |
| SS11769 | 0.020810831 | 0.026875173 | 1.37E-13 |
| SS01798 | 0.026801752 | 0.016435569 | 1.47E-13 |
| SS06141 | 0.026801752 | 0.016435569 | 1.47E-13 |
| SS13474 | 0 | 0.002285997 | 1.48E-13 |
| SS01560 | 0.00179993 | 0.00766625 | 1.55E-13 |
| SS13126 | 0 | 0.001148748 | 1.57E-13 |
| SS02928 | 0 | 0.000868354 | 1.61E-13 |
| SS07019 | 5.45E-06 | 0.000660584 | 1.62E-13 |
| SS12944 | 5.45E-06 | 0.000660584 | 1.62E-13 |
| SS02366 | 0.009853284 | 0.014491019 | 1.64E-13 |
| SS03309 | 0.009853284 | 0.014491019 | 1.64E-13 |
| SS08050 | 0 | 0.000477771 | 1.64E-13 |
| SS08074 | 0 | 0.000477771 | 1.64E-13 |
| SS04266 | 0.004754523 | 0.008763815 | 1.65E-13 |
| SS01376 | 0.002593529 | 0.000818817 | 1.70E-13 |
| SS02900 | 0.002593529 | 0.000818817 | 1.70E-13 |
| SS09513 | 0.002593529 | 0.000818817 | 1.70E-13 |
| SS00949 | 0.000443912 | 0.001675125 | 1.81E-13 |
| SS00999 | 0.00085221 | 0.002620507 | 1.85E-13 |

| | | | |
|---|---|---|---|
| SS03605 | 0.00085221 | 0.002620507 | 1.85E-13 |
| SS05232 | 0.00085221 | 0.002620507 | 1.85E-13 |
| SS10430 | 0.00085221 | 0.002620507 | 1.85E-13 |
| SS10449 | 0.00085221 | 0.002620507 | 1.85E-13 |
| SS02355 | 0.009021376 | 0.005528547 | 1.89E-13 |
| SS08945 | 0.009021376 | 0.005528547 | 1.89E-13 |
| SS09618 | 0 | 0.000301476 | 1.92E-13 |
| SS01393 | 5.37E-06 | 0.000266224 | 1.93E-13 |
| SS13325 | 2.76E-06 | 0.000222758 | 1.93E-13 |
| SS03350 | 0 | 0.000927186 | 1.93E-13 |
| SS13877 | 0 | 0.000927186 | 1.93E-13 |
| SS00094 | 0 | 0.000342406 | 1.95E-13 |
| SS13513 | 0 | 0.000342406 | 1.95E-13 |
| SS04493 | 0 | 0.000669593 | 1.97E-13 |
| SS06619 | 0 | 0.002441814 | 2.02E-13 |
| SS03113 | 0.000525636 | 0.007192361 | 2.04E-13 |
| SS02734 | 0.006122631 | 0.004095182 | 2.05E-13 |
| SS13564 | 0.000449521 | 0.00011088 | 2.07E-13 |
| SS06625 | 1.90E-05 | 0.000470849 | 2.09E-13 |
| SS01406 | 0 | 0.001280019 | 2.12E-13 |
| SS06988 | 0.001199097 | 0.003166716 | 2.13E-13 |
| SS07742 | 1.63E-05 | 0.001012366 | 2.13E-13 |
| SS14499 | 2.68E-06 | 0.000141082 | 2.14E-13 |
| SS02011 | 0.000560688 | 0.001601322 | 2.18E-13 |
| SS04503 | 0.013948764 | 0.019056004 | 2.18E-13 |
| SS04573 | 0.013948764 | 0.019056004 | 2.18E-13 |
| SS04676 | 0.013948764 | 0.019056004 | 2.18E-13 |
| SS07235 | 0.020689123 | 0.001604318 | 2.19E-13 |
| SS04208 | 0 | 0.000524689 | 2.21E-13 |
| SS02845 | 4.33E-05 | 0.00068733 | 2.22E-13 |
| SS02219 | 0.000138971 | 0.000577149 | 2.26E-13 |
| SS05761 | 0 | 0.000658084 | 2.27E-13 |
| SS11812 | 0 | 0.000269272 | 2.28E-13 |
| SS09322 | 0.01551193 | 0.009819965 | 2.32E-13 |
| SS10940 | 0.01551193 | 0.009819965 | 2.32E-13 |
| SS07530 | 0.000518788 | 0.003478171 | 2.33E-13 |
| SS03959 | 0.008130432 | 0.011395074 | 2.35E-13 |
| SS11780 | 0.008130432 | 0.011395074 | 2.35E-13 |
| SS14261 | 0.008130432 | 0.011395074 | 2.35E-13 |
| SS08461 | 0.000617453 | 0.002251841 | 2.36E-13 |
| SS12402 | 0 | 0.000285448 | 2.36E-13 |
| SS06621 | 0 | 0.000722706 | 2.37E-13 |
| SS02483 | 0.010971541 | 0.01880028 | 2.41E-13 |
| SS15330 | 0.010971541 | 0.01880028 | 2.41E-13 |
| SS09982 | 0 | 0.000354398 | 2.41E-13 |
| SS00954 | 0.000754382 | 0.00234765 | 2.45E-13 |
| SS03808 | 0 | 0.001652802 | 2.50E-13 |

| | | | |
|---|---|---|---|
| SS00095 | 0 | 0.000542722 | 2.57E-13 |
| SS13522 | 0 | 0.000542722 | 2.57E-13 |
| SS04483 | 0.001931606 | 0.000443451 | 2.60E-13 |
| SS00867 | 0 | 0.001107531 | 2.63E-13 |
| SS03175 | 0 | 0.001107531 | 2.63E-13 |
| SS03712 | 0.000247855 | 0.000720316 | 2.71E-13 |
| SS07234 | 0.020691647 | 0.002112908 | 2.80E-13 |
| SS03458 | 0.026970189 | 0.008651951 | 2.80E-13 |
| SS07477 | 0.026970189 | 0.008651951 | 2.80E-13 |
| SS15093 | 0.026970189 | 0.008651951 | 2.80E-13 |
| SS04534 | 0.022093178 | 0.030052303 | 2.88E-13 |
| SS05793 | 0 | 0.000665574 | 2.90E-13 |
| SS03265 | 0.001184955 | 0.000210773 | 2.96E-13 |
| SS04006 | 0.001184955 | 0.000210773 | 2.96E-13 |
| SS08878 | 0.001184955 | 0.000210773 | 2.96E-13 |
| SS00312 | 0.015788071 | 0.010027437 | 3.08E-13 |
| SS06931 | 0.015788071 | 0.010027437 | 3.08E-13 |
| SS09949 | 0.015788071 | 0.010027437 | 3.08E-13 |
| SS10659 | 0 | 0.000373123 | 3.25E-13 |
| SS03804 | 0 | 0.001587307 | 3.32E-13 |
| SS09119 | 0.000912703 | 0.000265476 | 3.40E-13 |
| SS02820 | 0 | 0.000206455 | 3.64E-13 |
| SS10865 | 0.027958938 | 0.036727206 | 3.64E-13 |
| SS15537 | 0.027958938 | 0.036727206 | 3.64E-13 |
| SS15470 | 0.009999171 | 0.019251151 | 3.71E-13 |
| SS05881 | 0.000337711 | 0.000138077 | 3.77E-13 |
| SS00139 | 0.000959695 | 0.00300806 | 3.80E-13 |
| SS06466 | 0.029567445 | 0.040171374 | 3.80E-13 |
| SS09553 | 0.029567445 | 0.040171374 | 3.80E-13 |
| SS10460 | 0.029567445 | 0.040171374 | 3.80E-13 |
| SS10960 | 0.002664195 | 0.001084548 | 3.80E-13 |
| SS02669 | 7.35E-05 | 0.000757973 | 3.83E-13 |
| SS09413 | 0 | 0.000192107 | 3.85E-13 |
| SS09420 | 0.002590523 | 0.000215383 | 3.87E-13 |
| SS10467 | 0.010884437 | 0.007520912 | 3.92E-13 |
| SS10658 | 0 | 0.000257265 | 4.06E-13 |
| SS01567 | 0.020790793 | 0.027853357 | 4.13E-13 |
| SS04011 | 0.020790793 | 0.027853357 | 4.13E-13 |
| SS15428 | 0.020790793 | 0.027853357 | 4.13E-13 |
| SS03133 | 0.020831213 | 0.012008329 | 4.25E-13 |
| SS04273 | 0.020831213 | 0.012008329 | 4.25E-13 |
| SS08772 | 0.020831213 | 0.012008329 | 4.25E-13 |
| SS07756 | 0 | 0.000749852 | 4.35E-13 |
| SS02270 | 0.009492013 | 0.007119536 | 4.36E-13 |
| SS03535 | 0.009492013 | 0.007119536 | 4.36E-13 |
| SS03600 | 0.009492013 | 0.007119536 | 4.36E-13 |
| SS03699 | 0.009492013 | 0.007119536 | 4.36E-13 |

| | | | |
|---|---|---|---|
| SS04448 | 0.009492013 | 0.007119536 | 4.36E-13 |
| SS05198 | 0.009492013 | 0.007119536 | 4.36E-13 |
| SS05499 | 0.009492013 | 0.007119536 | 4.36E-13 |
| SS08045 | 0.009492013 | 0.007119536 | 4.36E-13 |
| SS10424 | 0.009492013 | 0.007119536 | 4.36E-13 |
| SS15354 | 0.009492013 | 0.007119536 | 4.36E-13 |
| SS00661 | 0.004479381 | 0.001735556 | 4.42E-13 |
| SS06939 | 0.004479381 | 0.001735556 | 4.42E-13 |
| SS06977 | 0.004479381 | 0.001735556 | 4.42E-13 |
| SS07187 | 0.004479381 | 0.001735556 | 4.42E-13 |
| SS11083 | 0.032282957 | 0.040831546 | 4.47E-13 |
| SS08433 | 0.002374314 | 0.000881578 | 4.49E-13 |
| SS08470 | 0.002374314 | 0.000881578 | 4.49E-13 |
| SS03421 | 0.001618051 | 0.003078073 | 4.52E-13 |
| SS05768 | 0 | 0.00085526 | 4.60E-13 |
| SS06192 | 0 | 0.001646816 | 4.69E-13 |
| SS02749 | 0.000280705 | 0.00078297 | 4.71E-13 |
| SS12294 | 0.000280705 | 0.00078297 | 4.71E-13 |
| SS04839 | 0 | 0.001020387 | 4.72E-13 |
| SS13874 | 0 | 0.001020387 | 4.72E-13 |
| SS15465 | 0.006994913 | 0.009584288 | 4.76E-13 |
| SS15473 | 0.006994913 | 0.009584288 | 4.76E-13 |
| SS06709 | 0.022838132 | 0.014408644 | 4.89E-13 |
| SS10963 | 0.012366488 | 0.008125896 | 4.94E-13 |
| SS03508 | 0.020868365 | 0.009812562 | 4.95E-13 |
| SS13912 | 0.020868365 | 0.009812562 | 4.95E-13 |
| SS14085 | 0 | 0.000236107 | 5.14E-13 |
| SS08292 | 0.002314624 | 0.006009668 | 5.20E-13 |
| SS03803 | 2.68E-06 | 0.001212132 | 5.23E-13 |
| SS02224 | 2.18E-05 | 0.000306549 | 5.39E-13 |
| SS11599 | 2.76E-06 | 0.0010951 | 5.47E-13 |
| SS07499 | 0.002842106 | 0.009326067 | 5.48E-13 |
| SS05773 | 0 | 0.000498679 | 5.57E-13 |
| SS14071 | 0 | 0.000187891 | 5.78E-13 |
| SS15037 | 3.87E-05 | 0.001251461 | 5.95E-13 |
| SS00902 | 0 | 0.001055807 | 6.05E-13 |
| SS15290 | 8.05E-06 | 0.000749715 | 6.07E-13 |
| SS00403 | 0.001253018 | 9.39E-05 | 6.28E-13 |
| SS13733 | 0.007472708 | 0.014127964 | 6.36E-13 |
| SS13638 | 0.001831301 | 0.010218422 | 6.67E-13 |
| SS15325 | 0 | 0.000281421 | 6.71E-13 |
| SS03502 | 0.013250309 | 0.019219201 | 6.87E-13 |
| SS04268 | 0.013250309 | 0.019219201 | 6.87E-13 |
| SS07475 | 0.013250309 | 0.019219201 | 6.87E-13 |
| SS08714 | 0.013250309 | 0.019219201 | 6.87E-13 |
| SS10697 | 0.013250309 | 0.019219201 | 6.87E-13 |
| SS00327 | 0.01507591 | 0.019690341 | 6.88E-13 |

| | | | |
|---|---|---|---|
| SS01813 | 0.01507591 | 0.019690341 | 6.88E-13 |
| SS03953 | 0.01507591 | 0.019690341 | 6.88E-13 |
| SS09041 | 0.01507591 | 0.019690341 | 6.88E-13 |
| SS13110 | 0.01507591 | 0.019690341 | 6.88E-13 |
| SS07995 | 0 | 0.000294989 | 7.00E-13 |
| SS04020 | 0.002165319 | 5.06E-05 | 7.05E-13 |
| SS05992 | 0.018629042 | 0.006518369 | 7.12E-13 |
| SS06007 | 0.018629042 | 0.006518369 | 7.12E-13 |
| SS08911 | 0.018629042 | 0.006518369 | 7.12E-13 |
| SS12996 | 0.018629042 | 0.006518369 | 7.12E-13 |
| SS02822 | 0 | 0.000904864 | 7.14E-13 |
| SS07016 | 0.012794457 | 0.004950983 | 7.18E-13 |
| SS07687 | 0.012794457 | 0.004950983 | 7.18E-13 |
| SS01185 | 0.002671605 | 0.001712209 | 7.33E-13 |
| SS00900 | 0 | 0.001337911 | 7.38E-13 |
| SS09959 | 0 | 0.001337911 | 7.38E-13 |
| SS15154 | 0.000247936 | 0.00069184 | 7.39E-13 |
| SS04560 | 0.001676697 | 0.005707513 | 7.61E-13 |
| SS03126 | 0.0170436 | 0.011175249 | 7.64E-13 |
| SS07288 | 0.0170436 | 0.011175249 | 7.64E-13 |
| SS06029 | 0.026385104 | 0.003993351 | 7.79E-13 |
| SS10716 | 0 | 0.001202194 | 7.94E-13 |
| SS07768 | 0 | 0.000312117 | 8.07E-13 |
| SS11759 | 0.001039733 | 0.002473366 | 8.15E-13 |
| SS14485 | 0.001039733 | 0.002473366 | 8.15E-13 |
| SS01544 | 8.13E-06 | 0.000426979 | 8.23E-13 |
| SS01682 | 0 | 0.000369644 | 8.27E-13 |
| SS01029 | 0 | 0.001658214 | 8.31E-13 |
| SS00883 | 2.76E-06 | 0.001505324 | 8.60E-13 |
| SS12461 | 2.76E-06 | 0.001505324 | 8.60E-13 |
| SS10580 | 0 | 0.000212444 | 8.69E-13 |
| SS11039 | 0 | 0.001556058 | 8.77E-13 |
| SS06720 | 0.025921441 | 0.00938301 | 8.81E-13 |
| SS07889 | 0.006709356 | 0.000262368 | 8.88E-13 |
| SS10043 | 0 | 0.000323909 | 8.91E-13 |
| SS01716 | 0.01145827 | 0.016074307 | 9.25E-13 |
| SS06062 | 0.01145827 | 0.016074307 | 9.25E-13 |
| SS13542 | 0.01145827 | 0.016074307 | 9.25E-13 |
| SS13561 | 0.01145827 | 0.016074307 | 9.25E-13 |
| SS11265 | 0.008024036 | 0.014260624 | 9.59E-13 |
| SS08384 | 0 | 0.000202181 | 9.64E-13 |
| SS02178 | 0.033934673 | 0.049597458 | 9.82E-13 |
| SS06883 | 0.033934673 | 0.049597458 | 9.82E-13 |
| SS12682 | 0.033934673 | 0.049597458 | 9.82E-13 |
| SS14222 | 0.033934673 | 0.049597458 | 9.82E-13 |
| SS00435 | 0.011068795 | 0.003060097 | 1.03E-12 |
| SS06594 | 0 | 0.00034541 | 1.06E-12 |

| | | | |
|---|---|---|---|
| SS01631 | 0.016530513 | 0.002073084 | 1.07E-12 |
| SS02373 | 0.016530513 | 0.002073084 | 1.07E-12 |
| SS10534 | 0.006755592 | 0.001829348 | 1.07E-12 |
| SS07513 | 0 | 0.000238104 | 1.07E-12 |
| SS06626 | 2.17E-05 | 0.002206876 | 1.09E-12 |
| SS02490 | 0 | 0.000650377 | 1.09E-12 |
| SS08727 | 0 | 0.000271058 | 1.10E-12 |
| SS09925 | 0 | 0.000271058 | 1.10E-12 |
| SS02449 | 0 | 0.000390148 | 1.10E-12 |
| SS03290 | 0 | 0.000390148 | 1.10E-12 |
| SS10582 | 0 | 0.000227333 | 1.11E-12 |
| SS02805 | 0 | 0.002390804 | 1.13E-12 |
| SS04811 | 0.009065811 | 0.015358026 | 1.15E-12 |
| SS11148 | 0.009065811 | 0.015358026 | 1.15E-12 |
| SS03377 | 0.041851797 | 0.02690135 | 1.18E-12 |
| SS04429 | 0.041851797 | 0.02690135 | 1.18E-12 |
| SS06288 | 0.041851797 | 0.02690135 | 1.18E-12 |
| SS11205 | 0.01393906 | 0.008357409 | 1.19E-12 |
| SS11208 | 0.01393906 | 0.008357409 | 1.19E-12 |
| SS06096 | 0 | 0.000594273 | 1.19E-12 |
| SS04388 | 0.024541037 | 0.017194779 | 1.19E-12 |
| SS04988 | 0.024541037 | 0.017194779 | 1.19E-12 |
| SS11109 | 0.024541037 | 0.017194779 | 1.19E-12 |
| SS04799 | 0.000286073 | 8.93E-05 | 1.26E-12 |
| SS02482 | 0.003590398 | 0.00210014 | 1.31E-12 |
| SS15326 | 0.003590398 | 0.00210014 | 1.31E-12 |
| SS08049 | 0 | 0.000268737 | 1.32E-12 |
| SS10300 | 0.001851522 | 0.000933103 | 1.34E-12 |
| SS15048 | 0.001440782 | 0.002966771 | 1.35E-12 |
| SS08597 | 0.013894843 | 0.00366318 | 1.35E-12 |
| SS13252 | 5.53E-06 | 0.001005999 | 1.38E-12 |
| SS03939 | 0.048171345 | 0.058271523 | 1.41E-12 |
| SS04693 | 0.048171345 | 0.058271523 | 1.41E-12 |
| SS10592 | 0.048171345 | 0.058271523 | 1.41E-12 |
| SS11034 | 1.63E-05 | 3.71E-07 | 1.46E-12 |
| SS03026 | 0 | 0.00037011 | 1.48E-12 |
| SS03747 | 0 | 0.00037011 | 1.48E-12 |
| SS00008 | 0.01765213 | 0.022722193 | 1.49E-12 |
| SS01207 | 0.01765213 | 0.022722193 | 1.49E-12 |
| SS11562 | 0.007318321 | 0.005079467 | 1.57E-12 |
| SS11767 | 0.007318321 | 0.005079467 | 1.57E-12 |
| SS01755 | 0.008326122 | 0.013019447 | 1.58E-12 |
| SS01796 | 0 | 0.000706557 | 1.60E-12 |
| SS06170 | 0 | 0.000706557 | 1.60E-12 |
| SS03781 | 0.024418136 | 0.031392917 | 1.60E-12 |
| SS07259 | 0.024418136 | 0.031392917 | 1.60E-12 |
| SS15404 | 0.024418136 | 0.031392917 | 1.60E-12 |

| | | | |
|---|---|---|---|
| SS13615 | 0.013758316 | 0.021054185 | 1.61E-12 |
| SS08115 | 0.001132835 | 0.002611856 | 1.66E-12 |
| SS08549 | 0.001132835 | 0.002611856 | 1.66E-12 |
| SS07779 | 8.21E-06 | 0.000740024 | 1.66E-12 |
| SS05246 | 5.19E-05 | 0.000999883 | 1.69E-12 |
| SS01449 | 3.81E-05 | 0.000493542 | 1.73E-12 |
| SS13050 | 3.81E-05 | 0.000493542 | 1.73E-12 |
| SS01914 | 0 | 0.000312716 | 1.77E-12 |
| SS05518 | 0 | 0.000909081 | 1.79E-12 |
| SS04390 | 0.004884719 | 0.001793939 | 1.80E-12 |
| SS13198 | 0 | 0.000398257 | 1.83E-12 |
| SS01461 | 0 | 0.000354823 | 1.97E-12 |
| SS06446 | 0 | 0.000354823 | 1.97E-12 |
| SS07298 | 0 | 0.000246508 | 2.11E-12 |
| SS05192 | 0.023111554 | 0.01803324 | 2.11E-12 |
| SS06497 | 0.023111554 | 0.01803324 | 2.11E-12 |
| SS12223 | 0.000969789 | 0.002377298 | 2.14E-12 |
| SS07993 | 0 | 0.000300337 | 2.18E-12 |
| SS04667 | 0.005780951 | 0.001273244 | 2.19E-12 |
| SS10922 | 0.005780951 | 0.001273244 | 2.19E-12 |
| SS11214 | 0.007649782 | 0.010131819 | 2.19E-12 |
| SS15107 | 2.45E-05 | 0.000229248 | 2.24E-12 |
| SS03346 | 0.001375322 | 0.002776713 | 2.32E-12 |
| SS03477 | 0.001375322 | 0.002776713 | 2.32E-12 |
| SS05671 | 0.001375322 | 0.002776713 | 2.32E-12 |
| SS00713 | 0 | 0.000906103 | 2.34E-12 |
| SS07877 | 0 | 0.000906103 | 2.34E-12 |
| SS07558 | 0.008374961 | 0.014141478 | 2.34E-12 |
| SS11286 | 0.008374961 | 0.014141478 | 2.34E-12 |
| SS05912 | 0 | 0.000643332 | 2.37E-12 |
| SS09996 | 0.029872834 | 0.046544424 | 2.48E-12 |
| SS03576 | 0.007284428 | 0.001527387 | 2.51E-12 |
| SS13880 | 0.000280786 | 0.00157419 | 2.63E-12 |
| SS15003 | 0.000280786 | 0.00157419 | 2.63E-12 |
| SS03473 | 0 | 0.000820411 | 2.67E-12 |
| SS05669 | 0 | 0.000820411 | 2.67E-12 |
| SS00885 | 0 | 0.00085547 | 2.71E-12 |
| SS04102 | 5.37E-06 | 0.000622212 | 2.79E-12 |
| SS01904 | 0 | 0.000144401 | 2.89E-12 |
| SS11143 | 0 | 0.000144401 | 2.89E-12 |
| SS06253 | 0.013519121 | 0.023743664 | 2.94E-12 |
| SS14360 | 0.013519121 | 0.023743664 | 2.94E-12 |
| SS12502 | 0.001356213 | 1.79E-05 | 2.98E-12 |
| SS08161 | 0.007904611 | 0.001017044 | 2.98E-12 |
| SS08843 | 0.007904611 | 0.001017044 | 2.98E-12 |
| SS10261 | 6.81E-05 | 2.07E-05 | 3.00E-12 |
| SS11123 | 0.000542141 | 0.001412644 | 3.02E-12 |

| | | | |
|---|---|---|---|
| SS08585 | 0 | 0.000574089 | 3.05E-12 |
| SS02126 | 0.027485215 | 0.01921094 | 3.05E-12 |
| SS05994 | 0.027485215 | 0.01921094 | 3.05E-12 |
| SS06571 | 0 | 0.000204424 | 3.11E-12 |
| SS13179 | 0 | 0.000367126 | 3.11E-12 |
| SS04901 | 0 | 0.00111365 | 3.15E-12 |
| SS13903 | 0 | 0.00111365 | 3.15E-12 |
| SS06635 | 0 | 0.000186372 | 3.15E-12 |
| SS05911 | 0.000871399 | 7.69E-05 | 3.20E-12 |
| SS15205 | 0.000463101 | 0.0019911 | 3.22E-12 |
| SS10671 | 0.011397984 | 0.016468782 | 3.31E-12 |
| SS04648 | 0.007101148 | 0.010517008 | 3.34E-12 |
| SS11803 | 0.007101148 | 0.010517008 | 3.34E-12 |
| SS02858 | 0.015494829 | 0.008982415 | 3.40E-12 |
| SS12030 | 0.015494829 | 0.008982415 | 3.40E-12 |
| SS08077 | 0 | 0.000290998 | 3.47E-12 |
| SS07926 | 0 | 0.001743198 | 3.48E-12 |
| SS10012 | 0.002410892 | 0.001371352 | 3.62E-12 |
| SS11589 | 0.002430597 | 0.005349174 | 3.64E-12 |
| SS02203 | 0.010776642 | 0.006461685 | 3.65E-12 |
| SS14877 | 6.81E-05 | 1.84E-05 | 3.68E-12 |
| SS04504 | 2.76E-06 | 0.000603809 | 3.73E-12 |
| SS09405 | 0.002722922 | 0.004682959 | 3.77E-12 |
| SS00490 | 0.038710347 | 0.024879277 | 3.77E-12 |
| SS03127 | 0.010313461 | 0.004886529 | 3.84E-12 |
| SS03437 | 0.011622991 | 0.019128338 | 3.86E-12 |
| SS05778 | 0.011622991 | 0.019128338 | 3.86E-12 |
| SS10417 | 0.011622991 | 0.019128338 | 3.86E-12 |
| SS10436 | 0.011622991 | 0.019128338 | 3.86E-12 |
| SS11814 | 0.011622991 | 0.019128338 | 3.86E-12 |
| SS08364 | 0.002272724 | 0.007363466 | 3.92E-12 |
| SS01861 | 0 | 0.000405291 | 3.93E-12 |
| SS06315 | 0 | 0.000405291 | 3.93E-12 |
| SS06415 | 0 | 0.000405291 | 3.93E-12 |
| SS13611 | 0 | 0.000405291 | 3.93E-12 |
| SS06478 | 0.02298239 | 0.031482567 | 4.03E-12 |
| SS09112 | 0.00580763 | 0.002284507 | 4.04E-12 |
| SS10829 | 0.00580763 | 0.002284507 | 4.04E-12 |
| SS15247 | 0 | 0.000250051 | 4.09E-12 |
| SS08175 | 0.014176352 | 0.008024192 | 4.24E-12 |
| SS10147 | 0.014176352 | 0.008024192 | 4.24E-12 |
| SS02806 | 0 | 0.000632449 | 4.38E-12 |
| SS04981 | 0.034891937 | 0.056559883 | 4.43E-12 |
| SS06135 | 0.034891937 | 0.056559883 | 4.43E-12 |
| SS06167 | 0.034891937 | 0.056559883 | 4.43E-12 |
| SS11915 | 0.034891937 | 0.056559883 | 4.43E-12 |
| SS13917 | 0.000890749 | 0.002645547 | 4.45E-12 |

| | | | |
|---|---|---|---|
| SS10521 | 0.004077174 | 0.002892149 | 4.50E-12 |
| SS03035 | 4.59E-05 | 0.002364552 | 4.51E-12 |
| SS09461 | 4.59E-05 | 0.002364552 | 4.51E-12 |
| SS11596 | 0 | 0.000722046 | 4.53E-12 |
| SS00488 | 0.008981518 | 0.004765847 | 4.59E-12 |
| SS03326 | 0 | 0.000736421 | 4.65E-12 |
| SS04662 | 0.003102498 | 0.000220051 | 4.70E-12 |
| SS10906 | 0.003102498 | 0.000220051 | 4.70E-12 |
| SS06613 | 0 | 0.000236931 | 4.81E-12 |
| SS00996 | 0.011611177 | 0.008001446 | 4.82E-12 |
| SS06785 | 0.011611177 | 0.008001446 | 4.82E-12 |
| SS09825 | 0 | 0.000867753 | 4.84E-12 |
| SS00549 | 0 | 0.000262666 | 4.85E-12 |
| SS06678 | 0 | 0.000262666 | 4.85E-12 |
| SS02181 | 0.000550273 | 0.002136351 | 4.88E-12 |
| SS06889 | 0.000550273 | 0.002136351 | 4.88E-12 |
| SS00854 | 0.008103466 | 0.004409571 | 4.96E-12 |
| SS11593 | 0 | 0.001156454 | 5.25E-12 |
| SS02273 | 0.007742127 | 0.004248946 | 5.31E-12 |
| SS03539 | 0.007742127 | 0.004248946 | 5.31E-12 |
| SS03606 | 0.007742127 | 0.004248946 | 5.31E-12 |
| SS03702 | 0.007742127 | 0.004248946 | 5.31E-12 |
| SS04449 | 0.007742127 | 0.004248946 | 5.31E-12 |
| SS05206 | 0.007742127 | 0.004248946 | 5.31E-12 |
| SS05506 | 0.007742127 | 0.004248946 | 5.31E-12 |
| SS08055 | 0.007742127 | 0.004248946 | 5.31E-12 |
| SS10431 | 0.007742127 | 0.004248946 | 5.31E-12 |
| SS01216 | 0 | 0.000215596 | 5.33E-12 |
| SS08607 | 6.81E-05 | 8.56E-06 | 5.56E-12 |
| SS03800 | 0 | 0.000596852 | 5.91E-12 |
| SS05934 | 0.004744223 | 7.57E-05 | 6.18E-12 |
| SS08197 | 0.027835636 | 0.036050298 | 6.23E-12 |
| SS11220 | 0.027835636 | 0.036050298 | 6.23E-12 |
| SS14127 | 0.027835636 | 0.036050298 | 6.23E-12 |
| SS14192 | 0.027835636 | 0.036050298 | 6.23E-12 |
| SS14259 | 0.027835636 | 0.036050298 | 6.23E-12 |
| SS14366 | 0.027835636 | 0.036050298 | 6.23E-12 |
| SS15423 | 0.027835636 | 0.036050298 | 6.23E-12 |
| SS07548 | 0 | 0.001365217 | 6.60E-12 |
| SS05898 | 0 | 0.000506004 | 6.61E-12 |
| SS15392 | 0 | 0.000506004 | 6.61E-12 |
| SS14571 | 0.003186665 | 0.001530005 | 6.69E-12 |
| SS14658 | 0.003186665 | 0.001530005 | 6.69E-12 |
| SS06444 | 0.014487017 | 0.00021544 | 7.18E-12 |
| SS03111 | 0.000396839 | 0.002240709 | 7.31E-12 |
| SS04609 | 0.014785111 | 0.001003964 | 7.42E-12 |
| SS02869 | 0.000133442 | 4.16E-05 | 7.51E-12 |

| | | | |
|---|---|---|---|
| SS02578 | 0.004882551 | 0.000501555 | 7.73E-12 |
| SS10756 | 0 | 0.000478645 | 7.79E-12 |
| SS00896 | 5.53E-06 | 0.001074326 | 7.81E-12 |
| SS07713 | 0.008096183 | 0.026488698 | 7.89E-12 |
| SS04238 | 0.020349336 | 0.015756054 | 7.89E-12 |
| SS02785 | 0.001833618 | 0.004483134 | 7.99E-12 |
| SS15518 | 0.000647941 | 0.0022714 | 8.02E-12 |
| SS07526 | 2.68E-06 | 0.000737536 | 8.06E-12 |
| SS13092 | 0 | 0.000287254 | 8.14E-12 |
| SS13940 | 0 | 0.000287254 | 8.14E-12 |
| SS03205 | 0 | 0.001322861 | 8.21E-12 |
| SS04606 | 0.02428825 | 0.003015914 | 8.30E-12 |
| SS04650 | 0.02428825 | 0.003015914 | 8.30E-12 |
| SS04824 | 0.02428825 | 0.003015914 | 8.30E-12 |
| SS01209 | 0.008359695 | 0.00235127 | 8.30E-12 |
| SS01274 | 0.008359695 | 0.00235127 | 8.30E-12 |
| SS09809 | 0 | 0.000313128 | 8.65E-12 |
| SS13576 | 0 | 0.000313128 | 8.65E-12 |
| SS00073 | 0.023883967 | 0.018609488 | 8.81E-12 |
| SS06069 | 0.023883967 | 0.018609488 | 8.81E-12 |
| SS07183 | 0.023883967 | 0.018609488 | 8.81E-12 |
| SS09630 | 0.023883967 | 0.018609488 | 8.81E-12 |
| SS13265 | 0.023883967 | 0.018609488 | 8.81E-12 |
| SS03713 | 4.08E-05 | 0.000440891 | 8.92E-12 |
| SS11861 | 0.000182476 | 0.00048749 | 8.97E-12 |
| SS13914 | 0.001141047 | 0.00387601 | 9.07E-12 |
| SS13506 | 0.01477153 | 0.002684111 | 9.20E-12 |
| SS00119 | 5.37E-06 | 0.000228284 | 9.26E-12 |
| SS08700 | 1.09E-05 | 0.0008773 | 9.48E-12 |
| SS00198 | 0 | 0.000366778 | 9.84E-12 |
| SS07634 | 0 | 0.000366778 | 9.84E-12 |
| SS09912 | 0 | 0.000366778 | 9.84E-12 |
| SS07782 | 0 | 0.00041096 | 9.89E-12 |
| SS13020 | 0 | 0.00041096 | 9.89E-12 |
| SS01842 | 0.001748889 | 0.00035969 | 1.04E-11 |
| SS10550 | 0.001748889 | 0.00035969 | 1.04E-11 |
| SS11497 | 6.26E-05 | 1.38E-05 | 1.06E-11 |
| SS00722 | 0 | 0.000369563 | 1.06E-11 |
| SS00727 | 0 | 0.000369563 | 1.06E-11 |
| SS14134 | 0 | 0.000369563 | 1.06E-11 |
| SS11570 | 0.014497156 | 0.006543188 | 1.11E-11 |
| SS11583 | 0.014497156 | 0.006543188 | 1.11E-11 |
| SS11777 | 0.014497156 | 0.006543188 | 1.11E-11 |
| SS07924 | 0.00068656 | 0.000231141 | 1.11E-11 |
| SS02861 | 2.76E-05 | 0.000878853 | 1.19E-11 |
| SS00120 | 5.37E-06 | 0.000219397 | 1.24E-11 |
| SS07574 | 0.000326494 | 0.001081838 | 1.36E-11 |

| | | | |
|---|---|---|---|
| SS08634 | 0.000326494 | 0.001081838 | 1.36E-11 |
| SS07613 | 0.00217898 | 6.59E-05 | 1.36E-11 |
| SS14809 | 0.049654761 | 0.012141416 | 1.37E-11 |
| SS15065 | 0.049654761 | 0.012141416 | 1.37E-11 |
| SS12341 | 0 | 0.002622211 | 1.38E-11 |
| SS13620 | 0.006189054 | 0.003989722 | 1.38E-11 |
| SS02428 | 0.01005378 | 0.013739108 | 1.39E-11 |
| SS03094 | 0.01005378 | 0.013739108 | 1.39E-11 |
| SS07165 | 0.01005378 | 0.013739108 | 1.39E-11 |
| SS08696 | 0.01005378 | 0.013739108 | 1.39E-11 |
| SS13274 | 2.68E-06 | 0.000188225 | 1.40E-11 |
| SS04489 | 0.055127558 | 0.069001384 | 1.43E-11 |
| SS04640 | 0.055127558 | 0.069001384 | 1.43E-11 |
| SS04706 | 0.055127558 | 0.069001384 | 1.43E-11 |
| SS11046 | 0.055127558 | 0.069001384 | 1.43E-11 |
| SS15307 | 0.005499007 | 0.008530337 | 1.46E-11 |
| SS08827 | 3.27E-05 | 5.84E-06 | 1.48E-11 |
| SS13417 | 0.007204906 | 0.002759586 | 1.48E-11 |
| SS13427 | 0.007204906 | 0.002759586 | 1.48E-11 |
| SS04785 | 0.000351212 | 0.000100374 | 1.48E-11 |
| SS05004 | 0 | 0.000189202 | 1.48E-11 |
| SS01839 | 0.006327864 | 0.001364609 | 1.49E-11 |
| SS02574 | 0.006327864 | 0.001364609 | 1.49E-11 |
| SS02019 | 0 | 0.000395309 | 1.50E-11 |
| SS08375 | 0 | 0.000395309 | 1.50E-11 |
| SS06714 | 0.001933487 | 0.004301564 | 1.51E-11 |
| SS10069 | 0.003528024 | 0.001697345 | 1.56E-11 |
| SS04728 | 0 | 0.000161855 | 1.56E-11 |
| SS10013 | 0.005497447 | 0.003070084 | 1.56E-11 |
| SS02035 | 0.000828295 | 0.000206863 | 1.61E-11 |
| SS03856 | 2.68E-06 | 0.001335783 | 1.68E-11 |
| SS03887 | 2.68E-06 | 0.001335783 | 1.68E-11 |
| SS04222 | 0 | 0.000157815 | 1.72E-11 |
| SS02047 | 0.00120911 | 0.000362452 | 1.73E-11 |
| SS05696 | 0.008074344 | 0.000970736 | 1.73E-11 |
| SS10576 | 0.008074344 | 0.000970736 | 1.73E-11 |
| SS08701 | 2.46E-05 | 0.000644732 | 1.80E-11 |
| SS06305 | 0 | 0.000404678 | 1.81E-11 |
| SS06321 | 0 | 0.000230928 | 1.82E-11 |
| SS06421 | 0 | 0.000230928 | 1.82E-11 |
| SS11055 | 0.000758592 | 0.003763096 | 1.84E-11 |
| SS11134 | 0.00127449 | 0.000765606 | 1.85E-11 |
| SS07570 | 0 | 0.000172662 | 1.87E-11 |
| SS00658 | 0.000160523 | 0.000938764 | 1.87E-11 |
| SS06028 | 0.000160523 | 0.000938764 | 1.87E-11 |
| SS11111 | 0.000160523 | 0.000938764 | 1.87E-11 |
| SS05046 | 0.000166212 | 2.36E-05 | 1.88E-11 |

| | | | |
|---|---|---|---|
| SS11597 | 2.76E-06 | 0.000487395 | 1.98E-11 |
| SS05072 | 9.27E-05 | 0.000949244 | 2.02E-11 |
| SS02832 | 0.001434932 | 0.002526592 | 2.03E-11 |
| SS01034 | 0.001860538 | 0.000248162 | 2.03E-11 |
| SS04918 | 0.001860538 | 0.000248162 | 2.03E-11 |
| SS03801 | 0 | 0.00048638 | 2.07E-11 |
| SS03659 | 0.005633573 | 0.001356825 | 2.07E-11 |
| SS07071 | 0.005633573 | 0.001356825 | 2.07E-11 |
| SS03807 | 0 | 0.000888005 | 2.08E-11 |
| SS00670 | 0 | 0.000419749 | 2.14E-11 |
| SS03559 | 0 | 0.000419749 | 2.14E-11 |
| SS09997 | 0.005594907 | 0.010453691 | 2.16E-11 |
| SS08627 | 0 | 0.000462302 | 2.18E-11 |
| SS14884 | 0.04229673 | 0.054372794 | 2.19E-11 |
| SS04491 | 0 | 0.000360429 | 2.21E-11 |
| SS15067 | 0.002355331 | 0.006508025 | 2.21E-11 |
| SS09411 | 0 | 0.000241758 | 2.24E-11 |
| SS06352 | 2.68E-06 | 0.000446929 | 2.25E-11 |
| SS06858 | 2.68E-06 | 0.000446929 | 2.25E-11 |
| SS02994 | 0 | 0.000282167 | 2.28E-11 |
| SS08359 | 0.000255907 | 0.001457256 | 2.35E-11 |
| SS03245 | 0.000825049 | 0.002191688 | 2.36E-11 |
| SS03338 | 0.000825049 | 0.002191688 | 2.36E-11 |
| SS01405 | 0 | 0.000226002 | 2.43E-11 |
| SS03022 | 0 | 0.00043229 | 2.47E-11 |
| SS03743 | 0 | 0.00043229 | 2.47E-11 |
| SS03725 | 0.007710997 | 0.005222337 | 2.47E-11 |
| SS05463 | 0.000805779 | 0.001943592 | 2.51E-11 |
| SS02459 | 2.76E-06 | 0.00483451 | 2.53E-11 |
| SS05324 | 0.014669057 | 0.024409695 | 2.56E-11 |
| SS00773 | 0.014468275 | 0.021713958 | 2.57E-11 |
| SS02650 | 0.022596527 | 0.029309313 | 2.64E-11 |
| SS04540 | 0.022596527 | 0.029309313 | 2.64E-11 |
| SS00822 | 0 | 0.000885463 | 2.69E-11 |
| SS11446 | 0.009486118 | 0.014157176 | 2.71E-11 |
| SS05775 | 0 | 0.00031011 | 2.72E-11 |
| SS05953 | 0.003854036 | 0.000116924 | 2.73E-11 |
| SS09977 | 0.003854036 | 0.000116924 | 2.73E-11 |
| SS05413 | 0.006535138 | 0.004107395 | 2.78E-11 |
| SS05422 | 0.006535138 | 0.004107395 | 2.78E-11 |
| SS02349 | 0 | 0.000176473 | 2.79E-11 |
| SS00606 | 0 | 0.000232309 | 2.96E-11 |
| SS13495 | 0.000624026 | 0.00224185 | 2.97E-11 |
| SS01286 | 0 | 0.000957356 | 3.04E-11 |
| SS06979 | 0 | 0.00024099 | 3.06E-11 |
| SS14426 | 1.91E-05 | 0.000423565 | 3.13E-11 |
| SS11262 | 0.008103948 | 0.011210341 | 3.24E-11 |

| | | | |
|---|---|---|---|
| SS07556 | 0.011447982 | 0.017753087 | 3.25E-11 |
| SS11256 | 0.011447982 | 0.017753087 | 3.25E-11 |
| SS00090 | 0.008705137 | 0.002850596 | 3.28E-11 |
| SS03590 | 0.008705137 | 0.002850596 | 3.28E-11 |
| SS13507 | 0.008705137 | 0.002850596 | 3.28E-11 |
| SS02210 | 0 | 0.000156462 | 3.32E-11 |
| SS10839 | 0.020024115 | 0.025312188 | 3.36E-11 |
| SS14238 | 0.020024115 | 0.025312188 | 3.36E-11 |
| SS14280 | 0.020024115 | 0.025312188 | 3.36E-11 |
| SS09069 | 0.01428651 | 0.001038728 | 3.38E-11 |
| SS01643 | 0.000329579 | 0.000145536 | 3.48E-11 |
| SS07722 | 0 | 0.000477934 | 3.49E-11 |
| SS05226 | 0.007604234 | 0.010543167 | 3.50E-11 |
| SS14813 | 0.007604234 | 0.010543167 | 3.50E-11 |
| SS10366 | 0.0020731 | 0.003577487 | 3.51E-11 |
| SS10223 | 0.004804682 | 0.012157834 | 3.57E-11 |
| SS07770 | 0 | 0.000133265 | 3.65E-11 |
| SS12974 | 0 | 0.000133265 | 3.65E-11 |
| SS06313 | 7.90E-05 | 0.000763974 | 3.72E-11 |
| SS13381 | 0.005036674 | 0.011758158 | 3.76E-11 |
| SS03383 | 0 | 0.00017028 | 3.81E-11 |
| SS09432 | 1.63E-05 | 0.00063851 | 3.90E-11 |
| SS06254 | 0.01509979 | 0.008783919 | 3.91E-11 |
| SS11241 | 0.01509979 | 0.008783919 | 3.91E-11 |
| SS07581 | 0 | 0.000942436 | 4.00E-11 |
| SS03006 | 0.02647387 | 0.039048984 | 4.05E-11 |
| SS14470 | 0.02647387 | 0.039048984 | 4.05E-11 |
| SS10249 | 0.000198901 | 5.69E-05 | 4.07E-11 |
| SS05789 | 0.001037611 | 0.003075485 | 4.10E-11 |
| SS05588 | 0 | 0.000461118 | 4.11E-11 |
| SS02221 | 0 | 0.000519017 | 4.16E-11 |
| SS04340 | 0.000829453 | 0.002926522 | 4.18E-11 |
| SS13035 | 0.000829453 | 0.002926522 | 4.18E-11 |
| SS01838 | 0.010457512 | 0.000768576 | 4.22E-11 |
| SS13139 | 0 | 0.000170026 | 4.26E-11 |
| SS15311 | 0 | 0.000170026 | 4.26E-11 |
| SS01880 | 2.76E-06 | 0.00018862 | 4.27E-11 |
| SS15210 | 5.53E-06 | 0.000467857 | 4.46E-11 |
| SS11037 | 1.08E-05 | 0.000203771 | 4.49E-11 |
| SS02057 | 0.00939547 | 0.014427585 | 4.63E-11 |
| SS02897 | 0.00939547 | 0.014427585 | 4.63E-11 |
| SS10360 | 0.00939547 | 0.014427585 | 4.63E-11 |
| SS07778 | 0 | 0.000639743 | 4.84E-11 |
| SS00224 | 0.001130874 | 0.00221387 | 4.91E-11 |
| SS05555 | 0.001010691 | 0.003863524 | 4.99E-11 |
| SS05095 | 0 | 0.00046739 | 5.04E-11 |
| SS12505 | 0.002336979 | 7.89E-05 | 5.06E-11 |

| | | | |
|---|---|---|---|
| SS13744 | 0.002336979 | 7.89E-05 | 5.06E-11 |
| SS02317 | 0.140361021 | 0.177267222 | 5.28E-11 |
| SS03460 | 0.140361021 | 0.177267222 | 5.28E-11 |
| SS09330 | 0.004676562 | 0.000153672 | 5.36E-11 |
| SS03971 | 0.016674163 | 0.013231267 | 5.40E-11 |
| SS04122 | 0.016674163 | 0.013231267 | 5.40E-11 |
| SS08141 | 0.016674163 | 0.013231267 | 5.40E-11 |
| SS15159 | 0.016674163 | 0.013231267 | 5.40E-11 |
| SS01831 | 0 | 0.000291757 | 5.43E-11 |
| SS03008 | 0.026545444 | 0.018884454 | 5.52E-11 |
| SS00877 | 0.005046882 | 0.003229875 | 5.54E-11 |
| SS09795 | 0 | 0.000881746 | 5.59E-11 |
| SS15188 | 0 | 0.000881746 | 5.59E-11 |
| SS03398 | 0 | 0.000198657 | 5.62E-11 |
| SS01642 | 0 | 0.00014724 | 5.63E-11 |
| SS02466 | 0.000943545 | 0.003695916 | 5.81E-11 |
| SS14455 | 0 | 0.000263136 | 5.82E-11 |
| SS03260 | 0.005355873 | 0.002452617 | 5.93E-11 |
| SS10051 | 0.021212247 | 0.011328194 | 5.94E-11 |
| SS13954 | 0.001759785 | 0.003642683 | 6.08E-11 |
| SS11160 | 0.016174048 | 0.020708975 | 6.30E-11 |
| SS09375 | 0.012292254 | 0.00827478 | 6.41E-11 |
| SS00943 | 0.003160629 | 0.005432458 | 6.42E-11 |
| SS14013 | 0.003160629 | 0.005432458 | 6.42E-11 |
| SS06065 | 0.013292737 | 0.022101505 | 6.46E-11 |
| SS13555 | 0.013292737 | 0.022101505 | 6.46E-11 |
| SS14072 | 0 | 0.000401508 | 6.50E-11 |
| SS05886 | 0.002380565 | 0.005976584 | 6.67E-11 |
| SS03444 | 0 | 0.000592145 | 6.82E-11 |
| SS10289 | 0.027157471 | 0.009888667 | 6.87E-11 |
| SS10805 | 0.027157471 | 0.009888667 | 6.87E-11 |
| SS01176 | 0.002165881 | 0.001050019 | 7.08E-11 |
| SS00925 | 0.032114543 | 0.037802225 | 7.38E-11 |
| SS03964 | 0.032114543 | 0.037802225 | 7.38E-11 |
| SS14018 | 0.032114543 | 0.037802225 | 7.38E-11 |
| SS15490 | 0.032114543 | 0.037802225 | 7.38E-11 |
| SS09005 | 0.006502965 | 0.000184743 | 7.50E-11 |
| SS07005 | 0 | 0.000399522 | 7.67E-11 |
| SS05759 | 0 | 0.00035709 | 7.71E-11 |
| SS01443 | 4.07E-05 | 0.000292199 | 7.73E-11 |
| SS13004 | 4.07E-05 | 0.000292199 | 7.73E-11 |
| SS02177 | 0.018423625 | 0.025611866 | 7.76E-11 |
| SS06882 | 0.018423625 | 0.025611866 | 7.76E-11 |
| SS14221 | 0.018423625 | 0.025611866 | 7.76E-11 |
| SS15402 | 0.018423625 | 0.025611866 | 7.76E-11 |
| SS06735 | 0 | 0.000535898 | 7.80E-11 |
| SS04624 | 0 | 0.000686209 | 7.81E-11 |

| | | | |
|---|---|---|---|
| SS12999 | 0 | 0.001064449 | 7.98E-11 |
| SS07024 | 0.015516461 | 0.009032315 | 8.27E-11 |
| SS12980 | 0.015516461 | 0.009032315 | 8.27E-11 |
| SS04004 | 0.014707183 | 0.0041919 | 8.38E-11 |
| SS04494 | 0 | 0.000479359 | 8.39E-11 |
| SS07519 | 0.00044928 | 0.001937687 | 8.52E-11 |
| SS08509 | 0 | 0.00046277 | 8.57E-11 |
| SS00389 | 0.000667371 | 4.04E-05 | 8.79E-11 |
| SS03110 | 0.004804682 | 0.011974425 | 8.99E-11 |
| SS10202 | 0.004804682 | 0.011974425 | 8.99E-11 |
| SS07903 | 0.017150455 | 0.007179843 | 8.99E-11 |
| SS07619 | 0.002751769 | 0.000887954 | 9.27E-11 |
| SS08612 | 0.016446254 | 0.004811694 | 9.32E-11 |
| SS12507 | 0.002140922 | 7.09E-05 | 9.34E-11 |
| SS13747 | 0.002140922 | 7.09E-05 | 9.34E-11 |
| SS10204 | 2.76E-06 | 0.000162193 | 9.34E-11 |
| SS04236 | 0 | 0.000379456 | 9.46E-11 |
| SS01454 | 0 | 0.000378475 | 9.69E-11 |
| SS13621 | 0 | 0.000378475 | 9.69E-11 |
| SS09716 | 0 | 0.000384253 | 9.72E-11 |
| SS03235 | 0.001151864 | 0.002634932 | 9.81E-11 |
| SS03330 | 0.001151864 | 0.002634932 | 9.81E-11 |
| SS03244 | 0.001645372 | 0.004381965 | 9.83E-11 |
| SS03337 | 0.001645372 | 0.004381965 | 9.83E-11 |
| SS00740 | 0.000863749 | 0.000151445 | 9.97E-11 |
| SS05690 | 0.013061708 | 0.021671694 | 1.01E-10 |
| SS08716 | 0.013061708 | 0.021671694 | 1.01E-10 |
| SS15023 | 0.013061708 | 0.021671694 | 1.01E-10 |
| SS15030 | 0.013061708 | 0.021671694 | 1.01E-10 |
| SS03459 | 0.000702985 | 0.001861054 | 1.02E-10 |
| SS09282 | 0 | 0.000226961 | 1.05E-10 |
| SS03248 | 0.006813997 | 0.005135261 | 1.09E-10 |
| SS03342 | 0.006813997 | 0.005135261 | 1.09E-10 |
| SS04247 | 0 | 0.000345029 | 1.09E-10 |
| SS12960 | 0.012597999 | 0.018435328 | 1.12E-10 |
| SS15280 | 0.012597999 | 0.018435328 | 1.12E-10 |
| SS11743 | 0 | 0.000172898 | 1.15E-10 |
| SS14166 | 0 | 0.000172898 | 1.15E-10 |
| SS13884 | 0.000117178 | 0.001201345 | 1.19E-10 |
| SS03687 | 0.001579672 | 0.003958154 | 1.22E-10 |
| SS07544 | 0.001579672 | 0.003958154 | 1.22E-10 |
| SS11054 | 5.37E-06 | 0.000495777 | 1.24E-10 |
| SS13668 | 0.000166132 | 0.000716643 | 1.25E-10 |
| SS14984 | 0 | 0.004000561 | 1.28E-10 |
| SS01806 | 0 | 0.000318072 | 1.33E-10 |
| SS05049 | 0.005577645 | 2.87E-05 | 1.33E-10 |
| SS02611 | 0.008939331 | 0.004529111 | 1.35E-10 |

| | | | |
|---|---|---|---|
| SS03112 | 0.005110541 | 0.01141224 | 1.35E-10 |
| SS03690 | 0 | 0.000252328 | 1.36E-10 |
| SS07099 | 0 | 0.000252328 | 1.36E-10 |
| SS01742 | 0.000564015 | 0.001217118 | 1.36E-10 |
| SS06509 | 0.000564015 | 0.001217118 | 1.36E-10 |
| SS01184 | 0.005216501 | 0.000897627 | 1.37E-10 |
| SS01250 | 0.005216501 | 0.000897627 | 1.37E-10 |
| SS02754 | 0.005216501 | 0.000897627 | 1.37E-10 |
| SS03565 | 0.005216501 | 0.000897627 | 1.37E-10 |
| SS10172 | 0.005216501 | 0.000897627 | 1.37E-10 |
| SS04563 | 0.035526422 | 0.007778363 | 1.37E-10 |
| SS04665 | 0.035526422 | 0.007778363 | 1.37E-10 |
| SS12212 | 0.000612968 | 5.76E-05 | 1.38E-10 |
| SS06086 | 0 | 0.00031919 | 1.38E-10 |
| SS04481 | 0.000266964 | 2.20E-05 | 1.40E-10 |
| SS02798 | 0 | 0.000215701 | 1.40E-10 |
| SS08065 | 0 | 0.000294105 | 1.41E-10 |
| SS08861 | 0 | 0.000152817 | 1.42E-10 |
| SS10033 | 0 | 0.000178659 | 1.44E-10 |
| SS03351 | 0 | 0.000242318 | 1.47E-10 |
| SS13879 | 0 | 0.000242318 | 1.47E-10 |
| SS11429 | 0.008015744 | 0.013737079 | 1.52E-10 |
| SS13661 | 2.68E-06 | 0.000289752 | 1.54E-10 |
| SS01118 | 0 | 0.000168242 | 1.54E-10 |
| SS05705 | 0.005409036 | 0.010004041 | 1.56E-10 |
| SS09861 | 0.005409036 | 0.010004041 | 1.56E-10 |
| SS00533 | 0 | 0.00019394 | 1.61E-10 |
| SS15507 | 0.00017158 | 0.000797884 | 1.62E-10 |
| SS04383 | 0.024309561 | 0.007041858 | 1.63E-10 |
| SS04984 | 0.024309561 | 0.007041858 | 1.63E-10 |
| SS12982 | 0.024309561 | 0.007041858 | 1.63E-10 |
| SS07616 | 0.012685285 | 0.016551829 | 1.65E-10 |
| SS07641 | 0.012685285 | 0.016551829 | 1.65E-10 |
| SS01047 | 0.064703785 | 0.053593602 | 1.66E-10 |
| SS01191 | 0.064703785 | 0.053593602 | 1.66E-10 |
| SS01254 | 0.064703785 | 0.053593602 | 1.66E-10 |
| SS01305 | 0.064703785 | 0.053593602 | 1.66E-10 |
| SS14961 | 0.064703785 | 0.053593602 | 1.66E-10 |
| SS05033 | 0 | 0.000228039 | 1.67E-10 |
| SS05999 | 0.00020435 | 4.38E-05 | 1.67E-10 |
| SS04586 | 0.003705247 | 0.007921718 | 1.67E-10 |
| SS06200 | 0.001612602 | 0.002773429 | 1.67E-10 |
| SS05790 | 2.19E-05 | 0.001076762 | 1.68E-10 |
| SS05781 | 0.000425686 | 0.001943531 | 1.69E-10 |
| SS03777 | 0 | 0.000283978 | 1.71E-10 |
| SS06653 | 0.001195209 | 0.000384724 | 1.81E-10 |
| SS02531 | 0.006592305 | 0.001931452 | 1.82E-10 |

| | | | |
|---|---|---|---|
| SS00328 | 0.013293173 | 0.020812416 | 1.83E-10 |
| SS03955 | 0.013293173 | 0.020812416 | 1.83E-10 |
| SS02379 | 0.001509086 | 0.000977178 | 1.86E-10 |
| SS08856 | 0.007515617 | 0.005148285 | 1.86E-10 |
| SS08249 | 0.007653991 | 0.004717949 | 1.86E-10 |
| SS00117 | 0 | 0.000280348 | 1.86E-10 |
| SS07334 | 0.000122546 | 3.85E-05 | 1.88E-10 |
| SS07950 | 0 | 0.00046833 | 1.90E-10 |
| SS10634 | 0 | 0.000971211 | 1.92E-10 |
| SS12411 | 0 | 0.000267913 | 1.95E-10 |
| SS00946 | 0.003166157 | 0.005432458 | 2.01E-10 |
| SS14033 | 0.003166157 | 0.005432458 | 2.01E-10 |
| SS00215 | 0.018587417 | 0.01313309 | 2.07E-10 |
| SS00738 | 0.018587417 | 0.01313309 | 2.07E-10 |
| SS09836 | 0.018587417 | 0.01313309 | 2.07E-10 |
| SS01830 | 1.37E-05 | 0.00143271 | 2.09E-10 |
| SS09972 | 1.37E-05 | 0.00143271 | 2.09E-10 |
| SS00216 | 0.027484561 | 0.015387331 | 2.09E-10 |
| SS00752 | 0.027484561 | 0.015387331 | 2.09E-10 |
| SS09839 | 0.027484561 | 0.015387331 | 2.09E-10 |
| SS06717 | 0.012317213 | 0.00904541 | 2.11E-10 |
| SS13002 | 0.012317213 | 0.00904541 | 2.11E-10 |
| SS12735 | 2.68E-06 | 0.000209658 | 2.14E-10 |
| SS03776 | 0 | 0.000293002 | 2.15E-10 |
| SS12904 | 0 | 0.000293915 | 2.15E-10 |
| SS12917 | 0 | 0.000293915 | 2.15E-10 |
| SS05868 | 0.002160513 | 0.000836529 | 2.17E-10 |
| SS05878 | 0.002160513 | 0.000836529 | 2.17E-10 |
| SS05947 | 0.002160513 | 0.000836529 | 2.17E-10 |
| SS03707 | 1.10E-05 | 0.000430104 | 2.17E-10 |
| SS07009 | 0 | 0.000191688 | 2.19E-10 |
| SS05771 | 0 | 0.000335404 | 2.21E-10 |
| SS01804 | 0.007326052 | 4.63E-05 | 2.24E-10 |
| SS01481 | 0 | 0.000310282 | 2.27E-10 |
| SS00913 | 0.002481719 | 0.000820112 | 2.27E-10 |
| SS03589 | 0.002481719 | 0.000820112 | 2.27E-10 |
| SS13503 | 0.002481719 | 0.000820112 | 2.27E-10 |
| SS01504 | 0.000111649 | 3.41E-05 | 2.28E-10 |
| SS01686 | 0.000111649 | 3.41E-05 | 2.28E-10 |
| SS08080 | 0.000111649 | 3.41E-05 | 2.28E-10 |
| SS13570 | 0.000111649 | 3.41E-05 | 2.28E-10 |
| SS13260 | 0 | 0.000269498 | 2.28E-10 |
| SS13149 | 5.45E-05 | 0.00023471 | 2.34E-10 |
| SS13093 | 0 | 0.000303704 | 2.36E-10 |
| SS13941 | 0 | 0.000303704 | 2.36E-10 |
| SS08242 | 0.012434264 | 0.015616667 | 2.40E-10 |
| SS13868 | 0.012434264 | 0.015616667 | 2.40E-10 |

| | | | |
|---|---|---|---|
| SS10247 | 0.00012523 | 3.34E-05 | 2.44E-10 |
| SS06731 | 0 | 0.000140531 | 2.46E-10 |
| SS06268 | 0 | 0.000157823 | 2.55E-10 |
| SS01818 | 0.019306666 | 0.025725791 | 2.56E-10 |
| SS06160 | 0.019306666 | 0.025725791 | 2.56E-10 |
| SS06185 | 0.019306666 | 0.025725791 | 2.56E-10 |
| SS08217 | 0.019306666 | 0.025725791 | 2.56E-10 |
| SS12301 | 0 | 0.000603556 | 2.58E-10 |
| SS02271 | 0 | 0.000821471 | 2.58E-10 |
| SS08962 | 0 | 0.000266017 | 2.62E-10 |
| SS09483 | 0 | 0.000194687 | 2.64E-10 |
| SS05327 | 0.00442101 | 0.007574248 | 2.66E-10 |
| SS11140 | 0 | 0.000212011 | 2.68E-10 |
| SS05005 | 0 | 0.000202286 | 2.69E-10 |
| SS01282 | 0.02665395 | 0.003660774 | 2.75E-10 |
| SS08675 | 0.02665395 | 0.003660774 | 2.75E-10 |
| SS01641 | 0.001116972 | 0.000147072 | 2.76E-10 |
| SS10011 | 0.002149376 | 0.000535321 | 2.77E-10 |
| SS02382 | 0.01641525 | 0.020967298 | 2.77E-10 |
| SS03303 | 0.01641525 | 0.020967298 | 2.77E-10 |
| SS07787 | 0 | 0.000760932 | 2.81E-10 |
| SS02416 | 0.013295409 | 0.010536912 | 2.92E-10 |
| SS15414 | 0.013295409 | 0.010536912 | 2.92E-10 |
| SS05785 | 0 | 0.000176299 | 2.97E-10 |
| SS01366 | 0.069878683 | 0.057641759 | 2.99E-10 |
| SS02113 | 0.069878683 | 0.057641759 | 2.99E-10 |
| SS12868 | 0.069878683 | 0.057641759 | 2.99E-10 |
| SS02352 | 0.009778362 | 0.003315348 | 3.00E-10 |
| SS05465 | 0 | 0.000332197 | 3.02E-10 |
| SS02628 | 0 | 0.000147453 | 3.03E-10 |
| SS01712 | 0.005041801 | 0.010123895 | 3.10E-10 |
| SS01736 | 0.005041801 | 0.010123895 | 3.10E-10 |
| SS07189 | 0.005041801 | 0.010123895 | 3.10E-10 |
| SS01048 | 0.007199297 | 0.00389977 | 3.14E-10 |
| SS01308 | 0.007199297 | 0.00389977 | 3.14E-10 |
| SS06808 | 0.007199297 | 0.00389977 | 3.14E-10 |
| SS08933 | 0.007199297 | 0.00389977 | 3.14E-10 |
| SS12571 | 0.007199297 | 0.00389977 | 3.14E-10 |
| SS13843 | 0.007199297 | 0.00389977 | 3.14E-10 |
| SS15383 | 0.007199297 | 0.00389977 | 3.14E-10 |
| SS15431 | 0.007199297 | 0.00389977 | 3.14E-10 |
| SS11619 | 2.42E-05 | 0.000524429 | 3.17E-10 |
| SS02871 | 0 | 0.001197762 | 3.19E-10 |
| SS13616 | 0 | 0.001197762 | 3.19E-10 |
| SS02268 | 0.015015463 | 0.005765101 | 3.19E-10 |
| SS02573 | 0.015015463 | 0.005765101 | 3.19E-10 |
| SS08335 | 0.015015463 | 0.005765101 | 3.19E-10 |

| | | | |
|---|---|---|---|
| SS04926 | 0 | 0.00030675 | 3.20E-10 |
| SS07766 | 0 | 0.00093729 | 3.23E-10 |
| SS04401 | 0 | 0.000208607 | 3.28E-10 |
| SS11375 | 5.45E-06 | 1.09E-07 | 3.29E-10 |
| SS06095 | 0.015847554 | 0.021583104 | 3.31E-10 |
| SS06110 | 0.015847554 | 0.021583104 | 3.31E-10 |
| SS07685 | 0.015847554 | 0.021583104 | 3.31E-10 |
| SS07735 | 0.015847554 | 0.021583104 | 3.31E-10 |
| SS13582 | 0 | 0.000120236 | 3.44E-10 |
| SS03120 | 0.004917122 | 0.008247685 | 3.50E-10 |
| SS14477 | 0.004917122 | 0.008247685 | 3.50E-10 |
| SS15424 | 0.004917122 | 0.008247685 | 3.50E-10 |
| SS00780 | 0.029808052 | 0.036260649 | 3.51E-10 |
| SS07023 | 0.018110253 | 0.011052214 | 3.75E-10 |
| SS12979 | 0.018110253 | 0.011052214 | 3.75E-10 |
| SS02745 | 0.025001982 | 0.030842433 | 3.77E-10 |
| SS06039 | 0.025001982 | 0.030842433 | 3.77E-10 |
| SS06068 | 0.025001982 | 0.030842433 | 3.77E-10 |
| SS09626 | 0.025001982 | 0.030842433 | 3.77E-10 |
| SS15361 | 0.025001982 | 0.030842433 | 3.77E-10 |
| SS14563 | 0.000193052 | 0.000768936 | 3.78E-10 |
| SS12545 | 0.00551347 | 0.000921983 | 3.81E-10 |
| SS10266 | 0.000114333 | 1.86E-05 | 3.87E-10 |
| SS09026 | 0 | 0.000130426 | 3.92E-10 |
| SS00245 | 0 | 0.000298899 | 3.98E-10 |
| SS07568 | 0 | 0.000298899 | 3.98E-10 |
| SS13796 | 0.004250038 | 0.012746735 | 4.00E-10 |
| SS05766 | 0 | 0.000219483 | 4.02E-10 |
| SS05008 | 0 | 0.000272437 | 4.14E-10 |
| SS00223 | 0.000544745 | 0.0011989 | 4.16E-10 |
| SS04632 | 0 | 0.000415293 | 4.31E-10 |
| SS04265 | 0.012461471 | 0.016304831 | 4.33E-10 |
| SS00875 | 0.001676743 | 0.005642534 | 4.36E-10 |
| SS11279 | 0.001676743 | 0.005642534 | 4.36E-10 |
| SS11588 | 0.001676743 | 0.005642534 | 4.36E-10 |
| SS06085 | 0 | 0.000419171 | 4.42E-10 |
| SS11246 | 0 | 0.00081315 | 4.51E-10 |
| SS06573 | 0 | 0.000313623 | 4.53E-10 |
| SS05013 | 0 | 0.00022672 | 4.55E-10 |
| SS06956 | 0.011581412 | 0.008015674 | 4.58E-10 |
| SS07287 | 0.011581412 | 0.008015674 | 4.58E-10 |
| SS12346 | 0 | 0.000354923 | 4.70E-10 |
| SS04976 | 0.01497643 | 0.008702448 | 4.70E-10 |
| SS09034 | 0.01497643 | 0.008702448 | 4.70E-10 |
| SS04118 | 0.000362028 | 0.000828405 | 4.70E-10 |
| SS14022 | 2.68E-06 | 0.000159368 | 4.74E-10 |
| SS02343 | 0.000114172 | 0.000623966 | 4.84E-10 |

| | | | |
|---|---|---|---|
| SS05064 | 0.003395339 | 0.002006937 | 4.86E-10 |
| SS12400 | 0.002830487 | 0.00072911 | 4.99E-10 |
| SS06464 | 0.026182842 | 0.02189102 | 5.07E-10 |
| SS09521 | 0.026182842 | 0.02189102 | 5.07E-10 |
| SS09548 | 0.026182842 | 0.02189102 | 5.07E-10 |
| SS10457 | 0.026182842 | 0.02189102 | 5.07E-10 |
| SS10506 | 0.022531424 | 0.010977137 | 5.10E-10 |
| SS00366 | 0.01707052 | 0.011976116 | 5.14E-10 |
| SS01016 | 0.01707052 | 0.011976116 | 5.14E-10 |
| SS03580 | 0.01707052 | 0.011976116 | 5.14E-10 |
| SS06871 | 0.01707052 | 0.011976116 | 5.14E-10 |
| SS08221 | 0.01707052 | 0.011976116 | 5.14E-10 |
| SS14152 | 0.01707052 | 0.011976116 | 5.14E-10 |
| SS14210 | 0.01707052 | 0.011976116 | 5.14E-10 |
| SS15372 | 0.01707052 | 0.011976116 | 5.14E-10 |
| SS04524 | 0.000732107 | 0.001804197 | 5.15E-10 |
| SS08052 | 0 | 0.000135855 | 5.19E-10 |
| SS01752 | 0.010399669 | 0.000641491 | 5.20E-10 |
| SS05228 | 0.006254467 | 0.008742665 | 5.26E-10 |
| SS07907 | 0.006254467 | 0.008742665 | 5.26E-10 |
| SS14815 | 0.006254467 | 0.008742665 | 5.26E-10 |
| SS12456 | 0 | 0.000507498 | 5.28E-10 |
| SS09990 | 0.008020423 | 0.005113681 | 5.38E-10 |
| SS02668 | 0 | 0.000153133 | 5.38E-10 |
| SS03074 | 0 | 0.001263188 | 5.41E-10 |
| SS05894 | 0.000313636 | 0.001671917 | 5.48E-10 |
| SS13391 | 0 | 0.002833137 | 5.49E-10 |
| SS02143 | 0.000157999 | 0.000596974 | 5.54E-10 |
| SS15046 | 0.000451803 | 0.001079981 | 5.72E-10 |
| SS05272 | 0.009622966 | 0.00620048 | 5.76E-10 |
| SS15206 | 0 | 0.000529696 | 5.88E-10 |
| SS10082 | 0 | 0.000482044 | 5.97E-10 |
| SS09154 | 2.68E-06 | 0.001891303 | 6.06E-10 |
| SS12819 | 2.68E-06 | 0.000134711 | 6.06E-10 |
| SS01868 | 0 | 0.000279898 | 6.07E-10 |
| SS13854 | 0 | 0.000279898 | 6.07E-10 |
| SS03501 | 0.015111914 | 0.020759098 | 6.16E-10 |
| SS05241 | 0.015111914 | 0.020759098 | 6.16E-10 |
| SS07904 | 0.015111914 | 0.020759098 | 6.16E-10 |
| SS07915 | 0.015111914 | 0.020759098 | 6.16E-10 |
| SS14827 | 0.015111914 | 0.020759098 | 6.16E-10 |
| SS02924 | 0 | 0.000400412 | 6.21E-10 |
| SS13774 | 0 | 0.000367184 | 6.28E-10 |
| SS01083 | 0 | 0.000175124 | 6.72E-10 |
| SS00311 | 0.01044509 | 0.015011912 | 6.82E-10 |
| SS06951 | 0.01044509 | 0.015011912 | 6.82E-10 |
| SS09454 | 0 | 0.000224418 | 6.84E-10 |

| | | | |
|---|---|---|---|
| SS07572 | 0 | 0.000278234 | 6.90E-10 |
| SS01952 | 0 | 0.000148957 | 6.98E-10 |
| SS06172 | 0 | 0.000172487 | 6.99E-10 |
| SS04625 | 0 | 0.00079479 | 7.12E-10 |
| SS01187 | 0.019219105 | 0.014632801 | 7.16E-10 |
| SS01251 | 0.019219105 | 0.014632801 | 7.16E-10 |
| SS02755 | 0.019219105 | 0.014632801 | 7.16E-10 |
| SS03566 | 0.019219105 | 0.014632801 | 7.16E-10 |
| SS00413 | 0.011635333 | 0.007969006 | 7.31E-10 |
| SS13258 | 0 | 0.000125726 | 7.34E-10 |
| SS03847 | 0.000117097 | 4.47E-05 | 7.39E-10 |
| SS02037 | 0 | 0.000311413 | 7.39E-10 |
| SS04203 | 0.001084844 | 0.002997711 | 7.44E-10 |
| SS13372 | 0 | 0.000144202 | 7.56E-10 |
| SS05802 | 0.008521066 | 0.00257804 | 7.81E-10 |
| SS06538 | 0.006734521 | 0.003820091 | 7.82E-10 |
| SS07559 | 0.006734521 | 0.003820091 | 7.82E-10 |
| SS11093 | 0 | 0.000351819 | 7.97E-10 |
| SS13604 | 0.007977399 | 0.012468077 | 8.05E-10 |
| SS02789 | 0 | 0.000586373 | 8.15E-10 |
| SS13701 | 0.011322305 | 0.01507461 | 8.18E-10 |
| SS00968 | 0.011727035 | 0.009375805 | 8.19E-10 |
| SS06762 | 0.011727035 | 0.009375805 | 8.19E-10 |
| SS08434 | 0.011727035 | 0.009375805 | 8.19E-10 |
| SS08471 | 0.011727035 | 0.009375805 | 8.19E-10 |
| SS10826 | 0.015920939 | 0.008827772 | 8.25E-10 |
| SS14228 | 0.015920939 | 0.008827772 | 8.25E-10 |
| SS09520 | 0.001065334 | 0.000394823 | 8.33E-10 |
| SS02264 | 0.000560929 | 0.001584227 | 8.85E-10 |
| SS01856 | 0.005821933 | 0.004087416 | 8.87E-10 |
| SS06312 | 0.005821933 | 0.004087416 | 8.87E-10 |
| SS06413 | 0.005821933 | 0.004087416 | 8.87E-10 |
| SS13609 | 0.005821933 | 0.004087416 | 8.87E-10 |
| SS10459 | 0.015876722 | 0.020676835 | 8.92E-10 |
| SS02071 | 0.050406172 | 0.061871775 | 8.92E-10 |
| SS02607 | 0.050406172 | 0.061871775 | 8.92E-10 |
| SS08820 | 0.050406172 | 0.061871775 | 8.92E-10 |
| SS13797 | 0.050406172 | 0.061871775 | 8.92E-10 |
| SS13819 | 0.050406172 | 0.061871775 | 8.92E-10 |
| SS14895 | 0.050406172 | 0.061871775 | 8.92E-10 |
| SS04731 | 0 | 0.000231702 | 8.97E-10 |
| SS10079 | 0 | 0.000231702 | 8.97E-10 |
| SS13979 | 0.012352621 | 0.021975785 | 9.14E-10 |
| SS15443 | 0.012352621 | 0.021975785 | 9.14E-10 |
| SS13383 | 0.001492179 | 0.003391854 | 9.29E-10 |
| SS09807 | 0 | 0.000310791 | 9.34E-10 |
| SS13574 | 0 | 0.000310791 | 9.34E-10 |

| | | | |
|---|---|---|---|
| SS00229 | 0.000386585 | 9.63E-05 | 9.55E-10 |
| SS03954 | 0.009867702 | 0.012616573 | 9.69E-10 |
| SS15483 | 0.009867702 | 0.012616573 | 9.69E-10 |
| SS07518 | 0.001549185 | 0.000780317 | 9.73E-10 |
| SS02346 | 0 | 0.000148651 | 9.96E-10 |
| SS11118 | 0 | 0.000148651 | 9.96E-10 |
| SS03998 | 0.017053304 | 0.001076287 | 1.00E-09 |
| SS01728 | 0.014364403 | 0.026125609 | 1.04E-09 |
| SS04198 | 0.014364403 | 0.026125609 | 1.04E-09 |
| SS06917 | 0.014364403 | 0.026125609 | 1.04E-09 |
| SS07182 | 0.014364403 | 0.026125609 | 1.04E-09 |
| SS15080 | 0.014364403 | 0.026125609 | 1.04E-09 |
| SS13942 | 0 | 0.000593786 | 1.05E-09 |
| SS09940 | 0.002888893 | 0.000757318 | 1.06E-09 |
| SS12002 | 0.002888893 | 0.000757318 | 1.06E-09 |
| SS13358 | 0 | 0.000599308 | 1.06E-09 |
| SS00889 | 0.007491461 | 0.009463052 | 1.06E-09 |
| SS02549 | 0.007491461 | 0.009463052 | 1.06E-09 |
| SS09294 | 0.007491461 | 0.009463052 | 1.06E-09 |
| SS05088 | 2.76E-06 | 0.000289331 | 1.08E-09 |
| SS05294 | 0.000310631 | 9.10E-05 | 1.10E-09 |
| SS08179 | 0.011353756 | 0.008284372 | 1.12E-09 |
| SS10153 | 0.011353756 | 0.008284372 | 1.12E-09 |
| SS11687 | 5.45E-06 | 1.46E-06 | 1.14E-09 |
| SS02694 | 0.015876722 | 0.020644568 | 1.19E-09 |
| SS09552 | 0.015876722 | 0.020644568 | 1.19E-09 |
| SS00265 | 0.000166132 | 4.44E-07 | 1.19E-09 |
| SS01545 | 0.000166132 | 4.44E-07 | 1.19E-09 |
| SS01067 | 0.009506271 | 0.001205541 | 1.20E-09 |
| SS01165 | 0 | 0.000195474 | 1.22E-09 |
| SS01006 | 0.010579863 | 0.013639115 | 1.23E-09 |
| SS02472 | 0.010579863 | 0.013639115 | 1.23E-09 |
| SS13528 | 0.010579863 | 0.013639115 | 1.23E-09 |
| SS03711 | 8.16E-05 | 0.000480685 | 1.23E-09 |
| SS05801 | 0.009522902 | 0.001989986 | 1.24E-09 |
| SS10861 | 0.00394489 | 0.001217317 | 1.24E-09 |
| SS08702 | 0 | 0.000196025 | 1.27E-09 |
| SS03184 | 0 | 0.000505988 | 1.28E-09 |
| SS02979 | 0.046060256 | 0.000951643 | 1.32E-09 |
| SS08560 | 0.046060256 | 0.000951643 | 1.32E-09 |
| SS14998 | 0 | 0.000252647 | 1.35E-09 |
| SS15006 | 0 | 0.000252647 | 1.35E-09 |
| SS11064 | 0.009786701 | 0.000826878 | 1.35E-09 |
| SS06003 | 0.00023143 | 0.000104194 | 1.35E-09 |
| SS00657 | 0.016991675 | 0.027243413 | 1.38E-09 |
| SS02029 | 0.016991675 | 0.027243413 | 1.38E-09 |
| SS06937 | 0.016991675 | 0.027243413 | 1.38E-09 |

| | | | |
|---|---|---|---|
| SS09938 | 0.016991675 | 0.027243413 | 1.38E-09 |
| SS05661 | 0.032687447 | 0.002489611 | 1.38E-09 |
| SS13094 | 0.032687447 | 0.002489611 | 1.38E-09 |
| SS01667 | 2.68E-06 | 0.0001565 | 1.39E-09 |
| SS13516 | 0 | 0.000690802 | 1.40E-09 |
| SS02724 | 0 | 0.000666685 | 1.42E-09 |
| SS06840 | 0.054043253 | 0.107517285 | 1.42E-09 |
| SS13635 | 0.054043253 | 0.107517285 | 1.42E-09 |
| SS01448 | 2.76E-06 | 0.000167944 | 1.43E-09 |
| SS13049 | 2.76E-06 | 0.000167944 | 1.43E-09 |
| SS00415 | 0.010923619 | 0.008775002 | 1.46E-09 |
| SS00429 | 0.010923619 | 0.008775002 | 1.46E-09 |
| SS07951 | 0 | 0.000183211 | 1.47E-09 |
| SS15209 | 0.000541981 | 0.001535845 | 1.49E-09 |
| SS01001 | 0 | 0.000577654 | 1.50E-09 |
| SS02511 | 0.000221176 | 0.001073755 | 1.50E-09 |
| SS15525 | 0.000980364 | 0.000601072 | 1.52E-09 |
| SS06624 | 2.18E-05 | 0.000582816 | 1.52E-09 |
| SS00368 | 0.02654604 | 0.032630003 | 1.54E-09 |
| SS01018 | 0.02654604 | 0.032630003 | 1.54E-09 |
| SS03582 | 0.02654604 | 0.032630003 | 1.54E-09 |
| SS06873 | 0.02654604 | 0.032630003 | 1.54E-09 |
| SS08223 | 0.02654604 | 0.032630003 | 1.54E-09 |
| SS14154 | 0.02654604 | 0.032630003 | 1.54E-09 |
| SS14211 | 0.02654604 | 0.032630003 | 1.54E-09 |
| SS15374 | 0.02654604 | 0.032630003 | 1.54E-09 |
| SS10710 | 0.000106201 | 0.000333295 | 1.57E-09 |
| SS14456 | 2.18E-05 | 0.000707433 | 1.58E-09 |
| SS01190 | 0.007326534 | 0.005402279 | 1.59E-09 |
| SS00414 | 0.019333747 | 0.005930111 | 1.61E-09 |
| SS00428 | 0.019333747 | 0.005930111 | 1.61E-09 |
| SS05689 | 0.004325189 | 0.008111429 | 1.61E-09 |
| SS15022 | 0.004325189 | 0.008111429 | 1.61E-09 |
| SS15029 | 0.004325189 | 0.008111429 | 1.61E-09 |
| SS13392 | 0 | 0.000771683 | 1.63E-09 |
| SS00421 | 0.007977365 | 0.002504881 | 1.63E-09 |
| SS00432 | 0.007977365 | 0.002504881 | 1.63E-09 |
| SS00167 | 0.043881919 | 0.00276016 | 1.63E-09 |
| SS01671 | 0.043881919 | 0.00276016 | 1.63E-09 |
| SS03063 | 0.043881919 | 0.00276016 | 1.63E-09 |
| SS05362 | 0.043881919 | 0.00276016 | 1.63E-09 |
| SS05370 | 0.043881919 | 0.00276016 | 1.63E-09 |
| SS10789 | 0.043881919 | 0.00276016 | 1.63E-09 |
| SS14461 | 0.043881919 | 0.00276016 | 1.63E-09 |
| SS15336 | 0 | 0.000242554 | 1.66E-09 |
| SS00042 | 0.001119335 | 0.002323479 | 1.70E-09 |
| SS03056 | 3.81E-05 | 0.000715213 | 1.72E-09 |

| | | | |
|---|---|---|---|
| SS05494 | 3.81E-05 | 0.000715213 | 1.72E-09 |
| SS09515 | 3.81E-05 | 0.000715213 | 1.72E-09 |
| SS03325 | 2.68E-06 | 0.000231169 | 1.74E-09 |
| SS02750 | 0.012666933 | 0.007058596 | 1.75E-09 |
| SS03923 | 0.012666933 | 0.007058596 | 1.75E-09 |
| SS05921 | 0.012666933 | 0.007058596 | 1.75E-09 |
| SS11823 | 0.012086058 | 0.009222425 | 1.75E-09 |
| SS11831 | 0.012086058 | 0.009222425 | 1.75E-09 |
| SS13741 | 0.000359103 | 0.0009983 | 1.75E-09 |
| SS11816 | 0.002645407 | 0.001260738 | 1.76E-09 |
| SS11282 | 0.013706001 | 0.019549039 | 1.76E-09 |
| SS10320 | 0.011758807 | 0.000893123 | 1.77E-09 |
| SS01283 | 0.026642893 | 0.003640678 | 1.77E-09 |
| SS05497 | 0.026642893 | 0.003640678 | 1.77E-09 |
| SS05871 | 0.026642893 | 0.003640678 | 1.77E-09 |
| SS14905 | 0 | 0.00015954 | 1.78E-09 |
| SS11129 | 0.013811652 | 0.02878876 | 1.78E-09 |
| SS14741 | 0.013811652 | 0.02878876 | 1.78E-09 |
| SS09227 | 0 | 9.53E-05 | 1.80E-09 |
| SS05884 | 0.003030387 | 0.00640777 | 1.80E-09 |
| SS15388 | 0.003030387 | 0.00640777 | 1.80E-09 |
| SS03373 | 0.000530763 | 0.001217922 | 1.82E-09 |
| SS04870 | 0.000530763 | 0.001217922 | 1.82E-09 |
| SS07262 | 0.003230733 | 0.000465879 | 1.83E-09 |
| SS03555 | 0.000190287 | 0.000944867 | 1.84E-09 |
| SS08897 | 0.012065756 | 0.001763584 | 1.85E-09 |
| SS05277 | 0.00089848 | 0.000120313 | 1.87E-09 |
| SS00170 | 0.001351407 | 0.002553078 | 1.88E-09 |
| SS00195 | 0.001351407 | 0.002553078 | 1.88E-09 |
| SS13180 | 0 | 0.00027605 | 1.90E-09 |
| SS13626 | 0 | 0.000199481 | 1.92E-09 |
| SS09776 | 0.000310791 | 0.001431883 | 1.92E-09 |
| SS10589 | 0.000310791 | 0.001431883 | 1.92E-09 |
| SS15168 | 0.000310791 | 0.001431883 | 1.92E-09 |
| SS13594 | 0.000386906 | 3.00E-05 | 1.94E-09 |
| SS02719 | 0 | 0.00054485 | 2.06E-09 |
| SS04035 | 0.001707861 | 0.006008 | 2.09E-09 |
| SS06556 | 0.001707861 | 0.006008 | 2.09E-09 |
| SS13161 | 0 | 0.000295319 | 2.10E-09 |
| SS02633 | 0.048305188 | 0.056343676 | 2.11E-09 |
| SS04604 | 0.048305188 | 0.056343676 | 2.11E-09 |
| SS08454 | 0.048305188 | 0.056343676 | 2.11E-09 |
| SS14081 | 0 | 0.000581614 | 2.12E-09 |
| SS06125 | 0.00025062 | 0.00066223 | 2.16E-09 |
| SS14189 | 0.027946504 | 0.018757377 | 2.28E-09 |
| SS14253 | 0.027946504 | 0.018757377 | 2.28E-09 |
| SS04424 | 0.000234114 | 4.82E-05 | 2.29E-09 |

| | | | |
|---|---|---|---|
| SS06300 | 0.000234114 | 4.82E-05 | 2.29E-09 |
| SS09442 | 0.009761295 | 0.005415582 | 2.30E-09 |
| SS01775 | 0.043846626 | 0.002501569 | 2.32E-09 |
| SS05117 | 8.72E-05 | 0.00141874 | 2.34E-09 |
| SS08750 | 8.72E-05 | 0.00141874 | 2.34E-09 |
| SS11068 | 8.72E-05 | 0.00141874 | 2.34E-09 |
| SS03949 | 0.011227242 | 0.014071287 | 2.35E-09 |
| SS04638 | 0.011227242 | 0.014071287 | 2.35E-09 |
| SS11189 | 0.011227242 | 0.014071287 | 2.35E-09 |
| SS02697 | 0.00700901 | 0.002145645 | 2.39E-09 |
| SS03572 | 0.00700901 | 0.002145645 | 2.39E-09 |
| SS10183 | 0.00700901 | 0.002145645 | 2.39E-09 |
| SS09823 | 0 | 0.000141796 | 2.40E-09 |
| SS11600 | 0 | 0.000221523 | 2.48E-09 |
| SS06861 | 0.000929128 | 0.002510411 | 2.48E-09 |
| SS05765 | 0 | 0.000226194 | 2.48E-09 |
| SS05325 | 0.010736165 | 0.019534841 | 2.54E-09 |
| SS06617 | 0 | 0.000585236 | 2.60E-09 |
| SS06980 | 0 | 0.000288833 | 2.62E-09 |
| SS03396 | 0 | 0.000241289 | 2.65E-09 |
| SS05089 | 0 | 0.000485139 | 2.66E-09 |
| SS05991 | 0.040372178 | 0.049492735 | 2.67E-09 |
| SS06032 | 0.040372178 | 0.049492735 | 2.67E-09 |
| SS08213 | 0.040372178 | 0.049492735 | 2.67E-09 |
| SS12995 | 0.040372178 | 0.049492735 | 2.67E-09 |
| SS09651 | 0 | 0.000183992 | 2.69E-09 |
| SS00109 | 0.029511552 | 0.012319909 | 2.71E-09 |
| SS07824 | 0.000163448 | 0.000773885 | 2.72E-09 |
| SS00746 | 0.000602072 | 7.48E-05 | 2.78E-09 |
| SS00407 | 0 | 9.54E-05 | 2.78E-09 |
| SS12265 | 0.000748934 | 8.87E-06 | 2.80E-09 |
| SS09777 | 0.000687959 | 0.002637669 | 2.84E-09 |
| SS10590 | 0.000687959 | 0.002637669 | 2.84E-09 |
| SS15169 | 0.000687959 | 0.002637669 | 2.84E-09 |
| SS15155 | 0 | 0.000145483 | 2.88E-09 |
| SS10718 | 0.001080956 | 0.000592313 | 2.89E-09 |
| SS05743 | 0.007392314 | 0.010778661 | 2.93E-09 |
| SS13708 | 1.10E-05 | 0.00034931 | 2.94E-09 |
| SS02953 | 0 | 0.000251592 | 3.15E-09 |
| SS01078 | 0.013000596 | 0.019974255 | 3.16E-09 |
| SS08904 | 0 | 0.00015063 | 3.19E-09 |
| SS01279 | 7.61E-05 | 0.000649301 | 3.27E-09 |
| SS13828 | 0.003517483 | 0.005419616 | 3.30E-09 |
| SS02269 | 0.005650675 | 0.007776617 | 3.32E-09 |
| SS02575 | 0.005650675 | 0.007776617 | 3.32E-09 |
| SS11188 | 0.005650675 | 0.007776617 | 3.32E-09 |
| SS06681 | 0 | 0.000318319 | 3.33E-09 |

| | | | |
|---|---|---|---|
| SS01364 | 0.016261862 | 0.021404463 | 3.39E-09 |
| SS02296 | 0.016261862 | 0.021404463 | 3.39E-09 |
| SS02519 | 0.016261862 | 0.021404463 | 3.39E-09 |
| SS03774 | 0.016261862 | 0.021404463 | 3.39E-09 |
| SS06022 | 0.016261862 | 0.021404463 | 3.39E-09 |
| SS03630 | 0.018631221 | 0.009708045 | 3.43E-09 |
| SS04557 | 0.018631221 | 0.009708045 | 3.43E-09 |
| SS04831 | 0.018631221 | 0.009708045 | 3.43E-09 |
| SS09781 | 0.000190368 | 0.000860303 | 3.45E-09 |
| SS15173 | 0.000190368 | 0.000860303 | 3.45E-09 |
| SS12043 | 0.004617755 | 0.002365061 | 3.47E-09 |
| SS06475 | 0.009589118 | 0.012306372 | 3.47E-09 |
| SS03072 | 0.001302293 | 0.000524264 | 3.63E-09 |
| SS15438 | 0.001302293 | 0.000524264 | 3.63E-09 |
| SS14082 | 0.042118383 | 0.049749263 | 3.64E-09 |
| SS14035 | 0 | 0.000323859 | 3.67E-09 |
| SS10214 | 0.007341353 | 0.010278394 | 3.68E-09 |
| SS10328 | 0.007341353 | 0.010278394 | 3.68E-09 |
| SS11376 | 5.45E-06 | 1.58E-06 | 3.68E-09 |
| SS01696 | 0.002895304 | 0.005295314 | 3.70E-09 |
| SS05769 | 0 | 0.000345571 | 3.74E-09 |
| SS14748 | 2.72E-05 | 0.000175949 | 3.79E-09 |
| SS13946 | 0.029916581 | 0.000702901 | 3.82E-09 |
| SS13947 | 0.029916581 | 0.000702901 | 3.82E-09 |
| SS10045 | 0 | 9.13E-05 | 3.82E-09 |
| SS14008 | 0.024911518 | 0.018820924 | 3.84E-09 |
| SS14136 | 0.024911518 | 0.018820924 | 3.84E-09 |
| SS00038 | 0.017931597 | 0.001476283 | 3.85E-09 |
| SS10205 | 0 | 0.000267198 | 4.10E-09 |
| SS10276 | 0 | 0.000267198 | 4.10E-09 |
| SS01058 | 0.034465883 | 0.023538886 | 4.13E-09 |
| SS01319 | 0.034465883 | 0.023538886 | 4.13E-09 |
| SS03962 | 0.034465883 | 0.023538886 | 4.13E-09 |
| SS05026 | 0 | 0.000343111 | 4.15E-09 |
| SS07788 | 2.68E-06 | 0.000358676 | 4.16E-09 |
| SS02873 | 0 | 0.001595381 | 4.17E-09 |
| SS03664 | 2.68E-06 | 0.000292093 | 4.38E-09 |
| SS07073 | 2.68E-06 | 0.000292093 | 4.38E-09 |
| SS09583 | 0.021517051 | 0.004198603 | 4.40E-09 |
| SS06452 | 0.002800723 | 0.000945619 | 4.42E-09 |
| SS05508 | 5.18E-05 | 0.000701338 | 4.42E-09 |
| SS10818 | 5.18E-05 | 0.000701338 | 4.42E-09 |
| SS02540 | 0.014012009 | 0.006369661 | 4.67E-09 |
| SS06934 | 0 | 0.00074024 | 4.71E-09 |
| SS03638 | 0.005472408 | 0.000713098 | 4.77E-09 |
| SS04836 | 0.005472408 | 0.000713098 | 4.77E-09 |
| SS01561 | 0.000152792 | 0.000662346 | 4.79E-09 |

| | | | |
|---|---|---|---|
| SS13403 | 0.000152792 | 0.000662346 | 4.79E-09 |
| SS09135 | 0.000258431 | 0.001247006 | 4.82E-09 |
| SS10656 | 0 | 0.000128583 | 4.82E-09 |
| SS08964 | 1.63E-05 | 0.000353011 | 4.87E-09 |
| SS04712 | 0.044896681 | 0.030226162 | 4.91E-09 |
| SS11048 | 0.044896681 | 0.030226162 | 4.91E-09 |
| SS04015 | 0.001031761 | 0.001956954 | 4.96E-09 |
| SS10158 | 0 | 7.91E-05 | 4.98E-09 |
| SS04442 | 0 | 0.000148098 | 4.98E-09 |
| SS08360 | 8.13E-06 | 0.000677078 | 5.01E-09 |
| SS05354 | 0 | 0.000127241 | 5.05E-09 |
| SS02796 | 0 | 7.24E-05 | 5.08E-09 |
| SS02661 | 0.012598607 | 0.021445375 | 5.47E-09 |
| SS11441 | 0.012598607 | 0.021445375 | 5.47E-09 |
| SS03190 | 0 | 0.000338927 | 5.48E-09 |
| SS13856 | 0.015966051 | 0.00219304 | 5.55E-09 |
| SS05458 | 4.37E-05 | 0.000518832 | 5.57E-09 |
| SS02451 | 0 | 0.000531226 | 5.58E-09 |
| SS03294 | 0 | 0.000531226 | 5.58E-09 |
| SS01415 | 0 | 0.000143245 | 5.65E-09 |
| SS08970 | 0.001033046 | 0.002962444 | 5.65E-09 |
| SS15422 | 0.001033046 | 0.002962444 | 5.65E-09 |
| SS15440 | 0.001033046 | 0.002962444 | 5.65E-09 |
| SS04300 | 0.001322285 | 0.003263156 | 5.66E-09 |
| SS00188 | 0.091472732 | 0.080366195 | 5.69E-09 |
| SS01782 | 0.091472732 | 0.080366195 | 5.69E-09 |
| SS05343 | 0.091472732 | 0.080366195 | 5.69E-09 |
| SS06907 | 0.091472732 | 0.080366195 | 5.69E-09 |
| SS08720 | 0.091472732 | 0.080366195 | 5.69E-09 |
| SS09904 | 0.091472732 | 0.080366195 | 5.69E-09 |
| SS15071 | 0.091472732 | 0.080366195 | 5.69E-09 |
| SS00906 | 0.000120825 | 0.003657016 | 5.71E-09 |
| SS09963 | 0.000120825 | 0.003657016 | 5.71E-09 |
| SS14030 | 0 | 0.000330146 | 5.85E-09 |
| SS02391 | 0.00025663 | 0.00110145 | 6.05E-09 |
| SS15198 | 0.000585326 | 0.000229736 | 6.26E-09 |
| SS01440 | 0 | 0.000268084 | 6.39E-09 |
| SS13860 | 0.038102104 | 0.04841858 | 6.40E-09 |
| SS00184 | 0 | 0.000323682 | 6.45E-09 |
| SS00602 | 0 | 0.000323682 | 6.45E-09 |
| SS10553 | 0.001836222 | 0.000972118 | 6.56E-09 |
| SS04653 | 0.00396424 | 0.002252764 | 6.72E-09 |
| SS03263 | 0.021299889 | 0.0180234 | 6.74E-09 |
| SS00950 | 0.046904586 | 0.073425753 | 6.87E-09 |
| SS05897 | 0.046904586 | 0.073425753 | 6.87E-09 |
| SS09878 | 0.046904586 | 0.073425753 | 6.87E-09 |
| SS15066 | 0.046904586 | 0.073425753 | 6.87E-09 |

| | | | |
|---|---|---|---|
| SS00144 | 0.000152471 | 7.64E-06 | 7.01E-09 |
| SS07719 | 0.005504696 | 0.003504017 | 7.10E-09 |
| SS08369 | 0 | 0.000283082 | 7.12E-09 |
| SS03650 | 0.020530287 | 0.017006272 | 7.16E-09 |
| SS05619 | 0.020530287 | 0.017006272 | 7.16E-09 |
| SS05718 | 0.020530287 | 0.017006272 | 7.16E-09 |
| SS07017 | 0.020530287 | 0.017006272 | 7.16E-09 |
| SS07893 | 0.020530287 | 0.017006272 | 7.16E-09 |
| SS12935 | 0.020530287 | 0.017006272 | 7.16E-09 |
| SS03552 | 0 | 0.000310579 | 7.16E-09 |
| SS04887 | 0.002641599 | 3.64E-05 | 7.36E-09 |
| SS13499 | 0.002641599 | 3.64E-05 | 7.36E-09 |
| SS07780 | 0 | 0.000490272 | 7.44E-09 |
| SS00498 | 0.030598014 | 0.025727264 | 7.45E-09 |
| SS01002 | 0.030598014 | 0.025727264 | 7.45E-09 |
| SS02470 | 0.030598014 | 0.025727264 | 7.45E-09 |
| SS01732 | 0.014968034 | 0.021798861 | 7.50E-09 |
| SS06904 | 0.014968034 | 0.021798861 | 7.50E-09 |
| SS06920 | 0.014968034 | 0.021798861 | 7.50E-09 |
| SS01817 | 0.014980881 | 0.012612459 | 7.53E-09 |
| SS09045 | 0.014980881 | 0.012612459 | 7.53E-09 |
| SS13122 | 0.014980881 | 0.012612459 | 7.53E-09 |
| SS12236 | 0.000874003 | 0.000491397 | 7.64E-09 |
| SS09356 | 1.36E-05 | 0.000661894 | 7.70E-09 |
| SS08410 | 5.19E-05 | 0.000398595 | 7.90E-09 |
| SS09971 | 2.44E-05 | 0.000841918 | 8.02E-09 |
| SS11261 | 0.007855221 | 0.012834141 | 8.02E-09 |
| SS13487 | 0.005431907 | 0.001033537 | 8.08E-09 |
| SS05955 | 0.024397915 | 0.010902284 | 8.21E-09 |
| SS06503 | 0.000266723 | 0.001358627 | 8.37E-09 |
| SS07177 | 0.000266723 | 0.001358627 | 8.37E-09 |
| SS00189 | 9.04E-05 | 0.000880074 | 8.59E-09 |
| SS02675 | 0.02138143 | 0.033576416 | 8.74E-09 |
| SS13977 | 0.02138143 | 0.033576416 | 8.74E-09 |
| SS15441 | 0.02138143 | 0.033576416 | 8.74E-09 |
| SS00739 | 0.008180831 | 0.012891418 | 8.85E-09 |
| SS00790 | 0.008180831 | 0.012891418 | 8.85E-09 |
| SS03557 | 0.002272885 | 0.00421603 | 8.88E-09 |
| SS05398 | 0.000816435 | 0.001682342 | 8.91E-09 |
| SS05464 | 0.000816435 | 0.001682342 | 8.91E-09 |
| SS06577 | 0 | 0.000181371 | 9.24E-09 |
| SS11715 | 0.015952 | 0.011233522 | 9.27E-09 |
| SS11761 | 0.015952 | 0.011233522 | 9.27E-09 |
| SS14487 | 0.015952 | 0.011233522 | 9.27E-09 |
| SS09132 | 0.001951633 | 0.003786987 | 9.49E-09 |
| SS07781 | 1.38E-05 | 0.000339978 | 9.55E-09 |
| SS07001 | 0 | 0.000463576 | 9.55E-09 |

| | | | |
|---|---|---|---|
| SS08710 | 0 | 0.000218275 | 9.81E-09 |
| SS13705 | 0.030948583 | 0.039905669 | 9.86E-09 |
| SS11833 | 0 | 0.000159711 | 9.95E-09 |
| SS01915 | 8.21E-06 | 0.000193446 | 1.03E-08 |
| SS08263 | 0.006048798 | 0.007898346 | 1.03E-08 |
| SS10957 | 0.006048798 | 0.007898346 | 1.03E-08 |
| SS03787 | 0.01197153 | 0.014840891 | 1.05E-08 |
| SS06260 | 0.01197153 | 0.014840891 | 1.05E-08 |
| SS12634 | 0.01197153 | 0.014840891 | 1.05E-08 |
| SS13838 | 0.01197153 | 0.014840891 | 1.05E-08 |
| SS13919 | 0.01197153 | 0.014840891 | 1.05E-08 |
| SS04446 | 0 | 0.000334887 | 1.05E-08 |
| SS09503 | 1.63E-05 | 0.000275266 | 1.05E-08 |
| SS14040 | 0.010990443 | 0.016970534 | 1.06E-08 |
| SS11622 | 0 | 0.000104087 | 1.08E-08 |
| SS08553 | 0.010151011 | 0.000163563 | 1.08E-08 |
| SS03044 | 0.011308277 | 0.002405542 | 1.11E-08 |
| SS10850 | 0.011308277 | 0.002405542 | 1.11E-08 |
| SS02365 | 0.004446496 | 0.002015582 | 1.11E-08 |
| SS10221 | 0.004446496 | 0.002015582 | 1.11E-08 |
| SS10943 | 5.45E-06 | 0.00082798 | 1.16E-08 |
| SS05893 | 0.00397189 | 0.008529731 | 1.17E-08 |
| SS08603 | 0 | 0.000561511 | 1.19E-08 |
| SS01430 | 0 | 0.000107316 | 1.20E-08 |
| SS12947 | 0 | 0.000107316 | 1.20E-08 |
| SS02509 | 0 | 0.0001023 | 1.22E-08 |
| SS10741 | 0.008412503 | 0.004591186 | 1.25E-08 |
| SS00060 | 0.007741197 | 0.004103096 | 1.27E-08 |
| SS00813 | 0.007741197 | 0.004103096 | 1.27E-08 |
| SS02836 | 0.007741197 | 0.004103096 | 1.27E-08 |
| SS11204 | 0.025702226 | 0.017406431 | 1.29E-08 |
| SS11207 | 0.025702226 | 0.017406431 | 1.29E-08 |
| SS04293 | 0.014152276 | 0.006831425 | 1.31E-08 |
| SS03311 | 0 | 0.000316718 | 1.32E-08 |
| SS08182 | 0.003903472 | 0.006584729 | 1.33E-08 |
| SS10155 | 0.003903472 | 0.006584729 | 1.33E-08 |
| SS10288 | 0.003903472 | 0.006584729 | 1.33E-08 |
| SS07203 | 0.015583067 | 0.008111451 | 1.34E-08 |
| SS07226 | 0.015583067 | 0.008111451 | 1.34E-08 |
| SS04136 | 0 | 0.000276101 | 1.35E-08 |
| SS07275 | 0.029283907 | 0.022889814 | 1.36E-08 |
| SS08920 | 0.029283907 | 0.022889814 | 1.36E-08 |
| SS13970 | 6.80E-05 | 0.00036715 | 1.36E-08 |
| SS12458 | 0 | 0.000319244 | 1.37E-08 |
| SS05753 | 0 | 0.000352106 | 1.38E-08 |
| SS09071 | 0.002759981 | 0.001399664 | 1.38E-08 |
| SS14857 | 0.002759981 | 0.001399664 | 1.38E-08 |

| | | | |
|---|---|---|---|
| SS02031 | 0 | 0.00026458 | 1.38E-08 |
| SS06554 | 0.00331638 | 0.004957765 | 1.40E-08 |
| SS03499 | 0.006341559 | 0.008655911 | 1.41E-08 |
| SS11430 | 0.006341559 | 0.008655911 | 1.41E-08 |
| SS10034 | 0 | 0.000121409 | 1.43E-08 |
| SS02073 | 4.90E-05 | 0.00022612 | 1.43E-08 |
| SS04376 | 0.005028817 | 0.001574886 | 1.43E-08 |
| SS06748 | 0.015024593 | 0.019293503 | 1.44E-08 |
| SS10087 | 0.015024593 | 0.019293503 | 1.44E-08 |
| SS10179 | 0.015024593 | 0.019293503 | 1.44E-08 |
| SS15446 | 0.015024593 | 0.019293503 | 1.44E-08 |
| SS06738 | 0.006857422 | 0.000252586 | 1.44E-08 |
| SS08180 | 0.002284859 | 0.001128078 | 1.44E-08 |
| SS08533 | 0.00118968 | 0.000390371 | 1.46E-08 |
| SS04101 | 0.001453685 | 0.004666529 | 1.46E-08 |
| SS02956 | 0 | 0.00013862 | 1.46E-08 |
| SS01822 | 0.001255541 | 9.80E-07 | 1.47E-08 |
| SS06521 | 0.007103029 | 0.010599184 | 1.47E-08 |
| SS08789 | 0.007103029 | 0.010599184 | 1.47E-08 |
| SS10904 | 0.007103029 | 0.010599184 | 1.47E-08 |
| SS11955 | 0 | 0.001755913 | 1.49E-08 |
| SS08112 | 0.057002251 | 0.02198004 | 1.50E-08 |
| SS01899 | 2.46E-05 | 0.000231241 | 1.52E-08 |
| SS01911 | 2.46E-05 | 0.000231241 | 1.52E-08 |
| SS00451 | 0.007335951 | 0.010366245 | 1.53E-08 |
| SS12269 | 0.000130678 | 4.18E-05 | 1.55E-08 |
| SS14165 | 0 | 0.000206648 | 1.57E-08 |
| SS10736 | 0.02291162 | 0.02875441 | 1.57E-08 |
| SS13228 | 0.000667852 | 0.002020233 | 1.57E-08 |
| SS04290 | 0.006704229 | 0.001121422 | 1.58E-08 |
| SS10248 | 0.00012531 | 4.09E-05 | 1.61E-08 |
| SS05333 | 0.029420641 | 0.014314837 | 1.62E-08 |
| SS09502 | 0 | 0.000216019 | 1.66E-08 |
| SS07243 | 0.014824005 | 0.021069972 | 1.66E-08 |
| SS02187 | 0.003332965 | 0.000536835 | 1.69E-08 |
| SS06275 | 0.003332965 | 0.000536835 | 1.69E-08 |
| SS02960 | 0.026871857 | 0.008240792 | 1.69E-08 |
| SS08548 | 0.026871857 | 0.008240792 | 1.69E-08 |
| SS15208 | 0 | 0.000233959 | 1.73E-08 |
| SS04184 | 0.005716455 | 5.26E-05 | 1.76E-08 |
| SS13320 | 2.76E-06 | 0.000166546 | 1.77E-08 |
| SS04296 | 4.09E-05 | 0.001040584 | 1.77E-08 |
| SS05922 | 0 | 8.97E-05 | 1.77E-08 |
| SS15250 | 0 | 4.04E-05 | 1.78E-08 |
| SS01222 | 0.013888902 | 0.008711407 | 1.78E-08 |
| SS05452 | 0.013888902 | 0.008711407 | 1.78E-08 |
| SS03488 | 0.006704195 | 0.009222879 | 1.79E-08 |

| | | | |
|---|---|---|---|
| SS11276 | 0.006704195 | 0.009222879 | 1.79E-08 |
| SS00058 | 0.004950614 | 0.000387389 | 1.80E-08 |
| SS00799 | 0.004950614 | 0.000387389 | 1.80E-08 |
| SS08782 | 0.019752861 | 0.01494019 | 1.80E-08 |
| SS12470 | 0.019752861 | 0.01494019 | 1.80E-08 |
| SS13591 | 0 | 0.000116622 | 1.90E-08 |
| SS04732 | 0 | 0.000107444 | 1.90E-08 |
| SS13143 | 0 | 7.36E-05 | 1.92E-08 |
| SS15315 | 0 | 7.36E-05 | 1.92E-08 |
| SS04628 | 0 | 0.000239708 | 1.93E-08 |
| SS04634 | 0 | 0.000239708 | 1.93E-08 |
| SS04497 | 0.035833429 | 0.028794605 | 1.94E-08 |
| SS04658 | 0.035833429 | 0.028794605 | 1.94E-08 |
| SS05265 | 0.02203834 | 0.013120014 | 1.95E-08 |
| SS08198 | 0 | 0.000223952 | 1.99E-08 |
| SS04733 | 0 | 0.000344978 | 2.01E-08 |
| SS02526 | 0.006469874 | 0.001047421 | 2.07E-08 |
| SS00496 | 0.025274498 | 0.008648616 | 2.09E-08 |
| SS05831 | 0.025274498 | 0.008648616 | 2.09E-08 |
| SS13253 | 0.006931931 | 0.002886453 | 2.17E-08 |
| SS02023 | 0 | 0.000306396 | 2.18E-08 |
| SS07731 | 0.025509439 | 0.008311469 | 2.21E-08 |
| SS07738 | 0.025509439 | 0.008311469 | 2.21E-08 |
| SS14365 | 4.65E-05 | 0.000397732 | 2.23E-08 |
| SS06641 | 0 | 0.000392883 | 2.29E-08 |
| SS03382 | 0 | 0.000185666 | 2.32E-08 |
| SS02853 | 0.032976021 | 0.007978078 | 2.34E-08 |
| SS00065 | 0.022229625 | 0.026415092 | 2.36E-08 |
| SS08110 | 0.022229625 | 0.026415092 | 2.36E-08 |
| SS08863 | 0.022229625 | 0.026415092 | 2.36E-08 |
| SS11902 | 0.022229625 | 0.026415092 | 2.36E-08 |
| SS15355 | 0.022229625 | 0.026415092 | 2.36E-08 |
| SS06079 | 0.000329659 | 3.31E-05 | 2.37E-08 |
| SS06732 | 0 | 0.000114847 | 2.38E-08 |
| SS06517 | 0.029505301 | 0.018876081 | 2.40E-08 |
| SS00406 | 0.00046871 | 0.001061673 | 2.46E-08 |
| SS04626 | 0 | 0.000153408 | 2.48E-08 |
| SS02438 | 0.049405712 | 0.071522367 | 2.51E-08 |
| SS08543 | 0.049405712 | 0.071522367 | 2.51E-08 |
| SS04272 | 0.052395943 | 0.003454043 | 2.53E-08 |
| SS08771 | 0.052395943 | 0.003454043 | 2.53E-08 |
| SS15257 | 0.052395943 | 0.003454043 | 2.53E-08 |
| SS14377 | 0 | 0.000276035 | 2.55E-08 |
| SS07746 | 0.000931571 | 0.005066429 | 2.59E-08 |
| SS03313 | 0.020480484 | 0.024993699 | 2.63E-08 |
| SS13905 | 0 | 0.00015115 | 2.65E-08 |
| SS13789 | 1.09E-05 | 2.11E-06 | 2.66E-08 |

| | | | |
|---|---|---|---|
| SS05969 | 2.68E-06 | 0.000275699 | 2.71E-08 |
| SS09629 | 2.68E-06 | 0.000275699 | 2.71E-08 |
| SS01114 | 0 | 8.64E-05 | 2.74E-08 |
| SS04358 | 0 | 0.00034514 | 2.74E-08 |
| SS13131 | 0 | 0.00034514 | 2.74E-08 |
| SS11598 | 0 | 0.000202164 | 2.75E-08 |
| SS00222 | 0.000408378 | 0.000149452 | 2.78E-08 |
| SS03296 | 0.025497716 | 0.016000439 | 2.84E-08 |
| SS10207 | 0 | 0.000264425 | 2.85E-08 |
| SS00844 | 0.01227014 | 0.018735373 | 2.87E-08 |
| SS00895 | 0.001078914 | 0.000567867 | 2.88E-08 |
| SS14376 | 0.000365113 | 0.001457936 | 2.90E-08 |
| SS10096 | 0.000691928 | 0.001531135 | 2.94E-08 |
| SS09434 | 0 | 0.000172961 | 3.04E-08 |
| SS05888 | 0.001427763 | 0.004391003 | 3.07E-08 |
| SS10811 | 0.001832494 | 0.000179812 | 3.13E-08 |
| SS00552 | 0.000346406 | 0.001413753 | 3.14E-08 |
| SS15362 | 0.000346406 | 0.001413753 | 3.14E-08 |
| SS02155 | 0.078602723 | 0.062669731 | 3.16E-08 |
| SS05993 | 0.005386956 | 3.70E-05 | 3.21E-08 |
| SS12997 | 0.005386956 | 3.70E-05 | 3.21E-08 |
| SS10270 | 0 | 0.000146872 | 3.25E-08 |
| SS11331 | 0.000253384 | 5.26E-05 | 3.30E-08 |
| SS04551 | 0 | 0.000384602 | 3.38E-08 |
| SS13993 | 0.000756183 | 0.002065081 | 3.42E-08 |
| SS05011 | 0 | 0.000207671 | 3.45E-08 |
| SS09995 | 0.027625103 | 0.038210212 | 3.45E-08 |
| SS05014 | 0 | 0.000109868 | 3.48E-08 |
| SS09572 | 0 | 0.001205762 | 3.49E-08 |
| SS08172 | 0.017475538 | 0.02174989 | 3.52E-08 |
| SS08851 | 0.017475538 | 0.02174989 | 3.52E-08 |
| SS12013 | 2.76E-06 | 0.000184923 | 3.62E-08 |
| SS00619 | 0 | 9.69E-05 | 3.68E-08 |
| SS11154 | 0 | 5.39E-05 | 3.68E-08 |
| SS02748 | 0.000255988 | 0.000707513 | 3.70E-08 |
| SS00664 | 0 | 7.92E-05 | 3.73E-08 |
| SS03556 | 0 | 7.92E-05 | 3.73E-08 |
| SS06942 | 0 | 7.92E-05 | 3.73E-08 |
| SS09594 | 0.005220423 | 0.000225032 | 3.74E-08 |
| SS11604 | 0 | 0.000142393 | 3.74E-08 |
| SS04057 | 0 | 7.80E-05 | 3.75E-08 |
| SS13270 | 0.002129464 | 0.000192811 | 3.78E-08 |
| SS07553 | 0.000380013 | 0.001980149 | 3.87E-08 |
| SS10019 | 0 | 0.000573546 | 3.88E-08 |
| SS02914 | 0 | 0.000528816 | 3.89E-08 |
| SS12455 | 0 | 0.001432794 | 4.03E-08 |
| SS10813 | 0 | 0.0002947 | 4.05E-08 |

| | | | |
|---|---|---|---|
| SS14132 | 0 | 0.0002947 | 4.05E-08 |
| SS11333 | 0.000108885 | 2.84E-05 | 4.07E-08 |
| SS05579 | 0 | 0.000161701 | 4.11E-08 |
| SS14106 | 0 | 7.90E-05 | 4.12E-08 |
| SS15000 | 0.001468344 | 0.000634287 | 4.15E-08 |
| SS02913 | 5.45E-06 | 0.000379124 | 4.17E-08 |
| SS06186 | 0.026540098 | 0.002779902 | 4.18E-08 |
| SS10151 | 0.026540098 | 0.002779902 | 4.18E-08 |
| SS13256 | 0.026540098 | 0.002779902 | 4.18E-08 |
| SS15356 | 0.026540098 | 0.002779902 | 4.18E-08 |
| SS03517 | 0 | 0.000136402 | 4.29E-08 |
| SS07676 | 0.005089344 | 0.00064888 | 4.31E-08 |
| SS02170 | 0 | 8.47E-05 | 4.31E-08 |
| SS09490 | 2.76E-06 | 0.000312783 | 4.40E-08 |
| SS01913 | 4.63E-05 | 0.000193967 | 4.44E-08 |
| SS11145 | 4.63E-05 | 0.000193967 | 4.44E-08 |
| SS09440 | 0.006799132 | 0.00490521 | 4.45E-08 |
| SS02457 | 0 | 6.74E-05 | 4.46E-08 |
| SS03106 | 0 | 0.000366048 | 4.47E-08 |
| SS13321 | 2.45E-05 | 0.001045585 | 4.58E-08 |
| SS00182 | 0 | 0.000153649 | 4.58E-08 |
| SS13136 | 1.62E-05 | 0.000218651 | 4.62E-08 |
| SS15306 | 1.62E-05 | 0.000218651 | 4.62E-08 |
| SS13524 | 0.006426815 | 0.003385631 | 4.79E-08 |
| SS15014 | 7.65E-05 | 0.00196608 | 4.86E-08 |
| SS01499 | 0 | 9.58E-05 | 4.87E-08 |
| SS00642 | 0 | 9.69E-05 | 4.94E-08 |
| SS07000 | 0.002610871 | 0.004024501 | 4.97E-08 |
| SS12277 | 0.010196558 | 0.014225127 | 5.00E-08 |
| SS00992 | 0.000228505 | 0.000639924 | 5.13E-08 |
| SS01857 | 0.007286148 | 0.004072662 | 5.21E-08 |
| SS08932 | 0.007286148 | 0.004072662 | 5.21E-08 |
| SS04795 | 0 | 5.07E-05 | 5.21E-08 |
| SS01497 | 0.001381413 | 0.000438874 | 5.27E-08 |
| SS01683 | 0.001381413 | 0.000438874 | 5.27E-08 |
| SS06582 | 0 | 9.83E-05 | 5.33E-08 |
| SS02706 | 0.000784227 | 0.001739619 | 5.37E-08 |
| SS09430 | 0 | 0.000304845 | 5.43E-08 |
| SS11152 | 0 | 0.000362081 | 5.47E-08 |
| SS06274 | 0.007923961 | 0.005366644 | 5.48E-08 |
| SS08905 | 0.007367952 | 0.002897586 | 5.49E-08 |
| SS02955 | 0 | 0.000355553 | 5.51E-08 |
| SS04061 | 0 | 0.000308111 | 5.76E-08 |
| SS07928 | 0.000713801 | 0.001290642 | 5.79E-08 |
| SS15510 | 6.53E-05 | 0.000230991 | 5.80E-08 |
| SS02109 | 0.00179556 | 0.000820964 | 5.81E-08 |
| SS12590 | 0.00179556 | 0.000820964 | 5.81E-08 |

| | | | |
|---|---|---|---|
| SS03115 | 0.000292084 | 0.001305062 | 6.05E-08 |
| SS10203 | 0.000292084 | 0.001305062 | 6.05E-08 |
| SS10222 | 0.000292084 | 0.001305062 | 6.05E-08 |
| SS00446 | 0.001979115 | 0.000788122 | 6.30E-08 |
| SS10951 | 0.001979115 | 0.000788122 | 6.30E-08 |
| SS09321 | 0.000326735 | 3.26E-05 | 6.39E-08 |
| SS00763 | 0.027380586 | 0.032891101 | 6.41E-08 |
| SS00787 | 0.027380586 | 0.032891101 | 6.41E-08 |
| SS00794 | 0.027380586 | 0.032891101 | 6.41E-08 |
| SS02529 | 0.027380586 | 0.032891101 | 6.41E-08 |
| SS09794 | 0 | 0.000148872 | 6.49E-08 |
| SS15187 | 0 | 0.000148872 | 6.49E-08 |
| SS08825 | 4.63E-05 | 9.47E-06 | 6.57E-08 |
| SS08563 | 0.006020674 | 0.010297328 | 6.61E-08 |
| SS02690 | 0.020535781 | 0.016862578 | 6.66E-08 |
| SS03992 | 0.020535781 | 0.016862578 | 6.66E-08 |
| SS14124 | 0.020535781 | 0.016862578 | 6.66E-08 |
| SS11187 | 0.000963779 | 0.00235513 | 6.73E-08 |
| SS05707 | 0 | 0.000350676 | 6.74E-08 |
| SS05930 | 0.012506411 | 0.008417693 | 6.78E-08 |
| SS08191 | 0.012506411 | 0.008417693 | 6.78E-08 |
| SS14176 | 0.01050322 | 0.001827737 | 6.81E-08 |
| SS05147 | 0.005987503 | 0.008181229 | 6.88E-08 |
| SS04382 | 0.009261902 | 0.000454991 | 6.97E-08 |
| SS04379 | 0.029459972 | 0.001109083 | 7.01E-08 |
| SS11293 | 0 | 8.22E-05 | 7.13E-08 |
| SS03849 | 0 | 0.000314877 | 7.22E-08 |
| SS14066 | 0.004948974 | 7.11E-05 | 7.44E-08 |
| SS13140 | 0 | 0.000117769 | 7.48E-08 |
| SS15312 | 0 | 0.000117769 | 7.48E-08 |
| SS04532 | 0.013017354 | 0.009774192 | 7.49E-08 |
| SS06279 | 0.027195494 | 0.031766824 | 7.58E-08 |
| SS15496 | 0.027195494 | 0.031766824 | 7.58E-08 |
| SS10843 | 0.026039548 | 0.031031043 | 7.81E-08 |
| SS14257 | 0.026039548 | 0.031031043 | 7.81E-08 |
| SS14286 | 0.026039548 | 0.031031043 | 7.81E-08 |
| SS13520 | 5.73E-05 | 0.000320093 | 7.89E-08 |
| SS07732 | 0.000135644 | 0.000770071 | 7.90E-08 |
| SS04533 | 0.007234912 | 0.001597467 | 8.05E-08 |
| SS12197 | 0.000890428 | 5.61E-05 | 8.05E-08 |
| SS12126 | 0.010093477 | 0.001639824 | 8.05E-08 |
| SS07989 | 0 | 9.64E-05 | 8.14E-08 |
| SS09593 | 0 | 0.000178137 | 8.17E-08 |
| SS13803 | 0.000155074 | 0.000547181 | 8.18E-08 |
| SS00343 | 0 | 7.99E-05 | 8.30E-08 |
| SS13943 | 0 | 0.000253753 | 8.31E-08 |
| SS06781 | 0.006391637 | 0.000425062 | 8.37E-08 |

| | | | |
|---|---|---|---|
| SS07585 | 0.009837203 | 0.018468333 | 8.46E-08 |
| SS09445 | 0.00581356 | 0.00429698 | 8.52E-08 |
| SS10307 | 0.006759996 | 0.009134161 | 8.53E-08 |
| SS02089 | 0.005120634 | 0.006740924 | 8.62E-08 |
| SS03485 | 0.005120634 | 0.006740924 | 8.62E-08 |
| SS06755 | 0.005120634 | 0.006740924 | 8.62E-08 |
| SS11266 | 0.005120634 | 0.006740924 | 8.62E-08 |
| SS00147 | 0.000372924 | 0.000776389 | 8.63E-08 |
| SS05989 | 0.01222114 | 0.002253012 | 8.73E-08 |
| SS14057 | 0 | 0.000204482 | 8.75E-08 |
| SS09083 | 0.026523043 | 0.031407327 | 8.76E-08 |
| SS09090 | 0.026523043 | 0.031407327 | 8.76E-08 |
| SS05389 | 0.025538802 | 0.018600166 | 8.92E-08 |
| SS05427 | 0.025538802 | 0.018600166 | 8.92E-08 |
| SS09621 | 0 | 0.000256363 | 8.96E-08 |
| SS14064 | 0.000874725 | 0.002958202 | 9.02E-08 |
| SS00566 | 0 | 0.000234008 | 9.16E-08 |
| SS00905 | 0.000120825 | 0.003040966 | 9.19E-08 |
| SS09962 | 0.000120825 | 0.003040966 | 9.19E-08 |
| SS14156 | 2.72E-05 | 0.000292817 | 9.23E-08 |
| SS15233 | 2.72E-05 | 0.000292817 | 9.23E-08 |
| SS06211 | 0.001440862 | 2.11E-05 | 9.27E-08 |
| SS07127 | 0.001440862 | 2.11E-05 | 9.27E-08 |
| SS03619 | 0.00545206 | 0.001508867 | 9.44E-08 |
| SS05215 | 0.00545206 | 0.001508867 | 9.44E-08 |
| SS10421 | 0.00545206 | 0.001508867 | 9.44E-08 |
| SS10443 | 0.00545206 | 0.001508867 | 9.44E-08 |
| SS04434 | 0.007540943 | 0.001885173 | 9.45E-08 |
| SS13703 | 0 | 9.72E-05 | 9.47E-08 |
| SS01474 | 5.53E-06 | 0.000122224 | 9.53E-08 |
| SS01951 | 0 | 0.000121058 | 9.86E-08 |
| SS01119 | 0.014160007 | 0.0100844 | 9.97E-08 |
| SS03017 | 0.014160007 | 0.0100844 | 9.97E-08 |
| SS14473 | 0.014160007 | 0.0100844 | 9.97E-08 |
| SS12905 | 0 | 0.00011845 | 1.02E-07 |
| SS12918 | 0 | 0.00011845 | 1.02E-07 |
| SS09494 | 2.69E-05 | 0.000404941 | 1.02E-07 |
| SS10801 | 0.001944865 | 0.003255822 | 1.03E-07 |
| SS11167 | 0 | 0.000126332 | 1.03E-07 |
| SS10787 | 0.00662387 | 4.86E-05 | 1.05E-07 |
| SS02596 | 0.008622094 | 0.006765252 | 1.06E-07 |
| SS08816 | 0.008622094 | 0.006765252 | 1.06E-07 |
| SS13694 | 0.008622094 | 0.006765252 | 1.06E-07 |
| SS04552 | 0 | 0.000463528 | 1.08E-07 |
| SS01040 | 0 | 0.00011822 | 1.08E-07 |
| SS15251 | 0 | 6.66E-05 | 1.08E-07 |
| SS05567 | 0.00741202 | 0.012623883 | 1.10E-07 |

| | | | |
|---|---|---|---|
| SS13751 | 0.00250287 | 5.24E-06 | 1.10E-07 |
| SS03251 | 0.00154049 | 0.003100582 | 1.10E-07 |
| SS03352 | 0.00154049 | 0.003100582 | 1.10E-07 |
| SS02024 | 0 | 0.000529459 | 1.13E-07 |
| SS07037 | 0 | 6.86E-05 | 1.13E-07 |
| SS13071 | 0 | 6.86E-05 | 1.13E-07 |
| SS12289 | 0 | 0.003225844 | 1.15E-07 |
| SS04927 | 0.000699739 | 9.46E-05 | 1.16E-07 |
| SS05186 | 0.006191417 | 0.000594512 | 1.19E-07 |
| SS06490 | 0.006191417 | 0.000594512 | 1.19E-07 |
| SS03670 | 0.007103511 | 0.001927526 | 1.20E-07 |
| SS07078 | 0.007103511 | 0.001927526 | 1.20E-07 |
| SS04326 | 0 | 0.000314266 | 1.25E-07 |
| SS00468 | 0 | 0.000366405 | 1.26E-07 |
| SS08136 | 0 | 0.00011484 | 1.27E-07 |
| SS05984 | 0.012242853 | 0.009098249 | 1.30E-07 |
| SS12991 | 0.012242853 | 0.009098249 | 1.30E-07 |
| SS02958 | 0 | 0.00080173 | 1.30E-07 |
| SS09472 | 8.21E-06 | 8.61E-05 | 1.31E-07 |
| SS13991 | 3.81E-05 | 0.000206267 | 1.33E-07 |
| SS09955 | 0.017759455 | 0.014550595 | 1.35E-07 |
| SS10017 | 0.017759455 | 0.014550595 | 1.35E-07 |
| SS15106 | 0.017759455 | 0.014550595 | 1.35E-07 |
| SS10817 | 0 | 8.21E-05 | 1.36E-07 |
| SS14144 | 0 | 8.21E-05 | 1.36E-07 |
| SS00420 | 0 | 0.000262175 | 1.36E-07 |
| SS03520 | 0 | 0.000100461 | 1.37E-07 |
| SS04620 | 0.014634842 | 0.02192768 | 1.37E-07 |
| SS10644 | 0.011920569 | 9.98E-05 | 1.38E-07 |
| SS03385 | 0 | 6.97E-05 | 1.38E-07 |
| SS04685 | 0.002807008 | 0.000661695 | 1.39E-07 |
| SS02247 | 0.00103737 | 0.000607628 | 1.41E-07 |
| SS04056 | 0.00103737 | 0.000607628 | 1.41E-07 |
| SS07197 | 0.010976576 | 0.008657187 | 1.43E-07 |
| SS07221 | 0.010976576 | 0.008657187 | 1.43E-07 |
| SS00825 | 0.002136999 | 0.004416342 | 1.49E-07 |
| SS08414 | 0.002136999 | 0.004416342 | 1.49E-07 |
| SS14464 | 0.002136999 | 0.004416342 | 1.49E-07 |
| SS12327 | 0 | 0.000105959 | 1.49E-07 |
| SS07552 | 0 | 0.000377677 | 1.49E-07 |
| SS09271 | 0 | 6.35E-05 | 1.50E-07 |
| SS14784 | 0.043157359 | 0.003934213 | 1.50E-07 |
| SS02235 | 2.68E-06 | 0.000156638 | 1.51E-07 |
| SS00918 | 0 | 0.000134893 | 1.52E-07 |
| SS11492 | 0.000171499 | 5.10E-05 | 1.52E-07 |
| SS10967 | 2.68E-06 | 0.000146749 | 1.53E-07 |
| SS08157 | 0.033421676 | 0.022845754 | 1.56E-07 |

| | | | |
|---|---|---|---|
| SS10760 | 0.033421676 | 0.022845754 | 1.56E-07 |
| SS02585 | 5.44E-05 | 0.000278178 | 1.58E-07 |
| SS02589 | 5.44E-05 | 0.000278178 | 1.58E-07 |
| SS07573 | 0 | 8.63E-05 | 1.58E-07 |
| SS05191 | 0.041810367 | 0.029201322 | 1.58E-07 |
| SS06493 | 0.041810367 | 0.029201322 | 1.58E-07 |
| SS00346 | 0 | 5.77E-05 | 1.59E-07 |
| SS04103 | 0.000116535 | 0.001432485 | 1.60E-07 |
| SS01248 | 0.001359058 | 0.000807944 | 1.60E-07 |
| SS03434 | 0 | 0.0001335 | 1.62E-07 |
| SS07055 | 0.000834546 | 0.002146318 | 1.64E-07 |
| SS02927 | 2.68E-06 | 0.001914934 | 1.64E-07 |
| SS11987 | 0 | 8.32E-05 | 1.64E-07 |
| SS02409 | 0 | 0.00034715 | 1.65E-07 |
| SS05791 | 0 | 0.000176543 | 1.70E-07 |
| SS04629 | 0 | 0.000122063 | 1.70E-07 |
| SS12397 | 0.004107661 | 0.000126734 | 1.73E-07 |
| SS12532 | 0.004107661 | 0.000126734 | 1.73E-07 |
| SS05931 | 0 | 0.000241299 | 1.75E-07 |
| SS04201 | 0.016142207 | 0.012992933 | 1.77E-07 |
| SS06918 | 0.016142207 | 0.012992933 | 1.77E-07 |
| SS07184 | 0.016142207 | 0.012992933 | 1.77E-07 |
| SS15081 | 0.016142207 | 0.012992933 | 1.77E-07 |
| SS13442 | 0.004263056 | 0.000489129 | 1.79E-07 |
| SS00532 | 5.45E-06 | 0.00033846 | 1.81E-07 |
| SS08882 | 0.041163551 | 0.006769486 | 1.82E-07 |
| SS03718 | 5.45E-06 | 0.000809159 | 1.86E-07 |
| SS07772 | 5.45E-06 | 0.000809159 | 1.86E-07 |
| SS08790 | 5.45E-06 | 0.000809159 | 1.86E-07 |
| SS04078 | 0.048627254 | 0.059399845 | 1.86E-07 |
| SS10669 | 0.048627254 | 0.059399845 | 1.86E-07 |
| SS04696 | 0 | 0.000159845 | 1.86E-07 |
| SS08598 | 0.030039046 | 0.000497972 | 1.88E-07 |
| SS07052 | 0.005886234 | 0.000286726 | 1.94E-07 |
| SS00317 | 0.027517618 | 0.033224247 | 1.94E-07 |
| SS00929 | 0.027517618 | 0.033224247 | 1.94E-07 |
| SS07471 | 0.027517618 | 0.033224247 | 1.94E-07 |
| SS13830 | 0.027517618 | 0.033224247 | 1.94E-07 |
| SS15479 | 0.027517618 | 0.033224247 | 1.94E-07 |
| SS02653 | 0 | 0.00017804 | 1.96E-07 |
| SS01081 | 0 | 8.22E-05 | 1.96E-07 |
| SS08177 | 0.010842985 | 0.005925675 | 1.97E-07 |
| SS10149 | 0.010842985 | 0.005925675 | 1.97E-07 |
| SS01384 | 0.008427035 | 0.002416952 | 1.98E-07 |
| SS02048 | 0 | 0.000433154 | 2.01E-07 |
| SS08976 | 0 | 0.000433154 | 2.01E-07 |
| SS12244 | 0.000577434 | 6.22E-06 | 2.03E-07 |

| | | | |
|---|---|---|---|
| SS04002 | 0.00101638 | 0.000432885 | 2.04E-07 |
| SS01970 | 0 | 0.000154521 | 2.06E-07 |
| SS05757 | 0 | 8.93E-05 | 2.07E-07 |
| SS06262 | 0.001952676 | 0.001365172 | 2.07E-07 |
| SS08692 | 0 | 5.77E-05 | 2.14E-07 |
| SS14362 | 0.006521753 | 0.00069375 | 2.16E-07 |
| SS06037 | 0.011513074 | 0.009440208 | 2.17E-07 |
| SS07822 | 0.001402965 | 0.000228138 | 2.20E-07 |
| SS09412 | 0.001402965 | 0.000228138 | 2.20E-07 |
| SS00600 | 0 | 7.77E-05 | 2.23E-07 |
| SS03081 | 0 | 0.00013163 | 2.25E-07 |
| SS15466 | 0 | 6.21E-05 | 2.26E-07 |
| SS07206 | 0 | 0.000441612 | 2.28E-07 |
| SS13007 | 0 | 0.000441612 | 2.28E-07 |
| SS15273 | 0 | 0.000441612 | 2.28E-07 |
| SS05784 | 5.53E-06 | 0.000147162 | 2.30E-07 |
| SS04808 | 0.003305242 | 0.001771963 | 2.31E-07 |
| SS07048 | 0.003305242 | 0.001771963 | 2.31E-07 |
| SS11173 | 1.10E-05 | 0.000432781 | 2.32E-07 |
| SS02946 | 9.53E-05 | 1.41E-06 | 2.34E-07 |
| SS09942 | 0.022934113 | 0.030331896 | 2.35E-07 |
| SS10345 | 0.022934113 | 0.030331896 | 2.35E-07 |
| SS13768 | 0.000326654 | 0.000810957 | 2.37E-07 |
| SS13791 | 0.000326654 | 0.000810957 | 2.37E-07 |
| SS14891 | 0.000326654 | 0.000810957 | 2.37E-07 |
| SS05717 | 0.002812456 | 0.001706439 | 2.37E-07 |
| SS06351 | 5.45E-06 | 0.00020336 | 2.45E-07 |
| SS06857 | 5.45E-06 | 0.00020336 | 2.45E-07 |
| SS05280 | 0.000473677 | 9.16E-05 | 2.47E-07 |
| SS14188 | 0 | 0.000127617 | 2.49E-07 |
| SS09633 | 1.09E-05 | 3.47E-07 | 2.50E-07 |
| SS09163 | 4.63E-05 | 0.000240122 | 2.55E-07 |
| SS09783 | 4.63E-05 | 0.000240122 | 2.55E-07 |
| SS15176 | 4.63E-05 | 0.000240122 | 2.55E-07 |
| SS10074 | 0.010146159 | 0.012825468 | 2.55E-07 |
| SS03509 | 0.008982516 | 0.003486205 | 2.57E-07 |
| SS08971 | 0.008982516 | 0.003486205 | 2.57E-07 |
| SS12591 | 0.008982516 | 0.003486205 | 2.57E-07 |
| SS13918 | 0.008982516 | 0.003486205 | 2.57E-07 |
| SS15499 | 0.008982516 | 0.003486205 | 2.57E-07 |
| SS04997 | 0.018778565 | 0.005715535 | 2.57E-07 |
| SS10765 | 0.000961014 | 0.000323906 | 2.57E-07 |
| SS02984 | 0.007461777 | 0.000373556 | 2.58E-07 |
| SS03457 | 0.014724526 | 0.008271812 | 2.61E-07 |
| SS06953 | 0.014724526 | 0.008271812 | 2.61E-07 |
| SS07296 | 0.014724526 | 0.008271812 | 2.61E-07 |
| SS07476 | 0.014724526 | 0.008271812 | 2.61E-07 |

| | | | |
|---|---|---|---|
| SS15092 | 0.014724526 | 0.008271812 | 2.61E-07 |
| SS10379 | 0 | 0.000120402 | 2.61E-07 |
| SS04416 | 0 | 0.000129547 | 2.64E-07 |
| SS01573 | 0.0070165 | 0.009298734 | 2.67E-07 |
| SS04630 | 0 | 0.000135788 | 2.69E-07 |
| SS02290 | 0.012503646 | 0.009367168 | 2.71E-07 |
| SS06973 | 0.012503646 | 0.009367168 | 2.71E-07 |
| SS04734 | 0 | 6.55E-05 | 2.73E-07 |
| SS03146 | 0.000370401 | 0.0007977 | 2.80E-07 |
| SS10558 | 0.000370401 | 0.0007977 | 2.80E-07 |
| SS06747 | 0.01855787 | 0.025289362 | 2.85E-07 |
| SS10086 | 0.01855787 | 0.025289362 | 2.85E-07 |
| SS10175 | 0.01855787 | 0.025289362 | 2.85E-07 |
| SS15391 | 0.01855787 | 0.025289362 | 2.85E-07 |
| SS00238 | 0 | 0.000222556 | 3.01E-07 |
| SS07561 | 0 | 0.000222556 | 3.01E-07 |
| SS03909 | 0 | 0.000655253 | 3.01E-07 |
| SS13618 | 0.002470662 | 0.003909203 | 3.03E-07 |
| SS01285 | 0 | 0.000154068 | 3.17E-07 |
| SS12466 | 0.002946621 | 6.40E-05 | 3.31E-07 |
| SS12483 | 0.002946621 | 6.40E-05 | 3.31E-07 |
| SS12519 | 0.002946621 | 6.40E-05 | 3.31E-07 |
| SS12562 | 0.002946621 | 6.40E-05 | 3.31E-07 |
| SS07729 | 2.73E-05 | 0.000303377 | 3.42E-07 |
| SS11602 | 2.68E-06 | 0.000146297 | 3.44E-07 |
| SS07484 | 0 | 8.37E-05 | 3.50E-07 |
| SS06804 | 0.019108969 | 0.000247036 | 3.54E-07 |
| SS12094 | 0.019108969 | 0.000247036 | 3.54E-07 |
| SS12131 | 0.019108969 | 0.000247036 | 3.54E-07 |
| SS03454 | 4.90E-05 | 2.07E-06 | 3.56E-07 |
| SS09274 | 0.005085616 | 0.002571731 | 3.56E-07 |
| SS06323 | 0 | 6.55E-05 | 3.61E-07 |
| SS06423 | 0 | 6.55E-05 | 3.61E-07 |
| SS04235 | 0.030254327 | 0.035164072 | 3.62E-07 |
| SS02901 | 0.000656795 | 0.000161102 | 3.68E-07 |
| SS03116 | 0.001490779 | 0.003242519 | 3.70E-07 |
| SS10206 | 0.001490779 | 0.003242519 | 3.70E-07 |
| SS15009 | 0.01100306 | 0.00879996 | 3.73E-07 |
| SS03686 | 0 | 4.40E-05 | 3.81E-07 |
| SS07097 | 0 | 4.40E-05 | 3.81E-07 |
| SS11136 | 0.000978323 | 0.000253847 | 3.83E-07 |
| SS09042 | 0.043983704 | 0.057379824 | 3.90E-07 |
| SS13116 | 0.043983704 | 0.057379824 | 3.90E-07 |
| SS08147 | 0.003372789 | 0.001296782 | 3.90E-07 |
| SS09989 | 0 | 0.000156356 | 3.94E-07 |
| SS12062 | 0.0002358 | 0.002650569 | 3.94E-07 |
| SS05928 | 0.003606582 | 0.005187576 | 4.00E-07 |

| | | | |
|---|---|---|---|
| SS04823 | 0.008618974 | 0.001489618 | 4.06E-07 |
| SS10950 | 0.008618974 | 0.001489618 | 4.06E-07 |
| SS09498 | 0.00046863 | 0.000820651 | 4.06E-07 |
| SS06615 | 0 | 0.000107674 | 4.07E-07 |
| SS15254 | 0 | 9.40E-05 | 4.08E-07 |
| SS09610 | 0.000517343 | 3.24E-05 | 4.11E-07 |
| SS04637 | 2.72E-05 | 0.00019341 | 4.12E-07 |
| SS03380 | 0.007860738 | 0.004021806 | 4.13E-07 |
| SS04405 | 0.006988823 | 3.09E-05 | 4.14E-07 |
| SS11467 | 0 | 6.95E-05 | 4.16E-07 |
| SS00563 | 0 | 0.000101063 | 4.18E-07 |
| SS06229 | 0 | 0.000101063 | 4.18E-07 |
| SS07940 | 0 | 0.000101063 | 4.18E-07 |
| SS13077 | 0 | 6.13E-05 | 4.22E-07 |
| SS02618 | 0.003002904 | 0.001417884 | 4.24E-07 |
| SS01198 | 0.002600055 | 0.001035568 | 4.37E-07 |
| SS01263 | 0.002600055 | 0.001035568 | 4.37E-07 |
| SS02377 | 0.005439237 | 0.000111335 | 4.41E-07 |
| SS02641 | 0.005439237 | 0.000111335 | 4.41E-07 |
| SS07817 | 0 | 9.42E-05 | 4.43E-07 |
| SS07734 | 1.09E-05 | 0.000510119 | 4.45E-07 |
| SS04384 | 0.010979467 | 0.013351929 | 4.52E-07 |
| SS04986 | 0.010979467 | 0.013351929 | 4.52E-07 |
| SS04716 | 5.45E-06 | 0.000124321 | 4.56E-07 |
| SS00974 | 2.76E-06 | 9.99E-05 | 4.58E-07 |
| SS13959 | 0 | 7.09E-05 | 4.59E-07 |
| SS06361 | 0.002644926 | 0.000141627 | 4.63E-07 |
| SS08366 | 0 | 0.000192708 | 4.66E-07 |
| SS00593 | 0 | 0.000161497 | 4.72E-07 |
| SS13156 | 0 | 0.000161497 | 4.72E-07 |
| SS07551 | 0.000538494 | 0.001372653 | 4.76E-07 |
| SS03855 | 5.45E-06 | 0.00013115 | 4.77E-07 |
| SS04995 | 0.001124622 | 0.002734523 | 5.00E-07 |
| SS15091 | 0 | 0.000165885 | 5.04E-07 |
| SS13742 | 0.001190564 | 0.000199891 | 5.12E-07 |
| SS09988 | 0.045616275 | 0.038292957 | 5.17E-07 |
| SS04610 | 0.000136447 | 0.000461124 | 5.21E-07 |
| SS04633 | 0.000136447 | 0.000461124 | 5.21E-07 |
| SS12158 | 0.001757262 | 0.000251988 | 5.22E-07 |
| SS12492 | 0 | 5.69E-05 | 5.25E-07 |
| SS03987 | 0 | 7.67E-05 | 5.26E-07 |
| SS08295 | 0.003580901 | 0.000284363 | 5.27E-07 |
| SS02975 | 0.001291396 | 0.002484545 | 5.28E-07 |
| SS04776 | 0.020596515 | 0.006174398 | 5.33E-07 |
| SS06450 | 0.02353724 | 0.028264304 | 5.36E-07 |
| SS14883 | 0.02353724 | 0.028264304 | 5.36E-07 |
| SS08490 | 0 | 0.000120722 | 5.36E-07 |

| | | | |
|---|---|---|---|
| SS00115 | 0.006044555 | 0.000488731 | 5.38E-07 |
| SS04241 | 0.021944114 | 0.025690109 | 5.38E-07 |
| SS15410 | 0.021944114 | 0.025690109 | 5.38E-07 |
| SS00400 | 0.001159996 | 5.57E-05 | 5.42E-07 |
| SS09643 | 8.21E-06 | 0.000168988 | 5.42E-07 |
| SS00834 | 0.014621216 | 0.012098393 | 5.52E-07 |
| SS08452 | 0.014621216 | 0.012098393 | 5.52E-07 |
| SS14466 | 0.014621216 | 0.012098393 | 5.52E-07 |
| SS13643 | 0 | 0.000348345 | 5.54E-07 |
| SS13603 | 0 | 3.58E-05 | 5.60E-07 |
| SS03490 | 0.003943812 | 0.006180452 | 5.60E-07 |
| SS11278 | 0.003943812 | 0.006180452 | 5.60E-07 |
| SS04889 | 0.000111729 | 1.94E-05 | 5.68E-07 |
| SS02954 | 0 | 0.000829725 | 5.73E-07 |
| SS04081 | 0 | 4.47E-05 | 5.75E-07 |
| SS06080 | 0.002432444 | 0.000141198 | 5.75E-07 |
| SS01827 | 0 | 0.000231706 | 5.81E-07 |
| SS07712 | 0.007000155 | 0.00304365 | 5.90E-07 |
| SS07737 | 0.007000155 | 0.00304365 | 5.90E-07 |
| SS08348 | 0 | 0.000389128 | 5.91E-07 |
| SS10730 | 0.011166703 | 0.004290887 | 5.92E-07 |
| SS06689 | 0.000590212 | 0.001436636 | 6.11E-07 |
| SS05693 | 0.01604454 | 0.012125383 | 6.13E-07 |
| SS10742 | 0.01604454 | 0.012125383 | 6.13E-07 |
| SS11107 | 0.01604454 | 0.012125383 | 6.13E-07 |
| SS00099 | 0.01149424 | 0.001787802 | 6.13E-07 |
| SS06139 | 0 | 0.000111366 | 6.22E-07 |
| SS06171 | 0 | 0.000111366 | 6.22E-07 |
| SS11592 | 5.72E-05 | 0.000232082 | 6.30E-07 |
| SS11635 | 5.72E-05 | 0.000232082 | 6.30E-07 |
| SS11639 | 5.72E-05 | 0.000232082 | 6.30E-07 |
| SS04477 | 0.000340476 | 6.09E-06 | 6.34E-07 |
| SS09320 | 0.005077404 | 0.002557646 | 6.40E-07 |
| SS10935 | 0.005077404 | 0.002557646 | 6.40E-07 |
| SS02851 | 0 | 0.000625431 | 6.41E-07 |
| SS01676 | 0.000634842 | 7.08E-05 | 6.47E-07 |
| SS06057 | 0.018005314 | 0.023937836 | 6.47E-07 |
| SS10646 | 0.018005314 | 0.023937836 | 6.47E-07 |
| SS11891 | 0.018005314 | 0.023937836 | 6.47E-07 |
| SS06088 | 0 | 0.000155377 | 6.51E-07 |
| SS10581 | 0.003582105 | 0.000449008 | 6.55E-07 |
| SS11229 | 0 | 0.000156521 | 6.58E-07 |
| SS11749 | 0 | 0.000156521 | 6.58E-07 |
| SS10071 | 3.82E-05 | 0.000169181 | 6.77E-07 |
| SS02919 | 0 | 0.000360075 | 6.94E-07 |
| SS11092 | 0 | 0.000179409 | 6.94E-07 |
| SS00003 | 0.005016074 | 0.000280521 | 6.96E-07 |

| | | | |
|---|---|---|---|
| SS01192 | 0.005016074 | 0.000280521 | 6.96E-07 |
| SS01255 | 0.005016074 | 0.000280521 | 6.96E-07 |
| SS02342 | 0.015282347 | 0.005762223 | 6.98E-07 |
| SS04679 | 0.015282347 | 0.005762223 | 6.98E-07 |
| SS09932 | 0.00013881 | 4.21E-05 | 6.99E-07 |
| SS13660 | 0.005494075 | 0.002492016 | 7.00E-07 |
| SS07625 | 0 | 0.000178771 | 7.19E-07 |
| SS05933 | 0.002910686 | 0.00027611 | 7.23E-07 |
| SS05546 | 0 | 0.00022903 | 7.29E-07 |
| SS02998 | 0.002590523 | 0.001566895 | 7.30E-07 |
| SS03259 | 5.37E-06 | 0.000102198 | 7.41E-07 |
| SS00037 | 0.009715472 | 0.000198476 | 7.43E-07 |
| SS04369 | 0.009715472 | 0.000198476 | 7.43E-07 |
| SS01923 | 3.81E-05 | 1.38E-05 | 7.45E-07 |
| SS03896 | 0 | 0.000968354 | 7.48E-07 |
| SS03397 | 0 | 5.07E-05 | 7.51E-07 |
| SS06683 | 0 | 0.000129435 | 7.54E-07 |
| SS03558 | 0 | 0.000109532 | 7.56E-07 |
| SS00768 | 0.007415507 | 0.000266642 | 7.82E-07 |
| SS06698 | 0.000397482 | 3.20E-05 | 7.98E-07 |
| SS07514 | 3.29E-05 | 0.00033892 | 8.00E-07 |
| SS09246 | 0 | 4.46E-05 | 8.02E-07 |
| SS06402 | 0.002856765 | 4.83E-05 | 8.06E-07 |
| SS10769 | 0.033617045 | 0.016354242 | 8.13E-07 |
| SS01369 | 0 | 9.28E-05 | 8.19E-07 |
| SS02896 | 0 | 9.28E-05 | 8.19E-07 |
| SS13813 | 3.81E-05 | 1.17E-05 | 8.28E-07 |
| SS03014 | 0.012552566 | 0.014915493 | 8.34E-07 |
| SS05516 | 0.012552566 | 0.014915493 | 8.34E-07 |
| SS09969 | 0.012552566 | 0.014915493 | 8.34E-07 |
| SS14472 | 0.012552566 | 0.014915493 | 8.34E-07 |
| SS09027 | 2.68E-06 | 0.000115941 | 8.39E-07 |
| SS02917 | 0 | 0.000498786 | 8.56E-07 |
| SS03554 | 0 | 0.000498786 | 8.56E-07 |
| SS03899 | 0 | 0.000372876 | 8.67E-07 |
| SS04289 | 0.002460488 | 0.000883878 | 8.70E-07 |
| SS00852 | 0.005687528 | 0.002351528 | 8.72E-07 |
| SS08367 | 5.48E-05 | 0.000403174 | 8.72E-07 |
| SS11056 | 8.13E-06 | 0.00033189 | 8.76E-07 |
| SS10796 | 0.003984312 | 3.46E-05 | 8.78E-07 |
| SS05738 | 0.000523434 | 0.001150525 | 8.87E-07 |
| SS12059 | 0.000523434 | 0.001150525 | 8.87E-07 |
| SS03363 | 0 | 8.95E-05 | 8.92E-07 |
| SS13896 | 0 | 8.95E-05 | 8.92E-07 |
| SS02916 | 0 | 8.96E-05 | 8.95E-07 |
| SS05896 | 1.63E-05 | 0.000156316 | 9.19E-07 |
| SS11834 | 0 | 8.77E-05 | 9.30E-07 |

| | | | |
|---|---|---|---|
| SS01066 | 0.009388852 | 0.001688027 | 9.33E-07 |
| SS11445 | 0.010794822 | 0.01345888 | 9.34E-07 |
| SS06332 | 0.01571074 | 0.011311726 | 9.38E-07 |
| SS06432 | 0.01571074 | 0.011311726 | 9.38E-07 |
| SS04495 | 0 | 0.000128684 | 9.40E-07 |
| SS04545 | 0.012181294 | 0.017271473 | 9.43E-07 |
| SS04582 | 0.012181294 | 0.017271473 | 9.43E-07 |
| SS04231 | 0.034817404 | 0.039644221 | 9.50E-07 |
| SS10472 | 0.034817404 | 0.039644221 | 9.50E-07 |
| SS14950 | 0 | 0.000110239 | 9.54E-07 |
| SS06848 | 0.004298751 | 0.000697498 | 9.57E-07 |
| SS03268 | 0.006401409 | 0.004487538 | 9.70E-07 |
| SS12989 | 0 | 7.80E-05 | 9.74E-07 |
| SS15272 | 0 | 7.80E-05 | 9.74E-07 |
| SS07333 | 0.000141574 | 6.40E-06 | 9.91E-07 |
| SS01456 | 0 | 7.48E-05 | 1.01E-06 |
| SS13055 | 0 | 7.48E-05 | 1.01E-06 |
| SS13832 | 0.024209084 | 0.017656942 | 1.01E-06 |
| SS01679 | 0 | 0.000220793 | 1.02E-06 |
| SS08486 | 0 | 0.000220793 | 1.02E-06 |
| SS08458 | 0 | 0.000134042 | 1.02E-06 |
| SS03117 | 0.00222369 | 0.003902507 | 1.04E-06 |
| SS10208 | 0.00222369 | 0.003902507 | 1.04E-06 |
| SS09108 | 0.004051813 | 0.000328676 | 1.05E-06 |
| SS00228 | 0.011648592 | 0.008894934 | 1.07E-06 |
| SS00994 | 0.011648592 | 0.008894934 | 1.07E-06 |
| SS12449 | 9.80E-05 | 5.48E-06 | 1.08E-06 |
| SS14577 | 0.000441068 | 4.40E-05 | 1.10E-06 |
| SS14664 | 0.000441068 | 4.40E-05 | 1.10E-06 |
| SS05665 | 0.005865485 | 0.001256446 | 1.10E-06 |
| SS04727 | 5.45E-06 | 0.000529514 | 1.11E-06 |
| SS10060 | 5.45E-06 | 0.000529514 | 1.11E-06 |
| SS01800 | 5.45E-06 | 0.000228738 | 1.11E-06 |
| SS06177 | 5.45E-06 | 0.000228738 | 1.11E-06 |
| SS14997 | 0 | 6.60E-05 | 1.11E-06 |
| SS15005 | 0 | 6.60E-05 | 1.11E-06 |
| SS00557 | 0.02015344 | 0.017362319 | 1.11E-06 |
| SS00850 | 0.02015344 | 0.017362319 | 1.11E-06 |
| SS07683 | 0.001688316 | 9.67E-05 | 1.18E-06 |
| SS00767 | 0.020657994 | 0.001866356 | 1.18E-06 |
| SS04849 | 0 | 0.000756515 | 1.21E-06 |
| SS04858 | 0 | 0.000756515 | 1.21E-06 |
| SS03869 | 0 | 3.57E-05 | 1.21E-06 |
| SS03900 | 0 | 0.000217513 | 1.22E-06 |
| SS01056 | 0.003377515 | 0.000109239 | 1.22E-06 |
| SS01316 | 0.003377515 | 0.000109239 | 1.22E-06 |
| SS14969 | 0.003377515 | 0.000109239 | 1.22E-06 |

| | | | |
|---|---|---|---|
| SS12211 | 0.000100672 | 0.000333476 | 1.22E-06 |
| SS02885 | 0 | 0.000197578 | 1.22E-06 |
| SS04631 | 0 | 8.71E-05 | 1.23E-06 |
| SS02128 | 0.013169538 | 0.018664559 | 1.24E-06 |
| SS13837 | 0.013169538 | 0.018664559 | 1.24E-06 |
| SS02685 | 0.01053591 | 0.000647019 | 1.25E-06 |
| SS04526 | 0.01053591 | 0.000647019 | 1.25E-06 |
| SS04082 | 3.55E-05 | 0.000463345 | 1.25E-06 |
| SS09078 | 0.009990041 | 0.005178552 | 1.26E-06 |
| SS00448 | 0.015060207 | 0.017652096 | 1.26E-06 |
| SS04286 | 0.015060207 | 0.017652096 | 1.26E-06 |
| SS05967 | 0.015060207 | 0.017652096 | 1.26E-06 |
| SS09624 | 0.015060207 | 0.017652096 | 1.26E-06 |
| SS02111 | 0.013066468 | 0.010382532 | 1.26E-06 |
| SS04571 | 0.013066468 | 0.010382532 | 1.26E-06 |
| SS05526 | 0 | 8.41E-05 | 1.28E-06 |
| SS07020 | 5.45E-06 | 8.28E-05 | 1.30E-06 |
| SS12945 | 5.45E-06 | 8.28E-05 | 1.30E-06 |
| SS05792 | 0 | 9.53E-05 | 1.30E-06 |
| SS14946 | 0 | 7.88E-05 | 1.30E-06 |
| SS08130 | 0.051927737 | 0.001864878 | 1.31E-06 |
| SS08153 | 0.051927737 | 0.001864878 | 1.31E-06 |
| SS12624 | 0.051927737 | 0.001864878 | 1.31E-06 |
| SS07555 | 0.012787518 | 0.010271977 | 1.31E-06 |
| SS11255 | 0.012787518 | 0.010271977 | 1.31E-06 |
| SS13945 | 0.029217392 | 0.002485376 | 1.32E-06 |
| SS07481 | 2.68E-06 | 0.000179851 | 1.32E-06 |
| SS00530 | 0 | 0.000424221 | 1.33E-06 |
| SS15253 | 0 | 2.49E-05 | 1.34E-06 |
| SS03384 | 0 | 4.28E-05 | 1.35E-06 |
| SS10530 | 0.000373005 | 0.000205975 | 1.36E-06 |
| SS00931 | 0.000853173 | 0.001656037 | 1.37E-06 |
| SS03463 | 0.000853173 | 0.001656037 | 1.37E-06 |
| SS01993 | 8.98E-05 | 0.000385737 | 1.37E-06 |
| SS09954 | 0.006630202 | 0.000834747 | 1.39E-06 |
| SS10016 | 0.006630202 | 0.000834747 | 1.39E-06 |
| SS01743 | 1.66E-05 | 0.000261661 | 1.40E-06 |
| SS10690 | 0.017184349 | 0.006956593 | 1.40E-06 |
| SS09247 | 0.017184349 | 0.006964823 | 1.40E-06 |
| SS12267 | 0.001271725 | 0.000911134 | 1.41E-06 |
| SS03547 | 0.006754823 | 0.003957629 | 1.42E-06 |
| SS03623 | 0.006754823 | 0.003957629 | 1.42E-06 |
| SS05220 | 0.006754823 | 0.003957629 | 1.42E-06 |
| SS08066 | 0.006754823 | 0.003957629 | 1.42E-06 |
| SS10836 | 0.006754823 | 0.003957629 | 1.42E-06 |
| SS14233 | 0.006754823 | 0.003957629 | 1.42E-06 |
| SS15498 | 0.006754823 | 0.003957629 | 1.42E-06 |

| | | | |
|---|---|---|---|
| SS03039 | 0.002105549 | 9.14E-05 | 1.42E-06 |
| SS09466 | 0.002105549 | 9.14E-05 | 1.42E-06 |
| SS02188 | 0.000870677 | 0.002211354 | 1.43E-06 |
| SS01799 | 0.033425312 | 0.043166838 | 1.43E-06 |
| SS04991 | 0.033425312 | 0.043166838 | 1.43E-06 |
| SS06146 | 0.033425312 | 0.043166838 | 1.43E-06 |
| SS10645 | 0.033425312 | 0.043166838 | 1.43E-06 |
| SS10976 | 0.033425312 | 0.043166838 | 1.43E-06 |
| SS03683 | 0 | 6.71E-05 | 1.44E-06 |
| SS07094 | 0 | 6.71E-05 | 1.44E-06 |
| SS11544 | 0.001308303 | 0.002374379 | 1.44E-06 |
| SS08357 | 1.37E-05 | 0.00094846 | 1.45E-06 |
| SS10469 | 0.01849576 | 0.023316017 | 1.46E-06 |
| SS04833 | 0.005242537 | 5.30E-05 | 1.46E-06 |
| SS10374 | 0.005242537 | 5.30E-05 | 1.46E-06 |
| SS05271 | 0.011875067 | 0.005733457 | 1.47E-06 |
| SS09352 | 0.002291351 | 0.000541778 | 1.47E-06 |
| SS04561 | 0.003017253 | 0.007044636 | 1.47E-06 |
| SS01442 | 0.001539332 | 0.000995738 | 1.47E-06 |
| SS12454 | 0 | 0.00044282 | 1.47E-06 |
| SS01088 | 0.008625982 | 0.000157791 | 1.48E-06 |
| SS02722 | 2.76E-06 | 0.001491163 | 1.49E-06 |
| SS03689 | 0.008198013 | 0.022155318 | 1.49E-06 |
| SS02503 | 0 | 0.000268881 | 1.51E-06 |
| SS00088 | 0.000814233 | 1.59E-05 | 1.52E-06 |
| SS04888 | 0.000814233 | 1.59E-05 | 1.52E-06 |
| SS13500 | 0.000814233 | 1.59E-05 | 1.52E-06 |
| SS03894 | 0 | 0.000127455 | 1.52E-06 |
| SS05860 | 0 | 5.81E-05 | 1.54E-06 |
| SS05876 | 0 | 5.81E-05 | 1.54E-06 |
| SS05945 | 0 | 5.81E-05 | 1.54E-06 |
| SS00573 | 0.000356901 | 8.89E-05 | 1.54E-06 |
| SS06310 | 0.000356901 | 8.89E-05 | 1.54E-06 |
| SS06412 | 0.000356901 | 8.89E-05 | 1.54E-06 |
| SS12132 | 0.000356901 | 8.89E-05 | 1.54E-06 |
| SS05274 | 0.008907651 | 0.001499285 | 1.55E-06 |
| SS12207 | 0.000503923 | 1.36E-05 | 1.57E-06 |
| SS09492 | 0 | 6.05E-05 | 1.57E-06 |
| SS01986 | 0.004667867 | 6.26E-05 | 1.58E-06 |
| SS01987 | 0.004667867 | 6.26E-05 | 1.58E-06 |
| SS13261 | 0 | 6.88E-05 | 1.60E-06 |
| SS04125 | 0.01241316 | 0.010338738 | 1.61E-06 |
| SS06695 | 0.01241316 | 0.010338738 | 1.61E-06 |
| SS05754 | 0 | 4.10E-05 | 1.62E-06 |
| SS08498 | 0 | 0.000109568 | 1.63E-06 |
| SS14864 | 8.13E-06 | 0.000172259 | 1.64E-06 |
| SS06642 | 0 | 0.000146554 | 1.65E-06 |

| | | | |
|---|---|---|---|
| SS00952 | 0.024895391 | 0.036118952 | 1.67E-06 |
| SS03611 | 0.024895391 | 0.036118952 | 1.67E-06 |
| SS05916 | 0.024895391 | 0.036118952 | 1.67E-06 |
| SS14958 | 0.000794883 | 0.001698394 | 1.70E-06 |
| SS09614 | 2.68E-06 | 6.47E-05 | 1.70E-06 |
| SS02655 | 0.002750048 | 0.004764301 | 1.73E-06 |
| SS14830 | 0.002750048 | 0.004764301 | 1.73E-06 |
| SS01432 | 0 | 5.32E-05 | 1.74E-06 |
| SS12949 | 0 | 5.32E-05 | 1.74E-06 |
| SS05840 | 0.002838539 | 0.004130615 | 1.76E-06 |
| SS00830 | 0.000223218 | 2.46E-05 | 1.76E-06 |
| SS03908 | 0 | 0.000218183 | 1.78E-06 |
| SS11887 | 0.023012441 | 0.028613319 | 1.80E-06 |
| SS01794 | 0.020571865 | 0.025893201 | 1.81E-06 |
| SS05242 | 0.020571865 | 0.025893201 | 1.81E-06 |
| SS06137 | 0.020571865 | 0.025893201 | 1.81E-06 |
| SS06168 | 0.020571865 | 0.025893201 | 1.81E-06 |
| SS05879 | 0.003941965 | 0.000288435 | 1.82E-06 |
| SS05948 | 0.003941965 | 0.000288435 | 1.82E-06 |
| SS11206 | 0.003941965 | 0.000288435 | 1.82E-06 |
| SS00831 | 0.000503281 | 0.001067975 | 1.82E-06 |
| SS13345 | 0.002184267 | 5.42E-07 | 1.84E-06 |
| SS10218 | 0 | 0.000114803 | 1.84E-06 |
| SS13872 | 0 | 3.16E-05 | 1.84E-06 |
| SS05581 | 0 | 5.88E-05 | 1.86E-06 |
| SS05664 | 0 | 0.000835356 | 1.86E-06 |
| SS06403 | 0.007429214 | 0.002336113 | 1.90E-06 |
| SS08909 | 0.008349762 | 0.00011008 | 1.91E-06 |
| SS09642 | 0 | 4.42E-05 | 1.93E-06 |
| SS10275 | 0 | 9.86E-05 | 1.94E-06 |
| SS04677 | 0.042331863 | 0.037162748 | 1.95E-06 |
| SS04521 | 8.13E-06 | 0.000299701 | 1.98E-06 |
| SS13653 | 8.13E-06 | 0.000299701 | 1.98E-06 |
| SS02199 | 0.009701846 | 0.00310764 | 2.00E-06 |
| SS07710 | 0.009701846 | 0.00310764 | 2.00E-06 |
| SS09404 | 3.80E-05 | 0.00024485 | 2.01E-06 |
| SS00453 | 3.27E-05 | 9.75E-07 | 2.06E-06 |
| SS15305 | 0.005771339 | 0.007723158 | 2.07E-06 |
| SS01218 | 0.007844324 | 0.011038612 | 2.07E-06 |
| SS03077 | 0.007844324 | 0.011038612 | 2.07E-06 |
| SS05387 | 0.007844324 | 0.011038612 | 2.07E-06 |
| SS14493 | 0.007844324 | 0.011038612 | 2.07E-06 |
| SS03395 | 0 | 4.42E-05 | 2.10E-06 |
| SS09349 | 0.042350891 | 0.037204262 | 2.11E-06 |
| SS08070 | 0 | 0.000131915 | 2.14E-06 |
| SS05038 | 0.010337456 | 0.001221837 | 2.14E-06 |
| SS02174 | 0 | 0.000208557 | 2.17E-06 |

| | | | |
|---|---|---|---|
| SS12334 | 0.000828054 | 1.29E-05 | 2.18E-06 |
| SS02030 | 0 | 9.33E-05 | 2.20E-06 |
| SS03974 | 0.003745071 | 2.86E-05 | 2.20E-06 |
| SS00588 | 0 | 0.000184702 | 2.23E-06 |
| SS02294 | 0.007126635 | 0.005291164 | 2.23E-06 |
| SS02386 | 0.007126635 | 0.005291164 | 2.23E-06 |
| SS02530 | 0.007126635 | 0.005291164 | 2.23E-06 |
| SS09008 | 0.003633422 | 5.99E-05 | 2.25E-06 |
| SS03194 | 0 | 0.000189164 | 2.29E-06 |
| SS10231 | 0.000310309 | 9.08E-05 | 2.29E-06 |
| SS11737 | 0 | 0.000119108 | 2.32E-06 |
| SS14147 | 0 | 0.000119108 | 2.32E-06 |
| SS11094 | 0 | 0.000206924 | 2.35E-06 |
| SS10723 | 0.00832244 | 0.000240463 | 2.36E-06 |
| SS00388 | 0.001848838 | 0.00020309 | 2.39E-06 |
| SS10303 | 0.00871523 | 0.005835903 | 2.41E-06 |
| SS01268 | 0 | 0.000107521 | 2.41E-06 |
| SS13686 | 0 | 0.000136036 | 2.42E-06 |
| SS03052 | 0.000231591 | 0.000802694 | 2.44E-06 |
| SS03871 | 0 | 2.72E-05 | 2.45E-06 |
| SS13712 | 0 | 7.32E-05 | 2.47E-06 |
| SS07036 | 0 | 5.72E-05 | 2.47E-06 |
| SS13070 | 0 | 5.72E-05 | 2.47E-06 |
| SS11721 | 0.020871451 | 0.017198685 | 2.49E-06 |
| SS14198 | 0.020871451 | 0.017198685 | 2.49E-06 |
| SS14264 | 0.020871451 | 0.017198685 | 2.49E-06 |
| SS14327 | 0.020871451 | 0.017198685 | 2.49E-06 |
| SS14347 | 0.020871451 | 0.017198685 | 2.49E-06 |
| SS14370 | 0.020871451 | 0.017198685 | 2.49E-06 |
| SS09772 | 2.16E-05 | 0.000298386 | 2.49E-06 |
| SS08189 | 5.99E-05 | 0.000247796 | 2.50E-06 |
| SS02717 | 0 | 0.000682287 | 2.52E-06 |
| SS05273 | 0.020324744 | 0.024724041 | 2.53E-06 |
| SS11591 | 0 | 9.00E-05 | 2.56E-06 |
| SS02980 | 0.006845448 | 0.000643523 | 2.60E-06 |
| SS02545 | 0.011615754 | 0.003439804 | 2.60E-06 |
| SS00897 | 2.68E-06 | 0.000890489 | 2.60E-06 |
| SS03682 | 0 | 4.38E-05 | 2.64E-06 |
| SS07093 | 0 | 4.38E-05 | 2.64E-06 |
| SS15165 | 0 | 6.73E-05 | 2.69E-06 |
| SS02922 | 8.29E-06 | 0.000572505 | 2.70E-06 |
| SS10217 | 0.001398962 | 0.003113683 | 2.72E-06 |
| SS10331 | 0.001398962 | 0.003113683 | 2.72E-06 |
| SS04225 | 0.001658471 | 0.001117324 | 2.74E-06 |
| SS03266 | 0.008146811 | 0.006228105 | 2.75E-06 |
| SS06954 | 0.007184639 | 0.009738431 | 2.78E-06 |
| SS03850 | 0 | 0.000151883 | 2.82E-06 |

| | | | |
|---|---|---|---|
| SS10057 | 0.006219862 | 0.0031347 | 2.87E-06 |
| SS04506 | 0.000133522 | 6.38E-05 | 2.91E-06 |
| SS10274 | 1.09E-05 | 9.95E-05 | 2.93E-06 |
| SS00206 | 5.53E-06 | 0.000269429 | 2.95E-06 |
| SS01233 | 0.002969664 | 0.006537127 | 2.96E-06 |
| SS06546 | 0.007757382 | 0.00986154 | 3.01E-06 |
| SS00244 | 0 | 8.88E-05 | 3.01E-06 |
| SS13022 | 0 | 8.13E-05 | 3.05E-06 |
| SS11041 | 0.010491372 | 0.013424845 | 3.14E-06 |
| SS06632 | 0 | 6.18E-05 | 3.17E-06 |
| SS11151 | 0.002220685 | 0.000542721 | 3.18E-06 |
| SS10575 | 0 | 5.64E-05 | 3.20E-06 |
| SS08499 | 0 | 6.22E-05 | 3.24E-06 |
| SS10762 | 0 | 6.22E-05 | 3.24E-06 |
| SS00399 | 0.018726606 | 0.010935321 | 3.25E-06 |
| SS07777 | 0.018726606 | 0.010935321 | 3.25E-06 |
| SS13018 | 0.018726606 | 0.010935321 | 3.25E-06 |
| SS02222 | 0 | 0.00025714 | 3.35E-06 |
| SS07622 | 0 | 0.000150083 | 3.37E-06 |
| SS13443 | 0.009719957 | 6.58E-05 | 3.44E-06 |
| SS06697 | 0.012825105 | 0.003012192 | 3.46E-06 |
| SS08237 | 0.010019897 | 0.007411112 | 3.48E-06 |
| SS08354 | 0 | 0.000217828 | 3.52E-06 |
| SS05982 | 0.00953147 | 0.000341812 | 3.56E-06 |
| SS04287 | 0.000944429 | 0.002518912 | 3.56E-06 |
| SS04311 | 0.000944429 | 0.002518912 | 3.56E-06 |
| SS04972 | 0.030012252 | 0.025641106 | 3.62E-06 |
| SS06131 | 0.030012252 | 0.025641106 | 3.62E-06 |
| SS06162 | 0.030012252 | 0.025641106 | 3.62E-06 |
| SS02038 | 0 | 0.000315971 | 3.64E-06 |
| SS00832 | 0.000142297 | 0.00079649 | 3.68E-06 |
| SS09284 | 0.000286154 | 0.000114384 | 3.69E-06 |
| SS12214 | 0.001032484 | 6.31E-05 | 3.70E-06 |
| SS09325 | 0.00590134 | 0.002917061 | 3.75E-06 |
| SS09131 | 0.000143776 | 0.001731908 | 3.75E-06 |
| SS13935 | 0.000643295 | 0.001285102 | 3.76E-06 |
| SS05576 | 1.63E-05 | 3.20E-07 | 3.77E-06 |
| SS03516 | 0.006193653 | 0.00358914 | 3.78E-06 |
| SS12457 | 0 | 0.000705421 | 3.81E-06 |
| SS02293 | 0.007126635 | 0.005369592 | 3.83E-06 |
| SS08355 | 0 | 0.000366658 | 3.83E-06 |
| SS06993 | 0.016364897 | 0.000419328 | 3.85E-06 |
| SS11108 | 0.016364897 | 0.000419328 | 3.85E-06 |
| SS03310 | 0 | 5.68E-05 | 3.85E-06 |
| SS12153 | 0 | 7.54E-05 | 3.85E-06 |
| SS03305 | 0.013684369 | 0.007488789 | 3.88E-06 |
| SS05138 | 4.36E-05 | 0.000184526 | 3.90E-06 |

| | | | |
|---|---|---|---|
| SS15140 | 4.36E-05 | 0.000184526 | 3.90E-06 |
| SS01458 | 0 | 9.64E-05 | 3.90E-06 |
| SS13057 | 0 | 9.64E-05 | 3.90E-06 |
| SS02075 | 0 | 4.26E-05 | 3.92E-06 |
| SS03464 | 0.014304976 | 0.017950292 | 4.01E-06 |
| SS02351 | 0.00666858 | 0.004546143 | 4.02E-06 |
| SS12156 | 0.001054518 | 0.000196358 | 4.04E-06 |
| SS00260 | 0 | 7.29E-05 | 4.06E-06 |
| SS09580 | 0.000335188 | 0.000115127 | 4.11E-06 |
| SS08895 | 0 | 0.000100319 | 4.15E-06 |
| SS10491 | 0.026384898 | 0.009586906 | 4.16E-06 |
| SS10825 | 0.026384898 | 0.009586906 | 4.16E-06 |
| SS03729 | 0.002689028 | 0.004767922 | 4.17E-06 |
| SS00966 | 0.003366939 | 0.000458481 | 4.19E-06 |
| SS07123 | 0.003366939 | 0.000458481 | 4.19E-06 |
| SS11434 | 0.005217659 | 0.00932683 | 4.21E-06 |
| SS12629 | 0.005217659 | 0.00932683 | 4.21E-06 |
| SS15504 | 0.000587769 | 0.001099184 | 4.23E-06 |
| SS06058 | 0.022767155 | 0.018612277 | 4.27E-06 |
| SS11893 | 0.022767155 | 0.018612277 | 4.27E-06 |
| SS05832 | 0.017705454 | 0.015140107 | 4.29E-06 |
| SS09059 | 0.017705454 | 0.015140107 | 4.29E-06 |
| SS09880 | 0.017705454 | 0.015140107 | 4.29E-06 |
| SS09943 | 0 | 0.000123663 | 4.36E-06 |
| SS14363 | 0.005509387 | 0.00808683 | 4.38E-06 |
| SS09398 | 0 | 2.41E-05 | 4.39E-06 |
| SS04531 | 0.028468677 | 0.023747857 | 4.45E-06 |
| SS04661 | 0.028468677 | 0.023747857 | 4.45E-06 |
| SS01563 | 0.008132669 | 0.005427153 | 4.46E-06 |
| SS12270 | 0.000100672 | 0.000248213 | 4.62E-06 |
| SS15337 | 0.003227166 | 4.89E-05 | 4.65E-06 |
| SS03898 | 0 | 0.000185202 | 4.67E-06 |
| SS04284 | 0.020221766 | 0.010329473 | 4.77E-06 |
| SS08777 | 0.020221766 | 0.010329473 | 4.77E-06 |
| SS10691 | 0.020221766 | 0.010329473 | 4.77E-06 |
| SS03065 | 0.000277781 | 0.000152886 | 4.85E-06 |
| SS00621 | 0 | 4.17E-05 | 4.88E-06 |
| SS01837 | 0.000114253 | 0.000315573 | 5.01E-06 |
| SS10547 | 0.000114253 | 0.000315573 | 5.01E-06 |
| SS15456 | 0 | 0.000148003 | 5.06E-06 |
| SS05439 | 0.005271901 | 0.00056332 | 5.08E-06 |
| SS14476 | 0.008160552 | 0.001001406 | 5.09E-06 |
| SS15335 | 0.008160552 | 0.001001406 | 5.09E-06 |
| SS04864 | 0 | 0.000248585 | 5.10E-06 |
| SS15255 | 0.000419355 | 1.14E-05 | 5.20E-06 |
| SS00561 | 0 | 0.000427171 | 5.21E-06 |
| SS06227 | 0 | 0.000427171 | 5.21E-06 |

| | | | |
|---|---|---|---|
| SS07938 | 0 | 0.000427171 | 5.21E-06 |
| SS07256 | 1.36E-05 | 0.000122467 | 5.24E-06 |
| SS09704 | 0 | 0.000177 | 5.26E-06 |
| SS04576 | 0 | 0.000111962 | 5.27E-06 |
| SS04613 | 0 | 0.000111962 | 5.27E-06 |
| SS07956 | 0 | 0.000389291 | 5.27E-06 |
| SS03050 | 0.00077918 | 0.000112844 | 5.35E-06 |
| SS03761 | 0.00077918 | 0.000112844 | 5.35E-06 |
| SS10854 | 0.00077918 | 0.000112844 | 5.35E-06 |
| SS02347 | 0 | 0.000250777 | 5.41E-06 |
| SS11119 | 0 | 0.000250777 | 5.41E-06 |
| SS05259 | 0 | 5.85E-05 | 5.42E-06 |
| SS12777 | 0 | 5.85E-05 | 5.42E-06 |
| SS09625 | 0.019702347 | 0.001436208 | 5.42E-06 |
| SS03324 | 0.000452285 | 0.001118489 | 5.45E-06 |
| SS04841 | 0.000452285 | 0.001118489 | 5.45E-06 |
| SS13875 | 0.000452285 | 0.001118489 | 5.45E-06 |
| SS01057 | 0.025587389 | 0.028771316 | 5.48E-06 |
| SS01318 | 0.025587389 | 0.028771316 | 5.48E-06 |
| SS03961 | 0.025587389 | 0.028771316 | 5.48E-06 |
| SS07652 | 0.025587389 | 0.028771316 | 5.48E-06 |
| SS14334 | 0.025587389 | 0.028771316 | 5.48E-06 |
| SS14350 | 0.025587389 | 0.028771316 | 5.48E-06 |
| SS15455 | 0.025587389 | 0.028771316 | 5.48E-06 |
| SS15486 | 0.025587389 | 0.028771316 | 5.48E-06 |
| SS01177 | 0.002930035 | 0.00013791 | 5.56E-06 |
| SS02970 | 0.002649571 | 0.000197643 | 5.61E-06 |
| SS08556 | 0.002649571 | 0.000197643 | 5.61E-06 |
| SS11433 | 0.006508849 | 0.009412744 | 5.63E-06 |
| SS11443 | 0.01170788 | 0.006015826 | 5.64E-06 |
| SS07199 | 0.003059635 | 0.000861215 | 5.70E-06 |
| SS07223 | 0.003059635 | 0.000861215 | 5.70E-06 |
| SS01588 | 0 | 7.95E-05 | 5.71E-06 |
| SS04183 | 0.000130758 | 5.07E-05 | 5.75E-06 |
| SS06955 | 0.007752301 | 0.010447677 | 5.76E-06 |
| SS03144 | 0.000681032 | 0.001363392 | 5.83E-06 |
| SS10556 | 0.000681032 | 0.001363392 | 5.83E-06 |
| SS05187 | 0.014642882 | 0.010696912 | 5.83E-06 |
| SS06491 | 0.014642882 | 0.010696912 | 5.83E-06 |
| SS01064 | 0.013327067 | 0.005330621 | 5.83E-06 |
| SS13282 | 0.043696632 | 0.004029587 | 5.84E-06 |
| SS13687 | 0 | 6.22E-05 | 5.85E-06 |
| SS03455 | 0 | 5.12E-05 | 5.90E-06 |
| SS08427 | 0.012711013 | 0.001487973 | 5.92E-06 |
| SS00122 | 0.033113444 | 0.044209899 | 5.92E-06 |
| SS02932 | 0 | 0.000201552 | 5.96E-06 |
| SS10803 | 0.007544269 | 0.002661192 | 5.98E-06 |

| | | | |
|---|---|---|---|
| SS03493 | 0 | 6.37E-05 | 5.98E-06 |
| SS07987 | 0 | 6.37E-05 | 5.98E-06 |
| SS07245 | 0.01315957 | 0.001235871 | 5.98E-06 |
| SS03708 | 7.90E-05 | 1.93E-05 | 6.03E-06 |
| SS00976 | 0.000245252 | 0.000118857 | 6.07E-06 |
| SS09994 | 0.01074791 | 0.000928906 | 6.10E-06 |
| SS13619 | 0.001026474 | 0.000638477 | 6.12E-06 |
| SS01711 | 0.028824877 | 0.034149803 | 6.12E-06 |
| SS03301 | 0.028824877 | 0.034149803 | 6.12E-06 |
| SS14169 | 0 | 0.000172996 | 6.21E-06 |
| SS06255 | 0.012197741 | 0.007568643 | 6.33E-06 |
| SS11568 | 0.012197741 | 0.007568643 | 6.33E-06 |
| SS11773 | 0.012197741 | 0.007568643 | 6.33E-06 |
| SS09703 | 0 | 0.000122886 | 6.34E-06 |
| SS01413 | 0 | 6.58E-05 | 6.36E-06 |
| SS06543 | 0.028712276 | 0.02440733 | 6.38E-06 |
| SS04283 | 0.021335022 | 0.018010267 | 6.39E-06 |
| SS04769 | 0.021335022 | 0.018010267 | 6.39E-06 |
| SS13103 | 0.021335022 | 0.018010267 | 6.39E-06 |
| SS05007 | 0 | 0.000139634 | 6.43E-06 |
| SS15529 | 0.00018508 | 4.59E-05 | 6.47E-06 |
| SS13787 | 0.000242487 | 0.000656226 | 6.47E-06 |
| SS14890 | 0.000242487 | 0.000656226 | 6.47E-06 |
| SS14039 | 6.53E-05 | 0.000171217 | 6.50E-06 |
| SS02965 | 0 | 5.59E-05 | 6.50E-06 |
| SS00004 | 0.008772489 | 0.005069834 | 6.51E-06 |
| SS01197 | 0.008772489 | 0.005069834 | 6.51E-06 |
| SS01261 | 0.008772489 | 0.005069834 | 6.51E-06 |
| SS00626 | 0 | 3.97E-05 | 6.56E-06 |
| SS07571 | 6.81E-05 | 0.000537321 | 6.57E-06 |
| SS01447 | 0.022213969 | 0.007910668 | 6.68E-06 |
| SS04447 | 0 | 7.64E-05 | 6.75E-06 |
| SS04138 | 0 | 0.000207169 | 6.76E-06 |
| SS00748 | 0.00832572 | 0.003701238 | 6.77E-06 |
| SS00779 | 0.00832572 | 0.003701238 | 6.77E-06 |
| SS00792 | 0.00832572 | 0.003701238 | 6.77E-06 |
| SS12038 | 0 | 4.13E-05 | 6.97E-06 |
| SS05523 | 0.003169999 | 7.22E-05 | 7.00E-06 |
| SS01220 | 0.013817524 | 0.017330811 | 7.00E-06 |
| SS14746 | 0 | 4.24E-05 | 7.09E-06 |
| SS12878 | 0 | 0.000236723 | 7.13E-06 |
| SS09709 | 0 | 9.32E-05 | 7.29E-06 |
| SS00978 | 0 | 6.67E-05 | 7.32E-06 |
| SS01543 | 2.18E-05 | 0.000369209 | 7.43E-06 |
| SS09833 | 0.001365148 | 0.000461549 | 7.48E-06 |
| SS00050 | 0.000496433 | 0.001176863 | 7.50E-06 |
| SS12459 | 0 | 0.000224247 | 7.53E-06 |

| | | | |
|---|---|---|---|
| SS11751 | 4.10E-05 | 0.000207474 | 7.57E-06 |
| SS14197 | 4.10E-05 | 0.000207474 | 7.57E-06 |
| SS14325 | 4.10E-05 | 0.000207474 | 7.57E-06 |
| SS14345 | 4.10E-05 | 0.000207474 | 7.57E-06 |
| SS14369 | 4.10E-05 | 0.000207474 | 7.57E-06 |
| SS00580 | 0 | 0.000288042 | 7.63E-06 |
| SS06586 | 0 | 0.000288042 | 7.63E-06 |
| SS01801 | 0.012317018 | 0.006923505 | 7.67E-06 |
| SS06147 | 0.012317018 | 0.006923505 | 7.67E-06 |
| SS10978 | 0.012317018 | 0.006923505 | 7.67E-06 |
| SS01746 | 0.007515904 | 0.001851993 | 7.71E-06 |
| SS06925 | 0.007515904 | 0.001851993 | 7.71E-06 |
| SS15089 | 0.007515904 | 0.001851993 | 7.71E-06 |
| SS10304 | 0.006094071 | 0.000681446 | 7.83E-06 |
| SS11191 | 1.35E-05 | 0.000186971 | 7.83E-06 |
| SS09656 | 0.007385822 | 0.005937177 | 7.85E-06 |
| SS10745 | 0.007385822 | 0.005937177 | 7.85E-06 |
| SS05310 | 0.005397738 | 0.003244011 | 7.91E-06 |
| SS09018 | 0 | 7.14E-05 | 7.93E-06 |
| SS12124 | 0.000632318 | 0.000208224 | 8.01E-06 |
| SS11248 | 0 | 1.96E-05 | 8.05E-06 |
| SS07040 | 5.37E-06 | 0.000146934 | 8.06E-06 |
| SS13074 | 5.37E-06 | 0.000146934 | 8.06E-06 |
| SS02883 | 0 | 0.000144988 | 8.09E-06 |
| SS00956 | 0 | 2.39E-05 | 8.17E-06 |
| SS00305 | 0 | 7.48E-05 | 8.42E-06 |
| SS02105 | 0.005185451 | 2.42E-05 | 8.45E-06 |
| SS13332 | 0.005185451 | 2.42E-05 | 8.45E-06 |
| SS14789 | 0.007243009 | 0.000785329 | 8.47E-06 |
| SS15053 | 0.007243009 | 0.000785329 | 8.47E-06 |
| SS04826 | 0.013212642 | 0.001075898 | 8.63E-06 |
| SS11238 | 0.013212642 | 0.001075898 | 8.63E-06 |
| SS11714 | 0.013212642 | 0.001075898 | 8.63E-06 |
| SS11760 | 0.013212642 | 0.001075898 | 8.63E-06 |
| SS14236 | 0.013212642 | 0.001075898 | 8.63E-06 |
| SS15460 | 0.013212642 | 0.001075898 | 8.63E-06 |
| SS02218 | 0 | 0.000128734 | 8.63E-06 |
| SS02714 | 0 | 0.000433478 | 8.79E-06 |
| SS13142 | 0 | 8.18E-05 | 8.89E-06 |
| SS15314 | 0 | 8.18E-05 | 8.89E-06 |
| SS04038 | 0 | 6.17E-05 | 8.93E-06 |
| SS07983 | 0 | 6.15E-05 | 9.00E-06 |
| SS12462 | 0 | 7.17E-05 | 9.02E-06 |
| SS11051 | 0 | 3.23E-05 | 9.04E-06 |
| SS09409 | 0 | 4.96E-05 | 9.07E-06 |
| SS14562 | 0 | 3.07E-05 | 9.07E-06 |
| SS13531 | 0 | 1.43E-05 | 9.10E-06 |

| | | | |
|---|---|---|---|
| SS07210 | 0.000215246 | 0.000584684 | 9.12E-06 |
| SS02345 | 0 | 2.78E-05 | 9.17E-06 |
| SS03051 | 5.45E-05 | 0.000150112 | 9.25E-06 |
| SS08483 | 5.45E-05 | 0.000150112 | 9.25E-06 |
| SS09471 | 5.45E-05 | 0.000150112 | 9.25E-06 |
| SS09514 | 5.45E-05 | 0.000150112 | 9.25E-06 |
| SS08127 | 0.01341651 | 0.00154154 | 9.34E-06 |
| SS05950 | 0.015206255 | 0.008945293 | 9.67E-06 |
| SS14953 | 2.46E-05 | 0.000698024 | 9.74E-06 |
| SS10278 | 1.91E-05 | 9.60E-05 | 9.82E-06 |
| SS02920 | 0 | 0.000705328 | 9.85E-06 |
| SS04861 | 0 | 8.99E-05 | 9.87E-06 |
| SS08114 | 0 | 0.00022953 | 9.89E-06 |
| SS14130 | 0 | 0.000225119 | 1.00E-05 |
| SS06666 | 0.013719765 | 0.000484268 | 1.01E-05 |
| SS11884 | 0.010576089 | 0.001268584 | 1.01E-05 |
| SS14240 | 0.010576089 | 0.001268584 | 1.01E-05 |
| SS00447 | 0.00500434 | 0.000847989 | 1.02E-05 |
| SS00324 | 0.002756575 | 6.10E-05 | 1.02E-05 |
| SS14933 | 0.000512216 | 0.001304584 | 1.03E-05 |
| SS02375 | 0.015452871 | 0.018357256 | 1.03E-05 |
| SS07591 | 0.015452871 | 0.018357256 | 1.03E-05 |
| SS12533 | 0.006187448 | 4.91E-05 | 1.03E-05 |
| SS11922 | 0 | 8.95E-05 | 1.03E-05 |
| SS13272 | 0 | 8.95E-05 | 1.03E-05 |
| SS03564 | 0 | 0.000105137 | 1.04E-05 |
| SS00423 | 0.004462118 | 0.00196862 | 1.04E-05 |
| SS01412 | 0 | 4.06E-05 | 1.05E-05 |
| SS08811 | 0.007171941 | 0.000441254 | 1.05E-05 |
| SS07006 | 0.00186859 | 0.003264587 | 1.05E-05 |
| SS07480 | 0.00446837 | 0.002565495 | 1.05E-05 |
| SS09091 | 0.014050904 | 0.007818545 | 1.06E-05 |
| SS10962 | 0.014050904 | 0.007818545 | 1.06E-05 |
| SS11729 | 0 | 0.000251215 | 1.06E-05 |
| SS09706 | 0 | 0.000243356 | 1.07E-05 |
| SS06711 | 0.012928782 | 0.001424778 | 1.08E-05 |
| SS00344 | 0 | 4.57E-05 | 1.08E-05 |
| SS12464 | 0.002711188 | 0.000928769 | 1.08E-05 |
| SS00716 | 0 | 4.13E-05 | 1.09E-05 |
| SS04935 | 0 | 4.13E-05 | 1.09E-05 |
| SS07880 | 0 | 4.13E-05 | 1.09E-05 |
| SS08572 | 0 | 4.13E-05 | 1.09E-05 |
| SS13328 | 0.003987558 | 6.49E-05 | 1.11E-05 |
| SS06802 | 0.007338394 | 0.000156653 | 1.11E-05 |
| SS12092 | 0.007338394 | 0.000156653 | 1.11E-05 |
| SS12129 | 0.007338394 | 0.000156653 | 1.11E-05 |
| SS01634 | 0.0101212 | 0.001651541 | 1.12E-05 |

| | | | |
|---|---|---|---|
| SS01650 | 0.0101212 | 0.001651541 | 1.12E-05 |
| SS00114 | 0.006385145 | 0.000331382 | 1.14E-05 |
| SS01134 | 0.002519214 | 4.13E-06 | 1.14E-05 |
| SS07999 | 0 | 3.04E-05 | 1.14E-05 |
| SS10236 | 0.000163287 | 5.28E-05 | 1.15E-05 |
| SS15249 | 0 | 3.61E-05 | 1.15E-05 |
| SS00138 | 0.002425114 | 0.001171723 | 1.15E-05 |
| SS09533 | 0.01229854 | 0.000621932 | 1.17E-05 |
| SS09427 | 0 | 0.000165351 | 1.17E-05 |
| SS05287 | 0.006159002 | 0.00824521 | 1.17E-05 |
| SS14794 | 0.006159002 | 0.00824521 | 1.17E-05 |
| SS10480 | 0.019898576 | 0.001090046 | 1.18E-05 |
| SS03895 | 0 | 4.99E-05 | 1.18E-05 |
| SS08620 | 0.002638193 | 0.001841276 | 1.19E-05 |
| SS00553 | 0.000395038 | 0.000762912 | 1.19E-05 |
| SS10350 | 0 | 2.95E-05 | 1.20E-05 |
| SS02942 | 0 | 8.34E-05 | 1.20E-05 |
| SS05828 | 0.012505252 | 0.000381659 | 1.20E-05 |
| SS13637 | 0 | 4.12E-05 | 1.21E-05 |
| SS03152 | 0.001592209 | 0.002503823 | 1.21E-05 |
| SS12036 | 0.001592209 | 0.002503823 | 1.21E-05 |
| SS01375 | 0.001484207 | 0.002037933 | 1.22E-05 |
| SS02062 | 0.001484207 | 0.002037933 | 1.22E-05 |
| SS02505 | 8.13E-06 | 0.00010617 | 1.22E-05 |
| SS02799 | 0 | 7.77E-05 | 1.23E-05 |
| SS02825 | 8.21E-06 | 9.47E-05 | 1.23E-05 |
| SS08482 | 8.21E-06 | 9.47E-05 | 1.23E-05 |
| SS00967 | 0.006458335 | 0.001699499 | 1.24E-05 |
| SS06209 | 0.006458335 | 0.001699499 | 1.24E-05 |
| SS07124 | 0.006458335 | 0.001699499 | 1.24E-05 |
| SS02406 | 0 | 0.000244635 | 1.25E-05 |
| SS03637 | 0 | 0.000244635 | 1.25E-05 |
| SS01082 | 0 | 2.31E-05 | 1.27E-05 |
| SS04048 | 0 | 0.000141336 | 1.27E-05 |
| SS09281 | 0 | 2.42E-05 | 1.27E-05 |
| SS11554 | 0.00954926 | 0.000119879 | 1.28E-05 |
| SS14219 | 0.00954926 | 0.000119879 | 1.28E-05 |
| SS10800 | 0.026410418 | 0.022003481 | 1.29E-05 |
| SS13267 | 0.026410418 | 0.022003481 | 1.29E-05 |
| SS13251 | 0.011108171 | 0.006572692 | 1.30E-05 |
| SS08442 | 0.002669483 | 0.002095393 | 1.31E-05 |
| SS05649 | 3.27E-05 | 1.00E-05 | 1.32E-05 |
| SS01919 | 0.001787107 | 0.000190354 | 1.33E-05 |
| SS03720 | 0.001787107 | 0.000190354 | 1.33E-05 |
| SS13141 | 0 | 0.000129292 | 1.33E-05 |
| SS15313 | 0 | 0.000129292 | 1.33E-05 |
| SS04334 | 0.000808784 | 0.001448513 | 1.34E-05 |

| | | | |
|---|---|---|---|
| SS00583 | 0 | 3.46E-05 | 1.34E-05 |
| SS02718 | 0 | 0.000164643 | 1.35E-05 |
| SS02921 | 0 | 0.000620446 | 1.35E-05 |
| SS03924 | 0.000136287 | 3.96E-05 | 1.35E-05 |
| SS14062 | 0 | 4.82E-05 | 1.36E-05 |
| SS04673 | 0.004744464 | 2.36E-06 | 1.37E-05 |
| SS04674 | 0.004744464 | 2.36E-06 | 1.37E-05 |
| SS09616 | 0 | 5.18E-05 | 1.39E-05 |
| SS08323 | 0.015784458 | 0.000674868 | 1.40E-05 |
| SS09392 | 0.000199062 | 0.000518906 | 1.41E-05 |
| SS10934 | 0.000199062 | 0.000518906 | 1.41E-05 |
| SS01214 | 0.000651668 | 0.001288603 | 1.41E-05 |
| SS00742 | 0.017951772 | 0.00154856 | 1.42E-05 |
| SS04394 | 0 | 5.71E-05 | 1.43E-05 |
| SS05006 | 0 | 6.63E-05 | 1.46E-05 |
| SS13858 | 0.015022299 | 0.001658939 | 1.46E-05 |
| SS03652 | 0.004215582 | 0.006612698 | 1.46E-05 |
| SS05620 | 0.004215582 | 0.006612698 | 1.46E-05 |
| SS07759 | 0.004215582 | 0.006612698 | 1.46E-05 |
| SS12939 | 0.004215582 | 0.006612698 | 1.46E-05 |
| SS00927 | 0.008011729 | 0.009896729 | 1.47E-05 |
| SS10821 | 0.008011729 | 0.009896729 | 1.47E-05 |
| SS14216 | 0.008011729 | 0.009896729 | 1.47E-05 |
| SS14275 | 0.008011729 | 0.009896729 | 1.47E-05 |
| SS03681 | 0 | 0.000170456 | 1.47E-05 |
| SS07092 | 0 | 0.000170456 | 1.47E-05 |
| SS12216 | 0.000345924 | 5.09E-06 | 1.47E-05 |
| SS03440 | 5.75E-05 | 0.000308783 | 1.48E-05 |
| SS00251 | 0.009479672 | 0.00040665 | 1.49E-05 |
| SS05182 | 0.009479672 | 0.00040665 | 1.49E-05 |
| SS03902 | 0 | 7.16E-05 | 1.49E-05 |
| SS10381 | 0.000273135 | 0.001061234 | 1.50E-05 |
| SS00140 | 0 | 7.85E-05 | 1.51E-05 |
| SS02091 | 0.002502147 | 0.001738287 | 1.51E-05 |
| SS04794 | 0.007142819 | 0.00156033 | 1.51E-05 |
| SS09810 | 0 | 0.000164275 | 1.51E-05 |
| SS13577 | 0 | 0.000164275 | 1.51E-05 |
| SS09705 | 2.68E-06 | 0.000242538 | 1.54E-05 |
| SS00029 | 0 | 5.97E-05 | 1.54E-05 |
| SS05923 | 5.45E-06 | 6.93E-08 | 1.55E-05 |
| SS04391 | 0.000174183 | 0.000332285 | 1.55E-05 |
| SS07792 | 0.016551583 | 0.000181826 | 1.55E-05 |
| SS12046 | 0.016551583 | 0.000181826 | 1.55E-05 |
| SS05767 | 0 | 6.03E-05 | 1.58E-05 |
| SS08184 | 1.08E-05 | 0.000162423 | 1.58E-05 |
| SS01936 | 0 | 0.00010876 | 1.59E-05 |
| SS08285 | 0.017694466 | 0.000499576 | 1.60E-05 |

| | | | |
|---|---|---|---|
| SS00575 | 0.016530478 | 0.020710457 | 1.63E-05 |
| SS15385 | 0.016530478 | 0.020710457 | 1.63E-05 |
| SS09785 | 0.000122786 | 0.000531308 | 1.64E-05 |
| SS15178 | 0.000122786 | 0.000531308 | 1.64E-05 |
| SS06553 | 0 | 0.000240561 | 1.65E-05 |
| SS04167 | 0.00019866 | 5.06E-05 | 1.66E-05 |
| SS09055 | 0.005757483 | 0.002306143 | 1.66E-05 |
| SS09088 | 0.005757483 | 0.002306143 | 1.66E-05 |
| SS13798 | 0.014163517 | 0.017888903 | 1.72E-05 |
| SS06020 | 0.011779912 | 0.001684938 | 1.77E-05 |
| SS11104 | 0.011779912 | 0.001684938 | 1.77E-05 |
| SS03433 | 0 | 0.000652666 | 1.79E-05 |
| SS05732 | 0 | 2.86E-05 | 1.82E-05 |
| SS08522 | 0.009753208 | 3.22E-05 | 1.83E-05 |
| SS08365 | 0 | 8.36E-05 | 1.84E-05 |
| SS05291 | 0.00034332 | 0.002669156 | 1.85E-05 |
| SS05292 | 0.00034332 | 0.002669156 | 1.85E-05 |
| SS05298 | 0.00034332 | 0.002669156 | 1.85E-05 |
| SS02398 | 0.030427925 | 0.03561168 | 1.86E-05 |
| SS03524 | 0.030427925 | 0.03561168 | 1.86E-05 |
| SS14880 | 0.030427925 | 0.03561168 | 1.86E-05 |
| SS09497 | 0 | 8.31E-05 | 1.88E-05 |
| SS11495 | 7.07E-05 | 0.000272964 | 1.91E-05 |
| SS01379 | 0.003925379 | 0.001070696 | 1.91E-05 |
| SS10075 | 0.000978323 | 0.00157613 | 1.92E-05 |
| SS10190 | 0.000978323 | 0.00157613 | 1.92E-05 |
| SS11291 | 0 | 4.71E-05 | 1.93E-05 |
| SS13505 | 0.007877243 | 0.001674635 | 1.95E-05 |
| SS12198 | 0 | 1.71E-05 | 1.96E-05 |
| SS00410 | 0 | 0.000247294 | 1.96E-05 |
| SS13473 | 0.012014233 | 0.000342287 | 1.99E-05 |
| SS03428 | 0.008388507 | 0.004211823 | 2.02E-05 |
| SS10725 | 0.008388507 | 0.004211823 | 2.02E-05 |
| SS11617 | 3.55E-05 | 0.000655746 | 2.03E-05 |
| SS13641 | 0 | 0.000112184 | 2.04E-05 |
| SS00231 | 0.00043001 | 0.001025714 | 2.04E-05 |
| SS09428 | 0 | 0.000133596 | 2.06E-05 |
| SS06456 | 0.004206968 | 0.003225097 | 2.07E-05 |
| SS09814 | 0.004206968 | 0.003225097 | 2.07E-05 |
| SS08853 | 0.010712697 | 0.000969926 | 2.08E-05 |
| SS04488 | 0.072459962 | 0.083241146 | 2.08E-05 |
| SS04705 | 0.072459962 | 0.083241146 | 2.08E-05 |
| SS11045 | 0.072459962 | 0.083241146 | 2.08E-05 |
| SS04688 | 0.003484954 | 0.001396434 | 2.09E-05 |
| SS04461 | 0.002505875 | 5.90E-05 | 2.10E-05 |
| SS02114 | 0.019834505 | 0.02450528 | 2.11E-05 |
| SS12869 | 0.019834505 | 0.02450528 | 2.11E-05 |

| | | | |
|---|---|---|---|
| SS02250 | 0.000310631 | 6.86E-05 | 2.11E-05 |
| SS04066 | 0.000310631 | 6.86E-05 | 2.11E-05 |
| SS02233 | 0.007443551 | 0.009466287 | 2.13E-05 |
| SS06518 | 0.007443551 | 0.009466287 | 2.13E-05 |
| SS11537 | 5.45E-06 | 6.77E-08 | 2.13E-05 |
| SS07618 | 0.012991787 | 0.00119062 | 2.14E-05 |
| SS06542 | 0.036819964 | 0.031921027 | 2.17E-05 |
| SS03655 | 0.005265936 | 0.006819192 | 2.17E-05 |
| SS15279 | 0.005265936 | 0.006819192 | 2.17E-05 |
| SS11449 | 0 | 2.57E-05 | 2.18E-05 |
| SS02394 | 0.04782003 | 0.034367089 | 2.19E-05 |
| SS05001 | 0.04782003 | 0.034367089 | 2.19E-05 |
| SS05975 | 0.04782003 | 0.034367089 | 2.19E-05 |
| SS06158 | 0.04782003 | 0.034367089 | 2.19E-05 |
| SS06183 | 0.04782003 | 0.034367089 | 2.19E-05 |
| SS10785 | 0.04782003 | 0.034367089 | 2.19E-05 |
| SS09113 | 0.002366342 | 0.000151015 | 2.20E-05 |
| SS07263 | 0.014786671 | 0.017457835 | 2.22E-05 |
| SS09144 | 0.014786671 | 0.017457835 | 2.22E-05 |
| SS11864 | 0.014786671 | 0.017457835 | 2.22E-05 |
| SS06715 | 0.001534606 | 0.002566922 | 2.25E-05 |
| SS07595 | 0 | 5.09E-05 | 2.26E-05 |
| SS02912 | 0 | 0.000361386 | 2.26E-05 |
| SS03551 | 0 | 0.000361386 | 2.26E-05 |
| SS15220 | 0.000393639 | 0.002175474 | 2.26E-05 |
| SS14073 | 0 | 3.32E-05 | 2.26E-05 |
| SS01910 | 0 | 4.45E-05 | 2.27E-05 |
| SS04046 | 0 | 0.00013146 | 2.27E-05 |
| SS03375 | 0.000239482 | 8.49E-05 | 2.29E-05 |
| SS04894 | 0.000239482 | 8.49E-05 | 2.29E-05 |
| SS13902 | 0.000239482 | 8.49E-05 | 2.29E-05 |
| SS01267 | 0.005034506 | 0.001161101 | 2.30E-05 |
| SS12551 | 0.005034506 | 0.001161101 | 2.30E-05 |
| SS08238 | 0.008719933 | 0.013844458 | 2.30E-05 |
| SS01482 | 0 | 0.000155618 | 2.37E-05 |
| SS07253 | 0.006603637 | 0.000166411 | 2.40E-05 |
| SS10647 | 0.015031727 | 0.003987161 | 2.41E-05 |
| SS02715 | 0 | 0.000296623 | 2.41E-05 |
| SS01107 | 0.019714288 | 0.022267782 | 2.41E-05 |
| SS03004 | 0.019714288 | 0.022267782 | 2.41E-05 |
| SS14468 | 0.019714288 | 0.022267782 | 2.41E-05 |
| SS02510 | 0 | 0.000139211 | 2.41E-05 |
| SS07293 | 0.00329383 | 0.000880891 | 2.43E-05 |
| SS11225 | 0.00329383 | 0.000880891 | 2.43E-05 |
| SS11329 | 7.90E-05 | 1.41E-05 | 2.44E-05 |
| SS05023 | 0 | 0.000185364 | 2.45E-05 |
| SS12560 | 0 | 9.47E-05 | 2.47E-05 |

| | | | |
|---|---|---|---|
| SS03999 | 0.000631756 | 0.001125934 | 2.47E-05 |
| SS11285 | 0.014207963 | 0.018043704 | 2.48E-05 |
| SS06066 | 0.021652317 | 0.01210182 | 2.49E-05 |
| SS13556 | 0.021652317 | 0.01210182 | 2.49E-05 |
| SS12557 | 0 | 4.80E-05 | 2.54E-05 |
| SS09354 | 0 | 4.87E-05 | 2.54E-05 |
| SS12475 | 0.007144861 | 0.00048412 | 2.54E-05 |
| SS12525 | 0.007144861 | 0.00048412 | 2.54E-05 |
| SS12553 | 0.007144861 | 0.00048412 | 2.54E-05 |
| SS03034 | 0.031819041 | 0.010823909 | 2.55E-05 |
| SS09460 | 0.031819041 | 0.010823909 | 2.55E-05 |
| SS09561 | 0 | 8.29E-05 | 2.55E-05 |
| SS05029 | 0 | 0.000151445 | 2.61E-05 |
| SS02032 | 0 | 5.49E-05 | 2.63E-05 |
| SS10342 | 0 | 5.49E-05 | 2.63E-05 |
| SS14142 | 0 | 6.10E-05 | 2.64E-05 |
| SS08972 | 0 | 3.40E-05 | 2.65E-05 |
| SS12154 | 0.01405056 | 0.002060203 | 2.69E-05 |
| SS13636 | 0.01405056 | 0.002060203 | 2.69E-05 |
| SS05748 | 0.000778538 | 0.000303268 | 2.74E-05 |
| SS11742 | 8.16E-05 | 3.03E-05 | 2.75E-05 |
| SS11306 | 0 | 9.24E-05 | 2.77E-05 |
| SS13232 | 2.18E-05 | 7.18E-06 | 2.82E-05 |
| SS06960 | 0.010704209 | 0.012857109 | 2.82E-05 |
| SS04187 | 0 | 9.69E-05 | 2.85E-05 |
| SS15329 | 0.001212597 | 0.00190242 | 2.90E-05 |
| SS02876 | 0 | 1.75E-05 | 2.91E-05 |
| SS13060 | 0 | 1.75E-05 | 2.91E-05 |
| SS05533 | 0.000674861 | 0.001436133 | 2.93E-05 |
| SS05545 | 0.000674861 | 0.001436133 | 2.93E-05 |
| SS11471 | 5.72E-05 | 1.12E-05 | 2.94E-05 |
| SS08243 | 8.13E-06 | 7.01E-05 | 2.94E-05 |
| SS13869 | 8.13E-06 | 7.01E-05 | 2.94E-05 |
| SS05847 | 0.011538388 | 0.000589267 | 2.98E-05 |
| SS05060 | 0.006473716 | 0.003660878 | 3.00E-05 |
| SS14383 | 7.36E-05 | 1.71E-05 | 3.02E-05 |
| SS12513 | 0.002148974 | 9.04E-06 | 3.02E-05 |
| SS05763 | 0 | 0.000114328 | 3.05E-05 |
| SS13891 | 0.000677785 | 0.000282142 | 3.09E-05 |
| SS10985 | 3.81E-05 | 1.08E-05 | 3.12E-05 |
| SS08546 | 0 | 9.78E-05 | 3.13E-05 |
| SS02577 | 0.010351117 | 0.008493324 | 3.13E-05 |
| SS08407 | 0.010351117 | 0.008493324 | 3.13E-05 |
| SS08786 | 0.010351117 | 0.008493324 | 3.13E-05 |
| SS05892 | 0.000968344 | 0.00217059 | 3.14E-05 |
| SS15411 | 0.000968344 | 0.00217059 | 3.14E-05 |
| SS01715 | 0.022655117 | 0.006679704 | 3.16E-05 |

| | | | |
|---|---|---|---|
| SS13250 | 0.010167792 | 0.00127436 | 3.19E-05 |
| SS07624 | 0.006463669 | 0.000162146 | 3.20E-05 |
| SS02017 | 0 | 0.00015537 | 3.20E-05 |
| SS02908 | 0 | 0.00015537 | 3.20E-05 |
| SS06757 | 0.032669394 | 0.029078851 | 3.21E-05 |
| SS09070 | 0.032669394 | 0.029078851 | 3.21E-05 |
| SS10185 | 0.032669394 | 0.029078851 | 3.21E-05 |
| SS12032 | 0.032669394 | 0.029078851 | 3.21E-05 |
| SS14856 | 0.032669394 | 0.029078851 | 3.21E-05 |
| SS12231 | 0.000373005 | 2.39E-05 | 3.23E-05 |
| SS05353 | 0 | 0.000222846 | 3.26E-05 |
| SS00363 | 0.020175978 | 0.005678623 | 3.27E-05 |
| SS06866 | 0.020175978 | 0.005678623 | 3.27E-05 |
| SS10293 | 0.020175978 | 0.005678623 | 3.27E-05 |
| SS14206 | 0.020175978 | 0.005678623 | 3.27E-05 |
| SS01074 | 0.013435951 | 0.001502157 | 3.27E-05 |
| SS11972 | 0 | 2.87E-05 | 3.28E-05 |
| SS11883 | 0.01635532 | 0.013401187 | 3.28E-05 |
| SS12985 | 0 | 4.20E-05 | 3.33E-05 |
| SS15268 | 0 | 4.20E-05 | 3.33E-05 |
| SS14588 | 0.000419275 | 5.16E-05 | 3.34E-05 |
| SS14682 | 0.000419275 | 5.16E-05 | 3.34E-05 |
| SS13951 | 0.005530859 | 0.002201003 | 3.34E-05 |
| SS09311 | 6.27E-05 | 1.22E-05 | 3.36E-05 |
| SS02620 | 0.000653308 | 0.000289759 | 3.38E-05 |
| SS03075 | 0.000653308 | 0.000289759 | 3.38E-05 |
| SS08688 | 0.000653308 | 0.000289759 | 3.38E-05 |
| SS04798 | 0.002443902 | 0.000600906 | 3.41E-05 |
| SS03790 | 0 | 4.22E-05 | 3.43E-05 |
| SS03862 | 0 | 4.22E-05 | 3.43E-05 |
| SS11461 | 0 | 4.85E-05 | 3.49E-05 |
| SS02546 | 0.025192568 | 0.032861398 | 3.50E-05 |
| SS07588 | 0.025192568 | 0.032861398 | 3.50E-05 |
| SS05382 | 0.006395238 | 0.002598175 | 3.50E-05 |
| SS05290 | 0.00037601 | 0.002652673 | 3.53E-05 |
| SS05296 | 0.00037601 | 0.002652673 | 3.53E-05 |
| SS05297 | 0.00037601 | 0.002652673 | 3.53E-05 |
| SS05302 | 0.00037601 | 0.002652673 | 3.53E-05 |
| SS10107 | 0.000236798 | 9.95E-05 | 3.54E-05 |
| SS12431 | 0.004687297 | 6.24E-06 | 3.56E-05 |
| SS13826 | 0.002997376 | 0.001022843 | 3.57E-05 |
| SS14738 | 0 | 2.59E-05 | 3.61E-05 |
| SS02265 | 0.010312692 | 0.012501094 | 3.63E-05 |
| SS02569 | 0.010312692 | 0.012501094 | 3.63E-05 |
| SS04656 | 0.010312692 | 0.012501094 | 3.63E-05 |
| SS05299 | 0.00034332 | 0.00261829 | 3.67E-05 |
| SS07614 | 0.001223012 | 0.001620551 | 3.74E-05 |

| | | | |
|---|---|---|---|
| SS01862 | 0 | 3.34E-05 | 3.74E-05 |
| SS06316 | 0 | 3.34E-05 | 3.74E-05 |
| SS06416 | 0 | 3.34E-05 | 3.74E-05 |
| SS07158 | 0 | 1.93E-05 | 3.80E-05 |
| SS11193 | 0 | 6.50E-05 | 3.83E-05 |
| SS00577 | 0 | 9.49E-05 | 3.83E-05 |
| SS03364 | 0 | 9.19E-05 | 3.83E-05 |
| SS13897 | 0 | 9.19E-05 | 3.83E-05 |
| SS09350 | 0 | 3.52E-05 | 3.85E-05 |
| SS15013 | 0.002916214 | 0.003745785 | 3.85E-05 |
| SS08555 | 2.76E-06 | 6.98E-05 | 3.86E-05 |
| SS02882 | 0.000150911 | 0.001073797 | 3.92E-05 |
| SS00341 | 0 | 1.56E-05 | 3.96E-05 |
| SS12051 | 0.000525796 | 9.49E-06 | 3.97E-05 |
| SS12052 | 0.000525796 | 9.49E-06 | 3.97E-05 |
| SS08453 | 0 | 0.000134179 | 4.08E-05 |
| SS01459 | 0 | 4.03E-05 | 4.13E-05 |
| SS13058 | 0 | 4.03E-05 | 4.13E-05 |
| SS07820 | 0.00018516 | 0.000358085 | 4.14E-05 |
| SS01035 | 0.011694793 | 0.003058967 | 4.14E-05 |
| SS04919 | 0.011694793 | 0.003058967 | 4.14E-05 |
| SS09110 | 0.011694793 | 0.003058967 | 4.14E-05 |
| SS04669 | 0.01237173 | 0.011061434 | 4.15E-05 |
| SS11805 | 0.01237173 | 0.011061434 | 4.15E-05 |
| SS09788 | 2.18E-05 | 0.000165902 | 4.18E-05 |
| SS15181 | 2.18E-05 | 0.000165902 | 4.18E-05 |
| SS00971 | 0 | 4.66E-05 | 4.21E-05 |
| SS13438 | 0 | 4.34E-05 | 4.26E-05 |
| SS01753 | 0.012601084 | 0.001434219 | 4.29E-05 |
| SS02915 | 0 | 0.000563427 | 4.30E-05 |
| SS03778 | 0 | 9.21E-05 | 4.30E-05 |
| SS11245 | 0.00668637 | 0.003505137 | 4.33E-05 |
| SS06051 | 0.055744473 | 0.044524169 | 4.35E-05 |
| SS06075 | 0.055744473 | 0.044524169 | 4.35E-05 |
| SS09646 | 0.055744473 | 0.044524169 | 4.35E-05 |
| SS13240 | 0 | 2.02E-05 | 4.44E-05 |
| SS07111 | 0.000653951 | 0.000102675 | 4.46E-05 |
| SS02716 | 0 | 0.000808614 | 4.47E-05 |
| SS13592 | 0 | 0.000102139 | 4.47E-05 |
| SS08805 | 0.000553439 | 0.000113846 | 4.49E-05 |
| SS00755 | 0.009341791 | 0.001479199 | 4.49E-05 |
| SS01535 | 0.000588492 | 0.000294114 | 4.50E-05 |
| SS06363 | 0.000588492 | 0.000294114 | 4.50E-05 |
| SS08973 | 0.000588492 | 0.000294114 | 4.50E-05 |
| SS05267 | 0.010903317 | 0.005797973 | 4.57E-05 |
| SS09451 | 0.004669829 | 0.00064572 | 4.57E-05 |
| SS02145 | 0 | 4.92E-05 | 4.62E-05 |

| | | | |
|---|---|---|---|
| SS03119 | 0.002456921 | 0.007835732 | 4.66E-05 |
| SS10210 | 0.002456921 | 0.007835732 | 4.66E-05 |
| SS02202 | 0.004822861 | 0.000135424 | 4.68E-05 |
| SS03001 | 0.000187844 | 0.000109324 | 4.68E-05 |
| SS15538 | 0.000343561 | 0.000747949 | 4.70E-05 |
| SS09373 | 0 | 2.57E-05 | 4.70E-05 |
| SS13444 | 0.001972267 | 0.000175092 | 4.72E-05 |
| SS03057 | 1.09E-05 | 0.000164032 | 4.72E-05 |
| SS05495 | 1.09E-05 | 0.000164032 | 4.72E-05 |
| SS09516 | 1.09E-05 | 0.000164032 | 4.72E-05 |
| SS10481 | 0.019844415 | 0.000949374 | 4.73E-05 |
| SS00359 | 0.008795601 | 0.000650016 | 4.75E-05 |
| SS10737 | 0.008795601 | 0.000650016 | 4.75E-05 |
| SS10925 | 0 | 3.85E-05 | 4.78E-05 |
| SS02557 | 0.003721959 | 0.005444689 | 4.79E-05 |
| SS04363 | 0.010247795 | 0.000339212 | 4.80E-05 |
| SS04854 | 0 | 3.66E-05 | 4.80E-05 |
| SS06897 | 0.020679901 | 0.006188533 | 4.81E-05 |
| SS08791 | 0.005516636 | 0.000943782 | 4.81E-05 |
| SS05194 | 0.008955722 | 0.007649471 | 4.88E-05 |
| SS06498 | 0.008955722 | 0.007649471 | 4.88E-05 |
| SS01576 | 0.007099956 | 0.004598505 | 4.91E-05 |
| SS09137 | 0.007099956 | 0.004598505 | 4.91E-05 |
| SS15196 | 0.002155225 | 0.001458504 | 4.92E-05 |
| SS15340 | 0.017478314 | 0.013588562 | 4.96E-05 |
| SS15162 | 0 | 5.31E-05 | 4.96E-05 |
| SS08296 | 0.012017766 | 0.00791587 | 5.02E-05 |
| SS13640 | 0.012017766 | 0.00791587 | 5.02E-05 |
| SS03903 | 0 | 5.51E-05 | 5.04E-05 |
| SS05750 | 0.001686595 | 0.001121754 | 5.04E-05 |
| SS01926 | 8.99E-05 | 3.71E-06 | 5.05E-05 |
| SS04299 | 0.00081952 | 0.001338484 | 5.08E-05 |
| SS06102 | 0.000212723 | 0.000438775 | 5.14E-05 |
| SS00067 | 0.008140491 | 0.003672702 | 5.14E-05 |
| SS05185 | 0.008140491 | 0.003672702 | 5.14E-05 |
| SS05500 | 0.008140491 | 0.003672702 | 5.14E-05 |
| SS08865 | 0.008140491 | 0.003672702 | 5.14E-05 |
| SS11904 | 0.008140491 | 0.003672702 | 5.14E-05 |
| SS04857 | 0 | 3.15E-05 | 5.17E-05 |
| SS08690 | 0 | 0.000297515 | 5.18E-05 |
| SS05779 | 0.006029013 | 0.007610065 | 5.19E-05 |
| SS11815 | 0.006029013 | 0.007610065 | 5.19E-05 |
| SS09841 | 0 | 2.80E-05 | 5.20E-05 |
| SS11976 | 2.76E-06 | 3.91E-05 | 5.23E-05 |
| SS01572 | 0.005707164 | 0.007476678 | 5.23E-05 |
| SS02184 | 0.005707164 | 0.007476678 | 5.23E-05 |
| SS09136 | 0.005707164 | 0.007476678 | 5.23E-05 |

| | | | |
|---|---|---|---|
| SS13165 | 0.005707164 | 0.007476678 | 5.23E-05 |
| SS00045 | 0 | 5.53E-05 | 5.24E-05 |
| SS06126 | 0 | 5.53E-05 | 5.24E-05 |
| SS10399 | 0.000234355 | 1.38E-05 | 5.24E-05 |
| SS11369 | 4.36E-05 | 1.05E-06 | 5.25E-05 |
| SS07743 | 0.016551904 | 0.000236362 | 5.34E-05 |
| SS00293 | 0.000808945 | 5.32E-06 | 5.36E-05 |
| SS02070 | 0.008446385 | 0.006914045 | 5.37E-05 |
| SS02604 | 0.008446385 | 0.006914045 | 5.37E-05 |
| SS08818 | 0.008446385 | 0.006914045 | 5.37E-05 |
| SS13795 | 0.008446385 | 0.006914045 | 5.37E-05 |
| SS02804 | 0 | 9.35E-05 | 5.43E-05 |
| SS11605 | 0 | 7.42E-05 | 5.59E-05 |
| SS07786 | 0 | 2.69E-05 | 5.60E-05 |
| SS13078 | 0 | 2.69E-05 | 5.60E-05 |
| SS13955 | 0.03171065 | 0.001271561 | 5.62E-05 |
| SS00759 | 0.00457925 | 0.000490603 | 5.66E-05 |
| SS05751 | 0.002232177 | 0.001016817 | 5.66E-05 |
| SS01289 | 0 | 5.00E-05 | 5.67E-05 |
| SS09702 | 0 | 5.00E-05 | 5.67E-05 |
| SS15530 | 0.003612593 | 0.002323179 | 5.68E-05 |
| SS00116 | 0.001653665 | 0.00010567 | 5.69E-05 |
| SS01086 | 0 | 4.93E-05 | 5.69E-05 |
| SS13496 | 0 | 2.18E-05 | 5.77E-05 |
| SS14590 | 0.000433417 | 0.000151086 | 5.79E-05 |
| SS14684 | 0.000433417 | 0.000151086 | 5.79E-05 |
| SS03368 | 2.99E-05 | 0.000167363 | 5.80E-05 |
| SS09058 | 0.032669394 | 0.029319109 | 5.81E-05 |
| SS10176 | 0.032669394 | 0.029319109 | 5.81E-05 |
| SS12029 | 0.032669394 | 0.029319109 | 5.81E-05 |
| SS14853 | 0.032669394 | 0.029319109 | 5.81E-05 |
| SS06878 | 0.012023202 | 0.010080302 | 5.84E-05 |
| SS15386 | 0.012023202 | 0.010080302 | 5.84E-05 |
| SS15432 | 0.012023202 | 0.010080302 | 5.84E-05 |
| SS05940 | 0.002701048 | 0.00032379 | 5.94E-05 |
| SS00033 | 0 | 3.32E-05 | 5.96E-05 |
| SS01288 | 0 | 5.63E-05 | 5.96E-05 |
| SS09701 | 0 | 5.63E-05 | 5.96E-05 |
| SS00617 | 0 | 2.77E-05 | 6.09E-05 |
| SS04162 | 0.008347192 | 0.0010993 | 6.10E-05 |
| SS13711 | 0 | 5.87E-05 | 6.11E-05 |
| SS10866 | 0.008195135 | 0.005568687 | 6.15E-05 |
| SS01638 | 0 | 8.27E-05 | 6.18E-05 |
| SS01654 | 0 | 8.27E-05 | 6.18E-05 |
| SS10377 | 0 | 9.93E-05 | 6.20E-05 |
| SS07901 | 0.016770007 | 0.011318184 | 6.21E-05 |
| SS04331 | 0.013028365 | 0.000461285 | 6.25E-05 |

| | | | |
|---|---|---|---|
| SS15469 | 0.003522737 | 0.000589789 | 6.29E-05 |
| SS02926 | 0 | 0.000349694 | 6.29E-05 |
| SS01037 | 0 | 6.67E-05 | 6.36E-05 |
| SS12077 | 0 | 1.60E-05 | 6.36E-05 |
| SS13178 | 0 | 2.62E-05 | 6.38E-05 |
| SS07305 | 0.013019189 | 0.010750216 | 6.39E-05 |
| SS10572 | 0.013019189 | 0.010750216 | 6.39E-05 |
| SS00347 | 0 | 8.84E-05 | 6.40E-05 |
| SS14589 | 0.003021371 | 0.000729171 | 6.44E-05 |
| SS14683 | 0.003021371 | 0.000729171 | 6.44E-05 |
| SS00540 | 0 | 7.88E-05 | 6.46E-05 |
| SS10305 | 0.018454101 | 0.015609319 | 6.46E-05 |
| SS02952 | 0.000261516 | 9.19E-06 | 6.48E-05 |
| SS05502 | 0 | 1.78E-05 | 6.51E-05 |
| SS05235 | 0.005026294 | 0.000388213 | 6.52E-05 |
| SS14820 | 0.005026294 | 0.000388213 | 6.52E-05 |
| SS04722 | 0.002978267 | 0.001016958 | 6.67E-05 |
| SS13800 | 0.002978267 | 0.001016958 | 6.67E-05 |
| SS13827 | 0.002978267 | 0.001016958 | 6.67E-05 |
| SS00360 | 0.010450378 | 0.001628019 | 6.77E-05 |
| SS10738 | 0.010450378 | 0.001628019 | 6.77E-05 |
| SS02692 | 0.009436923 | 0.0069011 | 6.78E-05 |
| SS11758 | 0.009436923 | 0.0069011 | 6.78E-05 |
| SS14125 | 0.009436923 | 0.0069011 | 6.78E-05 |
| SS01140 | 0.003036798 | 2.47E-06 | 6.78E-05 |
| SS03195 | 0.010911725 | 0.000405682 | 6.79E-05 |
| SS03202 | 0.010911725 | 0.000405682 | 6.79E-05 |
| SS01562 | 0.004881989 | 0.003852781 | 6.83E-05 |
| SS07315 | 1.63E-05 | 0.00010891 | 6.90E-05 |
| SS02423 | 0.023993161 | 0.020521331 | 6.93E-05 |
| SS03462 | 0.023993161 | 0.020521331 | 6.93E-05 |
| SS05600 | 0 | 7.26E-05 | 6.94E-05 |
| SS07771 | 0 | 0.000161267 | 7.02E-05 |
| SS11426 | 0.007399804 | 0.013815474 | 7.07E-05 |
| SS07485 | 5.45E-06 | 6.54E-05 | 7.13E-05 |
| SS02026 | 0.000606958 | 0.000138937 | 7.13E-05 |
| SS09184 | 0.000144419 | 1.37E-05 | 7.25E-05 |
| SS10066 | 0.058013147 | 0.003076632 | 7.27E-05 |
| SS01055 | 0.007590849 | 0.005846652 | 7.27E-05 |
| SS01315 | 0.007590849 | 0.005846652 | 7.27E-05 |
| SS03950 | 0.007590849 | 0.005846652 | 7.27E-05 |
| SS11243 | 0.007590849 | 0.005846652 | 7.27E-05 |
| SS15482 | 0.007590849 | 0.005846652 | 7.27E-05 |
| SS10879 | 0 | 0.000204483 | 7.32E-05 |
| SS09821 | 3.29E-05 | 0.000256247 | 7.39E-05 |
| SS04527 | 0.019197117 | 0.015977151 | 7.41E-05 |
| SS06289 | 0 | 5.74E-05 | 7.41E-05 |

| | | | |
|---|---|---|---|
| SS14845 | 0.000127673 | 0.000373684 | 7.44E-05 |
| SS15137 | 0 | 0.000137993 | 7.50E-05 |
| SS05080 | 0.002302408 | 0.000716116 | 7.53E-05 |
| SS11070 | 0 | 4.33E-05 | 7.62E-05 |
| SS04635 | 0 | 2.46E-05 | 7.65E-05 |
| SS08900 | 0.005887232 | 0.000178423 | 7.69E-05 |
| SS06680 | 0 | 6.59E-05 | 7.71E-05 |
| SS08294 | 3.02E-05 | 0.000296222 | 7.72E-05 |
| SS12196 | 0 | 6.53E-05 | 7.73E-05 |
| SS13893 | 0.000795686 | 0.000276585 | 7.77E-05 |
| SS14096 | 0 | 6.10E-05 | 7.77E-05 |
| SS10989 | 0 | 6.92E-05 | 7.79E-05 |
| SS09429 | 0 | 4.19E-05 | 7.79E-05 |
| SS06273 | 0.034727296 | 0.038078116 | 7.85E-05 |
| SS14037 | 0.034727296 | 0.038078116 | 7.85E-05 |
| SS04402 | 0.009103272 | 0.003349066 | 7.89E-05 |
| SS04301 | 2.45E-05 | 0.000330736 | 7.89E-05 |
| SS04700 | 0.003107144 | 0.000117741 | 7.95E-05 |
| SS09174 | 0.003107144 | 0.000117741 | 7.95E-05 |
| SS06800 | 0.017360769 | 0.000263973 | 8.00E-05 |
| SS04466 | 0.005664416 | 7.53E-05 | 8.02E-05 |
| SS08358 | 2.72E-05 | 0.001314302 | 8.09E-05 |
| SS01748 | 0.010385252 | 0.007374956 | 8.17E-05 |
| SS14009 | 0 | 2.02E-05 | 8.18E-05 |
| SS01867 | 0 | 5.08E-05 | 8.23E-05 |
| SS07691 | 0.00844307 | 0.001409109 | 8.30E-05 |
| SS07072 | 0 | 2.38E-05 | 8.36E-05 |
| SS06769 | 0.009460609 | 0.007917274 | 8.39E-05 |
| SS06805 | 0.009460609 | 0.007917274 | 8.39E-05 |
| SS13840 | 0.009460609 | 0.007917274 | 8.39E-05 |
| SS10265 | 0.00010612 | 3.52E-05 | 8.39E-05 |
| SS04544 | 0.014546592 | 0.017065214 | 8.46E-05 |
| SS04574 | 0.014546592 | 0.017065214 | 8.46E-05 |
| SS10636 | 0.014546592 | 0.017065214 | 8.46E-05 |
| SS09531 | 0.000602313 | 0.000104422 | 8.50E-05 |
| SS13758 | 0.000602313 | 0.000104422 | 8.50E-05 |
| SS15103 | 0 | 4.78E-05 | 8.52E-05 |
| SS06246 | 0.009666759 | 0.008300295 | 8.56E-05 |
| SS10298 | 0.009666759 | 0.008300295 | 8.56E-05 |
| SS01433 | 0 | 3.19E-05 | 8.57E-05 |
| SS12950 | 0 | 3.19E-05 | 8.57E-05 |
| SS00331 | 0 | 2.72E-05 | 8.58E-05 |
| SS08347 | 0 | 0.000672777 | 8.64E-05 |
| SS00335 | 0 | 2.33E-05 | 8.79E-05 |
| SS04253 | 0.004696072 | 0.000123344 | 8.84E-05 |
| SS00098 | 0.009597377 | 0.001706167 | 8.84E-05 |
| SS06644 | 0 | 2.79E-05 | 8.85E-05 |

| | | | |
|---|---|---|---|
| SS03157 | 0 | 0.000399467 | 8.86E-05 |
| SS08954 | 0 | 6.22E-05 | 8.88E-05 |
| SS11797 | 0 | 6.22E-05 | 8.88E-05 |
| SS04010 | 0.000253063 | 0.000670828 | 8.92E-05 |
| SS07536 | 0.003518091 | 0.001903021 | 8.98E-05 |
| SS04305 | 5.72E-05 | 2.68E-05 | 9.00E-05 |
| SS08536 | 0.001776291 | 0.003018518 | 9.01E-05 |
| SS08795 | 0.012576171 | 0.006620106 | 9.03E-05 |
| SS13044 | 0 | 2.42E-05 | 9.16E-05 |
| SS05490 | 0.021960688 | 0.001151119 | 9.24E-05 |
| SS04924 | 0 | 3.68E-05 | 9.24E-05 |
| SS06481 | 0.023923263 | 0.021379507 | 9.24E-05 |
| SS12303 | 0.007324412 | 0.000268827 | 9.28E-05 |
| SS05974 | 0.015317044 | 0.019159024 | 9.36E-05 |
| SS06124 | 0.015317044 | 0.019159024 | 9.36E-05 |
| SS09655 | 0.015317044 | 0.019159024 | 9.36E-05 |
| SS08723 | 0.028446413 | 0.017494293 | 9.43E-05 |
| SS06084 | 0 | 5.52E-05 | 9.45E-05 |
| SS04126 | 0.018755281 | 0.021837013 | 9.51E-05 |
| SS06696 | 0.018755281 | 0.021837013 | 9.51E-05 |
| SS14978 | 0.018755281 | 0.021837013 | 9.51E-05 |
| SS01084 | 0 | 3.72E-05 | 9.57E-05 |
| SS09022 | 0 | 2.76E-05 | 9.57E-05 |
| SS05701 | 0.006497643 | 0.001171662 | 9.58E-05 |
| SS09381 | 0.006497643 | 0.001171662 | 9.58E-05 |
| SS02206 | 0.005159414 | 0.000593 | 9.61E-05 |
| SS11190 | 0.005159414 | 0.000593 | 9.61E-05 |
| SS00837 | 0 | 0.000156452 | 9.80E-05 |
| SS11088 | 0.051514014 | 0.05751498 | 9.84E-05 |
| SS11175 | 0.010661621 | 0.012794216 | 9.87E-05 |
| SS11184 | 0.010661621 | 0.012794216 | 9.87E-05 |
| SS00248 | 0.001127306 | 5.35E-05 | 9.89E-05 |
| SS10900 | 0.001127306 | 5.35E-05 | 9.89E-05 |
| SS11138 | 0.001127306 | 5.35E-05 | 9.89E-05 |
| SS13159 | 0 | 8.92E-05 | 9.96E-05 |
