## Supplementary figures and images for "Exploring metal resistance genes and mechanisms in copper enriched metal ore metagenome"

### Figure S-level2.pdf

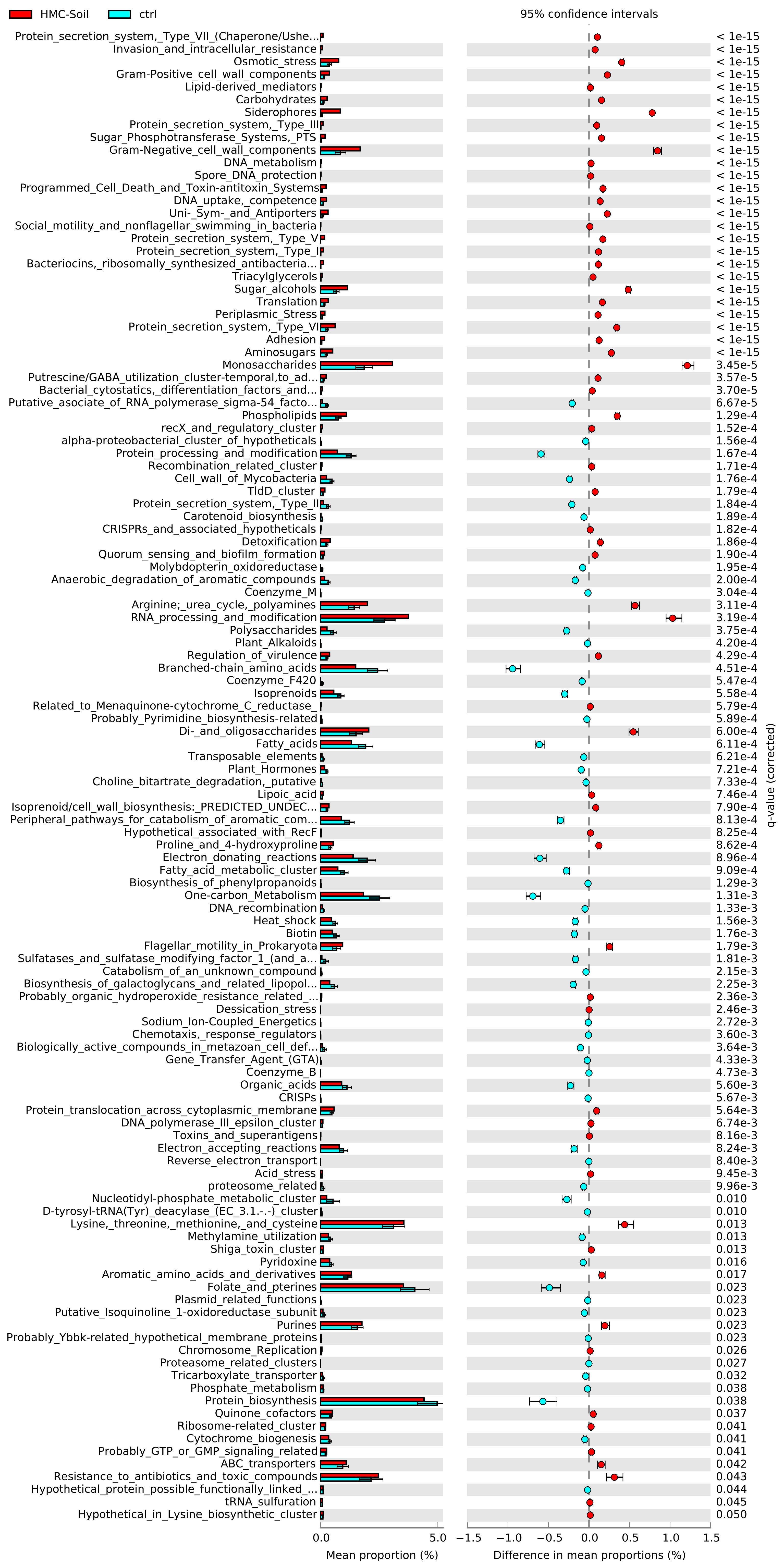

### Figure S-level3.pdf

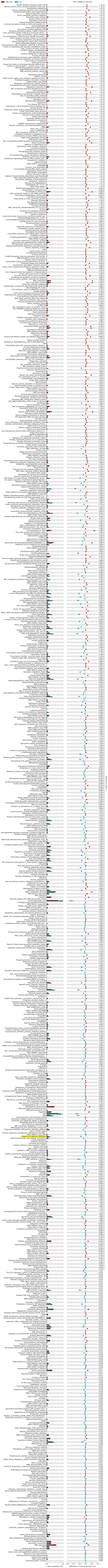
